# Toward Sustainable Biocultural Tourism: An Integrated Spatial Analysis of Cultural and Biodiversity Richness in Colombia

**DOI:** 10.1101/2024.02.01.578429

**Authors:** Alejandra Echeverri, Natasha M. Batista, Stacie Wolny, Guido A. Herrera-R, Federico Andrade-Rivas, Allison Bailey, Anaid Cardenas-Navarrete, Armando Dávila Arenas, Andres Felipe Díaz-Salazar, Katherine Victoria Hernandez, Kelley E. Langhans, Bryam E. Mateus-Aguilar, Dallas Levey, Andrew Neill, Oliver Nguyen, Andres Felipe Suárez-Castro, Felipe Zapata, Natalia Ocampo-Peñuela

**Affiliations:** Department of Environmental Science, Policy & Management, University of California Berkeley, Berkeley, CA, USA; The Natural Capital Project, Stanford University, Stanford, CA, USA; Department of Ecology and Evolutionary Biology, The University of Knoxville, TN, USA; School of Public Health and Social Policy, University of Victoria, Victoria, BC, Canada; Instituto de Salud y Ambiente, Universidad El Bosque, Bogotá, Colombia; Washington State Department of Natural Resources, WA, USA; Department of Integrative Biology, University of California Berkeley, Berkeley, CA, USA; Independent musician and tourist operator. Bogota, Colombia; Departamento de Ciencias Biológicas, Universidad de los Andes, Bogotá, Colombia; Institute of the Environment and Sustainability, University of California Los Angeles, Los Angeles, CA, USA; Department of Fish and Wildlife Conservation, Virginia Tech, VG, USA; Department of Biology, Stanford University, Stanford, CA, USA; Botany Department, School of Natural Sciences, Trinity College Dublin, Dublin, Ireland; Australian Rivers Institute, Griffith University, Australia; Department of Ecology and Evolutionary Biology and Center for Tropical Research, Institute of the Environment and Sustainability, University of California, Los Angeles, CA, USA; Environmental Studies Department, University of California Santa Cruz, Santa Cruz, CA, USA

**Keywords:** Biocultural diversity, Birdwatching tourism, Conservation planning, Cultural heritage, Ecotourism, Recreation services, Sports fishing, Sustainable tourism

## Abstract

1. Tourism plays a vital role in both economic development and depending on the scale, it can also aid environmental conservation. Tourism planning often considers culture-based and nature-based tourism separately, failing to recognize the synergies between them, with the potential to market locations as biocultural destinations.
2. Using Colombia as a case study, we created metrics of taxonomic biological diversity as measured by vertebrate species richness (including birds, mammals, freshwater fishes, reptiles, and amphibians) and institutionalized cultural richness (by counting the number of UNESCO world heritage sites, intangible cultural heritage sites, museums, endemic music festivals, Afro-Colombian territories, and Indigenous reserves), and evaluated the spatial correlations between them.
3. To determine biocultural tourism potential, we evaluated whether biocultural richness was accessible, and mapped potential biocultural tourism supply. By mapping areas of sports fisheries, birdwatching destinations, and airport arrivals we also estimated spatial demand. We also analyzed the difference between demand and supply to assess the realized and untapped potential for biocultural destinations.
4. While biocultural richness is high in the Amazon, Pacific, and Caribbean regions, we found that there are no win-win-win locations where culture, species richness, and accessibility are all high. Areas with great potential for biocultural tourism development largely coincide with designated Indigenous reserves and Afro-Colombian territories.
5. This study underscores the power of integrating cultural and biological variables to reshape the tourism sector. Our paper offers practical recommendations for policymakers, conservation organizations, and local communities seeking to create transformative and inclusive tourism experiences.

## 1 Introduction

Tourism can in theory generate win-win-win approaches to biodiversity conservation, cultural heritage protection, and economic development given its potential to be non-extractive (Echeverri et al., 2022). Tourism accounts for ∼10% of global Gross Domestic Product (GDP), with ecotourism being its fastest growing sector, growing at a rate of 15.2% from 2022 to 2030 (Balmford et al., 2009; OECD, 2020). A territory’s cultural and natural heritage determines its level of attractiveness to tourists (Canale et al., 2019; Cuccia et al., 2017), and places that offer unique experiences become desired destinations. Tourists are often drawn to sites with rich biodiversity and ecosystems (e.g., the Great Barrier Reef), or to places that offer unique insights on human cultures, including UNESCO World Heritage Sites (e.g., Egyptian pyramids) (Buckley, 2011).

According to the push-pull theory (Dann, 1976), a place is perceived as attractive if it has “pull factors” that determine the level of uniqueness of a destination. Pull factors influence when, where, and how people travel to that destination (Klenosky, 2002). The tourism industry often segments tourist markets based on types of pull factors (Prayag, 2010). For example, places are branded as creative destinations (e.g., New York City’s Broadway shows), cultural destinations (e.g., Machu Picchu in Peru), or nature destinations (e.g., Serengeti plains in Africa) (Al-Ababneh, 2019; D’Auria, 2009; Echeverri et al., 2022).

Conversely, “push factors” are specific forces that influence a person’s decision on where *not* to go on vacation (Kim et al., 2003). People might not travel to a location if it has, or they perceive it to have, high violence and crime rates, unsurmountable poverty, disease risk, or lacks accessibility and infrastructure (Echeverri et al., 2022). The interaction between push and pull factors is well documented for national park tourism patterns (Kim et al., 2003; Klenosky, 2002). However, these studies tend to focus narrowly on single types of tourism (i.e., nature-based), omitting synergies and antagonisms between multiple forms of tourism experiences that could comprehensively indicate how push and pull factors interact to make a destination desirable.

Cultural manifestations, including art, music, traditional dances, and artisan crafts, are often inextricably linked with local biodiversity, and both can act as pull factors driving tourism patterns (Aguado et al., 2021; Berkes and Berkes, 2009; Satterfield et al., 2013). For instance, fish and bird migrations are celebrated with folkloric music festivals among the Tangkhul Naga in Northeast India (Varah and Varah, 2022); and national parks such as the Great Barrier Reef often intersect with areas of endangered Indigenous languages (Romaine and Gorenflo, 2017). Cultural practices, such as Indigenous fire stewardship in North America, Australia, and South America, or traditional whaling in the circumpolar region—significantly impact ecological community composition (Firestone and Lilley, 2004; Hidasi-Neto et al., 2012). This occurs because the cultivation of culturally revered species results in a hyperdominance of certain species, thereby altering ecological networks (ter Steege et al., 2013). While research has documented how biodiversity interacts with infrastructure and crime to pull or push people to and from destinations (Ocampo-Peñuela and Winton, 2017), it has overlooked the potential overlap between cultural and biological pull factors, and their interaction with push factors in driving tourism patterns.

A useful framing for tourism planning could be one that focuses on biocultural diversity, which refers to the interdependence between biodiversity, cultural diversity, and linguistic diversity in the web of life (Maffi, 2010). In the context of tourism, a biocultural tourism framing posits that biodiversity and cultural diversity act as pull factors drawing people to destinations. For example, archaeological sites might coincide with areas for birdwatching (Figure 1). Of course, they can also interact as push factors, if certain species and cultural practices are perceived as dangerous, unattractive, supremacist or discriminatory. Yet, the interaction between cultural and biological diversity has mostly been overlooked. A few studies have shown the overlap between linguistic diversity and biodiversity (Gorenflo et al., 2012; Loh and Harmon, 2005), but this research has not examined the potential of these as interacting to form tourist destinations.

**Figure 1.**
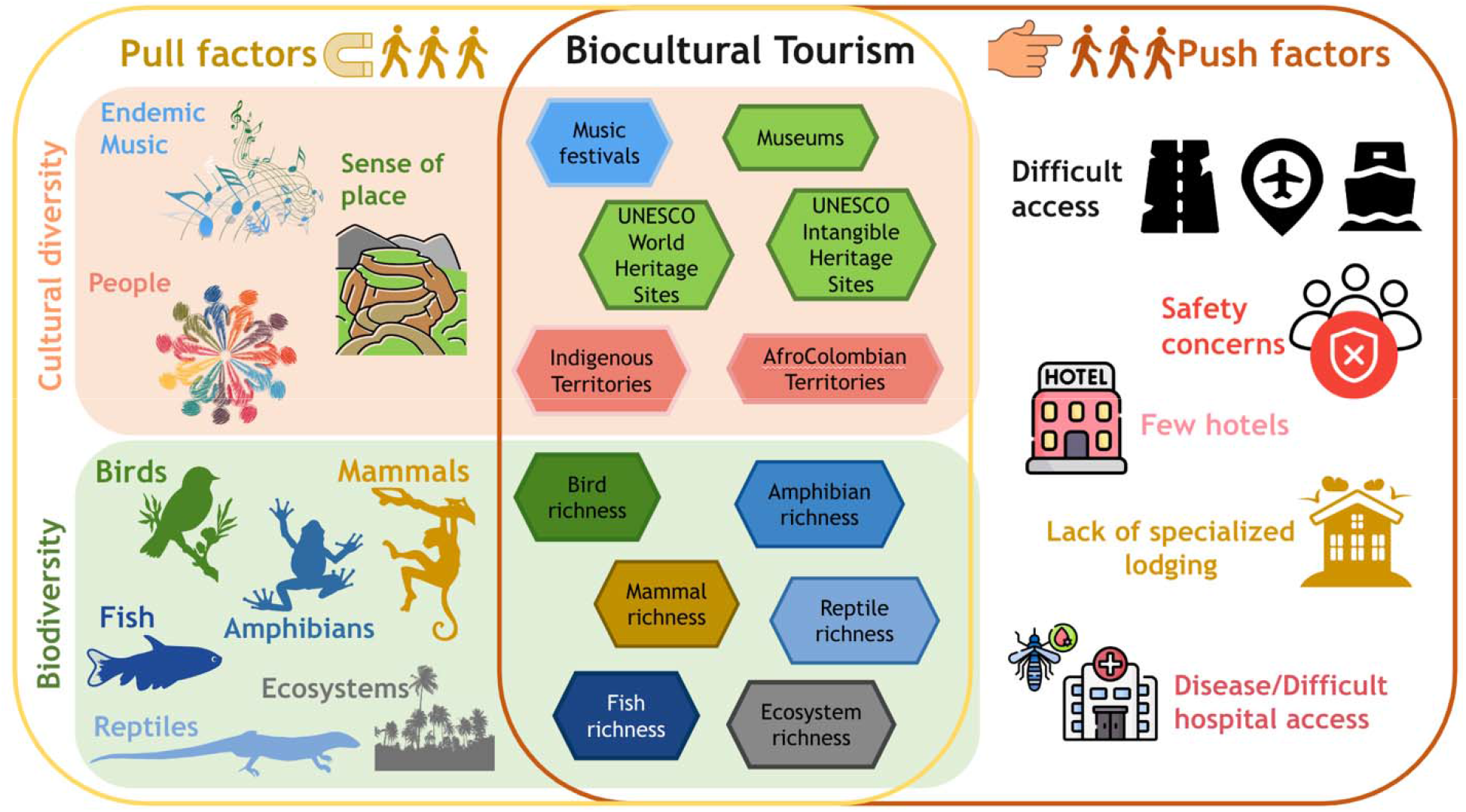
Biocultural tourism conceptual figure explaining the push-pull factors that may draw people to a destination or prevent them from going. Here, we propose that a biocultural framing recognizes the intricacies between biodiversity and cultural diversity and carefully balances them with accessibility. Within the “Biocultural tourism” panel, we show the variables used in the study.

**Figure 2.**
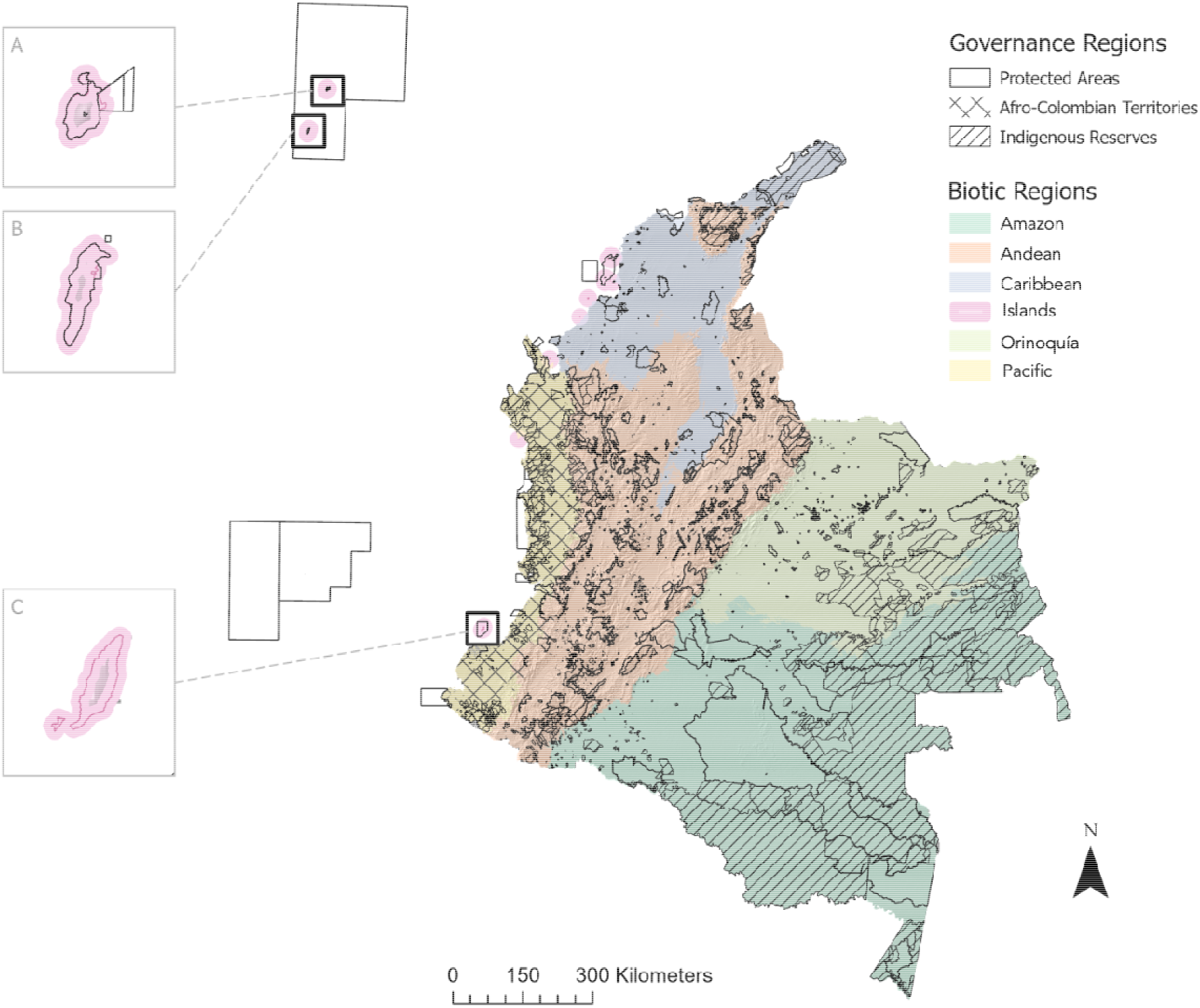
Biotic regions of Colombia overlaid with national protected areas, Afro-Colombian territories, and Indigenous reserves. Providencia (A), San Andrés (B), and Gorgona (C) islands are displayed in the insets. Malpelo Island is within a south-western marine protected area. Governance regions are displayed with textures, while biotic regions in colors. Islands were excluded from our analyses, and only continental Colombia was used

As countries develop, both their biological diversity and cultural diversity can become threatened. Recently, we have experienced unparalleled losses in both, arising from a common set of drivers (Pretty et al., 2009). Urbanization and intensive agricultural practices, for example, are destroying natural habitats while eroding the rural connections of people and modifying the landscapes and seascapes in which their culture is embedded (Berkes, 2009). Roads, and the accessibility they grant to previously inaccessible places threaten ecosystems and biodiversity (Benítez-López et al., 2010), as well as rural communities who end up shifting their cultural practices as a result to being more connected to other communities and to markets (Stronza and Gordillo, 2008). Moreover, the tourism sector has a history of local community exploitation, where their knowledge and sacred sites are exploited to benefit tourists (Stronza and Gordillo, 2008). Ecotourism often creates a foothold in culturally and biologically sensitive areas that are later exploited through mass tourism development eroding all sense of local ownership and local perspectives of what makes the place unique (Barros et al., 2015; Fletcher and Neves, 2012).

For countries to develop sustainably and ethically, it is imperative to mainstream both biodiversity and cultural diversity in economic development pathways, including those linked to the tourism sector (Folke et al., 2019). Importantly, this must be done through a process that is led by local communities. Successful examples of co-management and co-production of sustainable ecotourism are those where Indigenous sovereignty is upheld by federal policies, and those where communities decide what is to be shared with tourists and what is to be kept privately (Stronza, 2007). In theory, tourism is a sector that promotes social mobility and economic growth because it is a labor-intensive sector and has the potential to generate alternative local rural livelihoods (Banco Interamericano de Desarrollo, 2017; Stronza et al., 2019). When local communities are prioritized in the tourism planning process, they are empowered to direct the outcome of the tourism they want to guide and share (Stronza et al., 2019). Although, in practice, ecotourism has failed to be an inclusive sector for many communities whose voices has been erased from planning efforts and development activities (Fletcher and Neves, 2012).

## 2 Case study selection, research questions and hypotheses

We chose Colombia as a case study for evaluating biocultural tourism potential for a few reasons including its biophysical and cultural richness, as well as recent political will and growth in the tourism sector. Colombia harbors approximately 10% of the world’s species richness, including the highest avian diversity of any country (n=1,933 of the ∼11,000 bird species in our planet) and ranks second highest in amphibian, plant, and mammal diversity (Instituto Alexander von Humboldt, 2022). More than half the territory is forested, and it is home to >90 natural ecosystems (Schrodt et al., 2019). Colombia is increasingly becoming an ecotourist destination, particularly for birdwatching, sports fisheries, hiking, and horseback riding (Lasso et al., 2019; Maldonado et al., 2018).

Colombia declared itself a multiethnic state in the national constitution (Corte Constitucional de Colombia, 1991). There are four major ethnic groups: Afro-Colombians (including Black, Raizales and Palenqueros, 6.8% of the population), Indigenous (representing over 170 nations, 4.4%), Rrom (0.006%), and Mestizos (mixed-ancestry of White-caucasian/Hispanic descent and Indigenous descent, 88.8%) (DANE, 2018). Colombia’s geographical location facilitated early human migration from Mesoamerica and the Caribbean to the Andes and Amazon basin, leading to significant archaeological sites, including UNESCO World Heritage and Intangible Cultural Heritage sites, that date back to 18,000-8,000 BCE (Sutter, 2021). Colombia is also rich in modern cultural practices and is known for its folkloric music festivals, including some of the largest carnivals in South America (Barranquilla Carnival and Petronio Alvarez) (Aguado et al., 2021). Across the country, music festivals coincide with religious holidays and celebrate annual signifiers of change, such as the harvest, the rains, and the change in seasons (Aguado et al., 2021).

There is a political determination in Colombia to prioritize tourism as part of a comprehensive, long-term sustainable development plan. Under the leadership of President Petro (2022-2026), the government aims to shift from a mining-centric economy to a bioeconomy predominantly reliant on agriculture and tourism. Presently, the tourism sector contributes 2% to Colombia’s GDP, sustaining 2 million jobs in 2018 and establishing businesses in 281 municipalities across the country (OECD, 2020). Petro’s administration is also dedicated to positioning Colombia as a sustainable tourist destination that actively empowers Afro-Colombian and Indigenous communities (Ministerio de Comercio, Industria y Turismo, 2022). Thus, for biodiversity, cultural diversity, and political reasons, Colombia presents the ideal case study to evaluate the potential of biocultural tourism to guide the sustainable development of the country.

In this study, we characterize and map biocultural richness for assessing tourism supply and demand in Colombia. We sought to answer three main questions: First, what is the spatial overlap between vertebrate species richness (hereafter ‘biodiversity’), institutionalized cultural richness, and observed tourism demand in Colombia? We tested two hypotheses: (H1) At the municipality level, we predicted cultural richness and biodiversity to be positively correlated because of previous studies suggesting that these are intricately related (Maffi, 2010). (H2) We predicted that areas of high biocultural richness coincide with places of high tourism demand, because institutionalized cultural richness and biodiversity act as pull factors drawing people to destinations.

Second, where in the country may biocultural pull factors be contributing more to observed tourism demand that accessibility push and pull factors (Figure 1)? We predicted (H3) that areas of both high accessibility and high biocultural richness have the largest observed patterns of tourism because prior research has indicated that biodiversity interacts with infrastructure to draw tourists to any given locations (Echeverri et al., 2022).

Third, which municipalities have the highest potential for developing tourism and defining themselves as biocultural destinations? (H4) We predicted that biologically and culturally sensitive areas, such as those that have been inaccessible due to the armed conflict (e.g., Amazon, Pacific region) are the ones with the highest potential to become biocultural destinations.

## 3 Methods

### 3.1 Research design and interdisciplinary collaboration

Our research project originated as a collaborative partnership among anthropologists, ethnomusicologists, lawyers, biologists, ecologists, and spatial analysts. The lead author (AE) convened the authorship team and other participants at Stanford University in March 2022. During this gathering, a two-day workshop was organized, featuring internal sessions with informative talks on our respective expertise (e.g., ADA presented on music, FAR on ethnic diversity in the country, NOP on birds, and GAHR on fishes). Half of the workshop attendees were Colombians who had grown up in diverse regions of the country, while the other half comprised students and research technicians engaged in various methods, including spatial analyses, statistical analyses, and environmental policy.

A full day was dedicated to research design, collectively formulating research questions and refining research design and analyses. To achieve this, we initiated discussions on our positionality and epistemology, striving to integrate diverse ways of knowing based on our academic and personal backgrounds. Following the workshop, virtual working groups were established to advance different components of the research. Although the authorship team primarily consists of academics and practitioners, we maintained communication with key personnel in the Colombian government. Lead author (AE) traveled to Bogota in November 2022 to conduct a training session on tourism planning for Colombia in collaboration with the Departamento Nacional de Planeación (National Planning Department), Instituto de Investigación de Recursos Biológicos Alexander von Humboldt (Science Institute), DANE (National Statistics Bureau), Artesanías de Colombia (Colombian Artisan Crafts), Ministerio de Ciencias, Tecnología e Innovación (Science Ministry), and Ministerio de Comercio, Industria y Turismo (Tourism Ministry). While government employees are not coauthors due to a conflict of interest, they were consulted for the policy recommendations section and are acknowledged accordingly.

### 3.2 Data collection

To analyze push and pull factors and map biocultural tourism supply, diverse variables were considered as proxies for biodiversity, institutionalized cultural richness, and accessibility. Tourism demand was mapped using proxies for ecotourism and culture-based tourism as well as general visitation rates (Table 1). Municipalities were selected as the units of analysis due to the granularity of cultural data provided by the national statistics bureau in its population census reports (DANE, 2018). Biocultural tourism supply and demand were mapped separately as its conventional in tourism and recreation studies (Peña et al., 2015).

**Table 1.** List of variables in the analysis describing the meaning of each one, with their respective transformations and indicated if they were included in composite indices.

| Metric | Push or pull factor | Predictor variable | Meaning | Transformations applied | Included in final metric | Data source |
| --- | --- | --- | --- | --- | --- | --- |
| **Tourism supply** |  |  |  |  |  |  |
| Biodiversity richness index | Pull factors | Bird species richness | Maximum number of bird species found in a municipality | Min max transformation | Yes | This study |
| Biodiversity richness index | Pull factors | Mammal species richness | Maximum number of mammal species found in a municipality | Min max transformation | No | This study |
| Biodiversity richness index | Pull factors | Amphibian species richness | Maximum number of fish species found in a municipality | Min max transformation | No | (Jenkins et al., 2013) |
| Biodiversity richness index | Pull factors | Reptile species richness | Maximum number of reptile species found in a municipality | Min max transformation | No | (Caetano et al., 2022; Roll et al., 2017) |
| Biodiversity richness index | Pull factors | Fish species richness | Maximum number of fish species found in a municipality | Min max transformation | Yes | This study |
| Biodiversity richness index | Pull factors | Ecosystems richness | Maximum number of ecosystems found in a municipality | Min max transformation | Yes | (IDEAM et al., 2017) |
| Institutionalized cultural richness index | Pull factors | World Heritage Sites | # of UNESCO world heritage sites in a municipality | Min max transformation | Yes | (UNESCO, 2023a) |
| Institutionalized cultural richness index | Pull factors | Intangible Cultural Heritage Sites | # of UNESCO intangible cultural heritage sites in a municipality | Min max transformation | Yes | (UNESCO, 2023b) |
| Institutionalized cultural richness index | Pull factors | Music festivals of endemic musical rhythms | # music festivals from endemic musical rhythms in a municipality | Min max transformation | Yes | (Abadía Morales, 1995; Ochoa Escobar, 2013; Ochoa et al., 2014) and this study |
| Institutionalized cultural richness index | Pull factors | Afro-Colombian Lands | # of Afro-Colombian territories in a municipality | Continuity correction, log transformation, min max transformation | Yes | (Agenda Nacional de Tierras, 2023a; Unidad de Restitución de Tierras, 2023) |
| Institutionalized cultural richness index | Pull factors | Indigenous Lands | # of Indigenous reserves in a municipality | Continuity correction, log transformation, min max transformation | Yes | (Agenda Nacional de Tierras, 2023b) |
| Institutionalized cultural richness index | Pull factors | Museums density | # of museums per km<sup>2</sup> in each municipality | Density calculation (# museums/ area of the municipality) + continuity correction + log + min max transformation | Yes | (Ministerio de Cultura, 2023) |
| Accessibility index | Pull factors | Bird lodging density | # of bird lodges per km<sup>2</sup> in each municipality | Density calculation (#bird lodges/ area of the municipality) + continuity correction + log transformation + min-max transformation | Yes | (Beckers and Flores, 2013) |

| **Metric** | **Push or pull factor** | **Predictor variable** | **Meaning** | **Transformations applied** | **Included in final metric** | **Data source** |
| --- | --- | --- | --- | --- | --- | --- |
| Accessibility index | Pull factors | Road density | built roads per km<sup>2</sup> in each municipality | Density calculation (km of roads/ area of the municipality) + continuity correction + log transformation + min-max transformation | Yes | This study, INVEST visitation model (Sharp et al., 2020; Wood et al., 2013) |
| Accessibility index | Push factor if inaccessible, reverse coded to a pull factor | Distance to nearest airport | Distance to nearest airport in each municipality, reversed | Distance to nearest airport (km) multiplied by -1 (to reverse it) + min max transformation | Yes | This study, our airports database (Megginson, 2023) |
| Accessibility index | Pull factors | Lodging density | # of hotels and lodges per km<sup>2</sup> in each municipality | Density calculation (# hotels/ area of the municipality) + min max transformation | Yes | (GeoNames, 2023) |
| Accessibility index | Push factor as a conflict zone (reverse coded to be peaceful as a pull factor) | Peaceful | # of armed conflict cases per municipality divided by the area, reversed | Density calculation (# Conflict events/ area of the municipality) + continuity correction + log + reversing it (multiplied by -1) + min max transformation | Yes | (Centro Nacional de Memoria Histórica, 2022) |
| **Tourism demand** |  |  |  |  |  |  |
| Demand index | NA | eBird hotspots | # eBird hotspots per km<sup>2</sup> in each municipality | Density calculation (# eBird hotspots/ area of the municipality) + continuity correction + log + min max transformation | Yes | (Levatich, 2019) |
| Demand index | NA | Sport fisheries | # sport fisheries per km<sup>2</sup> in each municipality | Density calculation (# sport fisheries/ area of the municipality) + continuity correction + log + min max transformation | Yes | (Heinsohn Mallarino, 2023), this study |
| Demand index | NA | Flickr PUD | # of Flickr Photo-User-Days | Continuity correction, log transformation, min max transformation | Yes | This study, INVEST visitation model (Sharp et al., 2020; Wood et al., 2013) |
| Demand index | NA | Airport visitors | # of passengers arrived at each airport | Continuity correction, log transformation, min max transformation | Yes | (Aeronáutica Civil, 2019) |
| Demand index | NA | Music festivals visitors | # of visitors at music festivals | Continuity correction, log transformation, min max transformation | Yes | Local newspapers, local Facebook pages, (DANE, 2018) |

Extensive data were gathered from Colombian government sources (e.g., national census), international databases (e.g., UNESCO), as well as archives, newspapers, and books. Data collection was complemented by interdisciplinary team expertise (i.e, ethnomusicology, anthropology, and biology) and collective fieldwork experiences across the country spanning multiple decades.

#### 3.2.1 Biodiversity pull factors

Biodiversity pull factors were conceptualized as the richness of vertebrate species and ecosystems. Prior studies have indicated that people travel to locations to find specific animals (Echeverri et al., 2019), to recreate in different ecosystems (e.g., hike in mountains vs. lay on the beach) (Castaño-Isaza et al., 2015; Mancini et al., 2018), and both sports fisheries and birdwatching are important economic tourism subsectors in Colombia (Lasso et al., 2019; Maldonado et al., 2018).

To calculate the biodiversity at each municipality, vertebrate species richness data were gathered including amphibians, reptiles, birds, mammals, and freshwater fish species for continental Colombia and excluded islands and marine ecosystems (Figures S7-S18). Species range maps for amphibians and reptiles were cropped from global data sources (Table 1, FigureS10, FigureS12). The maximum number of species found in each pixel was counted and aggregated to municipality. To estimate the species richness of birds and mammals, a few methods were combined: expert range maps, ecoregions, elevation data, and validated crowdsourced information from eBird (Levatich, 2019) and the Global Biodiversity Information Facility (GBIF, 2023). After individual species range maps were created, these were rasterized to get a final species count, maximum number of species at each municipality was extracted (See supporting information for detailed methods, TableS3).

The distribution range for 92% of Colombian freshwater fish species was estimated by calculating a minimum spanning tree approach (Grill et al., 2014) among the river reaches with known published occurrences for each species. Fish occurrences from GBIF, AmazonFish (Jézéquel et al., 2020), and FishNet2 (Fishnet2, 2023) were matched to the nearest river reach in this simplified river network. For each species, the shortest path among all possible pairs of river reaches where a given fish species is present was calculated (FigureS15, TableS3).

A variable of ecosystem richness was added because tourism studies have documented that visitation patterns are determined by ecosystem type (e.g., beaches vs. forests) (Hale et al., 2019; Wood et al., 2013). As such, we used a government-led national map with several different ecosystem classifications for each municipality (IDEAM et al., 2017). Coarse ecosystem classification were used and we counted the maximum number of ecosystems at each municipality. All data processing and analyses were done in ArcGIS Pro, QGIS, and Statistical software R (R Development Core Team, 2023). Individual packages used for each variable are cited in the supporting methods.

#### 3.2.2 Institutionalized cultural richness pull factors

Cultural richness was conceptualized as anything that would draw tourists to a location, and as anything that Indigenous peoples and local communities would be willing to share with visitors. Variables that were not deemed as sharable, such as sacred sites, or that were not relevant for tourism per se, such as native languages, were excluded. Here we coin the term “institutionalized cultural richness” because most of the information we were able to find regarding Indigenous territories, Afro-Colombian lands, archaeological sites, museums, and music festivals, reflected the cultural diversity that has been validated by international and national institutions like UNESCO and the Colombian Ministry of Culture.

Colombia’s history of cultural mixing gave way to communities like the *campesinos* (rural dwellers) and *mestizos*, shaped by the intermixing of Indigenous, Caucasian, and Afro-Colombian heritages. Originating from Spanish colonization and the blending of European settlers, Indigenous peoples, and African slaves, these groups embody diverse cultural intersections. Rather than constituting defined ethnic groups, they symbolize cultural groups that exhibit rich cultures through their artisan crafts, food, dances. Unfortunately, these cultures are not captured in available data.

To map institutionalized cultural richness, six variables were identified (Table 1). UNESCO World Heritage Sites, and UNESCO Intangible Cultural Heritage sites were collected from the UNESCO world platform and include information on communities’ Traditional Ecological Knowledge and archaeological sites (UNESCO, 2023a, 2023b). Those sites were mapped to municipalities or to polygons delimiting the area (e.g., protected areas) (Figures S19-S30). Information for all the registered museums in the country, Indigenous reserves and Afro-Colombian territories were collected from government sources and mapped (Table 1). Lastly, coauthor ADA, an ethnomusicologist identified all the music festivals that celebrate endemic music genres and mapped them to municipalities (See Supporting Information for detailed methods, and Figures S19-S30 for spatial distribution of cultural layers).

#### 3.2.3 Accessibility push and pull factors

Building on previous studies that determine accessibility to tourist destinations (e.g., Echeverri et al., 2022; Ocampo-Peñuela and Winton, 2017), we considered the following variables that make locations accessible: proximity to airports, density of built roads per municipality, lodging and birdwatching lodging locations (Table 1). Information about the proximity to airports and road density were downloaded from national government sources (See Supporting Information). Georeferenced hotels and lodges were downloaded from global datasets. Birdwatching locations were taken from a national birdwatching guide, and downloaded as points that were mapped to municipalities (Table 1, Figures S31-S40).

Violence and armed conflict were conceptualized as push factors that would preclude people from going to a location. The total number of violent events that happened in each municipality between 1985 and 2022 were calculated, and a metric was developed and reverse coded to estimate peacefulness in a municipality, so it became a pull factor (Table 1, Figure S39). All analyses were done in R and Arc GIS Pro.

#### 3.2.4 Tourism demand variables

Proxies for tourism demand included indicators of ecotourism, general tourism and cultural tourism. Five variables for tourism demand were calculated (Table 1, Figures S41-S50). eBird hotspots were mapped to determine desired destinations for birdwatchers sensu Echeverri et al (2022). Presence and absence of recreational freshwater fisheries was also mapped as a metric of demand for this activity (Lasso et al., 2019). The InVEST Visitation model (Sharp et al., 2020) was used to estimate visitation from geotagged photos posted to the photo sharing site Flickr. The model calculates the number of “photo user days” (PUD) within each municipality.

Moreover, the median arrival of passengers at each airport was calculated from airport arrival data (2004-2019) from national government sources (Table 1). Based on information available in local newspapers and Facebook pages, music festival attendance for 19 festivals (of the 105 festivals) was recorded. For the rest of the festivals (n=86) we estimated that attendance was 10% of the municipality’s population because music festivals are often intended to celebrate the place and the community (Aguado et al., 2021) (Table 1).

### 3.3 Data Analysis

Individual data layers were condensed into four composite indices as follows: 1) biodiversity richness index, 2) institutionalized cultural richness index, 3) accessibility index, and 4) tourism potential index, and 5) tourism demand index (Figure S1). To do so, variables were transformed into densities by municipality, and we attempted to make the distribution of each variable normal. Continuity corrections (by replacing the 0s with the smallest number of the vector divided by 2) were performed, to then apply logarithmic transformations (when appropriate). In all cases we performed a min-max transformation that would allow us to rescale variables to comparable units. A complete list of the variables with their transformations is shown in Table 1.

Data distributions were inspected visually of both untransformed and transformed variables to decide on which transformations to perform (a complete list of the variable histograms is included in Figures S5-S50). All statistical analyses, data transformation and subsequent calculations were performed in R (R Development Core Team, 2023).

To create the composite indices, we first calculated pairwise correlations between individual variables that we wanted to include in the indices (e.g., between all biodiversity variables, all cultural variables, etc). Spearman’s correlations were used because many of our variables were ordinal/categorical (e.g., cultural variables), and because we expected non-linear relationships between variables (Bonett and Wright, 2000). The results of all four correlograms with Spearman’s r are presented in the supporting information (Figures S3-S6). We dropped variables that were correlated with a Spearman’s r> 0.5 as in (Echeverri et al., 2020) and only included uncorrelated variables in composite indices to avoid collinearity. When a pair of variables were highly correlated, we dropped one of them. Our final list of variables included in composite indices is shown in Table 1.

Finally, with the transformed, uncorrelated variables (Table 1), five composite indices were calculated using the equations below. All variables were given equal weight because there was no reason to expect that some would be more important than others, making the most parsimonious analysis possible.

**Equation 1. Biodiversity richness index** = ( ⅓ * Fishes sp. richness) + ( ⅓ * Bird sp. richness) + ( ⅓ * Ecosystems richness)

**Equation 2. Institutionalized cultural richness index** = ( ⅙ * UNESCO World Heritage Sites) + ( ⅙ * UNESCO Intangible Cultural Heritage) + ( ⅙ * Endemic music festivals) + ( ⅙ * Afro-Colombian territories) + ( ⅙ * Indigenous Reserves) + ( ⅙ * Museums)

**Equation 3. Accessibility index** = ( ¼ * Bird lodging density) + ( ¼ * Road density) + (¼ * Distance to nearest airport) + (¼ * Lodging density) + (¼ * Peaceful)

**Equation 4. Biocultural tourism potential index** = ( ⅓ * Biodiversity richness index) + ( ⅓ * Institutionalized cultural richness index) + ( ⅓ * Accessibility index)

**Equation 5. Tourism demand index** = ( ⍰ * eBird hotspots) + (⍰ * Sports fisheries) + (⍰ * Flickr PUD) + (⍰* Airport visitors) + (⍰ * Music festivals visitors)

Finally, to test hypotheses, Spearman’s correlations were used to determine the statistical association between composite indices at national and regional scales (Figure S2). We also used qualitative interpretation of spatial patterns to describe the association between indices by analyzing municipalities in the top and bottom quartiles of the distributions. Workflow analyses (Figure S1), descriptive statistics (Table S2), and results of correlations (FigureS2) are in the supporting information file.

## 4 Results

### 4.1 Biocultural richness supply and demand

Our analysis indicates that across the country, there are variations in biodiversity richness, institutionalized cultural richness and accessibility (Figure 3). Biodiversity richness is greatest in the Amazon region and lowest in the high-elevation municipalities across the Andean (Figure 3a). Bird diversity is highest at the Andes-Amazon transition, whereas fish diversity is highest in the downstream Amazonian River basin (Figure S8). Ecosystem richness is greatest in the northernmost Caribbean region (Figure S18), with 19 ecosystem types found in the Uribia municipality in Guajira, the northernmost state in the continental country (Table S1).

**Figure 3.**
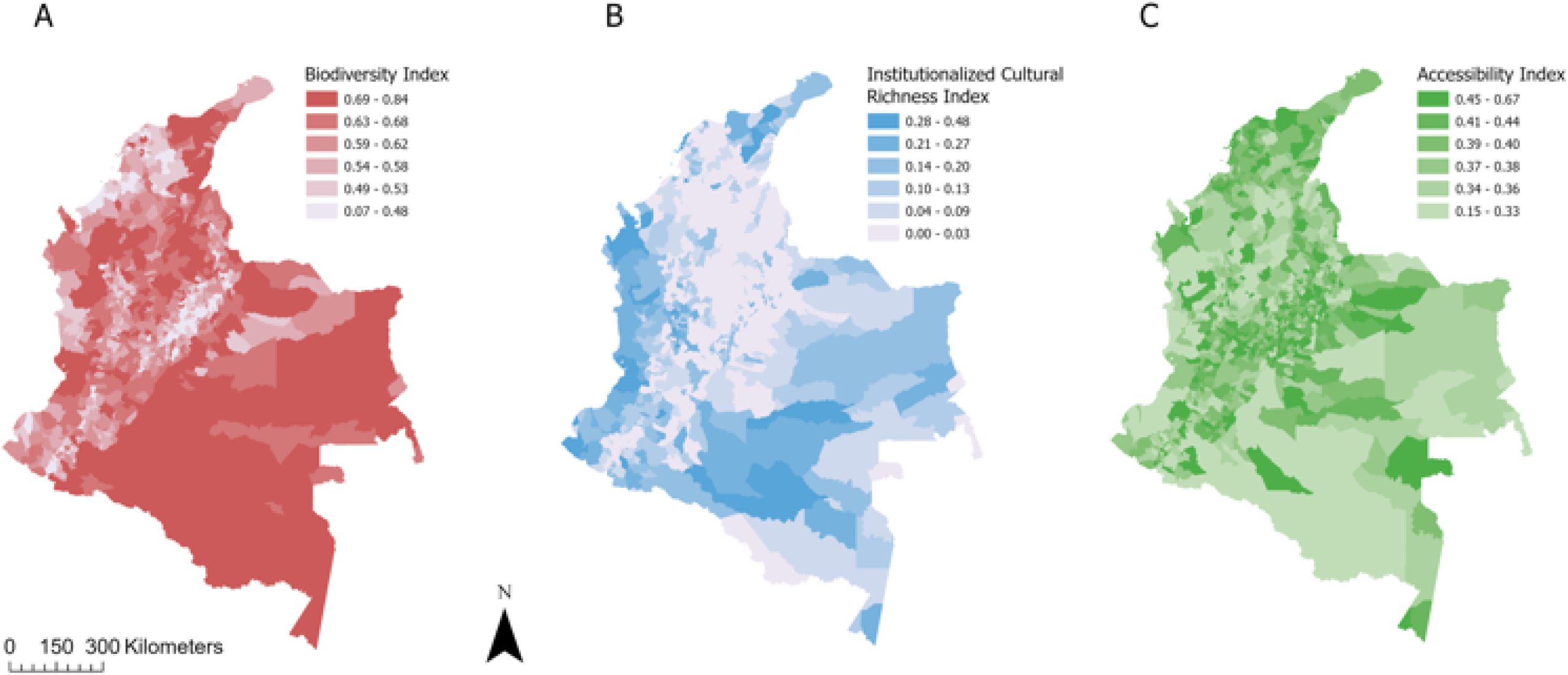
In Colombia, biodiversity richness (A), institutionalized cultural richness (B), and accessibility (C) vary spatially across the country. The Amazon is the most biodiverse region (A), while the institutionalized cultural richness shows that municipalities in the South Pacific, and Amazon are the richest (B). Accessibility is spatially heterogenous throughout Colombia, with higher values around the Andes and especially in the largest urban municipalities (e.g., Bogotá and Cartagena) (C). Index values are scaled and presented here in six breaks to show the spatial variations.

Institutionalized cultural richness is greatest within municipalities in the Pacific, Amazon, and northern Caribbean regions (Figure 3b). Afro-Colombian communities are clustered along the Pacific coast, and Indigenous reserves are mostly concentrated in the western Amazon followed by the Pacific coast (Figures S26, S28). UNESCO World Heritage Sites are in the central Amazon and northern Pacific regions and UNESCO intangible cultural heritage sites are primarily in the northern Caribbean region, around the Sierra Nevada de Santa Marta (Figures S20, S22). Music festivals are distributed throughout all geographic regions, with a higher concentration in the Pacific and Caribbean regions (Figure S24).

Our results suggest that the biodiversity richness index and the institutionalized cultural richness index are positively and significantly correlated at a national level (Spearman’s R=0.22, p=2.5e-14, Figure S2). They are also positively and significantly correlated at the regional level for all biotic regions except Amazon and Orinoquia (Figure S2). Therefore, we have partial evidence to support our first hypothesis predicting that biodiversity and cultural richness would be correlated (H1). However, while some of these correlations are positive and significant, there are spatial variations that cannot be overlooked. Areas of high biodiversity richness and areas of high institutionalized cultural richness do not always overlap (Figure 3, Figure 4). In some municipalities both are high (e.g., San Vicente del Caguán in the Amazon region, Table S1), but some have high biodiversity only (e.g., Puerto Rico in the Amazon Region), while some have high institutionalized cultural richness only (e.g., Riosucio in the Andean region) (Table S1).

**Figure 4.**
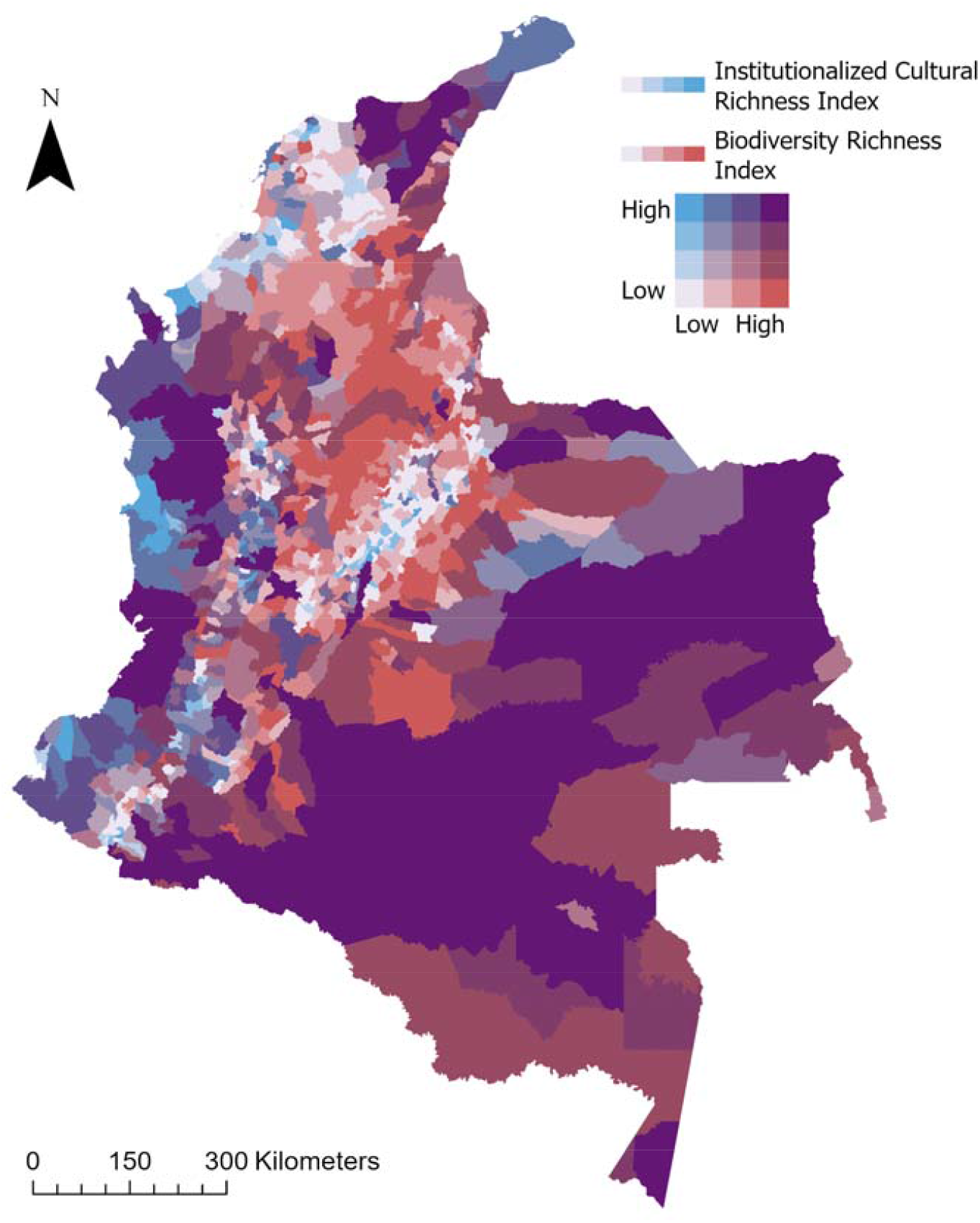
Biocultural pull factors (i.e., areas where biodiversity and institutionalized cultural richness coincide) are higher in the central Amazon, Orinoquia, central Pacific, and northern Caribbean regions. The Pacific and Caribbean regions are also rich in culture, while the Andean foothills are rich in biodiversity. The Andean peaks show low biodiversity and low institutionalized cultural richness. Standardized indices are visualized in quartiles, where purple pixels represent municipalities that are in the highest quartiles for both biodiversity and cultural richness, whereas red pixels are those that are only rich in biodiversity, and blue pixels are those that are only rich in culture.

We find evidence to support our second hypothesis predicting that areas of biocultural tourism potential coincide with areas of high tourism demand at a national scale (Spearman’s R=0.39, p<2.2e-16) (H2) (Figure 5a, b). Moreover, while doing pairwise correlations between individual indices, we find positive and significant correlations between biodiversity richness and tourism demand (Spearman’s R=0.13 p=1.08e-5), and between institutionalized cultural richness and demand (Spearman’s R=0.33 p<2.2e-16). Spatial patterns indicate that tourism demand in Colombia is greatest in the northern Caribbean and Orinoquía regions, as well as some municipalities in the Pacific and Andean regions (Figure 5). The patterns in the Orinoquia region seem to be driven by recreational fisheries and fish biodiversity (Figures S14, S44).

**Figure 5.**
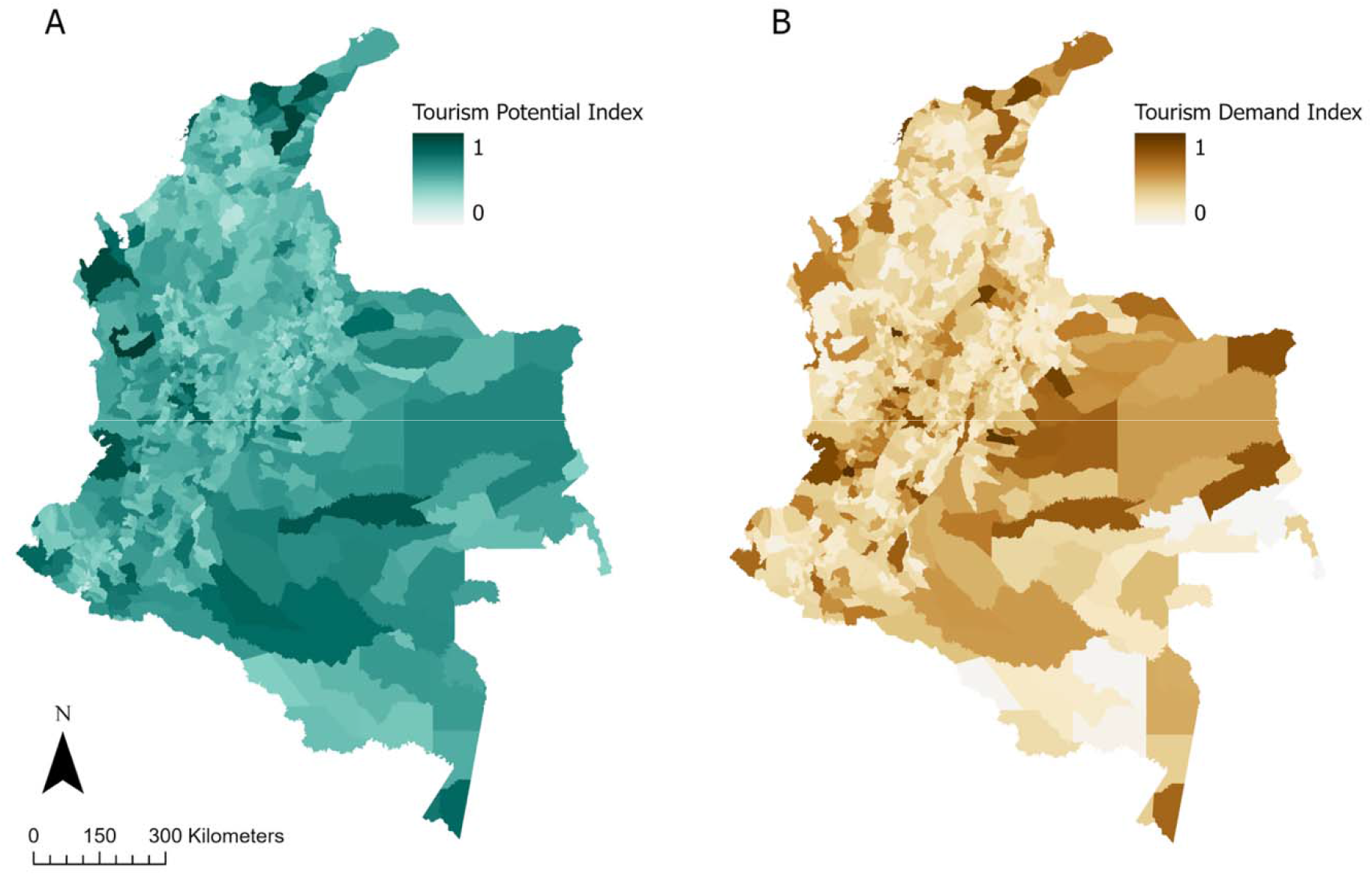
Biocultural tourism potential is correlated with tourism demand at a national scale. The Caribbean and Pacific regions have municipalities that are biocultural destinations that are somewhat accessible offering a tourism supply (A) while the Orinoquia and Caribbean regions show the greatest tourism demand (B). The standardized index of 0 (lowest) to 1 (highest) in the distribution is shown for each index.

### 4.2 Accessibility supply and tourism demand

Correlation tests give evidence to support our third hypothesis predicting that areas of high accessibility are also areas of high tourism demand (Spearman’s R=0.23, p<1.15 e-15) (H3). However, we do not find evidence to support locations where all three are high: accessibility, biodiversity, and cultural richness. No municipalities are in the top quartile for all three tourism pull factors indices: biodiversity richness, institutionalized cultural richness, and accessibility. Instead, there are spatial trade-offs between biocultural richness and accessibility. For example, the municipalities with the greatest accessibility are in the Andean region and include Subachoque and Sabaneta (I=0.67, I=0.6, respectively) (Figure 3c, Table S1). These municipalities have medium tourism demand (I= 0.38, I=0.49, respectively) (Figure 5b, Table S1). Subachoque (Biodiversity I=0.46, Cultural I=0) and Sabaneta (Biodiversity I=0.47, Cultural I=0) are in the lowest quartile for both biodiversity richness and institutionalized cultural richness.

### 4.3 Areas for potential biocultural tourism development

We do find evidence to support our fourth hypothesis predicting that most areas where biocultural richness is high are already protected. Either in the form of national parks, Indigenous reserves, or Afro-Colombian territories (H4). Municipalities with the greatest potential for biocultural tourism development are in the Amazon region (such as Mirití-Paraná, Puerto Colombia) and in the Pacific region (Argelia, and Carmen del Darién) which lie within Indigenous reservations and Afro-Colombian territories (Figure 6). While our findings suggest that most municipalities where tourism supply (as measured by biodiversity, cultural richness, and accessibility) is higher than tourism demand (Figure 5c, and Figure 6) lie within Afro-Colombian territories and Indigenous reserves, we find that only a few of them are under the jurisdiction of the Colombian government, such as San Juan del Cesar and Aracataca.

**Figure 6.**
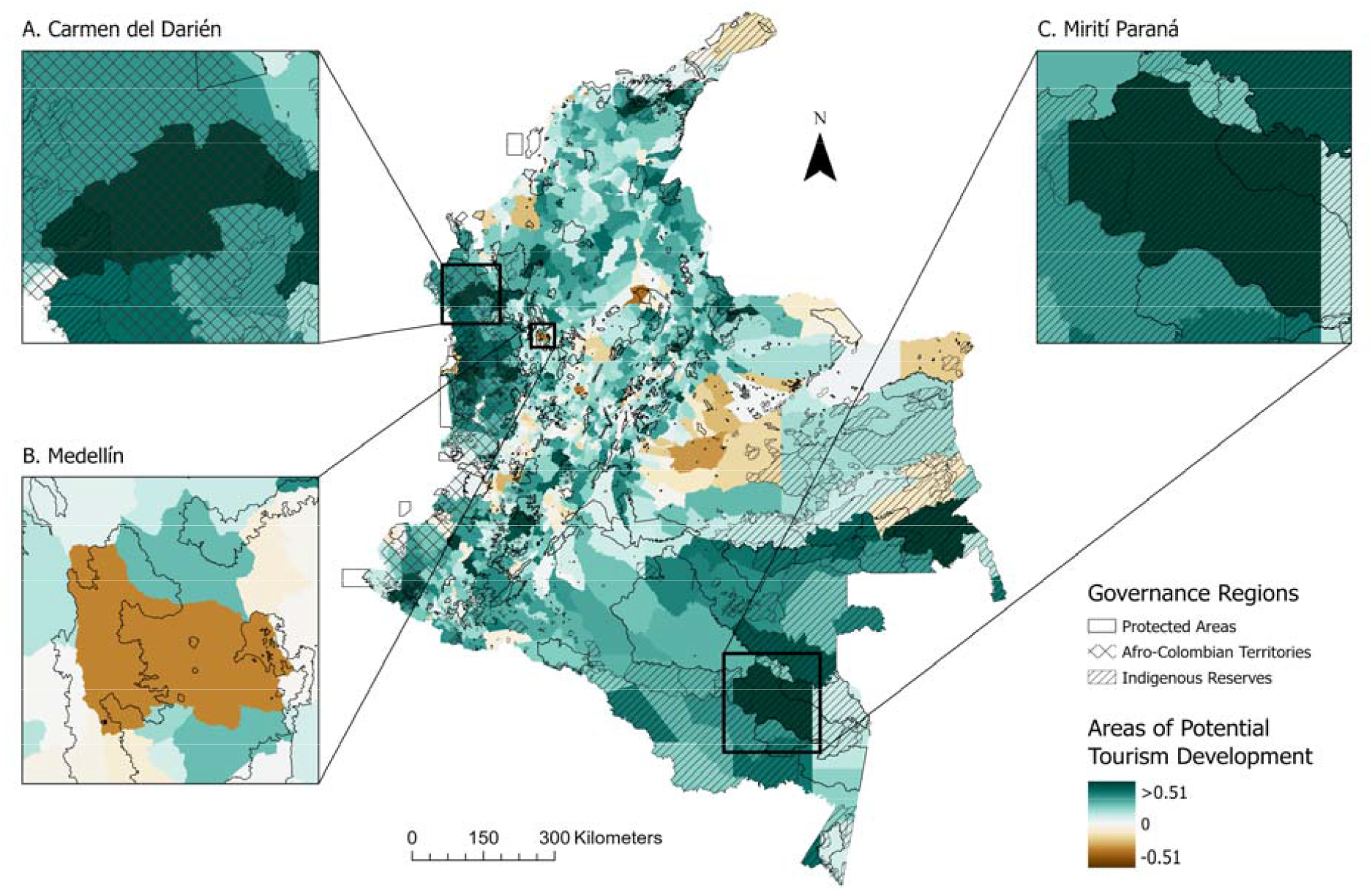
Areas of potential tourism development lie within Afro-Colombian territories and Indigenous reserves. This map shows the difference between tourism supply and tourism demand, where a high value (green) indicates greater supply of biocultural diversity and accessibility relative to current tourism demand, and a negative value (brown) means more demand is observed than the biocultural tourism supply. Three municipalities stand out as areas where biocultural tourism could be developed. Carmen del Darién (Pacific region, Afro-Colombian territories) is on the top 25% for biodiversity and institutionalized cultural richness but is in the top 25% for accessibility. It has low tourism demand (I=0.09) and has a low ranking for peacefulness and proximity to an airport, little road infrastructure, and no lodging (A). Mirití-Paraná (Amazon region, Indigenous reserves) is also on the top 25% for biodiversity and institutionalized cultural richness but has low tourism demand (I=0.02), is close to an airport and has a high peacefulness rating, though it has no lodging or roads (B). These municipalities have the greatest potential for infrastructure development. Conversely, Medellín has high tourism demand (I=0.98) but is in the bottom 50% for accessibility mostly because it ranks the lowest in peacefulness (highest in violence and crime rates). It has the third highest density of roads and lodges in the country is in the top 25% for richness of cultural attractions and top 50% for biodiversity richness (C).

## 5 Discussion

By combining sociocultural and ecological data at the municipality scale, our work provides a roadmap for integrating biocultural values into spatial planning for the conservation and tourism sectors. We show that biodiversity richness and institutionalized cultural richness are correlated at a national scale and in some regional scales, and that both are also correlated with tourism demand. We also demonstrate how this method and analysis can be used for Colombia but believe this framework can be adapted to other countries and be applied at different scales (local to global). To our knowledge, our work is the first empirical study to operationalize the push-pull theory (Dann, 1976) in a spatially explicit manner for biocultural tourism approaches.

The tourism industry is a growing sector internationally (Balmford et al., 2009). In our attempts to identify Colombian destinations that preserve biodiversity, institutionalized cultural richness, and are accessible simultaneously, we encountered significant spatial trade-offs: there are no win-win-win locations for biodiversity, culture, and accessibility in the country, as we could not find a single municipality that was in the top quartile for all three. Nonetheless, the correlations between biodiversity and institutionalized cultural richness suggest that indeed, in some contexts (nationally and in some regions) these variables might be intertwined, as previously suggested (Maffi, 2005; Pretty et al., 2009; Sterling et al., 2017), although with regional variations.

### 5.1 Strengths of our methodological approach and key findings

The main contribution of this paper is methodological: a biocultural tourism framework implemented through geospatial analysis is a useful tool for guiding tourism planning at national and regional scales that are trying to optimize for different objectives, such as protecting both cultural and biological diversity. The framework (Figure 1) and the methods developed here resulted from a collaborative effort stemming from both the in-person workshop and virtual working group meetings. Judging from all our personal experiences doing research in ecology and environmental sciences, workshops that intentionally bring together social scientists, natural scientists, artists, and humanists is rare. Doing so, elevated the interdisciplinary integration of diverse data sets, methods, analytical tools, and approaches to conduct this research study.

Our results reveal that while some municipalities have high biodiversity and institutionalized cultural richness, others excel in only one, particularly in the Andean region. Several systemic social and institutional factors could be contributing to the spatial mismatch between biodiversity, institutionalized cultural richness, and accessibility. Colombia’s history of colonization and Constitutional amendments in 1991 recognize Colombia as a multiethnic state (Velasco, 2011). Law 70 of 1993 further acknowledged the rights of Afro-Colombian and Indigenous communities to collective land ownership, emphasizing the preservation of cultural identity and participation in development decisions (Bonilla-Mejía and Higuera-Mendieta, 2019; Echeverri et al., 2023).

Positive and significant correlations between tourism potential and tourism demand, indicate that at least in Colombia, tourism might be a mixture of ecotourism, culture-based tourism, and opportunistic tourism driven mostly by accessibility. Looking at national and regional spatial patterns (Figure 6) it is evident that as a country, Colombia caters to the different types of tourism. Importantly, regions like the Pacific and Caribbean are the ones that show most evidence for biocultural tourism potential and demand.

Indigenous reserves, Afro-Colombian territories, and land under national and subnational governments have experienced divergent development pathways, leading to unequal accessibility to planning resources and uneven funding distribution (Van De Sandt, 2003; Velasco, 2011). This historical trajectory has also resulted in different land-use impacts that together shape the institutionalized cultural richness, biodiversity richness, and accessibility to these lands. For example, it is evident that the Andean region is comparatively more accessible than Afro-Colombian and Indigenous territories.

### 5.2 Limitations: A critical perspective on national studies without field work

The tourism sector, while offering economic benefits, poses threats to nature, culture, and Indigenous Peoples (Stronza et al., 2019). Exploitation occurs when local communities are excluded from tourism projects, denying them rights and voices (Hunt and Stronza, 2011). Our study has a major limitation as we did not consult with Indigenous peoples and local communities to prioritize data sources that would represent their cultural richness. Nevertheless, our research is not prescriptive; we identify potential biocultural richness areas to guide future community consultations and fieldwork analyses for a nuanced understanding at local scales.

Our analysis faces limitations in cultural representation given data availability constraints. The islands of San Andrés, Malpelo, Gorgona, and Providencia are excluded from our analysis due to incomplete data sets across all index factors. However, they hold important cultural and social significance; the island of San Andrés is home to the largest population of Raizales, an ethnic group recognized by the Colombian constitution (Portz et al., 2022). The lack of representation of this vibrant community further demonstrates the need to build robust data sets in collaboration with Indigenous peoples and local communities, to accurately identify multi-sectoral partners, local interest groups and rights holders for equitable tourism development.

Our study recognizes institutionalized cultural diversity but fails to encompass non-geospatially linked forms, like beliefs, livelihoods, and norms (Pretty et al., 2009). It overlooks the diversity within blended communities, neglecting Andean communities (e.g., campesinos, mestizos, blancos, colonos). This study, authored by individuals from these communities, emphasizes the need for institutions like UNESCO to develop protocols for eliciting cultural diversity in blended communities. For example, we feel like our individual cultures and nuances in our music and traditional foods are not represented in this study. Finally, to enhance tourism representation, our cultural index should encompass gastronomic diversity, artisanal crafts, literature, art, clothing, and other sought-after cultural expressions by tourists.

Lastly, a main caveat of our analysis is the scale. We used municipality as the unit of analysis because socio-cultural data, when collected, are available for this scale as determined by census tracts. Unfortunately, the variation between municipality areas leads to biases especially as they pertain to the variables that are counts (e.g., species richness, number of music festivals).

### 5.3 Future research directions

Future research could incorporate data from additional taxa, such as butterflies, orchids, and fungi, which are currently excluded, but are known to be sought out by tourists in this context (Caputo et al., 2008; Millican, 1891). We could also add island and marine ecosystem, as they constitute a key part of the tourism sector of the country, activities like whale watching and scuba diving are important for island and coastal communities but were not included in our study. Further, expanding the cultural index to include gastronomic and artisanal destinations could cater to tourists seeking unique experiences. These layers could be co-developed through a participatory approach with Indigenous peoples and local communities, to address nuances of cultural richness that are not represented by our analyses. Field work, and ethnographic methods could be integrated with spatial analyses to better address the limitations of our current analysis.

Incorporating social media data from platforms like Instagram and Tripadvisor could enhance tourism demand estimation and market segmentation. Further research might develop methods for urban contexts, acknowledging cities as significant cultural hubs. Our current method overlooks urban cultural richness, except for museums, omitting the idea that cities are places of cultural convergence, blending communities, and exhibiting richness not captured here. Future methods can integrate these aspects into the exploration of biocultural heritage.

### 5.4 Policy recommendations

Improving accessibility and increasing tourism development in certain regions is not a trivial effort, which leads us to provide a set of policy recommendations. Our study suggests that a regional approach to biocultural tourism development would be more appropriate than a national one, given the regional differences found here. Doing so will require roundtables that convene actors representing the three jurisdictions: national state, Afro-Colombians, and Indigenous people so that they can together align their visions for development and articulate concerted efforts between spatial ordering plans (government tools), and “planes de vida” (community planning tools). While Indigenous and Afro-Colombian people have the right to govern their own territories (via planes de vida), our study show that their lands are ideal for biocultural tourism.

Free Prior and Informed Consent should be upheld when consulting with any Indigenous or local community whose territory is being under consideration for a tourism destination (McGee, 2009). Both biodiversity and cultural diversity are at risk if tourism planning efforts fail to include local voices, and follow guidelines for cumulative environmental impact assessments (Singh et al., 2020).

Second, we also point to the need of reporting cultural data in similar formats as biodiversity data. We were unable to add artisan crafts, gastronomic diversity, and other forms of cultural expressions because these data exist in narrative forms and oral traditions. Having an account of the cultural and biological heritages in tabular and spatial formats might lead to a better understanding of biocultural richness through the integration between biodiversity and cultural data. Gathering this information can be achieved through participatory approaches with Indigenous peoples and local communities, and through roundtables between the environment, culture, and tourism ministries, and the national statistics bureau and planning department. Doing so also implies more inter-institutional coordination, and fieldwork with local communities. At the global scale, we also recommend an integration of biological and cultural data in portals like the UNESCO world heritage sites, including ones that also recognize blended communities.

Lastly, tourism planning efforts can be paired with community outreach and education programs that celebrate the biodiversity and cultural diversity of each region or municipality. Such programs could highlight the importance of preserving and restoring local traditions and culturally important species, particularly with children, so these places get to celebrate and share their biocultural heritage with visitors in perpetuity.

## 6 Conclusion

This paper underscores the critical importance of assessing the biological, cultural, and access dimensions of tourism. Sustainable and equitable tourism development can be a vehicle for protecting biodiversity, preserving cultural heritage, and improving livelihoods. Mapping biological and cultural tourism attractions allows us to identify specific locations with the greatest tourism potential. Our methods and results can help optimize national and regional tourism development strategies that seek to align conservation goals with economic development that improves social mobility. Tourism strategies must be developed in partnership with Indigenous peoples and local communities at all planning stages to ensure their autonomy in dictating which parts of their culture and landscape, if any, they are willing to share and commodify. Biocultural tourism might be a pathway to develop Colombia in a non-extractive way if communities are consulted in the process. We hope this approach serves to guide not just Colombia’s national tourism strategy, but also guide international tourism strategies that are aiming to bring together countries’ cultural and biological heritages while improving social-ecological resilience and well-being.

## 7 Acknowledgements

We wrote this manuscript from the unceded territory of the Muwekma-Ohlone peoples. We celebrate the perseverance of all Indigenous Peoples across the world who stand strong in preserving their cultural traditions and connection to the land, such as the ones we write about in this manuscript. We are grateful to Lori Epsworth for helping us organize the workshop that led to this publication. We are indebted to Tenzin Norzin, Sergio Sánchez Lopez, Dr. Hector Angarita, Dr. Tong Wu, and profs. Gretchen Daily, Tadashi Fukami, and Rodolfo Dirzo for insightful conversations. We are also grateful to conversations with the Departamento Nacional de Planeación, Artesanías de Colombia, Instituto de Investigación de Recursos Biológicos Alexander von Humboldt, Ministerio de Ciencias, and Ministerio de Comercio, Industria y Turismo that influenced our research design and policy recommendations.

## 8 Positionality statement

We are 18 scientists and practitioners who deeply value biodiversity and cultural diversity. Nine of us grew up in Colombia and together we represent many places including the United States, Mexico, Ireland, Chile, Brazil, Germany, and Viet Nam. None of us identify as Afro-Colombian or Indigenous. Our academic backgrounds include ecology, evolutionary biology, botany, music, anthropology, geography, and public health. We all have postgraduate academic education (Masters or PhDs underway). We span different career stages. Many of us have traveled to and within Colombia as international or domestic tourists. We acknowledge the prevalence of a Western scientific and postpositivist worldview in writing this paper. We also acknowledge that here we are treating nature and culture as entities that we can measure, quantify, and map.

## 9 Funding

We acknowledge funding from the Science, Ethics and Technology fund awarded to AE to host a workshop titled: Biocultural values for sustainable tourism in Colombia, where we convened the author group and where this idea was born. GAHR acknowledges funding from the Ministry of Sciences, Technology, and Innovation of Colombia (Call for doctorates abroad No. 860 from 2019) and through the Graduate School of the University of Tennessee, Knoxville.

## 10 Conflict of interest statement

The authors declare no conflict of interest.

## 11 Data availability statement

Post-processed data (such as shapefiles and results by municipalities) and R scripts will become available once the manuscript gets published. The authors will make data available in Dryad, a public repository. Raw data, particularly as it pertains to fishes, mammals, and birds is available upon request to coauthors GAHR, AFSC, and NOP.

## 12 Author contributions

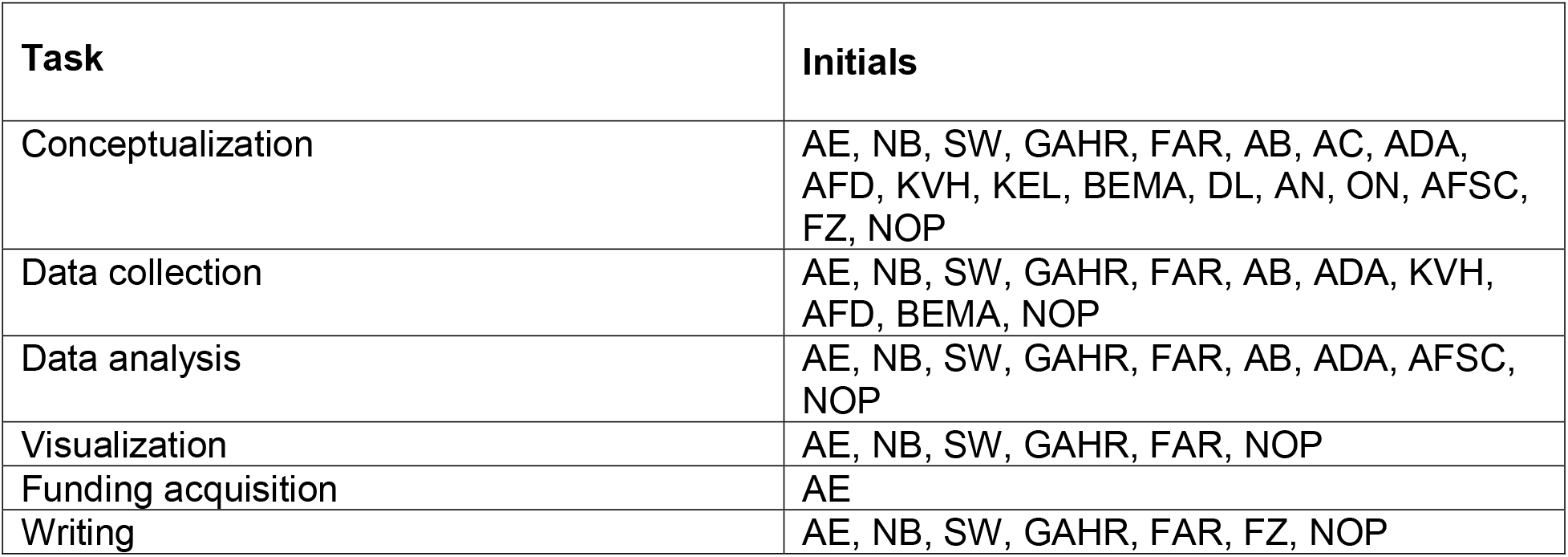

**13 References**

## Notes

### Competing Interest Statement

The authors have declared no competing interest.

