## Supporting Information for "Toward Sustainable Biocultural Tourism: An Integrated Spatial Analysis of Cultural and Biodiversity Richness in Colombia"

**Supplementary material**

### 1. Supporting methods and analysis workflow

In this paper, we use the “push-pull” theory to determine the variables to map biocultural tourism supply and demand (Figure 1). We comprise our “pull” factors, or tourist attractions, of biodiversity richness and institutionalized cultural richness variables. Our “push” factors, or variables that keep tourists away from a place, are accessibility barriers. To determine Colombia’s biocultural tourism supply, we combine our “pull” factors and the inverse of our “push” factors, which proxy for accessibility.

#### 1.1. Data collection and processing

To map pull factors (tourism supply) and tourism demand, we considered variables that would act as proxies for biodiversity richness, institutionalized cultural richness, accessibility, and demand. Our interdisciplinary team of anthropologists, musicians, biologists, ecologists, and GIS analysts reviewed Colombian government data sources (e.g., DANE- National Statistics Bureau), international data sources (e.g., UNESCO), archives of newspapers and books, and leveraged our own expertise and personal fieldwork spanning decades to gather information for each municipality. Most cultural data exist in narrative form (e.g., music genres), so we tabulated these data in spreadsheets and then mapped them to data points or polygons, which were aggregated by municipality. Some variables were transformed into densities to normalize variable distribution.

We chose municipality (i.e., *municipios*) as our unit of analysis because that is the minimum spatial unit that we could identify for the cultural layers (n=1120 municipalities, median area=1017.8 km^2^), as determined by the national population census that reports national data at the level of municipalities departments (DANE, 2018). The following sections describe how each variable’s data layer was created. To conduct our analysis, we used ArcGIS Pro, QGIS, Rstudio, and Microsoft excel.

##### 1.1.1. Biodiversity variables

Vertebrates and ecosystems were the focus of our biodiversity richness analysis, which was based on the availability of existing data sets and known fauna and serve as an accurate representation of the biodiversity that tourists. We included birds, mammals, amphibians, reptiles, and freshwater fish species. Ecosystem types were included as a proxy for habitat and plant diversity. We excluded invertebrates, plants, fungi, and marine species due to the lack of national range maps.

###### 1.1.1.1. Birds

We produced range maps for 1610 terrestrial birds in Colombia using a combination of expert-driven range maps, ecoregions, elevation, and data from the citizen platform eBird (Sullivan et al., 2009). We started with expert drawn maps from (Ayerbe-Quiñones, 2018) (available in vector format (Vélez et al., 2021). We then created a 20km buffer around the polygon and intersected it with a map of Colombian ecoregions, which is based on combining information from the Terrestrial Ecoregions of the World map (Olson et al., 2001) with a national map of biotic units produced by the Instituto de Investigación de Recursos Biológicos Alexander von Humboldt (Londoño et al., 2015). The buffered expert drawn map was overlapped with the ecoregions map and if 60% of an ecoregion coincides with the range map, that ecoregion becomes part of that species’ distribution. This process results in more natural boundaries to species distribution maps. Following the ecoregion refining, we use elevational data from (Ayerbe-Quiñones, 2018) to select only areas that fell within the species preferred elevation. Resulting range maps were then enhanced by ecoregion and elevational data and better represented species distributions as confirmed by our validation with presence points from eBird which yielded 85% accuracy (% of known presence points that fall within our range maps). We used these individual species maps to produce a bird species richness layer by adding up all ranges for Colombia. We used the ArcGIS Pro tool Zonal Statistics as Table to extract the maximum species richness value per Colombian municipality.

###### 1.1.1.2. Mammals

We used 369 species range maps obtained from two different sources. We included 242 models that follow the “Biomodelos” methodology which relies on an expert elicitation process (Velásquez-Tibatá et al., 2019). In “Biomodelos”, experts validate models that are developed using machine learning algorithms such a Maximum Entropy algorithm (MaxEnt) (Phillips et al., 2006). Maps and statistics of models for primates (Mammalia: Primates, n = 35), Terrestrial carnivores (Mammalia: Carnivora, n = 27), and large rodents (Mammalia: Rodentia, n = 11) are freely available in the Biomodelos database (Velásquez-Tibatá et al., 2019). Models for bats (Mammalia: Chiroptera, n = 169) were developed by (Astorquiza Onofre, 2022) and are available upon request. In addition, we added 127 maps of mammal species that are included in the IUCN database (IUCN, 2023) but that were not considered in the 242 set of models described above. All models were filtered by elevation using information of elevational ranges from the official list of mammals of Colombia (Ramírez-Chaves et al., 2021). Mammal data was provided as a separate raster layer for each species, with a value of 1 within the species’ range, and 0 outside of their range. These layers were overlaid in GIS and summed to create a map of total species richness for each pixel. We then extracted the maximum species richness value within each municipality.

###### 1.1.1.3. Amphibians

Biodiversity Mapping (Jenkins et al., 2013) provides a global raster of amphibian richness, based on 2017 IUCN map data representing native, extant species only. From this, we also extracted the maximum species richness value per municipality, and then we raster spatial resolutions maximums across taxa. The raster provides has a spatial resolution of 10 km^2^ and represents native, extant species only. We clipped the raster to continental Colombia and used the ArcGIS Pro tool Zonal Statistics as Table to extract the maximum species richness value per Colombian municipality.

###### 1.1.1.4. Reptiles

Reptile data was derived from Global Assessment of Reptile Distributions (GARD) version 1.7 (Caetano et al., 2022; Roll et al., 2017). Their dataset provides a distribution area polygon for each species, with 626 species within Colombia. The ArcGIS Pro tool Count Overlapping Features was used to sum all overlapping species area polygons, and the resulting summary vector was converted to raster, whose values represent the total number of reptile species in each pixel. We then extracted the maximum species richness value within each municipality.

###### 1.1.1.5. Freshwater Fishes

We estimated the distribution range for 92% of Colombian freshwater fish species by calculating a minimum spanning tree approach (Grill et al., 2014) among the river reaches with known published occurrences for each species. First, we gathered all the occurrences from the Global Biodiversity Information Facility (GBIF, 2023), AmazonFish (Jézéquel et al., 2020) and FishNet2 (Fishnet2, 2023). Given that the natural distribution of many fish species in Colombia extends beyond the political boundaries, we also included occurrences from transnational basins (i.e., Venezuela, Panama, Peru, Ecuador, Bolivia, and Brazil).

We excluded occurrences missing specific epithets, with missing geographical coordinates, with high coordinate uncertainty (>5000 m), and marine occurrences extending beyond the continental shelf. We validated the taxonomic validity of the remaining occurrences and retrieved the most updated names using FishBase through the “rfishbase” package in R v4.2 (Boettiger et al., 2012). We used the most recent version of the freshwater fish species of Colombia checklist, with updated species distributions from the main hydrogeographic regions of (Amazonas, Orinoco, Caribbean, Pacific, and Magdalena-Cauca) (DoNascimiento et al., 2017), listing 1606 valid species. We excluded unverified species occurrences outside their hydrographic regions, which may include misidentifications or species translocations. A total of 1480 species with at least one occurrence were included in our final database.

We used a simplified river network for the Northern South America basins from RiverATLAS (Linke et al., 2019) sensu HydroRIVERS (Lehner and Grill, 2013), excluding river reaches <50 kilometers upstream. We matched fish occurrences to the nearest river reach in this simplified river network. For each species, we calculated the shortest path among all possible pairs of river reaches where a given fish species is present using the *sfnetwork*, *stplanr* and *hydrocode* R packages (Lovelace and Ellison, 2018). Given that many species may display disjunct distributions across independent rivers (e.g., headwaters), we only allowed calculating paths between river reaches with downstream-upstream connectivity based on Pfafstetter codes provided by HydroBASINS (Lehner and Grill, 2013). The minimum spanning tree is constructed from all the calculated paths, where at least two river reaches are needed. Species ranges were summarized at HydroBASINS level 8 which offers a coarser and more conservative resolution where biases due to inventorying effort and uncertainty in the location of occurrences are limited. Then, we calculated the maximum number of species per municipality.

###### 1.1.1.6. Ecosystems

Colombian government institutions including the meteorology institute, the ministry of environment, and Humboldt institute among others, provide a national map with several different ecosystem classifications for each area polygon (IDEAM et al., 2017). We used data from the field “ECOS_SINTE”, which includes 30 coarse ecosystems - examples include shrub, forest, and savannah. This coarse classification was chosen assuming that tourists are more likely to be interested in a “forest” ecosystem, rather than a particular forest sub-type. GIS tools were used to count the number of different ecosystem types intersecting each Colombian municipality. We counted total diversity per municipality, unlike the species lists because of the coarser resolution of ecosystems vs. species

##### 1.1.2. Institutionalized cultural richness variables

Cultural variables were selected based on data availability of culturally significant sites, areas, and events. These variables represent quantifiable cultural elements that have been registered, recognized, or promoted by national and international institutions. We included UNESCO World Heritage Sites, UNESCO Intangible Cultural Heritage locations, endemic music festivals, Afro-Colombian territories, Indigenous reserves, and museums. We excluded edible plants, ethnobotany, and native languages due to the lack of municipality-level data. We excluded the municipality of San Andrés and Providencia archipelago, due to the lack of data across several variables (mainly biodiversity). We recognize these islands are home to many Raizales, an Afro-Colombian ethnic groups, who hold a vibrant culture. We also acknowledge San Andrés and Providencia as a key touristic area and excluding them is a limitation of our study (Portz et al., 2022).

###### 1.1.2.1. UNESCO World Heritage Sites

There are eight UNESCO World Heritage Sites in Colombia supplied as latitude/longitude points (UNESCO, 2023a). Three of these sites are national parks, so we mapped them to polygons from Sistema Nacional de Áreas Protegidas (UNEP- WCMC and IUCN, 2023). One Site, the “Coffee Cultural Landscape of Colombia” is spread over a large, less-defined area, so we used the national ecosystems map (IDEAM et al., 2017) to locate areas classified as coffee agro-ecosystem. The Spatial Join GIS tool was used to count the total number of World Heritage Sites occurring within each municipality.

###### 1.1.2.2. UNESCO Intangible Cultural Heritage

The UNESCO Intangible Cultural Heritage (ICH) list contains written descriptions of 13 cultural elements within Colombia (UNESCO, 2023b). Some of these include specific location information about towns or departments, which were mapped to existing spatial data (DANE, 2018; IDEAM et al., 2017). Two were related to Indigenous peoples, which were mapped to Indigenous Reserves from the government’s public land database (ANT) (Agenda Nacional de Tierras, 2023a). Several intangible cultural elements are related to music, which were already covered in our music dataset, so we excluded those entries to avoid duplication. Ultimately, 8 out of the 13 elements were included in our analysis. The Spatial Join GIS tool was used to count the total number of ICH Sites occurring within each municipality.

###### 1.1.2.3. Endemic music festivals

A music festival is defined as a musical event that consists of performances of several bands and artists in a limited period of time and limited space. These events allow socializing and reuniting family and friends while expressing the cultural norms of a place. It also allows an intergenerational transfer of culture, which interacts positively with communities of different ethnic backgrounds, and attracts tourists (Aguado et al., 2021).

We only included festivals that celebrate endemic musical genres and excluded festivals that covered international genres (e.g., rock, pop, jazz, salsa). Specifically, coauthor ADA is a Colombian ethnomusicologist and has done extensive fieldwork with local communities as well as archival work in the national library (Biblioteca Luis Angel Arango). Based on his knowledge and expertise, we were able to identify the endemic music genres. In collaboration with local communities across the country, and by reviewing history and ethnomusicology books (Abadía Morales, 1995; Ochoa Escobar, 2013; Ochoa et al., 2014), we were able to identify 105 music festivals spanning over 20 genres (e.g., Bullerengue, Chirimía, Marimba). The number of endemic festivals were mapped as their count per municipality.

###### 1.1.2.4. Afro-Colombian territories

Two spatial layers were combined to represent Afro-Colombian territories: Open data from the national land restitution unit, which aims to repatriate land to Afro-Colombian communities (Unidad de Restitucion de Tierras, 2023), and community council areas from the ANT (Agenda Nacional de Tierras, 2023b). Each polygon in these layers represents an Afro-Colombian territory, and the two datasets have some overlap, with community council data covering more area. Community council was used as a base layer, and polygons were added to it from the land restitution unit data that fall outside of the ANT area. The Spatial Join GIS tool was used to count the total number of Afro-Colombian territories that have an area within each municipality.

###### 1.1.2.5. Indigenous reserves

ANT provides the national dataset Resguardos Indígenas, containing a polygon for each reserve (Agenda Nacional de Tierras, 2023a). The Spatial Join GIS tool was used to count the total number of Indigenous reserves that have area within each municipality.

###### 1.1.2.6. Museums

Data were downloaded from the national museum database in the country (Ministerio de Cultura, 2023), which has a list of all museums registered to the national government (Ministerio de Cultura, 2023). Each museum (n=439) reported on the SIMCO website up to July 2023 was georeferenced, then total number of museums in each municipality were counted and the transformed variable was used in subsequent analyses.

##### 1.1.3. Accessibility variables

Tourism push factors (Figure 1) are defined as variables that would preclude people from visiting a location, such as lack of infrastructure. For this analysis, push factors were reversed to create an accessibility pull factor composite index that would dictate how accessible a municipality was to tourists. Publicly available data on birding lodges, roads, distance to the nearest airport, general lodging, and armed conflict (reversed) were used as a proxy for how peaceful an area was. All variables were evaluated at the municipality level.

###### 1.1.3.1. Bird lodging

Coordinates for bird lodging locations were downloaded from the Birdwatching in Colombia book (Beckers and Flores, 2013), which describes in detail the most accessible birding sites in Colombia, covering 127 sites spread across almost every department of the country. The book covers all lodging locations in Colombia up to 2013 and includes hostels, ecolodges, and camping grounds. To date, this is the most complete data source available for birdwatching in Colombia. Sites outlined in the book were mapped to points, which were later summed to create a count of bird lodges per municipality.

###### 1.1.3.2. Roads

The InVEST Visitation model (Sharp et al., 2020; Wood et al., 2020, 2013) was used to sum the length of roads (DANE, 2018) within each municipality. Road density was calculated for each municipality.

###### 1.1.3.3. Distance to nearest airport

The centroid for each municipality was defined in GIS, then a geodesic distance between each municipality centroid and its nearest airport in the Our Airports database was calculated (Megginson, 2023).

###### 1.1.3.4. Lodging

Lodging location data was taken from Geonames (GeoNames, 2023), as it was found to have the most complete data on lodging sites across Colombia. Its tabular data was imported into GIS and converted to a vector layer. The Geoname field feature co-indicates the general type classification for each point, and points with a value of “HTL” (988 in total) were selected. These were points defined as “a building providing lodging and/or meals for the public”. Lodging sites were summed up to create a count of sites per municipality.

###### 1.1.3.5. Armed conflict data

Armed conflict data is provided by national government data sources (Centro Nacional de Memoria Histórica, 2022), were used to calculate the total number of violent events that happened in each municipality between 1985 and June 30, 2022. Violent events include: 1) military actions, 2) selective assassinations, 3) attacks on towns, 4) terrorist attacks, 5) damage to civilian property, 6) forced disappearances, 7) massacres, 8) antipersonnel mine events, 9) unexploded munitions and improvised explosive devices, 10) recruitment and use of children and adolescents, kidnappings and 11) sexual violence.

This variable was calculated as a density of the total number of violent conflicts divided by the area of each municipality (km^2^), where area data was extracted from the national census (DANE, 2018). The variable was inverted by multiplying it by -1, to penalize violence as a push factor in a destination, hereafter referred to as “Peaceful” (Nikjoo and Ketabi, 2015). This variable was transformed and used in subsequent analyses.

##### 1.1.4. Tourism demand variables

Tourism demand was assessed by evaluating the density and distribution of foreign and national visitors participating in tourism activities throughout Colombia. Proxies that would indicate participation in ecotourism activities such as birdwatching (eBird data), and fishing (sports fisheries) were sought for. Culture-based tourism with photographic data (Flickr), and attendance at music festivals were estimated. Tourism demand for a given municipality was estimated by looking at the arrival of tourists at each airport. Given limited data, datasets span different time ranges, but were all included given the desire to maximize limited data and no reason to limit time range.

###### 1.1.4.1. eBird hotspots

The eBird hotspot data table was accessed from the eBird API 2.0 in September 2022 (Sullivan et al., 2009). ArcGIS Pro tool XY Table to Point was used to display each eBird hotspot at its given latitude and longitude listed in the table and imported into a ArcGIS Pro. The tool Spatial Join was used to count the total number of eBird hotspots within each municipality.

###### 1.1.4.2. Recreational freshwater fisheries

Recreational freshwater fisheries target over 75 fish species (n=69 native, n=5 non-native) in continental Colombia, with growing popularity over recent years (Lasso et al., 2019). The most recent snapshot of recreational fisheries in Colombia from the National Aquaculture and Fisheries Authority (Heinsohn Mallarino, 2023) were used and rasterized to the map of the main sport fisheries zones of Colombia from Heinsohn Mallarino (2023) using QGIS v3.24. Later, using GIMP 2.10.34 data were extracted from a map image on the rivers and lakes where fisheries are documented to occur. This was done by picking all the ares marked in red in the digital image using GIMP, then converted to a geo.tiff. After manually excluding marine recreational fisheries areas along the Pacific and Caribbean coasts, the municipalities where recreational fisheries occur by intersecting with our rasterized layer were identified.

###### 1.1.4.3. Flickr Photo User Days (PUD)

Lacking empirical data on visitation rates to tourism sites, the InVEST Visitation model (Sharp et al., 2020; Wood et al., 2013) was used to estimate visitation from geotagged photos posted to the photo sharing site Flickr. The model calculates the number of “photo user days” (PUD) within each polygon of interest (municipalities). One PUD equals one unique photographer who took at least one photo on a specific day at a particular location. Data are provided for the years 2005-2017, and data from all years were used to calculate the annual average PUD within each municipality.

###### 1.1.4.4. Airport visitors

Yearly airport arrival data were available for years 2004-2019 in the statistics of the national aeronautics website (Aeronáutica Civil, 2019). The median arrival of passengers at each airport was calculated. Airport names from the OurAirports database (Megginson, 2023) were combined with the median passenger arrival and summed the total number of passengers arriving at all airports within each municipality.

###### 1.1.4.5. Music festivals visitors

The number of visitors to music festivals was estimated by looking at national newspapers, event Facebook pages, and blog posts from the municipalities. Local newspapers tend to cover festivals and often report the total number of attendees for a given year. We looked for recent data sources (2017-2023) and found attendance data for 19 festivals. We used actual attendance data for the 19 that had it, and 10% of the town population of the town otherwise. We did so because music festivals are often intended to celebrate the place and the community (Aguado et al., 2021). While this is an imperfect metric, this was used as an estimate of demand to avoid having no data for these festivals. Logarithmic transformations were applied, and the transformed variable was used in subsequent analyses (Table 1).

##### 1.1.5. References

Abadía Morales, G., 1995. ABC del folklore Colombiano. Panamericana Editorial, University of Texas.

Aeronáutica Civil, 2019. Estadísticas operacionales [WWW Document]. URL https://www.aerocivil.gov.co/atencion/estadisticas-de-las-actividades-aeronauticas/estadisticas-operacionales (accessed 10.12.23).

Agenda Nacional de Tierras, 2023a. Resguardos Indígenas- Portal de datos abiertos.

Agenda Nacional de Tierras, 2023b. Consejos comunitarios.

Aguado, L.F., Arbona, A., Palma, L., Heredia-Carroza, J., 2021. How to value a cultural festival? The case of Petronio Álvarez pacific music festival in Colombia. Development Studies Research 8, 309–316. https://doi.org/10.1080/21665095.2021.1979417

Astorquiza Onofre, J.M., 2022. Patrones biogeográficos de diversidad Alfa, Beta y funcional de especies de murciélagos (mammalia, chiroptera) y su representatividad en el sistema nacional de áreas protegidas en Colombia. Universidad de Nariño.

Ayerbe-Quiñones, F., 2018. Guia ilustrada de la avifauna colombiana, First edition. ed. Wildlife Conservation Society, Colombian Programme. PuntoAparte, Bogotá D.C.

Beckers, J., Flores, P., 2013. Birdwatching in Colombia, 1st ed. Buteo Books.

Boettiger, C., Lang, D.T., Wainwright, P.C., 2012. rfishbase: exploring, manipulating and visualizing FishBase data from R. Journal of Fish Biology 81, 2030–2039. https://doi.org/10.1111/j.1095-8649.2012.03464.x

Caetano, G.H.D.O., Chapple, D.G., Grenyer, R., Raz, T., Rosenblatt, J., Tingley, R., Böhm, M., Meiri, S., Roll, U., 2022. Automated assessment reveals that the extinction risk of reptiles is widely underestimated across space and phylogeny. PLoS Biol 20, e3001544. https://doi.org/10.1371/journal.pbio.3001544

Centro Nacional de Memoria Histórica, 2022. El conflicto armado en cifras.

DANE, 2018. Censo nacional de población y vivienda. Estadísticas para grupos étnicos.

DoNascimiento, C., Herrera-Collazos, E.E., Herrera-R, G.A., Ortega-Lara, A., Villa-Navarro, F.A., Oviedo, J.S.U., Maldonado-Ocampo, J.A., 2017. Checklist of the freshwater fishes of Colombia: a Darwin core alternative to the updating problem. ZooKeys 708, 25–138. https://doi.org/10.3897/zookeys.708.13897

Fishnet2, 2023. Fishnet2.

GBIF, 2023. GBIF occurrence download.

GeoNames, 2023. GeoNames.

Grill, G., Ouellet Dallaire, C., Fluet Chouinard, E., Sindorf, N., Lehner, B., 2014. Development of new indicators to evaluate river fragmentation and flow regulation at large scales: A case study for the Mekong River Basin. Ecological Indicators 45, 148–159. https://doi.org/10.1016/j.ecolind.2014.03.026

Heinsohn Mallarino, C.R., 2023. Panorama de la pesca deportiva en Colombia 2022. Autoridad Nacional de Acuicultura y Pesca (AUNAP), Bogotá D.C.

IDEAM, IaVH, Invemar, IGAC, 2017. Mapa de ecosistemas continentales, costeros y marinos de Colombia. Escala 1:100.000.

IUCN, 2023. Mammal range maps. The IUCN Red List of Threatened Species.

Jenkins, C.N., Pimm, S.L., Joppa, L.N., 2013. Global patterns of terrestrial vertebrate diversity and conservation. Proc. Natl. Acad. Sci. U.S.A. 110. https://doi.org/10.1073/pnas.1302251110

Jézéquel, C., Tedesco, P.A., Bigorne, R., Maldonado-Ocampo, J.A., Ortega, H., Hidalgo, M., Martens, K., Torrente-Vilara, G., Zuanon, J., Acosta, A., Agudelo, E., Barrera Maure, S., Bastos, D.A., Bogotá Gregory, J., Cabeceira, F.G., Canto, A.L.C., Carvajal-Vallejos, F.M., Carvalho, L.N., Cella-Ribeiro, A., Covain, R., Donascimiento, C., Dória, C.R.C., Duarte, C., Ferreira, E.J.G., Galuch, A.V., Giarrizzo, T., Leitão, R.P., Lundberg, J.G., Maldonado, M., Mojica, J.I., Montag, L.F.A., Ohara, W.M., Pires, T.H.S., Pouilly, M., Prada-Pedreros, S., de Queiroz, L.J., Rapp Py-Daniel, L., Ribeiro, F.R.V., Ríos Herrera, R., Sarmiento, J., Sousa, L.M., Stegmann, L.F., Valdiviezo-Rivera, J., Villa, F., Yunoki, T., Oberdorff, T., 2020. A database of freshwater fish species of the Amazon Basin. Sci Data 7, 96. https://doi.org/10.1038/s41597-020-0436-4

Lasso, C.A., Heinsohn, C.R., Jensen, S., Morales-Betancourt, M.A., 2019. La pesca deportiva continental en Colombia: guía de las especies de agua dulce. Serie Editorial Recursos Hidrobiológicos y Pesqueros Continentales de Colombia. Instituto de Investigación de Recursos Alexander von Humboldt, Bogotá D.C.

Lehner, B., Grill, G., 2013. Global river hydrography and network routing: baseline data and new approaches to study the world’s large river systems. Hydrological Processes 27, 2171–2186. https://doi.org/10.1002/hyp.9740

Linke, S., Lehner, B., Ouellet Dallaire, C., Ariwi, J., Grill, G., Anand, M., Beames, P., Burchard-Levine, V., Maxwell, S., Moidu, H., Tan, F., Thieme, M., 2019. Global hydro-environmental sub-basin and river reach characteristics at high spatial resolution. Sci Data 6, 283. https://doi.org/10.1038/s41597-019-0300-6

Londoño, M., Olaya, M.H., Bello, C., González, I., Gutiérrez, C., López, D., Velásquez, J., Laboratorio de Biogeografía Aplicada y Bioacústica, Instituto Alexander von Humboldt, 2015. Regiones bióticas de Colombia. Un mapa para cada grupo taxonómico: aves, mamíferos y herpetos y un mapa consenso.

Lovelace, R., Ellison, R., 2018. The R Journal: stplanr: A Package for Transport Planning. The R Journal 10, 7–23. https://doi.org/10.32614/RJ-2018-053

Megginson, D., 2023. OurAirports database.

Ministerio de Cultura, 2023. Sistema de información de museos colombianos [WWW Document]. URL http://simco.museoscolombianos.gov.co/ (accessed 7.12.23).

Nikjoo, A.H., Ketabi, M., 2015. The role of push and pull factors in the way tourists choose their destination. Anatolia 26, 588–597. https://doi.org/10.1080/13032917.2015.1041145

Ochoa Escobar, F., 2013. El libro de las gaitas largas, tradición de los Montes de María. Editorial Pontificia Universidad Javeriana, Bogotá D.C.

Ochoa, J.S., Convers, L., Hernández, O., 2014. Arrullos y currulaos. Editorial Pontificia Universidad Javeriana, Bogotá D.C.

Olson, D.M., Dinerstein, E., Wikramanayake, E.D., Burgess, N.D., Powell, G.V.N., Underwood, E.C., D’amico, J.A., Itoua, I., Strand, H.E., Morrison, J.C., Loucks, C.J., Allnutt, T.F., Ricketts, T.H., Kura, Y., Lamoreux, J.F., Wettengel, W.W., Hedao, P., Kassem, K.R., 2001. Terrestrial Ecoregions of the World: A New Map of Life on Earth: A new global map of terrestrial ecoregions provides an innovative tool for conserving biodiversity. BioScience 51, 933–938. https://doi.org/10.1641/0006-3568(2001)051[0933:TEOTWA]2.0.CO;2

Phillips, S.J., Anderson, R.P., Schapire, R.E., 2006. Maximum entropy modeling of species geographic distributions. Ecological Modelling 190, 231–259. https://doi.org/10.1016/j.ecolmodel.2005.03.026

Portz, L., Manzolli, R.P., Villate-Daza, D.A., Fontán-Bouzas, Á., 2022. Where does marine litter hide? The Providencia and Santa Catalina Island problem, SEAFLOWER Reserve (Colombia). Science of The Total Environment 813, 151878. https://doi.org/10.1016/j.scitotenv.2021.151878

Ramírez-Chaves, H.E., Suárez-Castro, A.F., Rodríguez-Posada, M.E., Zurc, D., Concha Osbahr, D.C., Trujillo, A., Noguera Urbano, E.A., Pantoja Peña, G.E., González Maya, J.F., Pérez Torres, J., Mantilla Meluk, H., López Castañeda, C., Velásquez Valencia, A., Zárate Charry, D., 2021. Mamíferos de Colombia. https://doi.org/10.15472/kl1whs

Roll, U., Feldman, A., Novosolov, M., Allison, A., Bauer, A.M., Bernard, R., Böhm, M., Castro-Herrera, F., Chirio, L., Collen, B., Colli, G.R., Dabool, L., Das, I., Doan, T.M., Grismer, L.L., Hoogmoed, M., Itescu, Y., Kraus, F., LeBreton, M., Lewin, A., Martins, M., Maza, E., Meirte, D., Nagy, Z.T., De C. Nogueira, C., Pauwels, O.S.G., Pincheira-Donoso, D., Powney, G.D., Sindaco, R., Tallowin, O.J.S., Torres-Carvajal, O., Trape, J.-F., Vidan, E., Uetz, P., Wagner, P., Wang, Y., Orme, C.D.L., Grenyer, R., Meiri, S., 2017. The global distribution of tetrapods reveals a need for targeted reptile conservation. Nat Ecol Evol 1, 1677–1682. https://doi.org/10.1038/s41559-017-0332-2

Sharp, R.P., Douglass, J., Wolny, S., Arkema, K.K., Bernhardt, J., Bierbower, W., Chaumont, N., Denu, D., Fisher, D., Glowinski, K., Griffin, R., Guannel, G., Guerry, A.D., Johnson, J.A., Hamel, P., Kennedy, C., Kim, C.K., Lacayo, M., Lonsdorf, E.V., Mandle, L., Rogers, L., Silver, J.M., Toft, J., Verutes, G., Vogl, A.L., Wood, S.A., Wyatt, K., 2020. InVEST 3.9.0.post255+ug.g85c93b8 User’s Guide.

Sullivan, B.L., Wood, C.L., Iliff, M.J., Bonney, R.E., Fink, D., Kelling, S., 2009. eBird: A citizen-based bird observation network in the biological sciences. Biological Conservation 142, 2282–2292. https://doi.org/10.1016/j.biocon.2009.05.006

UNEP- WCMC, IUCN, 2023. Protected planet: The world database on protected areas (WDPA).

UNESCO, 2023a. Colombia - UNESCO World Heritage Convention [WWW Document]. UNESCO World Heritage Centre. URL https://whc.unesco.org/en/statesparties/co

UNESCO, 2023b. UNESCO Intangible Cultural Heritage 1992-2023.

Unidad de Restitucion de Tierras, 2023. Unidad administrativa especial de gestión de restitución de tierras despojadas.

Velásquez-Tibatá, J., Olaya-Rodríguez, M.H., López-Lozano, D., Gutiérrez, C., González, I., Londoño-Murcia, M.C., 2019. BioModelos: A collaborative online system to map species distributions. PLOS ONE 14, e0214522. https://doi.org/10.1371/journal.pone.0214522

Vélez, D., Tamayo, E., Ayerbe-Quiñones, F., Torres, J., Rey, J., Castro-Moreno, C., Ramírez, B., Ochoa-Quintero, J.M., 2021. Distribution of birds in Colombia. Biodiversity Data Journal 9, e59202. https://doi.org/10.3897/BDJ.9.e59202

Wood, S.A., Guerry, A.D., Silver, J.M., Lacayo, M., 2013. Using social media to quantify nature-based tourism and recreation. Scientific Reports 3, 2976. https://doi.org/10.1038/srep02976

Wood, S.A., Winder, S.G., Lia, E.H., White, E.M., Crowley, C.S.L., Milnor, A.A., 2020. Next-generation visitation models using social media to estimate recreation on public lands. Sci Rep 10, 15419. https://doi.org/10.1038/s41598-020-70829-x

#### 1.2. Analysis workflow

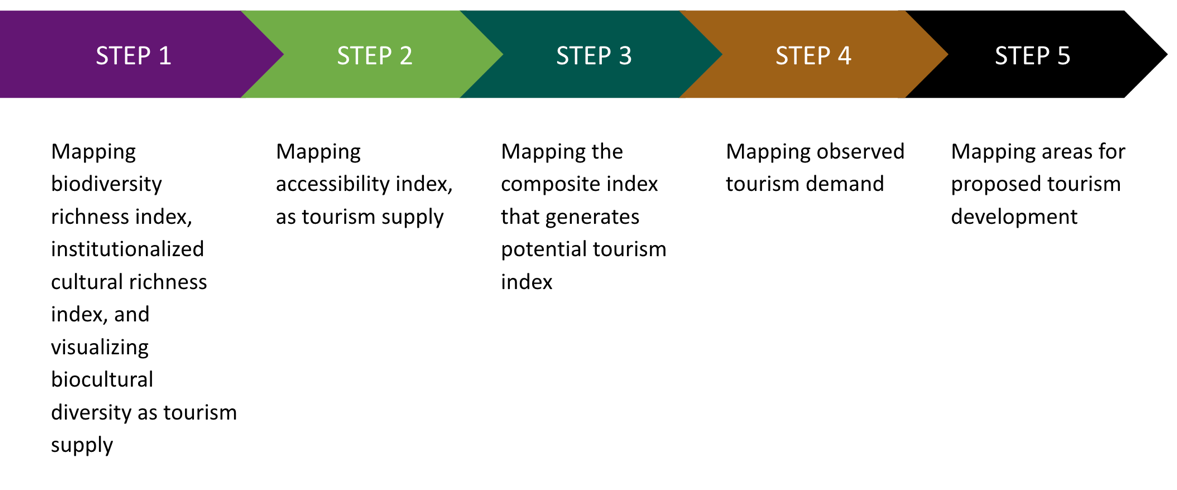

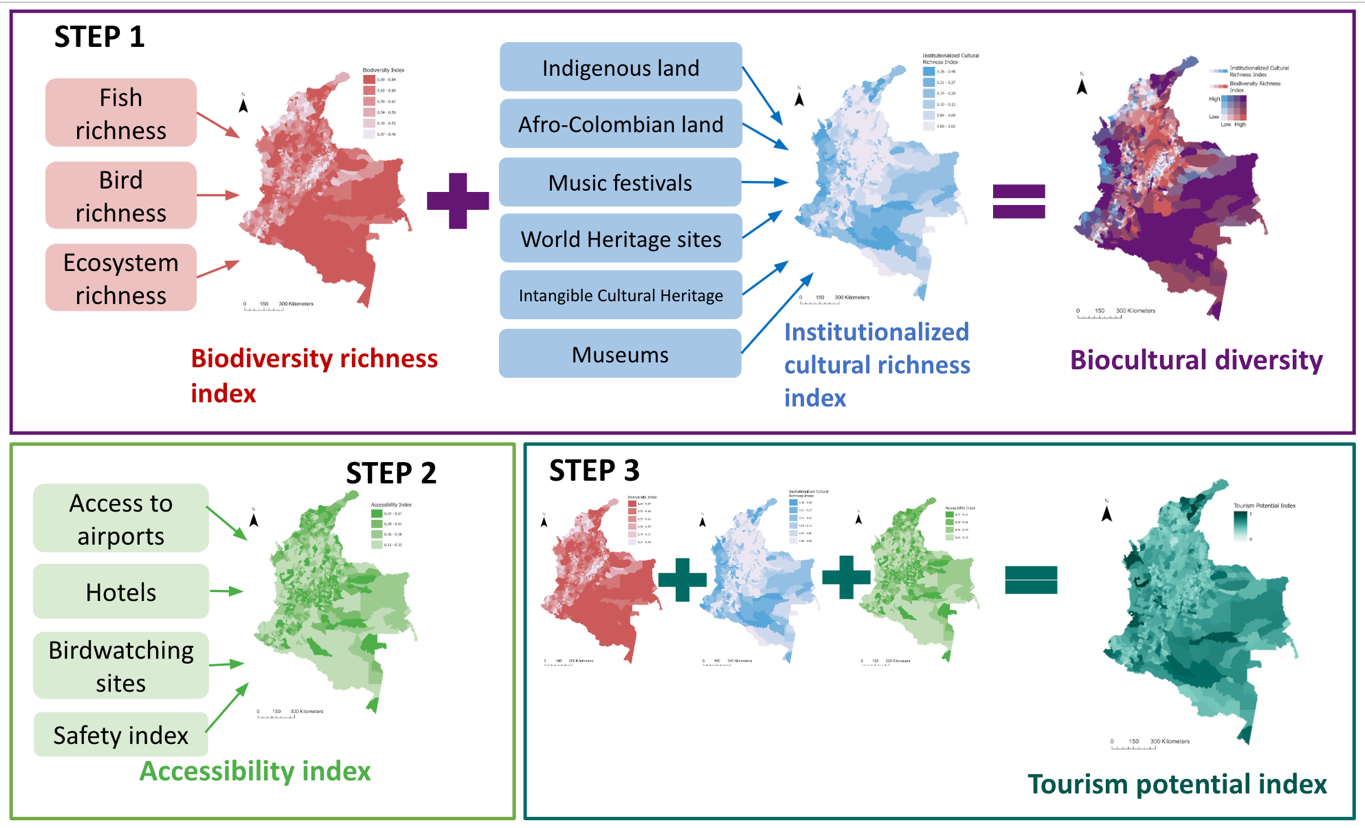

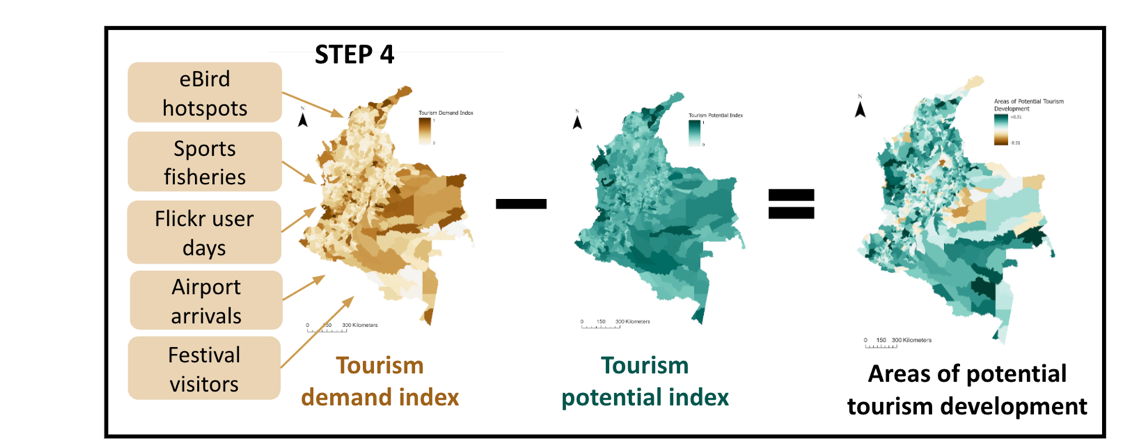

**Figure S1. Analysis workflow for the research project presented here**

### 2. Supporting results

#### 2.1. Correlations at the national scale and regional scales

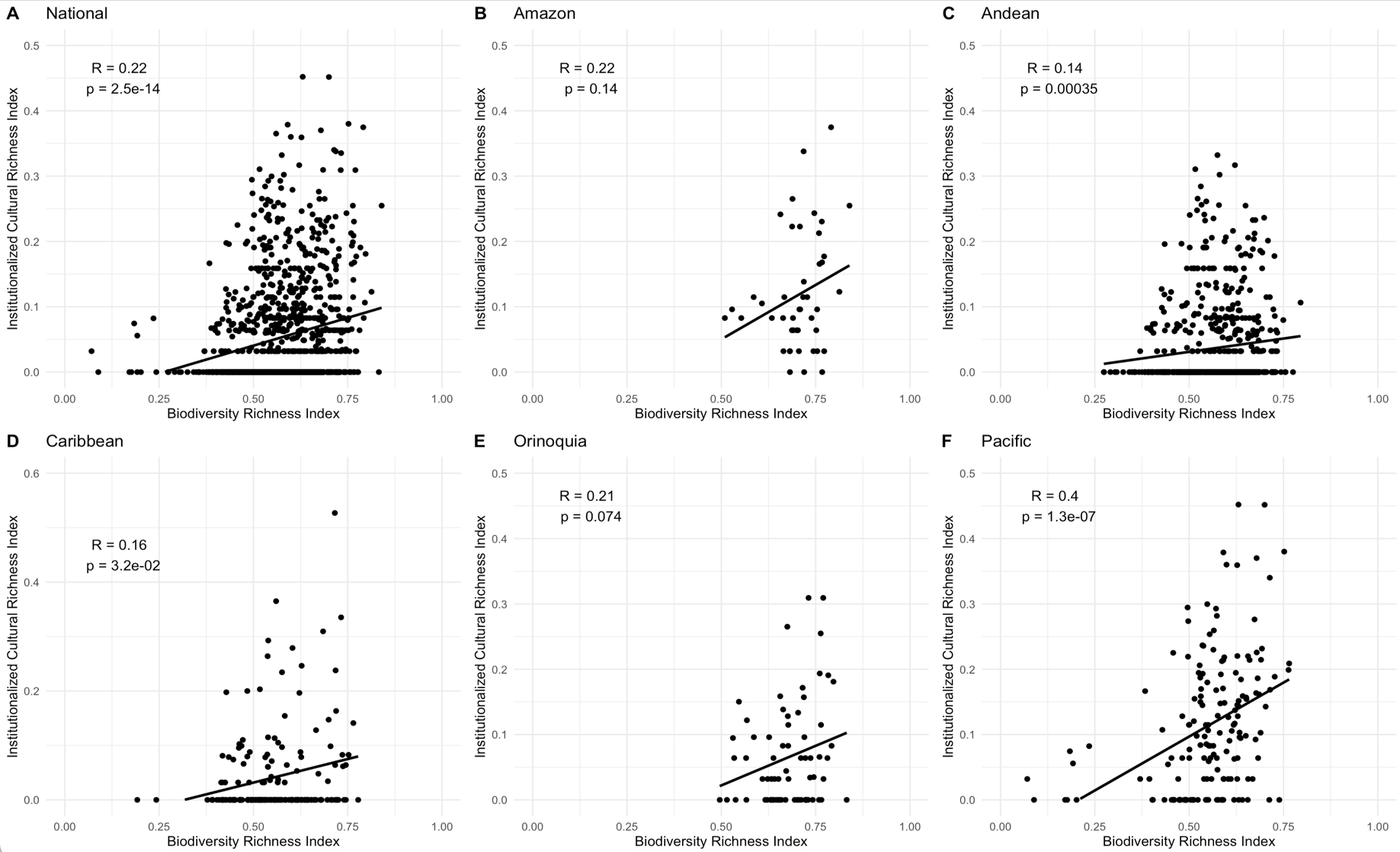

**Figure S2. Correlations between the Biodiversity Richness Index and the Institutionalized Cultural Richness Index at the national scale (A), and at regional scales (B-F).**

#### 2.2. Results by municipality

**Table S1. Results of the composite indices by each municipality**

| **FID** | **Department** | **Municipio** | **Region** | **Biodiversity Index** | **Cultural Index** | **Biodiversity Richness Index Scaled** | **Cultural Diversity Index Scaled** | **Demand Index Scaled** | **Pull Index Scaled** | **Tourism Potential Index Normalized** | **Tourism Demand Index Normalized** | **Proposed areas of Tourism Development** |
| --- | --- | --- | --- | --- | --- | --- | --- | --- | --- | --- | --- | --- |
| 0 | ANTIOQUIA | MEDELLÍN | Andean | 117 | 5 | 0.62 | 0.21 | 0.8 | 0.4 | 0.68 | 0.98 | -0.3 |
| 1 | ANTIOQUIA | ABEJORRAL | Andean | 110 | 0 | 0.49 | 0 | 0.21 | 0.32 | 0.32 | 0.18 | 0.14 |
| 2 | ANTIOQUIA | ABRIAQUÍ | Andean | 107.33 | 0 | 0.52 | 0 | 0.3 | 0.35 | 0.37 | 0.3 | 0.07 |
| 3 | ANTIOQUIA | ALEJANDRÍA | Andean | 133.67 | 0 | 0.61 | 0 | 0.33 | 0.28 | 0.36 | 0.35 | 0.02 |
| 4 | ANTIOQUIA | AMAGÁ | Andean | 114 | 0 | 0.53 | 0 | 0.27 | 0.36 | 0.38 | 0.27 | 0.12 |
| 5 | ANTIOQUIA | AMALFI | Andean | 142 | 0.33 | 0.68 | 0.09 | 0.38 | 0.47 | 0.66 | 0.41 | 0.25 |
| 6 | ANTIOQUIA | ANDES | Andean | 126.5 | 0.5 | 0.63 | 0.07 | 0.42 | 0.37 | 0.52 | 0.46 | 0.05 |
| 7 | ANTIOQUIA | ANGELÓPOLIS | Andean | 105.67 | 0 | 0.46 | 0 | 0.36 | 0.37 | 0.35 | 0.38 | -0.03 |
| 8 | ANTIOQUIA | ANGOSTURA | Andean | 129.17 | 0.33 | 0.58 | 0.1 | 0.2 | 0.31 | 0.46 | 0.17 | 0.29 |
| 9 | ANTIOQUIA | ANORÍ | Andean | 141.83 | 0.5 | 0.68 | 0.07 | 0.31 | 0.44 | 0.61 | 0.31 | 0.3 |
| 10 | ANTIOQUIA | SANTA FÉ DE ANTIOQUIA | Andean | 118.33 | 0.5 | 0.63 | 0.1 | 0.42 | 0.37 | 0.55 | 0.47 | 0.08 |
| 11 | ANTIOQUIA | ANZÁ | Andean | 116.83 | 0 | 0.54 | 0 | 0.31 | 0.33 | 0.37 | 0.32 | 0.05 |
| 12 | ANTIOQUIA | APARTADÓ | Andean | 120.83 | 0.5 | 0.57 | 0.08 | 0.31 | 0.33 | 0.46 | 0.32 | 0.14 |
| 13 | ANTIOQUIA | ARBOLETES | Andean | 83.5 | 0.17 | 0.43 | 0.03 | 0.26 | 0.31 | 0.31 | 0.25 | 0.06 |
| 14 | ANTIOQUIA | ARGELIA | Pacific | 149.5 | 0 | 0.64 | 0 | 0.32 | 0.24 | 0.35 | 0.33 | 0.01 |
| 15 | ANTIOQUIA | ARMENIA | Andean | 118.5 | 0 | 0.54 | 0 | 0.33 | 0.45 | 0.47 | 0.35 | 0.12 |
| 16 | ANTIOQUIA | BARBOSA | Andean | 129.67 | 0 | 0.62 | 0 | 0.25 | 0.32 | 0.41 | 0.24 | 0.17 |
| 17 | ANTIOQUIA | BELMIRA | Andean | 96.5 | 0 | 0.5 | 0 | 0.23 | 0.33 | 0.34 | 0.21 | 0.12 |
| 18 | ANTIOQUIA | BELLO | Andean | 112 | 0.33 | 0.57 | 0.12 | 0.39 | 0.52 | 0.67 | 0.43 | 0.24 |
| 19 | ANTIOQUIA | BETANIA | Andean | 116.67 | 0.33 | 0.6 | 0.13 | 0.23 | 0.36 | 0.55 | 0.2 | 0.35 |
| 20 | ANTIOQUIA | BETULIA | Andean | 150.83 | 0 | 0.66 | 0 | 0.21 | 0.31 | 0.42 | 0.19 | 0.23 |
| 21 | ANTIOQUIA | CIUDAD BOLÍVAR | Andean | 118.83 | 0.17 | 0.61 | 0.03 | 0.34 | 0.34 | 0.44 | 0.36 | 0.08 |
| 22 | ANTIOQUIA | BRICEÑO | Andean | 141.17 | 0 | 0.66 | 0 | 0.29 | 0.49 | 0.58 | 0.29 | 0.29 |
| 23 | ANTIOQUIA | BURITICÁ | Andean | 117.5 | 0.17 | 0.61 | 0.09 | 0.2 | 0.34 | 0.5 | 0.17 | 0.32 |
| 24 | ANTIOQUIA | CÁCERES | Andean | 137.67 | 1 | 0.7 | 0.1 | 0.26 | 0.33 | 0.55 | 0.25 | 0.3 |
| 25 | ANTIOQUIA | CAICEDO | Andean | 117.67 | 0.17 | 0.58 | 0.1 | 0.21 | 0.34 | 0.49 | 0.18 | 0.31 |
| 26 | ANTIOQUIA | CALDAS | Andean | 114 | 0 | 0.51 | 0 | 0.39 | 0.37 | 0.38 | 0.42 | -0.04 |
| 27 | ANTIOQUIA | CAMPAMENTO | Andean | 131 | 0 | 0.61 | 0 | 0.21 | 0.31 | 0.39 | 0.18 | 0.21 |
| 28 | ANTIOQUIA | CAÑASGORDAS | Andean | 115.83 | 0 | 0.54 | 0 | 0.31 | 0.35 | 0.38 | 0.31 | 0.07 |
| 29 | ANTIOQUIA | CARACOLÍ | Andean | 124.5 | 0.17 | 0.58 | 0.03 | 0.41 | 0.35 | 0.43 | 0.45 | -0.02 |
| 30 | ANTIOQUIA | CARAMANTA | Andean | 120 | 0.17 | 0.54 | 0.03 | 0.25 | 0.38 | 0.44 | 0.23 | 0.21 |
| 31 | ANTIOQUIA | CAREPA | Andean | 121.67 | 0.33 | 0.53 | 0.05 | 0.48 | 0.37 | 0.45 | 0.54 | -0.09 |
| 32 | ANTIOQUIA | EL CARMEN DE VIBORAL | Andean | 135.5 | 0.17 | 0.63 | 0.08 | 0.33 | 0.32 | 0.49 | 0.34 | 0.15 |
| 33 | ANTIOQUIA | CAROLINA | Andean | 132.83 | 0 | 0.64 | 0 | 0.33 | 0.32 | 0.42 | 0.34 | 0.08 |
| 34 | ANTIOQUIA | CAUCASIA | Andean | 124.5 | 0 | 0.68 | 0 | 0.33 | 0.35 | 0.46 | 0.35 | 0.12 |
| 35 | ANTIOQUIA | CHIGORODÓ | Andean | 123.5 | 0.5 | 0.57 | 0.08 | 0.38 | 0.35 | 0.48 | 0.4 | 0.07 |
| 36 | ANTIOQUIA | CISNEROS | Andean | 136.83 | 0 | 0.57 | 0 | 0.37 | 0.29 | 0.35 | 0.39 | -0.05 |
| 37 | ANTIOQUIA | COCORNÁ | Andean | 137.33 | 0.17 | 0.66 | 0.1 | 0.37 | 0.31 | 0.51 | 0.39 | 0.11 |
| 38 | ANTIOQUIA | CONCEPCIÓN | Andean | 130.5 | 0 | 0.6 | 0 | 0.33 | 0.47 | 0.52 | 0.34 | 0.18 |
| 39 | ANTIOQUIA | CONCORDIA | Andean | 119.83 | 0 | 0.58 | 0 | 0.24 | 0.31 | 0.37 | 0.22 | 0.16 |
| 40 | ANTIOQUIA | COPACABANA | Andean | 104.33 | 0 | 0.51 | 0 | 0.4 | 0.38 | 0.39 | 0.44 | -0.05 |
| 41 | ANTIOQUIA | DABEIBA | Andean | 154 | 3 | 0.71 | 0.16 | 0.17 | 0.3 | 0.59 | 0.12 | 0.47 |
| 42 | ANTIOQUIA | DONMATÍAS | Andean | 136.5 | 0.17 | 0.68 | 0.1 | 0.24 | 0.34 | 0.55 | 0.22 | 0.33 |
| 43 | ANTIOQUIA | EBÉJICO | Andean | 113.83 | 0 | 0.51 | 0 | 0.23 | 0.31 | 0.33 | 0.2 | 0.13 |
| 44 | ANTIOQUIA | EL BAGRE | Andean | 126 | 1.5 | 0.67 | 0.16 | 0.31 | 0.34 | 0.61 | 0.32 | 0.29 |
| 45 | ANTIOQUIA | ENTRERRÍOS | Andean | 93.67 | 0 | 0.5 | 0 | 0.31 | 0.38 | 0.39 | 0.32 | 0.07 |
| 46 | ANTIOQUIA | ENVIGADO | Andean | 103.5 | 0.67 | 0.53 | 0.14 | 0.42 | 0.57 | 0.7 | 0.47 | 0.24 |
| 47 | ANTIOQUIA | FREDONIA | Andean | 114.67 | 0 | 0.51 | 0 | 0.25 | 0.48 | 0.48 | 0.23 | 0.25 |
| 48 | ANTIOQUIA | FRONTINO | Andean | 143.83 | 1.83 | 0.69 | 0.14 | 0.37 | 0.31 | 0.57 | 0.4 | 0.17 |
| 49 | ANTIOQUIA | GIRALDO | Andean | 115 | 0 | 0.53 | 0 | 0.24 | 0.36 | 0.39 | 0.23 | 0.16 |
| 50 | ANTIOQUIA | GIRARDOTA | Andean | 105.83 | 0 | 0.53 | 0 | 0.36 | 0.35 | 0.38 | 0.39 | -0.01 |
| 51 | ANTIOQUIA | GÓMEZ PLATA | Andean | 139.5 | 0 | 0.65 | 0 | 0.33 | 0.34 | 0.44 | 0.34 | 0.1 |
| 52 | ANTIOQUIA | GRANADA | Andean | 162.67 | 0.33 | 0.67 | 0.11 | 0.23 | 0.29 | 0.52 | 0.21 | 0.31 |
| 53 | ANTIOQUIA | GUADALUPE | Andean | 154.5 | 0 | 0.62 | 0 | 0.34 | 0.47 | 0.54 | 0.35 | 0.19 |
| 54 | ANTIOQUIA | GUARNE | Andean | 97 | 0 | 0.52 | 0 | 0.38 | 0.39 | 0.4 | 0.41 | -0.01 |
| 55 | ANTIOQUIA | GUATAPÉ | Andean | 120.17 | 0.17 | 0.57 | 0.12 | 0.39 | 0.32 | 0.48 | 0.42 | 0.07 |
| 56 | ANTIOQUIA | HELICONIA | Andean | 112.67 | 0 | 0.49 | 0 | 0.34 | 0.35 | 0.35 | 0.35 | 0 |
| 57 | ANTIOQUIA | HISPANIA | Andean | 115.67 | 0 | 0.51 | 0 | 0.27 | 0.38 | 0.39 | 0.26 | 0.13 |
| 58 | ANTIOQUIA | ITAGÜÍ | Andean | 99 | 0.5 | 0.43 | 0.17 | 0.33 | 0.39 | 0.51 | 0.35 | 0.16 |
| 59 | ANTIOQUIA | ITUANGO | Andean | 150.83 | 0.67 | 0.72 | 0.08 | 0.29 | 0.34 | 0.56 | 0.29 | 0.27 |
| 60 | ANTIOQUIA | JARDÍN | Andean | 121.17 | 0.33 | 0.59 | 0.05 | 0.39 | 0.51 | 0.6 | 0.43 | 0.17 |
| 61 | ANTIOQUIA | JERICÓ | Andean | 122 | 0.17 | 0.58 | 0.1 | 0.35 | 0.39 | 0.53 | 0.38 | 0.16 |
| 62 | ANTIOQUIA | LA CEJA | Andean | 110.17 | 0.17 | 0.51 | 0.11 | 0.4 | 0.36 | 0.47 | 0.44 | 0.03 |
| 63 | ANTIOQUIA | LA ESTRELLA | Andean | 98.33 | 0 | 0.49 | 0 | 0.41 | 0.39 | 0.39 | 0.45 | -0.06 |
| 64 | ANTIOQUIA | LA PINTADA | Andean | 109.33 | 0 | 0.5 | 0 | 0.29 | 0.33 | 0.34 | 0.28 | 0.05 |
| 65 | ANTIOQUIA | LA UNIÓN | Caribbean | 125.83 | 0 | 0.56 | 0 | 0.26 | 0.33 | 0.37 | 0.24 | 0.13 |
| 66 | ANTIOQUIA | LIBORINA | Andean | 117.5 | 0 | 0.6 | 0 | 0.31 | 0.35 | 0.42 | 0.32 | 0.1 |
| 67 | ANTIOQUIA | MACEO | Andean | 123.17 | 0 | 0.57 | 0 | 0.32 | 0.33 | 0.39 | 0.33 | 0.06 |
| 68 | ANTIOQUIA | MARINILLA | Andean | 111.67 | 0 | 0.49 | 0 | 0.36 | 0.36 | 0.37 | 0.38 | -0.02 |
| 69 | ANTIOQUIA | MONTEBELLO | Andean | 106.83 | 0 | 0.47 | 0 | 0.24 | 0.34 | 0.33 | 0.23 | 0.11 |
| 70 | ANTIOQUIA | MURINDÓ | Andean | 142 | 2 | 0.67 | 0.19 | 0.26 | 0.25 | 0.55 | 0.26 | 0.3 |
| 71 | ANTIOQUIA | MUTATÁ | Andean | 150.5 | 1 | 0.71 | 0.1 | 0.35 | 0.35 | 0.58 | 0.38 | 0.2 |
| 72 | ANTIOQUIA | NARIÑO | Andean | 149.33 | 0 | 0.66 | 0 | 0.31 | 0.32 | 0.43 | 0.32 | 0.11 |
| 73 | ANTIOQUIA | NECOCLÍ | Andean | 84.5 | 0.67 | 0.49 | 0.16 | 0.56 | 0.35 | 0.51 | 0.65 | -0.14 |
| 74 | ANTIOQUIA | NECHÍ | Andean | 129.83 | 0.17 | 0.68 | 0.03 | 0.18 | 0.31 | 0.46 | 0.14 | 0.32 |
| 75 | ANTIOQUIA | OLAYA | Andean | 117 | 0 | 0.57 | 0 | 0.26 | 0.38 | 0.42 | 0.25 | 0.18 |
| 76 | ANTIOQUIA | PEÑOL | Andean | 119 | 0.17 | 0.57 | 0.11 | 0.27 | 0.33 | 0.49 | 0.27 | 0.22 |
| 77 | ANTIOQUIA | PEQUE | Andean | 117 | 0 | 0.63 | 0 | 0.18 | 0.22 | 0.32 | 0.15 | 0.18 |
| 78 | ANTIOQUIA | PUEBLORRICO | Andean | 122.33 | 0.17 | 0.57 | 0.03 | 0.26 | 0.37 | 0.45 | 0.25 | 0.19 |
| 79 | ANTIOQUIA | PUERTO BERRÍO | Andean | 130.83 | 0.17 | 0.69 | 0.03 | 0.51 | 0.49 | 0.62 | 0.58 | 0.03 |
| 80 | ANTIOQUIA | PUERTO NARE | Andean | 131.17 | 0 | 0.65 | 0 | 0.53 | 0.37 | 0.46 | 0.62 | -0.16 |
| 81 | ANTIOQUIA | PUERTO TRIUNFO | Andean | 130.5 | 0 | 0.59 | 0 | 0.47 | 0.38 | 0.44 | 0.53 | -0.09 |
| 82 | ANTIOQUIA | REMEDIOS | Andean | 125.83 | 0.17 | 0.62 | 0.03 | 0.29 | 0.35 | 0.46 | 0.29 | 0.16 |
| 83 | ANTIOQUIA | RETIRO | Andean | 105.5 | 0 | 0.56 | 0 | 0.38 | 0.38 | 0.42 | 0.41 | 0.01 |
| 84 | ANTIOQUIA | RIONEGRO | Andean | 141 | 0.33 | 0.61 | 0.11 | 0.56 | 0.56 | 0.72 | 0.65 | 0.07 |
| 85 | ANTIOQUIA | SABANALARGA | Andean | 148 | 0.17 | 0.68 | 0.09 | 0.2 | 0.3 | 0.51 | 0.16 | 0.35 |
| 86 | ANTIOQUIA | SABANETA | Andean | 98.83 | 0 | 0.47 | 0 | 0.44 | 0.6 | 0.56 | 0.49 | 0.07 |
| 87 | ANTIOQUIA | SALGAR | Andean | 124 | 0 | 0.61 | 0 | 0.31 | 0.32 | 0.39 | 0.31 | 0.08 |
| 88 | ANTIOQUIA | SAN ANDRÉS DE CUERQUÍA | Andean | 130 | 0 | 0.59 | 0 | 0.22 | 0.36 | 0.42 | 0.2 | 0.22 |
| 89 | ANTIOQUIA | SAN CARLOS | Andean | 136.5 | 0 | 0.68 | 0 | 0.43 | 0.31 | 0.43 | 0.48 | -0.05 |
| 90 | ANTIOQUIA | SAN FRANCISCO | Andean | 157.33 | 0 | 0.67 | 0 | 0.42 | 0.45 | 0.55 | 0.47 | 0.08 |
| 91 | ANTIOQUIA | SAN JERÓNIMO | Andean | 117.83 | 0 | 0.55 | 0 | 0.35 | 0.38 | 0.41 | 0.37 | 0.04 |
| 92 | ANTIOQUIA | SAN JOSÉ DE LA MONTAÑA | Andean | 102.17 | 0 | 0.51 | 0 | 0.33 | 0.35 | 0.37 | 0.34 | 0.03 |
| 93 | ANTIOQUIA | SAN JUAN DE URABÁ | Andean | 81.33 | 0.17 | 0.41 | 0.03 | 0.27 | 0.37 | 0.35 | 0.27 | 0.08 |
| 94 | ANTIOQUIA | SAN LUIS | Andean | 141 | 0 | 0.64 | 0 | 0.41 | 0.32 | 0.41 | 0.45 | -0.04 |
| 95 | ANTIOQUIA | SAN PEDRO DE LOS MILAGROS | Andean | 103.83 | 0.17 | 0.5 | 0.1 | 0.35 | 0.36 | 0.47 | 0.37 | 0.1 |
| 96 | ANTIOQUIA | SAN PEDRO DE URABÁ | Andean | 113.33 | 0.17 | 0.58 | 0.03 | 0.38 | 0.33 | 0.42 | 0.41 | 0.01 |
| 97 | ANTIOQUIA | SAN RAFAEL | Andean | 137.33 | 0 | 0.65 | 0 | 0.34 | 0.29 | 0.4 | 0.36 | 0.04 |
| 98 | ANTIOQUIA | SAN ROQUE | Andean | 139.17 | 0 | 0.68 | 0 | 0.2 | 0.28 | 0.4 | 0.17 | 0.24 |
| 99 | ANTIOQUIA | SAN VICENTE FERRER | Andean | 112.5 | 0.17 | 0.53 | 0.1 | 0.23 | 0.35 | 0.47 | 0.21 | 0.26 |
| 100 | ANTIOQUIA | SANTA BÁRBARA | Andean | 126.83 | 0 | 0.58 | 0 | 0.33 | 0.31 | 0.37 | 0.34 | 0.02 |
| 101 | ANTIOQUIA | SANTA ROSA DE OSOS | Andean | 138.17 | 0.33 | 0.68 | 0.08 | 0.21 | 0.36 | 0.56 | 0.18 | 0.38 |
| 102 | ANTIOQUIA | SANTO DOMINGO | Andean | 138.5 | 0 | 0.65 | 0 | 0.22 | 0.29 | 0.39 | 0.2 | 0.2 |
| 103 | ANTIOQUIA | EL SANTUARIO | Andean | 102.5 | 0.33 | 0.49 | 0.13 | 0.26 | 0.35 | 0.47 | 0.25 | 0.22 |
| 104 | ANTIOQUIA | SEGOVIA | Andean | 122.33 | 0.33 | 0.57 | 0.05 | 0.16 | 0.31 | 0.42 | 0.12 | 0.3 |
| 105 | ANTIOQUIA | SONSÓN | Andean | 149.83 | 1 | 0.73 | 0.1 | 0.33 | 0.3 | 0.55 | 0.35 | 0.2 |
| 106 | ANTIOQUIA | SOPETRÁN | Andean | 119 | 0.17 | 0.57 | 0.03 | 0.34 | 0.37 | 0.44 | 0.36 | 0.08 |
| 107 | ANTIOQUIA | TÁMESIS | Andean | 121.83 | 0.5 | 0.6 | 0.15 | 0.35 | 0.35 | 0.56 | 0.38 | 0.18 |
| 108 | ANTIOQUIA | TARAZÁ | Andean | 142.33 | 0.17 | 0.72 | 0.03 | 0.16 | 0.32 | 0.49 | 0.12 | 0.37 |
| 109 | ANTIOQUIA | TARSO | Andean | 121 | 0 | 0.54 | 0 | 0.34 | 0.38 | 0.41 | 0.35 | 0.05 |
| 110 | ANTIOQUIA | TITIRIBÍ | Andean | 113.67 | 0 | 0.55 | 0 | 0.36 | 0.35 | 0.39 | 0.38 | 0 |
| 111 | ANTIOQUIA | TOLEDO | Andean | 136.83 | 0 | 0.58 | 0 | 0.22 | 0.36 | 0.41 | 0.19 | 0.23 |
| 112 | ANTIOQUIA | TURBO | Andean | 129.17 | 1.67 | 0.62 | 0.29 | 0.51 | 0.44 | 0.79 | 0.59 | 0.2 |
| 113 | ANTIOQUIA | URAMITA | Andean | 134.33 | 0.17 | 0.37 | 0.03 | 0.2 | 0.33 | 0.29 | 0.17 | 0.12 |
| 114 | ANTIOQUIA | URRAO | Andean | 150.33 | 1.67 | 0.73 | 0.16 | 0.39 | 0.45 | 0.73 | 0.43 | 0.3 |
| 115 | ANTIOQUIA | VALDIVIA | Andean | 143.67 | 0 | 0.67 | 0 | 0.31 | 0.33 | 0.44 | 0.32 | 0.12 |
| 116 | ANTIOQUIA | VALPARAÍSO | Amazon | 163.17 | 0.33 | 0.61 | 0.14 | 0.24 | 0.49 | 0.67 | 0.23 | 0.45 |
| 117 | ANTIOQUIA | VEGACHÍ | Andean | 136.83 | 0 | 0.67 | 0 | 0.17 | 0.35 | 0.46 | 0.13 | 0.33 |
| 118 | ANTIOQUIA | VENECIA | Andean | 122.33 | 0.17 | 0.6 | 0.11 | 0.35 | 0.33 | 0.5 | 0.37 | 0.13 |
| 119 | ANTIOQUIA | VIGÍA DEL FUERTE | Andean | 142.5 | 1.5 | 0.69 | 0.15 | 0.28 | 0.27 | 0.55 | 0.28 | 0.26 |
| 120 | ANTIOQUIA | YALÍ | Andean | 122.5 | 0 | 0.57 | 0 | 0.27 | 0.33 | 0.38 | 0.27 | 0.12 |
| 121 | ANTIOQUIA | YARUMAL | Andean | 141.17 | 0.33 | 0.66 | 0.09 | 0.32 | 0.32 | 0.51 | 0.33 | 0.17 |
| 122 | ANTIOQUIA | YOLOMBÓ | Andean | 143.5 | 0.17 | 0.67 | 0.03 | 0.28 | 0.32 | 0.46 | 0.27 | 0.19 |
| 123 | ANTIOQUIA | YONDÓ | Andean | 127.5 | 0.17 | 0.67 | 0.03 | 0.39 | 0.31 | 0.45 | 0.42 | 0.03 |
| 124 | ANTIOQUIA | ZARAGOZA | Andean | 122.83 | 1.5 | 0.63 | 0.17 | 0.26 | 0.37 | 0.61 | 0.25 | 0.36 |
| 125 | ATLÁNTICO | BARRANQUILLA | Caribbean | 91.33 | 2 | 0.54 | 0.26 | 0.62 | 0.44 | 0.71 | 0.73 | -0.03 |
| 126 | ATLÁNTICO | BARANOA | Caribbean | 92 | 0.17 | 0.45 | 0.11 | 0.34 | 0.39 | 0.47 | 0.35 | 0.11 |
| 127 | ATLÁNTICO | CAMPO DE LA CRUZ | Caribbean | 88.17 | 0 | 0.44 | 0 | 0.22 | 0.43 | 0.39 | 0.2 | 0.2 |
| 128 | ATLÁNTICO | CANDELARIA | Caribbean | 104 | 0 | 0.52 | 0 | 0.21 | 0.4 | 0.41 | 0.18 | 0.23 |
| 129 | ATLÁNTICO | GALAPA | Caribbean | 90.33 | 0.5 | 0.43 | 0.2 | 0.49 | 0.41 | 0.55 | 0.56 | 0 |
| 130 | ATLÁNTICO | JUAN DE ACOSTA | Caribbean | 88 | 0.17 | 0.24 | 0.06 | 0.47 | 0.39 | 0.3 | 0.53 | -0.23 |
| 131 | ATLÁNTICO | LURUACO | Caribbean | 90.83 | 0 | 0.51 | 0 | 0.31 | 0.4 | 0.41 | 0.31 | 0.1 |
| 132 | ATLÁNTICO | MALAMBO | Caribbean | 91.33 | 0.17 | 0.48 | 0.11 | 0.34 | 0.42 | 0.52 | 0.36 | 0.16 |
| 133 | ATLÁNTICO | MANATÍ | Caribbean | 95.83 | 0 | 0.56 | 0 | 0.21 | 0.45 | 0.48 | 0.18 | 0.31 |
| 134 | ATLÁNTICO | PALMAR DE VARELA | Caribbean | 92.33 | 0 | 0.47 | 0 | 0.24 | 0.4 | 0.38 | 0.22 | 0.16 |
| 135 | ATLÁNTICO | PIOJÓ | Caribbean | 89.33 | 0 | 0.49 | 0 | 0.33 | 0.41 | 0.41 | 0.35 | 0.06 |
| 136 | ATLÁNTICO | POLONUEVO | Caribbean | 91.67 | 0 | 0.39 | 0 | 0.22 | 0.4 | 0.34 | 0.2 | 0.15 |
| 137 | ATLÁNTICO | PONEDERA | Caribbean | 92.83 | 0 | 0.49 | 0 | 0.21 | 0.39 | 0.38 | 0.18 | 0.2 |
| 138 | ATLÁNTICO | PUERTO COLOMBIA | Caribbean | 150.67 | 0.33 | 0.6 | 0.18 | 0.4 | 0.55 | 0.77 | 0.44 | 0.33 |
| 139 | ATLÁNTICO | REPELÓN | Caribbean | 95.17 | 0.17 | 0.54 | 0.03 | 0.29 | 0.41 | 0.46 | 0.29 | 0.17 |
| 140 | ATLÁNTICO | SABANAGRANDE | Caribbean | 91.33 | 0 | 0.47 | 0 | 0.34 | 0.4 | 0.39 | 0.36 | 0.03 |
| 141 | ATLÁNTICO | SABANALARGA | Caribbean | 148 | 0 | 0.68 | 0 | 0.3 | 0.53 | 0.63 | 0.3 | 0.33 |
| 142 | ATLÁNTICO | SANTA LUCÍA | Caribbean | 93 | 0 | 0.42 | 0 | 0.24 | 0.43 | 0.38 | 0.22 | 0.16 |
| 143 | ATLÁNTICO | SANTO TOMÁS | Caribbean | 92 | 0 | 0.47 | 0 | 0.34 | 0.4 | 0.38 | 0.36 | 0.02 |
| 144 | ATLÁNTICO | SOLEDAD | Caribbean | 90.83 | 0.33 | 0.48 | 0.11 | 0.71 | 0.43 | 0.52 | 0.85 | -0.33 |
| 145 | ATLÁNTICO | SUAN | Caribbean | 89.67 | 0 | 0.43 | 0 | 0.23 | 0.41 | 0.37 | 0.2 | 0.17 |
| 146 | ATLÁNTICO | TUBARÁ | Caribbean | 88.33 | 0 | 0.49 | 0 | 0.25 | 0.41 | 0.41 | 0.24 | 0.17 |
| 147 | ATLÁNTICO | USIACURÍ | Caribbean | 90.17 | 0.17 | 0.43 | 0.11 | 0.34 | 0.4 | 0.47 | 0.35 | 0.12 |
| 148 | BOGOTÁ, D.C. | BOGOTÁ, D.C. | Andean | 88.5 | 11 | 0.66 | 0.14 | 0.61 | 0.5 | 0.72 | 0.73 | -0.01 |
| 149 | BOLÍVAR | CARTAGENA DE INDIAS | Caribbean | 92.33 | 3 | 0.56 | 0.39 | 0.75 | 0.45 | 0.85 | 0.91 | -0.05 |
| 150 | BOLÍVAR | ACHÍ | Caribbean | 119.17 | 0 | 0.6 | 0 | 0.16 | 0.15 | 0.25 | 0.11 | 0.13 |
| 151 | BOLÍVAR | ALTOS DEL ROSARIO | Caribbean | 115 | 0.17 | 0.62 | 0.03 | 0.2 | 0.31 | 0.42 | 0.17 | 0.25 |
| 152 | BOLÍVAR | ARENAL | Caribbean | 127.33 | 0 | 0.65 | 0 | 0.16 | 0.31 | 0.42 | 0.11 | 0.31 |
| 153 | BOLÍVAR | ARJONA | Caribbean | 95 | 0 | 0.58 | 0 | 0.3 | 0.39 | 0.44 | 0.31 | 0.13 |
| 154 | BOLÍVAR | ARROYOHONDO | Caribbean | 96.67 | 0 | 0.56 | 0 | 0.21 | 0.39 | 0.43 | 0.18 | 0.25 |
| 155 | BOLÍVAR | BARRANCO DE LOBA | Caribbean | 115.67 | 0.17 | 0.63 | 0.03 | 0.2 | 0.32 | 0.43 | 0.17 | 0.27 |
| 156 | BOLÍVAR | CALAMAR | Caribbean | 171.17 | 0 | 0.74 | 0 | 0.3 | 0.5 | 0.64 | 0.3 | 0.33 |
| 157 | BOLÍVAR | CANTAGALLO | Caribbean | 130.67 | 0 | 0.66 | 0 | 0.27 | 0.32 | 0.43 | 0.26 | 0.17 |
| 158 | BOLÍVAR | CICUCO | Caribbean | 88.83 | 0 | 0.42 | 0 | 0.24 | 0.42 | 0.38 | 0.23 | 0.15 |
| 159 | BOLÍVAR | CÓRDOBA | Caribbean | 121.5 | 0 | 0.64 | 0 | 0.26 | 0.49 | 0.57 | 0.25 | 0.31 |
| 160 | BOLÍVAR | CLEMENCIA | Caribbean | 89.5 | 0 | 0.4 | 0 | 0.31 | 0.4 | 0.35 | 0.31 | 0.04 |
| 161 | BOLÍVAR | EL CARMEN DE BOLÍVAR | Caribbean | 97.33 | 0.5 | 0.56 | 0.12 | 0.26 | 0.38 | 0.53 | 0.25 | 0.29 |
| 162 | BOLÍVAR | EL GUAMO | Caribbean | 94.67 | 0 | 0.53 | 0 | 0.17 | 0.38 | 0.4 | 0.13 | 0.27 |
| 163 | BOLÍVAR | EL PEÑÓN | Andean | 141.17 | 0 | 0.68 | 0 | 0.2 | 0.34 | 0.46 | 0.16 | 0.3 |
| 164 | BOLÍVAR | MAHATES | Caribbean | 97.17 | 0.33 | 0.58 | 0.14 | 0.34 | 0.4 | 0.58 | 0.36 | 0.21 |
| 165 | BOLÍVAR | MARGARITA | Caribbean | 108.67 | 0 | 0.53 | 0 | 0.17 | 0.38 | 0.4 | 0.13 | 0.27 |
| 166 | BOLÍVAR | MARÍA LA BAJA | Caribbean | 96.67 | 0.5 | 0.58 | 0.11 | 0.4 | 0.4 | 0.55 | 0.44 | 0.11 |
| 167 | BOLÍVAR | MONTECRISTO | Caribbean | 123.83 | 0 | 0.61 | 0 | 0.13 | 0.41 | 0.48 | 0.08 | 0.4 |
| 168 | BOLÍVAR | SANTA CRUZ DE MOMPOX | Caribbean | 109.67 | 0.33 | 0.52 | 0.21 | 0.33 | 0.39 | 0.6 | 0.34 | 0.25 |
| 169 | BOLÍVAR | MORALES | Caribbean | 133.33 | 0 | 0.72 | 0 | 0.26 | 0.3 | 0.45 | 0.24 | 0.2 |
| 170 | BOLÍVAR | NOROSÍ | Caribbean | 130 | 0.33 | 0.62 | 0 | 0.17 | 0.38 | 0.45 | 0.13 | 0.32 |
| 171 | BOLÍVAR | PINILLOS | Caribbean | 112.83 | 0 | 0.58 | 0 | 0.16 | 0.36 | 0.41 | 0.12 | 0.3 |
| 172 | BOLÍVAR | REGIDOR | Caribbean | 93.67 | 0 | 0.54 | 0 | 0.2 | 0.34 | 0.38 | 0.17 | 0.21 |
| 173 | BOLÍVAR | RÍO VIEJO | Caribbean | 133.17 | 0 | 0.72 | 0 | 0.17 | 0.32 | 0.46 | 0.12 | 0.34 |
| 174 | BOLÍVAR | SAN CRISTÓBAL | Caribbean | 93.67 | 0 | 0.46 | 0 | 0.17 | 0.42 | 0.4 | 0.13 | 0.27 |
| 175 | BOLÍVAR | SAN ESTANISLAO | Caribbean | 95.17 | 0.17 | 0.54 | 0.03 | 0.2 | 0.36 | 0.42 | 0.16 | 0.26 |
| 176 | BOLÍVAR | SAN JACINTO | Caribbean | 97.33 | 0.67 | 0.54 | 0.19 | 0.43 | 0.39 | 0.6 | 0.48 | 0.12 |
| 177 | BOLÍVAR | SAN JACINTO DEL CAUCA | Caribbean | 122.17 | 0 | 0.59 | 0 | 0.15 | 0.24 | 0.32 | 0.1 | 0.22 |
| 178 | BOLÍVAR | SAN JUAN NEPOMUCENO | Caribbean | 95.67 | 0.33 | 0.57 | 0.05 | 0.3 | 0.42 | 0.51 | 0.31 | 0.2 |
| 179 | BOLÍVAR | SAN MARTÍN DE LOBA | Caribbean | 120.17 | 0 | 0.64 | 0.06 | 0.31 | 0.31 | 0.46 | 0.32 | 0.14 |
| 180 | BOLÍVAR | SAN PABLO | Caribbean | 130.17 | 0 | 0.68 | 0 | 0.25 | 0.31 | 0.43 | 0.23 | 0.2 |
| 181 | BOLÍVAR | SANTA CATALINA | Caribbean | 90.5 | 0 | 0.51 | 0 | 0.35 | 0.37 | 0.38 | 0.37 | 0.02 |
| 182 | BOLÍVAR | SANTA ROSA | Caribbean | 158.5 | 0 | 0.71 | 0 | 0.32 | 0.38 | 0.51 | 0.33 | 0.18 |
| 183 | BOLÍVAR | SANTA ROSA DEL SUR | Caribbean | 132.83 | 0 | 0.63 | 0 | 0.35 | 0.33 | 0.42 | 0.37 | 0.05 |
| 184 | BOLÍVAR | SIMITÍ | Caribbean | 134.83 | 0 | 0.74 | 0 | 0.21 | 0.35 | 0.51 | 0.19 | 0.32 |
| 185 | BOLÍVAR | SOPLAVIENTO | Caribbean | 94.67 | 0 | 0.5 | 0 | 0.15 | 0.38 | 0.38 | 0.1 | 0.28 |
| 186 | BOLÍVAR | TALAIGUA NUEVO | Caribbean | 88.5 | 0 | 0.42 | 0 | 0.21 | 0.35 | 0.31 | 0.19 | 0.13 |
| 187 | BOLÍVAR | TIQUISIO | Andean | 130.33 | 0 | 0.66 | 0 | 0.15 | 0.32 | 0.42 | 0.11 | 0.32 |
| 188 | BOLÍVAR | TURBACO | Caribbean | 90.17 | 0.17 | 0.42 | 0.1 | 0.34 | 0.43 | 0.47 | 0.36 | 0.11 |
| 189 | BOLÍVAR | TURBANÁ | Caribbean | 90 | 0 | 0.51 | 0 | 0.31 | 0.44 | 0.45 | 0.31 | 0.14 |
| 190 | BOLÍVAR | VILLANUEVA | Caribbean | 178.5 | 0 | 0.69 | 0.06 | 0.35 | 0.3 | 0.47 | 0.37 | 0.1 |
| 191 | BOLÍVAR | ZAMBRANO | Caribbean | 95.5 | 0 | 0.58 | 0 | 0.36 | 0.38 | 0.44 | 0.39 | 0.05 |
| 192 | BOLÍVAR | HATILLO DE LOBA | Caribbean | 92 | 0 | 0.52 | 0 | 0.3 | 0.35 | 0.37 | 0.3 | 0.08 |
| 193 | BOLÍVAR | SAN FERNANDO | Caribbean | 90.67 | 0 | 0.44 | 0 | 0.18 | 0.5 | 0.46 | 0.14 | 0.32 |
| 194 | BOLÍVAR | MAGANGUÉ | Andean | 113.83 | 0 | 0.64 | 0 | 0.39 | 0.38 | 0.47 | 0.42 | 0.05 |
| 195 | BOYACÁ | TUNJA | Andean | 74.17 | 1 | 0.43 | 0.2 | 0.62 | 0.44 | 0.58 | 0.74 | -0.16 |
| 196 | BOYACÁ | ALMEIDA | Andean | 116.5 | 0 | 0.57 | 0 | 0.23 | 0.36 | 0.41 | 0.21 | 0.2 |
| 197 | BOYACÁ | AQUITANIA | Andean | 136.67 | 0 | 0.72 | 0 | 0.42 | 0.5 | 0.63 | 0.46 | 0.17 |
| 198 | BOYACÁ | ARCABUCO | Andean | 83.5 | 0.17 | 0.48 | 0.11 | 0.36 | 0.58 | 0.65 | 0.39 | 0.26 |
| 199 | BOYACÁ | BELÉN | Andean | 94.83 | 0 | 0.49 | 0 | 0.22 | 0.5 | 0.48 | 0.2 | 0.29 |
| 200 | BOYACÁ | BERBEO | Andean | 132.67 | 0 | 0.6 | 0 | 0.22 | 0.37 | 0.44 | 0.19 | 0.24 |
| 201 | BOYACÁ | BETÉITIVA | Andean | 78 | 0 | 0.4 | 0 | 0.2 | 0.38 | 0.33 | 0.17 | 0.16 |
| 202 | BOYACÁ | BOAVITA | Andean | 92 | 0 | 0.48 | 0 | 0.22 | 0.42 | 0.41 | 0.19 | 0.21 |
| 203 | BOYACÁ | BOYACÁ | Andean | 75.67 | 0 | 0.31 | 0 | 0.29 | 0.43 | 0.32 | 0.28 | 0.03 |
| 204 | BOYACÁ | BRICEÑO | Andean | 140.83 | 0 | 0.62 | 0 | 0.16 | 0.41 | 0.48 | 0.11 | 0.37 |
| 205 | BOYACÁ | BUENAVISTA | Andean | 131.67 | 0.17 | 0.59 | 0.06 | 0.33 | 0.49 | 0.59 | 0.34 | 0.24 |
| 206 | BOYACÁ | BUSBANZÁ | Andean | 76 | 0 | 0.33 | 0 | 0.26 | 0.49 | 0.38 | 0.25 | 0.13 |
| 207 | BOYACÁ | CALDAS | Andean | 114 | 0 | 0.51 | 0 | 0.24 | 0.48 | 0.48 | 0.23 | 0.25 |
| 208 | BOYACÁ | CAMPOHERMOSO | Andean | 136.33 | 0 | 0.61 | 0 | 0.18 | 0.24 | 0.33 | 0.14 | 0.18 |
| 209 | BOYACÁ | CERINZA | Andean | 75.67 | 0 | 0.4 | 0 | 0.24 | 0.51 | 0.44 | 0.23 | 0.21 |
| 210 | BOYACÁ | CHINAVITA | Andean | 124.5 | 0 | 0.62 | 0 | 0.22 | 0.35 | 0.43 | 0.2 | 0.23 |
| 211 | BOYACÁ | CHIQUINQUIRÁ | Andean | 86.33 | 0.17 | 0.47 | 0.1 | 0.26 | 0.36 | 0.44 | 0.25 | 0.19 |
| 212 | BOYACÁ | CHITA | Andean | 136 | 0 | 0.73 | 0 | 0.19 | 0.36 | 0.51 | 0.15 | 0.36 |
| 213 | BOYACÁ | CHITARAQUE | Andean | 95.17 | 0 | 0.5 | 0 | 0.22 | 0.33 | 0.34 | 0.2 | 0.14 |
| 214 | BOYACÁ | CHIVATÁ | Andean | 68.67 | 0 | 0.27 | 0 | 0.26 | 0.42 | 0.29 | 0.24 | 0.05 |
| 215 | BOYACÁ | CIÉNEGA | Andean | 95 | 0 | 0.49 | 0 | 0.25 | 0.45 | 0.44 | 0.23 | 0.22 |
| 216 | BOYACÁ | CÓMBITA | Andean | 75 | 0 | 0.47 | 0 | 0.24 | 0.52 | 0.49 | 0.22 | 0.26 |
| 217 | BOYACÁ | COPER | Andean | 128 | 0 | 0.53 | 0 | 0.21 | 0.43 | 0.45 | 0.18 | 0.27 |
| 218 | BOYACÁ | CORRALES | Andean | 76.83 | 0 | 0.38 | 0 | 0.27 | 0.41 | 0.34 | 0.26 | 0.09 |
| 219 | BOYACÁ | LABRANZAGRANDE | Andean | 140.33 | 0 | 0.66 | 0 | 0.27 | 0.36 | 0.47 | 0.26 | 0.21 |
| 220 | BOYACÁ | LA CAPILLA | Andean | 89.33 | 0 | 0.44 | 0 | 0.24 | 0.35 | 0.32 | 0.22 | 0.1 |
| 221 | BOYACÁ | LA VICTORIA | Andean | 157.5 | 0 | 0.58 | 0 | 0.18 | 0.22 | 0.29 | 0.14 | 0.14 |
| 222 | BOYACÁ | LA UVITA | Andean | 91.17 | 0 | 0.53 | 0 | 0.22 | 0.51 | 0.52 | 0.19 | 0.33 |
| 223 | BOYACÁ | PESCA | Andean | 96 | 0 | 0.55 | 0 | 0.32 | 0.42 | 0.45 | 0.32 | 0.13 |
| 224 | BOYACÁ | PISBA | Andean | 137.67 | 0 | 0.66 | 0 | 0.17 | 0.3 | 0.41 | 0.12 | 0.28 |
| 225 | BOYACÁ | PUERTO BOYACÁ | Andean | 147.83 | 0 | 0.67 | 0 | 0.54 | 0.36 | 0.47 | 0.63 | -0.16 |
| 226 | BOYACÁ | QUÍPAMA | Andean | 138.83 | 0 | 0.62 | 0 | 0.3 | 0.34 | 0.42 | 0.3 | 0.12 |
| 227 | BOYACÁ | SOCOTÁ | Andean | 127.67 | 0 | 0.67 | 0 | 0.26 | 0.36 | 0.46 | 0.25 | 0.22 |
| 228 | BOYACÁ | SOCHA | Andean | 95.67 | 0.17 | 0.52 | 0.11 | 0.21 | 0.4 | 0.52 | 0.18 | 0.34 |
| 229 | BOYACÁ | SOGAMOSO | Andean | 96.67 | 0.83 | 0.52 | 0.18 | 0.49 | 0.47 | 0.65 | 0.56 | 0.09 |
| 230 | BOYACÁ | SOMONDOCO | Andean | 115.83 | 0 | 0.52 | 0 | 0.25 | 0.37 | 0.39 | 0.23 | 0.16 |
| 231 | BOYACÁ | ÚMBITA | Andean | 85.5 | 0 | 0.48 | 0 | 0.3 | 0.41 | 0.4 | 0.3 | 0.1 |
| 232 | BOYACÁ | VENTAQUEMADA | Andean | 75.5 | 0.17 | 0.4 | 0.1 | 0.25 | 0.39 | 0.43 | 0.23 | 0.2 |
| 233 | BOYACÁ | VIRACACHÁ | Andean | 93.33 | 0 | 0.47 | 0 | 0.24 | 0.3 | 0.3 | 0.22 | 0.07 |
| 234 | BOYACÁ | ZETAQUIRA | Andean | 127.5 | 0 | 0.63 | 0 | 0.3 | 0.42 | 0.49 | 0.31 | 0.19 |
| 235 | BOYACÁ | COVARACHÍA | Andean | 127.17 | 0 | 0.62 | 0 | 0.22 | 0.33 | 0.41 | 0.2 | 0.21 |
| 236 | BOYACÁ | CUCAITA | Andean | 74 | 0 | 0.39 | 0 | 0.27 | 0.37 | 0.31 | 0.26 | 0.05 |
| 237 | BOYACÁ | CUÍTIVA | Andean | 90.17 | 0.17 | 0.49 | 0.13 | 0.48 | 0.42 | 0.54 | 0.55 | -0.02 |
| 238 | BOYACÁ | CHÍQUIZA | Andean | 78.33 | 0 | 0.46 | 0 | 0.34 | 0.42 | 0.4 | 0.35 | 0.04 |
| 239 | BOYACÁ | CHIVOR | Andean | 122.17 | 0 | 0.61 | 0 | 0.34 | 0.41 | 0.47 | 0.36 | 0.11 |
| 240 | BOYACÁ | DUITAMA | Andean | 82.83 | 0 | 0.52 | 0 | 0.35 | 0.5 | 0.5 | 0.37 | 0.13 |
| 241 | BOYACÁ | EL COCUY | Andean | 73.17 | 0.17 | 0.42 | 0.03 | 0.25 | 0.37 | 0.35 | 0.23 | 0.12 |
| 242 | BOYACÁ | EL ESPINO | Andean | 90.67 | 0 | 0.42 | 0 | 0.24 | 0.37 | 0.33 | 0.22 | 0.11 |
| 243 | BOYACÁ | FIRAVITOBA | Andean | 77.5 | 0 | 0.38 | 0 | 0.24 | 0.56 | 0.47 | 0.22 | 0.25 |
| 244 | BOYACÁ | FLORESTA | Andean | 76.17 | 0.17 | 0.39 | 0.12 | 0.26 | 0.39 | 0.44 | 0.24 | 0.19 |
| 245 | BOYACÁ | GACHANTIVÁ | Andean | 89 | 0 | 0.51 | 0 | 0.36 | 0.43 | 0.43 | 0.39 | 0.04 |
| 246 | BOYACÁ | GÁMEZA | Andean | 92.67 | 0.17 | 0.55 | 0 | 0.21 | 0.31 | 0.36 | 0.18 | 0.18 |
| 247 | BOYACÁ | GARAGOA | Andean | 127.67 | 0 | 0.66 | 0 | 0.32 | 0.4 | 0.5 | 0.33 | 0.16 |
| 248 | BOYACÁ | GUACAMAYAS | Andean | 87.5 | 0 | 0.42 | 0 | 0.25 | 0.48 | 0.43 | 0.24 | 0.19 |
| 249 | BOYACÁ | GUATEQUE | Andean | 110 | 0 | 0.43 | 0 | 0.28 | 0.39 | 0.35 | 0.27 | 0.08 |
| 250 | BOYACÁ | GUAYATÁ | Andean | 95.17 | 0 | 0.49 | 0 | 0.33 | 0.36 | 0.36 | 0.35 | 0.02 |
| 251 | BOYACÁ | IZA | Andean | 91.33 | 0 | 0.46 | 0 | 0.37 | 0.44 | 0.41 | 0.4 | 0.01 |
| 252 | BOYACÁ | JENESANO | Andean | 82.33 | 0 | 0.35 | 0 | 0.27 | 0.45 | 0.36 | 0.26 | 0.11 |
| 253 | BOYACÁ | JERICÓ | Andean | 122 | 0 | 0.58 | 0 | 0.21 | 0.39 | 0.45 | 0.18 | 0.26 |
| 254 | BOYACÁ | VILLA DE LEYVA | Andean | 87.83 | 1.83 | 0.54 | 0.26 | 0.58 | 0.42 | 0.7 | 0.67 | 0.02 |
| 255 | BOYACÁ | MACANAL | Andean | 132.17 | 0 | 0.62 | 0 | 0.22 | 0.4 | 0.47 | 0.2 | 0.28 |
| 256 | BOYACÁ | MARIPÍ | Andean | 129.17 | 0.17 | 0.55 | 0 | 0.21 | 0.46 | 0.48 | 0.18 | 0.31 |
| 257 | BOYACÁ | MIRAFLORES | Andean | 152.67 | 0 | 0.66 | 0 | 0.21 | 0.48 | 0.56 | 0.18 | 0.39 |
| 258 | BOYACÁ | MONGUA | Andean | 129.67 | 0 | 0.67 | 0 | 0.21 | 0.31 | 0.42 | 0.18 | 0.24 |
| 259 | BOYACÁ | MONGUÍ | Andean | 89.83 | 0 | 0.44 | 0 | 0.27 | 0.5 | 0.46 | 0.27 | 0.19 |
| 260 | BOYACÁ | MONIQUIRÁ | Andean | 94.83 | 0 | 0.5 | 0 | 0.23 | 0.44 | 0.44 | 0.21 | 0.23 |
| 261 | BOYACÁ | MOTAVITA | Andean | 66.83 | 0 | 0.36 | 0 | 0.26 | 0.4 | 0.32 | 0.26 | 0.06 |
| 262 | BOYACÁ | MUZO | Andean | 129.17 | 0 | 0.57 | 0 | 0.32 | 0.44 | 0.48 | 0.33 | 0.15 |
| 263 | BOYACÁ | NOBSA | Andean | 77 | 0 | 0.39 | 0 | 0.28 | 0.41 | 0.35 | 0.28 | 0.07 |
| 264 | BOYACÁ | NUEVO COLÓN | Andean | 80.17 | 0 | 0.34 | 0 | 0.25 | 0.41 | 0.32 | 0.24 | 0.08 |
| 265 | BOYACÁ | OICATÁ | Andean | 68.17 | 0 | 0.27 | 0 | 0.26 | 0.37 | 0.24 | 0.24 | -0.01 |
| 266 | BOYACÁ | OTANCHE | Andean | 140 | 0 | 0.62 | 0 | 0.29 | 0.36 | 0.44 | 0.29 | 0.16 |
| 267 | BOYACÁ | PACHAVITA | Andean | 110.5 | 0 | 0.52 | 0 | 0.24 | 0.36 | 0.38 | 0.22 | 0.16 |
| 268 | BOYACÁ | PÁEZ | Andean | 139.67 | 0 | 0.62 | 0 | 0.11 | 0.49 | 0.55 | 0.05 | 0.5 |
| 269 | BOYACÁ | PAIPA | Andean | 89.5 | 0.5 | 0.6 | 0.16 | 0.59 | 0.43 | 0.64 | 0.69 | -0.04 |
| 270 | BOYACÁ | PAJARITO | Andean | 138.67 | 0 | 0.67 | 0 | 0.3 | 0.38 | 0.49 | 0.3 | 0.19 |
| 271 | BOYACÁ | PANQUEBA | Andean | 76.67 | 0 | 0.4 | 0 | 0.27 | 0.39 | 0.34 | 0.26 | 0.08 |
| 272 | BOYACÁ | PAUNA | Andean | 132.17 | 0 | 0.61 | 0 | 0.2 | 0.41 | 0.48 | 0.17 | 0.3 |
| 273 | BOYACÁ | PAYA | Andean | 135.67 | 0 | 0.64 | 0 | 0.17 | 0.35 | 0.44 | 0.13 | 0.32 |
| 274 | BOYACÁ | PAZ DE RÍO | Andean | 83.5 | 0 | 0.48 | 0 | 0.23 | 0.49 | 0.47 | 0.2 | 0.27 |
| 275 | BOYACÁ | RAMIRIQUÍ | Andean | 99.83 | 0 | 0.59 | 0 | 0.24 | 0.44 | 0.49 | 0.23 | 0.26 |
| 276 | BOYACÁ | RÁQUIRA | Andean | 81.67 | 0 | 0.57 | 0 | 0.36 | 0.38 | 0.43 | 0.38 | 0.05 |
| 277 | BOYACÁ | RONDÓN | Andean | 124.17 | 0 | 0.6 | 0 | 0.2 | 0.39 | 0.46 | 0.17 | 0.29 |
| 278 | BOYACÁ | SABOYÁ | Andean | 84.33 | 0 | 0.47 | 0 | 0.22 | 0.39 | 0.37 | 0.19 | 0.18 |
| 279 | BOYACÁ | SÁCHICA | Andean | 79.83 | 0 | 0.44 | 0 | 0.39 | 0.41 | 0.37 | 0.42 | -0.05 |
| 280 | BOYACÁ | SAMACÁ | Andean | 75.67 | 0 | 0.52 | 0 | 0.33 | 0.39 | 0.41 | 0.34 | 0.06 |
| 281 | BOYACÁ | SAN EDUARDO | Andean | 126.5 | 0 | 0.62 | 0 | 0.14 | 0.25 | 0.34 | 0.09 | 0.25 |
| 282 | BOYACÁ | SAN JOSÉ DE PARE | Andean | 95 | 0 | 0.44 | 0 | 0.24 | 0.36 | 0.34 | 0.22 | 0.11 |
| 283 | BOYACÁ | SAN LUIS DE GACENO | Andean | 144.33 | 0 | 0.64 | 0 | 0.19 | 0.36 | 0.45 | 0.15 | 0.3 |
| 284 | BOYACÁ | SAN MATEO | Andean | 92.17 | 0 | 0.45 | 0 | 0.21 | 0.4 | 0.37 | 0.18 | 0.19 |
| 285 | BOYACÁ | SAN MIGUEL DE SEMA | Andean | 78.17 | 0 | 0.42 | 0 | 0.24 | 0.41 | 0.37 | 0.22 | 0.14 |
| 286 | BOYACÁ | SAN PABLO DE BORBUR | Andean | 132.5 | 0 | 0.61 | 0 | 0.18 | 0.35 | 0.43 | 0.15 | 0.28 |
| 287 | BOYACÁ | SANTANA | Andean | 94.67 | 0 | 0.44 | 0 | 0.24 | 0.34 | 0.32 | 0.23 | 0.09 |
| 288 | BOYACÁ | SANTA MARÍA | Andean | 140 | 0 | 0.63 | 0 | 0.34 | 0.41 | 0.49 | 0.36 | 0.13 |
| 289 | BOYACÁ | SANTA ROSA DE VITERBO | Andean | 81.33 | 0.17 | 0.47 | 0.11 | 0.24 | 0.57 | 0.64 | 0.23 | 0.41 |
| 290 | BOYACÁ | SANTA SOFÍA | Andean | 89.5 | 0.17 | 0.53 | 0.12 | 0.26 | 0.36 | 0.5 | 0.25 | 0.24 |
| 291 | BOYACÁ | SATIVANORTE | Andean | 93.33 | 0 | 0.54 | 0 | 0.21 | 0.32 | 0.35 | 0.18 | 0.17 |
| 292 | BOYACÁ | SATIVASUR | Andean | 88 | 0 | 0.46 | 0 | 0.22 | 0.34 | 0.32 | 0.19 | 0.13 |
| 293 | BOYACÁ | SIACHOQUE | Andean | 81 | 0 | 0.39 | 0 | 0.22 | 0.37 | 0.31 | 0.19 | 0.12 |
| 294 | BOYACÁ | SOATÁ | Andean | 96.33 | 0 | 0.58 | 0 | 0.36 | 0.43 | 0.48 | 0.38 | 0.1 |
| 295 | BOYACÁ | SORA | Andean | 73.67 | 0 | 0.39 | 0 | 0.26 | 0.44 | 0.37 | 0.25 | 0.12 |
| 296 | BOYACÁ | SOTAQUIRÁ | Andean | 80.33 | 0 | 0.52 | 0 | 0.22 | 0.52 | 0.52 | 0.19 | 0.33 |
| 297 | BOYACÁ | SORACÁ | Andean | 74 | 0 | 0.3 | 0 | 0.27 | 0.38 | 0.27 | 0.26 | 0.01 |
| 298 | BOYACÁ | SUSACÓN | Andean | 95.83 | 0 | 0.58 | 0 | 0.22 | 0.39 | 0.45 | 0.19 | 0.25 |
| 299 | BOYACÁ | SUTAMARCHÁN | Andean | 86.67 | 0 | 0.49 | 0 | 0.35 | 0.38 | 0.37 | 0.37 | 0 |
| 300 | BOYACÁ | SUTATENZA | Andean | 115.67 | 0 | 0.48 | 0 | 0.27 | 0.37 | 0.37 | 0.26 | 0.11 |
| 301 | BOYACÁ | TASCO | Andean | 85.83 | 0 | 0.54 | 0 | 0.21 | 0.33 | 0.37 | 0.17 | 0.19 |
| 302 | BOYACÁ | TENZA | Andean | 90.33 | 0 | 0.35 | 0 | 0.26 | 0.39 | 0.31 | 0.25 | 0.06 |
| 303 | BOYACÁ | TIBANÁ | Andean | 86.5 | 0 | 0.5 | 0 | 0.24 | 0.41 | 0.41 | 0.22 | 0.19 |
| 304 | BOYACÁ | TIBASOSA | Andean | 77 | 0 | 0.39 | 0 | 0.35 | 0.42 | 0.36 | 0.38 | -0.02 |
| 305 | BOYACÁ | TINJACÁ | Andean | 81.5 | 0 | 0.45 | 0 | 0.26 | 0.41 | 0.38 | 0.25 | 0.13 |
| 306 | BOYACÁ | TIPACOQUE | Andean | 95.83 | 0 | 0.54 | 0 | 0.24 | 0.42 | 0.45 | 0.23 | 0.22 |
| 307 | BOYACÁ | TOCA | Andean | 70.67 | 0 | 0.37 | 0 | 0.32 | 0.42 | 0.34 | 0.34 | 0 |
| 308 | BOYACÁ | TOGÜÍ | Andean | 93.17 | 0 | 0.48 | 0 | 0.34 | 0.28 | 0.28 | 0.36 | -0.08 |
| 309 | BOYACÁ | TÓPAGA | Andean | 76.83 | 0 | 0.38 | 0 | 0.27 | 0.42 | 0.35 | 0.26 | 0.09 |
| 310 | BOYACÁ | TOTA | Andean | 89.33 | 0 | 0.53 | 0 | 0.46 | 0.38 | 0.4 | 0.52 | -0.12 |
| 311 | BOYACÁ | TUNUNGUÁ | Andean | 123.83 | 0 | 0.5 | 0 | 0.18 | 0.25 | 0.27 | 0.14 | 0.13 |
| 312 | BOYACÁ | TURMEQUÉ | Andean | 77.5 | 0 | 0.4 | 0 | 0.24 | 0.44 | 0.38 | 0.23 | 0.16 |
| 313 | BOYACÁ | TUTA | Andean | 74.5 | 0 | 0.44 | 0 | 0.33 | 0.35 | 0.33 | 0.35 | -0.02 |
| 314 | BOYACÁ | TUTAZÁ | Andean | 78.33 | 0 | 0.43 | 0 | 0.22 | 0.39 | 0.35 | 0.2 | 0.16 |
| 315 | BOYACÁ | CHISCAS | Andean | 91.17 | 0.17 | 0.52 | 0.03 | 0.17 | 0.32 | 0.37 | 0.13 | 0.24 |
| 316 | BOYACÁ | CUBARÁ | Andean | 123.67 | 0.5 | 0.66 | 0.07 | 0.24 | 0.47 | 0.63 | 0.22 | 0.41 |
| 317 | BOYACÁ | GÜICÁN DE LA SIERRA | Andean | 90.67 | 0.5 | 0.56 | 0.07 | 0.21 | 0.37 | 0.47 | 0.18 | 0.29 |
| 318 | CALDAS | MANIZALES | Andean | 119.83 | 1 | 0.56 | 0.3 | 0.75 | 0.57 | 0.87 | 0.91 | -0.04 |
| 319 | CALDAS | AGUADAS | Andean | 119 | 0.5 | 0.56 | 0.27 | 0.46 | 0.34 | 0.64 | 0.52 | 0.13 |
| 320 | CALDAS | NEIRA | Andean | 120.17 | 0.17 | 0.57 | 0.13 | 0.32 | 0.34 | 0.51 | 0.34 | 0.17 |
| 321 | CALDAS | MARULANDA | Andean | 137.33 | 0 | 0.61 | 0 | 0.32 | 0.35 | 0.43 | 0.33 | 0.09 |
| 322 | CALDAS | MARQUETALIA | Andean | 147.83 | 0.17 | 0.61 | 0.13 | 0.45 | 0.37 | 0.57 | 0.51 | 0.06 |
| 323 | CALDAS | MARMATO | Andean | 118.17 | 0.33 | 0.54 | 0.13 | 0.27 | 0.39 | 0.54 | 0.27 | 0.28 |
| 324 | CALDAS | MANZANARES | Andean | 142.67 | 0.17 | 0.62 | 0.13 | 0.33 | 0.37 | 0.57 | 0.34 | 0.23 |
| 325 | CALDAS | LA MERCED | Andean | 115.5 | 0.17 | 0.51 | 0.13 | 0.24 | 0.37 | 0.51 | 0.23 | 0.28 |
| 326 | CALDAS | LA DORADA | Andean | 135.17 | 0.17 | 0.62 | 0.08 | 0.44 | 0.38 | 0.53 | 0.5 | 0.03 |
| 327 | CALDAS | FILADELFIA | Andean | 115.83 | 0.17 | 0.52 | 0.13 | 0.21 | 0.36 | 0.5 | 0.19 | 0.32 |
| 328 | CALDAS | CHINCHINÁ | Andean | 115.5 | 0.17 | 0.54 | 0.13 | 0.39 | 0.4 | 0.55 | 0.42 | 0.13 |
| 329 | CALDAS | BELALCÁZAR | Andean | 118.33 | 0.33 | 0.54 | 0.16 | 0.35 | 0.37 | 0.55 | 0.37 | 0.18 |
| 330 | CALDAS | ARANZAZU | Andean | 116.17 | 0.67 | 0.54 | 0.26 | 0.22 | 0.37 | 0.65 | 0.2 | 0.45 |
| 331 | CALDAS | ANSERMA | Andean | 120.67 | 0.17 | 0.56 | 0.13 | 0.34 | 0.39 | 0.56 | 0.35 | 0.2 |
| 332 | CALDAS | NORCASIA | Andean | 135.83 | 0 | 0.56 | 0 | 0.43 | 0.35 | 0.39 | 0.48 | -0.09 |
| 333 | CALDAS | VITERBO | Andean | 110.33 | 0.17 | 0.51 | 0.03 | 0.35 | 0.39 | 0.42 | 0.38 | 0.05 |
| 334 | CALDAS | VILLAMARÍA | Andean | 118.67 | 0.17 | 0.69 | 0.19 | 0.51 | 0.54 | 0.82 | 0.59 | 0.23 |
| 335 | CALDAS | VICTORIA | Andean | 147 | 0.17 | 0.65 | 0.13 | 0.44 | 0.39 | 0.61 | 0.49 | 0.12 |
| 336 | CALDAS | SUPÍA | Andean | 119.67 | 0.5 | 0.54 | 0.27 | 0.24 | 0.36 | 0.64 | 0.22 | 0.42 |
| 337 | CALDAS | SAN JOSÉ | Andean | 118 | 0.33 | 0.5 | 0.16 | 0.35 | 0.36 | 0.52 | 0.37 | 0.15 |
| 338 | CALDAS | SAMANÁ | Andean | 153 | 0 | 0.68 | 0 | 0.4 | 0.34 | 0.46 | 0.44 | 0.02 |
| 339 | CALDAS | SALAMINA | Andean | 124.83 | 0.17 | 0.61 | 0.13 | 0.22 | 0.34 | 0.54 | 0.2 | 0.34 |
| 340 | CALDAS | RISARALDA | Andean | 119 | 0.33 | 0.54 | 0.16 | 0.25 | 0.38 | 0.56 | 0.23 | 0.33 |
| 341 | CALDAS | PENSILVANIA | Andean | 147.33 | 0.33 | 0.68 | 0.21 | 0.5 | 0.36 | 0.67 | 0.57 | 0.1 |
| 342 | CALDAS | PALESTINA | Andean | 115.67 | 0.17 | 0.52 | 0.13 | 0.44 | 0.51 | 0.64 | 0.49 | 0.15 |
| 343 | CALDAS | PÁCORA | Andean | 118.17 | 0.33 | 0.6 | 0.23 | 0.22 | 0.38 | 0.66 | 0.19 | 0.47 |
| 344 | CALDAS | RIOSUCIO | Andean | 141.83 | 1.17 | 0.58 | 0.35 | 0.48 | 0.35 | 0.73 | 0.55 | 0.18 |
| 345 | CAQUETÁ | FLORENCIA | Amazon | 184.67 | 1 | 0.81 | 0.24 | 0.63 | 0.33 | 0.76 | 0.74 | 0.01 |
| 346 | CAQUETÁ | ALBANIA | Andean | 161.83 | 0.17 | 0.72 | 0.03 | 0.16 | 0.3 | 0.47 | 0.12 | 0.35 |
| 347 | CAQUETÁ | BELÉN DE LOS ANDAQUÍES | Amazon | 171.33 | 0.5 | 0.74 | 0.07 | 0.3 | 0.33 | 0.55 | 0.31 | 0.24 |
| 348 | CAQUETÁ | EL DONCELLO | Amazon | 161.83 | 0 | 0.72 | 0 | 0.37 | 0.23 | 0.38 | 0.39 | -0.01 |
| 349 | CAQUETÁ | EL PAUJÍL | Amazon | 183.67 | 0 | 0.77 | 0 | 0.16 | 0.24 | 0.42 | 0.11 | 0.3 |
| 350 | CAQUETÁ | LA MONTAÑITA | Amazon | 186.33 | 0.17 | 0.77 | 0.03 | 0.26 | 0.36 | 0.56 | 0.25 | 0.31 |
| 351 | CAQUETÁ | MILÁN | Amazon | 164 | 1.67 | 0.72 | 0.12 | 0.34 | 0.31 | 0.57 | 0.36 | 0.21 |
| 352 | CAQUETÁ | MORELIA | Amazon | 182.5 | 0.33 | 0.75 | 0.08 | 0.41 | 0.3 | 0.54 | 0.46 | 0.08 |
| 353 | CAQUETÁ | SAN JOSÉ DEL FRAGUA | Amazon | 169.5 | 1 | 0.72 | 0.1 | 0.27 | 0.31 | 0.54 | 0.26 | 0.28 |
| 354 | CAQUETÁ | VALPARAÍSO | Andean | 163.83 | 0.17 | 0.68 | 0.03 | 0.22 | 0.35 | 0.49 | 0.2 | 0.29 |
| 355 | CAQUETÁ | CARTAGENA DEL CHAIRÁ | Amazon | 147.5 | 0.5 | 0.71 | 0.24 | 0.47 | 0.47 | 0.81 | 0.53 | 0.28 |
| 356 | CAQUETÁ | SAN VICENTE DEL CAGUÁN | Amazon | 187.33 | 0.83 | 0.84 | 0.21 | 0.41 | 0.26 | 0.68 | 0.45 | 0.23 |
| 357 | CAQUETÁ | SOLANO | Amazon | 193.33 | 4 | 0.79 | 0.31 | 0.46 | 0.29 | 0.78 | 0.52 | 0.26 |
| 358 | CAQUETÁ | SOLITA | Amazon | 164.67 | 0.17 | 0.68 | 0.03 | 0.27 | 0.25 | 0.4 | 0.27 | 0.14 |
| 359 | CAQUETÁ | CURILLO | Amazon | 151.17 | 0 | 0.68 | 0 | 0.28 | 0.3 | 0.42 | 0.27 | 0.15 |
| 360 | CAQUETÁ | PUERTO RICO | Amazon | 190.17 | 0.83 | 0.8 | 0.09 | 0.43 | 0.27 | 0.55 | 0.48 | 0.08 |
| 361 | CAUCA | POPAYÁN | Pacific | 104.17 | 2.5 | 0.55 | 0.27 | 0.55 | 0.41 | 0.69 | 0.64 | 0.05 |
| 362 | CAUCA | ALMAGUER | Pacific | 93.17 | 0 | 0.5 | 0 | 0.21 | 0.37 | 0.37 | 0.18 | 0.19 |
| 363 | CAUCA | ARGELIA | Andean | 149.5 | 0.83 | 0.64 | 0.15 | 0.16 | 0.44 | 0.66 | 0.12 | 0.55 |
| 364 | CAUCA | BALBOA | Pacific | 120.5 | 0.33 | 0.65 | 0.05 | 0.3 | 0.34 | 0.48 | 0.3 | 0.18 |
| 365 | CAUCA | BOLÍVAR | Andean | 151.83 | 0.17 | 0.73 | 0.03 | 0.18 | 0.49 | 0.64 | 0.14 | 0.5 |
| 366 | CAUCA | BUENOS AIRES | Pacific | 120.33 | 1.33 | 0.59 | 0.16 | 0.29 | 0.33 | 0.54 | 0.29 | 0.25 |
| 367 | CAUCA | CAJIBÍO | Pacific | 107.83 | 0.67 | 0.51 | 0.1 | 0.3 | 0.32 | 0.43 | 0.3 | 0.13 |
| 368 | CAUCA | CALDONO | Pacific | 94.67 | 1.33 | 0.48 | 0.11 | 0.21 | 0.33 | 0.43 | 0.18 | 0.26 |
| 369 | CAUCA | CALOTO | Pacific | 102.5 | 1 | 0.53 | 0.13 | 0.44 | 0.39 | 0.54 | 0.49 | 0.05 |
| 370 | CAUCA | CORINTO | Pacific | 97.5 | 0.33 | 0.55 | 0.12 | 0.3 | 0.35 | 0.5 | 0.3 | 0.2 |
| 371 | CAUCA | FLORENCIA | Andean | 184 | 0.33 | 0.74 | 0.11 | 0.46 | 0.4 | 0.65 | 0.53 | 0.12 |
| 372 | CAUCA | GUACHENÉ | Pacific | 98.83 | 0.5 | 0.48 | 0.08 | 0.33 | 0.48 | 0.54 | 0.34 | 0.2 |
| 373 | CAUCA | INZÁ | Pacific | 104.5 | 1.5 | 0.59 | 0.31 | 0.29 | 0.39 | 0.75 | 0.29 | 0.45 |
| 374 | CAUCA | JAMBALÓ | Pacific | 95 | 0.33 | 0.5 | 0.05 | 0.22 | 0.38 | 0.43 | 0.19 | 0.24 |
| 375 | CAUCA | LA SIERRA | Pacific | 95.83 | 1 | 0.53 | 0.2 | 0.22 | 0.46 | 0.66 | 0.19 | 0.47 |
| 376 | CAUCA | LA VEGA | Pacific | 130.33 | 0.17 | 0.66 | 0.03 | 0.2 | 0.35 | 0.48 | 0.17 | 0.31 |
| 377 | CAUCA | MERCADERES | Pacific | 91.5 | 0.17 | 0.52 | 0.03 | 0.29 | 0.35 | 0.4 | 0.28 | 0.12 |
| 378 | CAUCA | MORALES | Pacific | 132.5 | 1.67 | 0.63 | 0.16 | 0.19 | 0.34 | 0.57 | 0.16 | 0.42 |
| 379 | CAUCA | PADILLA | Pacific | 91.5 | 0.17 | 0.39 | 0.03 | 0.33 | 0.39 | 0.36 | 0.34 | 0.02 |
| 380 | CAUCA | PÁEZ | Andean | 141 | 1.33 | 0.76 | 0.15 | 0.17 | 0.32 | 0.63 | 0.13 | 0.5 |
| 381 | CAUCA | PATÍA | Pacific | 108.33 | 0.83 | 0.62 | 0.13 | 0.53 | 0.35 | 0.56 | 0.61 | -0.05 |
| 382 | CAUCA | PIAMONTE | Pacific | 165.67 | 1.83 | 0.7 | 0.12 | 0.28 | 0.29 | 0.54 | 0.27 | 0.27 |
| 383 | CAUCA | PIENDAMÓ - TUNÍA | Pacific | 103.5 | 1 | 0.5 | 0.1 | 0.33 | 0.41 | 0.51 | 0.35 | 0.16 |
| 384 | CAUCA | PUERTO TEJADA | Pacific | 98.17 | 0 | 0.48 | 0 | 0.44 | 0.38 | 0.37 | 0.49 | -0.12 |
| 385 | CAUCA | PURACÉ | Pacific | 94 | 1 | 0.62 | 0.1 | 0.34 | 0.36 | 0.53 | 0.36 | 0.17 |
| 386 | CAUCA | ROSAS | Pacific | 95 | 0.17 | 0.47 | 0.03 | 0.22 | 0.35 | 0.37 | 0.19 | 0.17 |
| 387 | CAUCA | SAN SEBASTIÁN | Pacific | 95 | 0.17 | 0.56 | 0.03 | 0.19 | 0.31 | 0.38 | 0.15 | 0.23 |
| 388 | CAUCA | SANTANDER DE QUILICHAO | Pacific | 102.33 | 1.33 | 0.56 | 0.18 | 0.58 | 0.38 | 0.59 | 0.68 | -0.09 |
| 389 | CAUCA | SANTA ROSA | Pacific | 158.67 | 1.5 | 0.73 | 0.13 | 0.26 | 0.31 | 0.59 | 0.25 | 0.34 |
| 390 | CAUCA | SILVIA | Pacific | 93 | 1 | 0.55 | 0.1 | 0.32 | 0.33 | 0.46 | 0.32 | 0.14 |
| 391 | CAUCA | SOTARÁ PAISPAMBA | Pacific | 96.83 | 0.83 | 0.55 | 0.09 | 0.32 | 0.33 | 0.46 | 0.33 | 0.13 |
| 392 | CAUCA | SUÁREZ | Pacific | 124.83 | 1 | 0.59 | 0.11 | 0.29 | 0.36 | 0.53 | 0.29 | 0.24 |
| 393 | CAUCA | SUCRE | Andean | 138.17 | 0 | 0.66 | 0 | 0.21 | 0.49 | 0.58 | 0.18 | 0.39 |
| 394 | CAUCA | TIMBÍO | Pacific | 95 | 0.33 | 0.45 | 0.05 | 0.33 | 0.36 | 0.39 | 0.35 | 0.04 |
| 395 | CAUCA | TORIBÍO | Pacific | 96.33 | 0.33 | 0.54 | 0.05 | 0.37 | 0.37 | 0.45 | 0.4 | 0.05 |
| 396 | CAUCA | TOTORÓ | Pacific | 93.33 | 1 | 0.54 | 0.1 | 0.31 | 0.31 | 0.44 | 0.32 | 0.12 |
| 397 | CAUCA | EL TAMBO | Pacific | 128.17 | 0.83 | 0.66 | 0.1 | 0.37 | 0.52 | 0.7 | 0.39 | 0.31 |
| 398 | CAUCA | GUAPI | Pacific | 123 | 2 | 0.62 | 0.14 | 0.31 | 0.31 | 0.53 | 0.31 | 0.21 |
| 399 | CAUCA | LÓPEZ DE MICAY | Pacific | 133.83 | 2.5 | 0.66 | 0.21 | 0.33 | 0.31 | 0.62 | 0.34 | 0.28 |
| 400 | CAUCA | TIMBIQUÍ | Pacific | 124 | 2.83 | 0.55 | 0.27 | 0.53 | 0.27 | 0.58 | 0.62 | -0.04 |
| 401 | CAUCA | VILLA RICA | Pacific | 97.17 | 0 | 0.48 | 0 | 0.35 | 0.4 | 0.39 | 0.37 | 0.02 |
| 402 | CAUCA | MIRANDA | Pacific | 110.83 | 0.17 | 0.59 | 0.03 | 0.31 | 0.41 | 0.5 | 0.31 | 0.18 |
| 403 | CESAR | ASTREA | Caribbean | 90.83 | 0 | 0.51 | 0 | 0.18 | 0.45 | 0.46 | 0.14 | 0.31 |
| 404 | CESAR | BOSCONIA | Caribbean | 89.33 | 0 | 0.52 | 0 | 0.28 | 0.37 | 0.39 | 0.28 | 0.11 |
| 405 | CESAR | CHIMICHAGUA | Caribbean | 122.33 | 0 | 0.7 | 0 | 0.24 | 0.38 | 0.5 | 0.23 | 0.27 |
| 406 | CESAR | CHIRIGUANÁ | Caribbean | 122.83 | 0 | 0.72 | 0 | 0.17 | 0.34 | 0.48 | 0.13 | 0.35 |
| 407 | CESAR | CURUMANÍ | Caribbean | 121.33 | 0.17 | 0.7 | 0.07 | 0.18 | 0.38 | 0.57 | 0.14 | 0.44 |
| 408 | CESAR | EL COPEY | Caribbean | 110 | 0.17 | 0.56 | 0.03 | 0.24 | 0.39 | 0.45 | 0.23 | 0.23 |
| 409 | CESAR | EL PASO | Caribbean | 91.67 | 0 | 0.52 | 0 | 0.41 | 0.38 | 0.39 | 0.45 | -0.05 |
| 410 | CESAR | GAMARRA | Andean | 113.5 | 0 | 0.62 | 0 | 0.28 | 0.37 | 0.45 | 0.28 | 0.17 |
| 411 | CESAR | GONZÁLEZ | Caribbean | 101.33 | 0 | 0.49 | 0 | 0.15 | 0.33 | 0.33 | 0.11 | 0.22 |
| 412 | CESAR | LA GLORIA | Caribbean | 117.33 | 0 | 0.67 | 0 | 0.35 | 0.37 | 0.47 | 0.37 | 0.1 |
| 413 | CESAR | LA JAGUA DE IBIRICO | Caribbean | 131.17 | 0.17 | 0.74 | 0.03 | 0.4 | 0.38 | 0.56 | 0.43 | 0.12 |
| 414 | CESAR | MANAURE BALCÓN DEL CESAR | Caribbean | 112 | 0.17 | 0.59 | 0.11 | 0.36 | 0.41 | 0.57 | 0.38 | 0.18 |
| 415 | CESAR | PAILITAS | Caribbean | 117.33 | 0 | 0.65 | 0 | 0.35 | 0.37 | 0.47 | 0.37 | 0.1 |
| 416 | CESAR | PELAYA | Caribbean | 114.67 | 0 | 0.6 | 0 | 0.27 | 0.41 | 0.47 | 0.27 | 0.2 |
| 417 | CESAR | RÍO DE ORO | Caribbean | 131.67 | 0.17 | 0.63 | 0 | 0.19 | 0.39 | 0.47 | 0.15 | 0.32 |
| 418 | CESAR | SAN ALBERTO | Caribbean | 133 | 0 | 0.64 | 0 | 0.29 | 0.5 | 0.57 | 0.29 | 0.28 |
| 419 | CESAR | SAN DIEGO | Caribbean | 115 | 0 | 0.6 | 0 | 0.23 | 0.35 | 0.42 | 0.21 | 0.22 |
| 420 | CESAR | TAMALAMEQUE | Caribbean | 93 | 0 | 0.52 | 0 | 0.18 | 0.36 | 0.38 | 0.14 | 0.24 |
| 421 | CESAR | VALLEDUPAR | Caribbean | 111.67 | 1.83 | 0.73 | 0.38 | 0.63 | 0.47 | 0.96 | 0.75 | 0.22 |
| 422 | CESAR | AGUSTÍN CODAZZI | Caribbean | 119.83 | 0.5 | 0.72 | 0.11 | 0.49 | 0.38 | 0.61 | 0.56 | 0.05 |
| 423 | CESAR | BECERRIL | Caribbean | 120.17 | 0.5 | 0.74 | 0.07 | 0.33 | 0.49 | 0.69 | 0.34 | 0.35 |
| 424 | CESAR | LA PAZ | Caribbean | 135.17 | 0.5 | 0.75 | 0.07 | 0.36 | 0.38 | 0.6 | 0.39 | 0.21 |
| 425 | CESAR | PUEBLO BELLO | Caribbean | 110 | 0.83 | 0.62 | 0.18 | 0.29 | 0.38 | 0.63 | 0.28 | 0.35 |
| 426 | CESAR | SAN MARTÍN | Caribbean | 180.17 | 0 | 0.78 | 0.06 | 0.39 | 0.53 | 0.73 | 0.43 | 0.31 |
| 427 | CESAR | AGUACHICA | Caribbean | 129 | 0 | 0.73 | 0 | 0.3 | 0.4 | 0.54 | 0.3 | 0.23 |
| 428 | CÓRDOBA | MONTERÍA | Caribbean | 90.17 | 0.67 | 0.54 | 0.07 | 0.56 | 0.35 | 0.45 | 0.66 | -0.21 |
| 429 | CÓRDOBA | AYAPEL | Caribbean | 121.83 | 0 | 0.6 | 0 | 0.35 | 0.4 | 0.47 | 0.37 | 0.09 |
| 430 | CÓRDOBA | BUENAVISTA | Caribbean | 132.17 | 0 | 0.65 | 0.06 | 0.4 | 0.49 | 0.62 | 0.44 | 0.18 |
| 431 | CÓRDOBA | CANALETE | Caribbean | 82.5 | 0 | 0.39 | 0 | 0.17 | 0.33 | 0.28 | 0.13 | 0.15 |
| 432 | CÓRDOBA | CERETÉ | Caribbean | 84 | 0 | 0.45 | 0.06 | 0.56 | 0.41 | 0.44 | 0.65 | -0.22 |
| 433 | CÓRDOBA | CHIMÁ | Caribbean | 83.67 | 0.17 | 0.49 | 0.03 | 0.17 | 0.51 | 0.52 | 0.13 | 0.39 |
| 434 | CÓRDOBA | CHINÚ | Andean | 109 | 0.17 | 0.56 | 0.03 | 0.28 | 0.37 | 0.43 | 0.28 | 0.15 |
| 435 | CÓRDOBA | CIÉNAGA DE ORO | Caribbean | 87.17 | 0.17 | 0.55 | 0.08 | 0.18 | 0.41 | 0.51 | 0.14 | 0.37 |
| 436 | CÓRDOBA | COTORRA | Caribbean | 82.17 | 0 | 0.45 | 0 | 0.28 | 0.43 | 0.4 | 0.28 | 0.12 |
| 437 | CÓRDOBA | LA APARTADA | Caribbean | 121.67 | 0 | 0.57 | 0 | 0.2 | 0.38 | 0.42 | 0.16 | 0.26 |
| 438 | CÓRDOBA | LORICA | Caribbean | 84.33 | 0.17 | 0.53 | 0.06 | 0.46 | 0.42 | 0.49 | 0.52 | -0.03 |
| 439 | CÓRDOBA | LOS CÓRDOBAS | Caribbean | 80.83 | 0 | 0.38 | 0 | 0.17 | 0.38 | 0.31 | 0.13 | 0.17 |
| 440 | CÓRDOBA | MOMIL | Caribbean | 91.83 | 0.17 | 0.52 | 0.03 | 0.21 | 0.4 | 0.44 | 0.18 | 0.26 |
| 441 | CÓRDOBA | MOÑITOS | Caribbean | 81.33 | 0 | 0.42 | 0 | 0.23 | 0.41 | 0.36 | 0.21 | 0.15 |
| 442 | CÓRDOBA | PLANETA RICA | Caribbean | 111.83 | 0 | 0.61 | 0 | 0.18 | 0.34 | 0.41 | 0.15 | 0.27 |
| 443 | CÓRDOBA | PUEBLO NUEVO | Caribbean | 112 | 0 | 0.59 | 0 | 0.17 | 0.38 | 0.44 | 0.12 | 0.31 |
| 444 | CÓRDOBA | PUERTO ESCONDIDO | Caribbean | 80 | 0.17 | 0.39 | 0.06 | 0.33 | 0.38 | 0.37 | 0.34 | 0.03 |
| 445 | CÓRDOBA | PURÍSIMA DE LA CONCEPCIÓN | Caribbean | 92.5 | 0.17 | 0.53 | 0.03 | 0.21 | 0.38 | 0.43 | 0.18 | 0.25 |
| 446 | CÓRDOBA | SAHAGÚN | Caribbean | 111.17 | 0 | 0.57 | 0 | 0.18 | 0.38 | 0.42 | 0.14 | 0.28 |
| 447 | CÓRDOBA | SAN ANDRÉS DE SOTAVENTO | Caribbean | 91 | 0.17 | 0.46 | 0.03 | 0.2 | 0.4 | 0.41 | 0.17 | 0.24 |
| 448 | CÓRDOBA | SAN ANTERO | Caribbean | 92 | 0.33 | 0.57 | 0.15 | 0.47 | 0.37 | 0.56 | 0.53 | 0.03 |
| 449 | CÓRDOBA | SAN BERNARDO DEL VIENTO | Caribbean | 94 | 0 | 0.51 | 0 | 0.31 | 0.36 | 0.37 | 0.32 | 0.05 |
| 450 | CÓRDOBA | SAN CARLOS | Caribbean | 136.5 | 0 | 0.68 | 0 | 0.19 | 0.38 | 0.49 | 0.15 | 0.34 |
| 451 | CÓRDOBA | SAN JOSÉ DE URÉ | Caribbean | 119.33 | 0.33 | 0.59 | 0.05 | 0.16 | 0.47 | 0.57 | 0.12 | 0.45 |
| 452 | CÓRDOBA | SAN PELAYO | Caribbean | 83.83 | 0 | 0.47 | 0.06 | 0.44 | 0.51 | 0.54 | 0.49 | 0.05 |
| 453 | CÓRDOBA | TUCHÍN | Caribbean | 83.17 | 0.17 | 0.41 | 0.03 | 0.22 | 0.41 | 0.39 | 0.2 | 0.19 |
| 454 | CÓRDOBA | VALENCIA | Caribbean | 115 | 0.17 | 0.58 | 0.03 | 0.28 | 0.37 | 0.45 | 0.27 | 0.18 |
| 455 | CÓRDOBA | MONTELÍBANO | Caribbean | 142.67 | 0.33 | 0.72 | 0.05 | 0.29 | 0.36 | 0.54 | 0.28 | 0.26 |
| 456 | CÓRDOBA | PUERTO LIBERTADOR | Caribbean | 141.67 | 0.33 | 0.74 | 0.05 | 0.13 | 0.34 | 0.54 | 0.08 | 0.47 |
| 457 | CÓRDOBA | TIERRALTA | Andean | 147.5 | 0.67 | 0.76 | 0.1 | 0.33 | 0.34 | 0.59 | 0.34 | 0.25 |
| 458 | CUNDINAMARCA | ALBÁN | Andean | 125.33 | 0.17 | 0.54 | 0.13 | 0.36 | 0.41 | 0.56 | 0.39 | 0.17 |
| 459 | CUNDINAMARCA | ANAPOIMA | Andean | 124.5 | 0 | 0.58 | 0 | 0.27 | 0.37 | 0.42 | 0.26 | 0.16 |
| 460 | CUNDINAMARCA | ANOLAIMA | Andean | 128.33 | 0 | 0.57 | 0 | 0.36 | 0.37 | 0.42 | 0.38 | 0.04 |
| 461 | CUNDINAMARCA | ARBELÁEZ | Andean | 115.83 | 0 | 0.63 | 0 | 0.24 | 0.38 | 0.47 | 0.22 | 0.25 |
| 462 | CUNDINAMARCA | BELTRÁN | Andean | 127 | 0 | 0.54 | 0 | 0.3 | 0.38 | 0.4 | 0.3 | 0.1 |
| 463 | CUNDINAMARCA | BITUIMA | Andean | 126.5 | 0 | 0.53 | 0 | 0.35 | 0.41 | 0.43 | 0.37 | 0.06 |
| 464 | CUNDINAMARCA | BOJACÁ | Andean | 107.33 | 0 | 0.52 | 0 | 0.26 | 0.38 | 0.4 | 0.25 | 0.15 |
| 465 | CUNDINAMARCA | CABRERA | Andean | 136.17 | 0 | 0.66 | 0 | 0.19 | 0.41 | 0.51 | 0.16 | 0.35 |
| 466 | CUNDINAMARCA | CACHIPAY | Andean | 113.17 | 0 | 0.47 | 0 | 0.27 | 0.36 | 0.35 | 0.27 | 0.08 |
| 467 | CUNDINAMARCA | CAJICÁ | Andean | 79.33 | 0.33 | 0.45 | 0.14 | 0.47 | 0.42 | 0.53 | 0.54 | -0.01 |
| 468 | CUNDINAMARCA | CAPARRAPÍ | Andean | 148.33 | 0 | 0.68 | 0 | 0.39 | 0.41 | 0.51 | 0.43 | 0.08 |
| 469 | CUNDINAMARCA | CÁQUEZA | Andean | 94.67 | 0 | 0.46 | 0 | 0.26 | 0.41 | 0.39 | 0.24 | 0.15 |
| 470 | CUNDINAMARCA | CARMEN DE CARUPA | Andean | 85.17 | 0 | 0.54 | 0 | 0.32 | 0.37 | 0.41 | 0.33 | 0.07 |
| 471 | CUNDINAMARCA | CHAGUANÍ | Andean | 142 | 0 | 0.59 | 0 | 0.24 | 0.38 | 0.44 | 0.22 | 0.23 |
| 472 | CUNDINAMARCA | CHIPAQUE | Andean | 87 | 0 | 0.46 | 0 | 0.24 | 0.4 | 0.38 | 0.23 | 0.15 |
| 473 | CUNDINAMARCA | CHOACHÍ | Andean | 112.83 | 0 | 0.54 | 0 | 0.37 | 0.42 | 0.45 | 0.4 | 0.05 |
| 474 | CUNDINAMARCA | CHOCONTÁ | Andean | 77.17 | 0.33 | 0.48 | 0.12 | 0.35 | 0.39 | 0.5 | 0.36 | 0.13 |
| 475 | CUNDINAMARCA | COGUA | Andean | 78 | 0 | 0.43 | 0 | 0.36 | 0.44 | 0.4 | 0.39 | 0.01 |
| 476 | CUNDINAMARCA | COTA | Andean | 80 | 0.5 | 0.43 | 0.18 | 0.4 | 0.38 | 0.51 | 0.44 | 0.08 |
| 477 | CUNDINAMARCA | CUCUNUBÁ | Andean | 76 | 0.17 | 0.44 | 0.11 | 0.45 | 0.44 | 0.51 | 0.5 | 0.01 |
| 478 | CUNDINAMARCA | EL COLEGIO | Andean | 131.5 | 0 | 0.6 | 0 | 0.34 | 0.6 | 0.63 | 0.36 | 0.27 |
| 479 | CUNDINAMARCA | EL PEÑÓN | Andean | 140.5 | 0 | 0.61 | 0 | 0.23 | 0.35 | 0.43 | 0.21 | 0.22 |
| 480 | CUNDINAMARCA | FACATATIVÁ | Andean | 84.33 | 0 | 0.48 | 0 | 0.36 | 0.38 | 0.38 | 0.38 | -0.01 |
| 481 | CUNDINAMARCA | FÓMEQUE | Andean | 116.33 | 0 | 0.65 | 0 | 0.34 | 0.48 | 0.56 | 0.36 | 0.2 |
| 482 | CUNDINAMARCA | FOSCA | Andean | 95.33 | 0 | 0.5 | 0 | 0.23 | 0.43 | 0.43 | 0.21 | 0.22 |
| 483 | CUNDINAMARCA | FUNZA | Andean | 79.67 | 0.17 | 0.4 | 0.12 | 0.38 | 0.42 | 0.48 | 0.41 | 0.06 |
| 484 | CUNDINAMARCA | FÚQUENE | Andean | 76.83 | 0 | 0.43 | 0 | 0.37 | 0.42 | 0.38 | 0.4 | -0.01 |
| 485 | CUNDINAMARCA | GACHALÁ | Andean | 141.67 | 0 | 0.69 | 0 | 0.3 | 0.35 | 0.47 | 0.3 | 0.17 |
| 486 | CUNDINAMARCA | GACHANCIPÁ | Andean | 75.5 | 0 | 0.4 | 0 | 0.28 | 0.43 | 0.37 | 0.28 | 0.1 |
| 487 | CUNDINAMARCA | GACHETÁ | Andean | 94 | 0 | 0.52 | 0 | 0.22 | 0.53 | 0.53 | 0.19 | 0.34 |
| 488 | CUNDINAMARCA | GAMA | Andean | 94.83 | 0 | 0.48 | 0 | 0.25 | 0.33 | 0.34 | 0.23 | 0.11 |
| 489 | CUNDINAMARCA | GRANADA | Andean | 162.33 | 0 | 0.64 | 0 | 0.26 | 0.41 | 0.49 | 0.25 | 0.24 |
| 490 | CUNDINAMARCA | GUACHETÁ | Andean | 77 | 0 | 0.5 | 0 | 0.33 | 0.41 | 0.41 | 0.34 | 0.07 |
| 491 | CUNDINAMARCA | GUADUAS | Andean | 147 | 0.17 | 0.62 | 0.07 | 0.31 | 0.34 | 0.49 | 0.32 | 0.17 |
| 492 | CUNDINAMARCA | GUASCA | Andean | 85.5 | 0 | 0.58 | 0 | 0.37 | 0.38 | 0.43 | 0.4 | 0.03 |
| 493 | CUNDINAMARCA | GUATAQUÍ | Andean | 118.17 | 0 | 0.51 | 0 | 0.24 | 0.4 | 0.4 | 0.22 | 0.18 |
| 494 | CUNDINAMARCA | GUATAVITA | Andean | 81.67 | 0 | 0.56 | 0 | 0.36 | 0.51 | 0.54 | 0.39 | 0.15 |
| 495 | CUNDINAMARCA | GUAYABAL DE SÍQUIMA | Andean | 127.67 | 0 | 0.53 | 0 | 0.36 | 0.36 | 0.39 | 0.38 | 0.01 |
| 496 | CUNDINAMARCA | GUAYABETAL | Andean | 139.5 | 0 | 0.64 | 0 | 0.34 | 0.38 | 0.47 | 0.36 | 0.11 |
| 497 | CUNDINAMARCA | GUTIÉRREZ | Andean | 112.83 | 0 | 0.61 | 0 | 0.19 | 0.53 | 0.58 | 0.16 | 0.42 |
| 498 | CUNDINAMARCA | JERUSALÉN | Andean | 128.5 | 0 | 0.55 | 0 | 0.19 | 0.39 | 0.42 | 0.16 | 0.26 |
| 499 | CUNDINAMARCA | JUNÍN | Andean | 113 | 0.17 | 0.57 | 0 | 0.32 | 0.37 | 0.42 | 0.33 | 0.1 |
| 500 | CUNDINAMARCA | LA CALERA | Andean | 88.17 | 0 | 0.56 | 0 | 0.39 | 0.37 | 0.41 | 0.42 | -0.01 |
| 501 | CUNDINAMARCA | LA MESA | Andean | 117.5 | 0.17 | 0.5 | 0.11 | 0.37 | 0.52 | 0.61 | 0.4 | 0.21 |
| 502 | CUNDINAMARCA | LA PALMA | Andean | 138 | 0 | 0.66 | 0 | 0.21 | 0.35 | 0.45 | 0.18 | 0.27 |
| 503 | CUNDINAMARCA | LA PEÑA | Andean | 131 | 0 | 0.55 | 0 | 0.23 | 0.4 | 0.43 | 0.21 | 0.22 |
| 504 | CUNDINAMARCA | LA VEGA | Andean | 129.67 | 0 | 0.58 | 0 | 0.37 | 0.39 | 0.44 | 0.4 | 0.04 |
| 505 | CUNDINAMARCA | LENGUAZAQUE | Andean | 75.5 | 0.33 | 0.42 | 0 | 0.33 | 0.58 | 0.52 | 0.35 | 0.17 |
| 506 | CUNDINAMARCA | MACHETÁ | Andean | 90.17 | 0 | 0.51 | 0 | 0.23 | 0.37 | 0.38 | 0.2 | 0.18 |
| 507 | CUNDINAMARCA | MADRID | Andean | 82 | 0.17 | 0.44 | 0.11 | 0.27 | 0.36 | 0.43 | 0.26 | 0.17 |
| 508 | CUNDINAMARCA | MANTA | Andean | 93.5 | 0 | 0.5 | 0 | 0.32 | 0.31 | 0.33 | 0.33 | -0.01 |
| 509 | CUNDINAMARCA | MEDINA | Andean | 154.17 | 0 | 0.71 | 0 | 0.38 | 0.6 | 0.71 | 0.41 | 0.29 |
| 510 | CUNDINAMARCA | NARIÑO | Andean | 148.83 | 0 | 0.6 | 0 | 0.25 | 0.47 | 0.52 | 0.23 | 0.29 |
| 511 | CUNDINAMARCA | NEMOCÓN | Andean | 76 | 0.17 | 0.41 | 0.11 | 0.38 | 0.34 | 0.4 | 0.41 | 0 |
| 512 | CUNDINAMARCA | NILO | Andean | 119.83 | 0.17 | 0.62 | 0.1 | 0.35 | 0.42 | 0.58 | 0.37 | 0.21 |
| 513 | CUNDINAMARCA | NIMAIMA | Andean | 129.5 | 0 | 0.56 | 0 | 0.27 | 0.35 | 0.4 | 0.26 | 0.14 |
| 514 | CUNDINAMARCA | NOCAIMA | Andean | 128.67 | 0 | 0.55 | 0 | 0.35 | 0.42 | 0.45 | 0.37 | 0.08 |
| 515 | CUNDINAMARCA | VENECIA | Andean | 122.17 | 0 | 0.58 | 0 | 0.22 | 0.52 | 0.56 | 0.19 | 0.37 |
| 516 | CUNDINAMARCA | PACHO | Andean | 126 | 0 | 0.57 | 0 | 0.22 | 0.52 | 0.56 | 0.2 | 0.36 |
| 517 | CUNDINAMARCA | PAIME | Andean | 128.5 | 0 | 0.57 | 0 | 0.2 | 0.39 | 0.44 | 0.16 | 0.28 |
| 518 | CUNDINAMARCA | PANDI | Andean | 112.83 | 0 | 0.52 | 0 | 0.26 | 0.58 | 0.57 | 0.25 | 0.32 |
| 519 | CUNDINAMARCA | PARATEBUENO | Andean | 151.83 | 0 | 0.68 | 0 | 0.38 | 0.32 | 0.44 | 0.41 | 0.03 |
| 520 | CUNDINAMARCA | PASCA | Andean | 89.33 | 0.17 | 0.52 | 0.09 | 0.22 | 0.34 | 0.45 | 0.19 | 0.26 |
| 521 | CUNDINAMARCA | PUERTO SALGAR | Andean | 141.5 | 0 | 0.65 | 0 | 0.4 | 0.49 | 0.57 | 0.44 | 0.13 |
| 522 | CUNDINAMARCA | PULÍ | Andean | 133.67 | 0 | 0.59 | 0 | 0.31 | 0.38 | 0.43 | 0.31 | 0.12 |
| 523 | CUNDINAMARCA | QUEBRADANEGRA | Andean | 131.5 | 0 | 0.57 | 0 | 0.25 | 0.38 | 0.43 | 0.24 | 0.19 |
| 524 | CUNDINAMARCA | QUETAME | Andean | 134.67 | 0 | 0.61 | 0 | 0.23 | 0.37 | 0.45 | 0.21 | 0.23 |
| 525 | CUNDINAMARCA | QUIPILE | Andean | 137.5 | 0 | 0.61 | 0 | 0.22 | 0.41 | 0.48 | 0.2 | 0.28 |
| 526 | CUNDINAMARCA | APULO | Andean | 117.33 | 0 | 0.55 | 0 | 0.35 | 0.36 | 0.4 | 0.38 | 0.02 |
| 527 | CUNDINAMARCA | RICAURTE | Andean | 131.5 | 0 | 0.57 | 0 | 0.35 | 0.41 | 0.46 | 0.37 | 0.09 |
| 528 | CUNDINAMARCA | SAN ANTONIO DEL TEQUENDAMA | Andean | 127.5 | 0.33 | 0.58 | 0.13 | 0.4 | 0.4 | 0.57 | 0.44 | 0.14 |
| 529 | CUNDINAMARCA | SAN BERNARDO | Andean | 114.83 | 0.17 | 0.6 | 0.1 | 0.21 | 0.33 | 0.49 | 0.19 | 0.3 |
| 530 | CUNDINAMARCA | SAN CAYETANO | Andean | 136.83 | 0 | 0.62 | 0 | 0.22 | 0.38 | 0.46 | 0.2 | 0.26 |
| 531 | CUNDINAMARCA | SAN FRANCISCO | Andean | 157.17 | 0 | 0.65 | 0 | 0.39 | 0.36 | 0.46 | 0.42 | 0.04 |
| 532 | CUNDINAMARCA | SASAIMA | Andean | 128.33 | 0 | 0.57 | 0 | 0.36 | 0.36 | 0.41 | 0.38 | 0.03 |
| 533 | CUNDINAMARCA | SESQUILÉ | Andean | 76 | 0.17 | 0.46 | 0.03 | 0.35 | 0.55 | 0.54 | 0.38 | 0.16 |
| 534 | CUNDINAMARCA | SILVANIA | Andean | 128.33 | 0 | 0.6 | 0 | 0.35 | 0.4 | 0.46 | 0.37 | 0.09 |
| 535 | CUNDINAMARCA | SIMIJACA | Andean | 79.83 | 0 | 0.48 | 0 | 0.25 | 0.48 | 0.47 | 0.24 | 0.23 |
| 536 | CUNDINAMARCA | SOPÓ | Andean | 77.17 | 0.17 | 0.45 | 0 | 0.38 | 0.52 | 0.48 | 0.42 | 0.06 |
| 537 | CUNDINAMARCA | SUBACHOQUE | Andean | 82 | 0 | 0.46 | 0 | 0.36 | 0.67 | 0.62 | 0.38 | 0.23 |
| 538 | CUNDINAMARCA | SUESCA | Andean | 76 | 0 | 0.46 | 0 | 0.46 | 0.4 | 0.38 | 0.52 | -0.14 |
| 539 | CUNDINAMARCA | SUPATÁ | Andean | 123.67 | 0 | 0.52 | 0 | 0.23 | 0.35 | 0.38 | 0.21 | 0.16 |
| 540 | CUNDINAMARCA | SUSA | Andean | 79.17 | 0 | 0.47 | 0 | 0.24 | 0.4 | 0.38 | 0.23 | 0.15 |
| 541 | CUNDINAMARCA | SUTATAUSA | Andean | 76.5 | 0.17 | 0.41 | 0.12 | 0.37 | 0.39 | 0.46 | 0.4 | 0.06 |
| 542 | CUNDINAMARCA | TABIO | Andean | 79.33 | 0 | 0.4 | 0.06 | 0.53 | 0.36 | 0.36 | 0.61 | -0.25 |
| 543 | CUNDINAMARCA | TAUSA | Andean | 72.33 | 0 | 0.43 | 0 | 0.46 | 0.4 | 0.36 | 0.51 | -0.15 |
| 544 | CUNDINAMARCA | TENA | Andean | 113.17 | 0 | 0.47 | 0 | 0.38 | 0.53 | 0.5 | 0.42 | 0.08 |
| 545 | CUNDINAMARCA | TENJO | Andean | 80 | 0.33 | 0.4 | 0.05 | 0.36 | 0.44 | 0.43 | 0.39 | 0.04 |
| 546 | CUNDINAMARCA | TIBACUY | Andean | 128.17 | 0 | 0.63 | 0 | 0.36 | 0.44 | 0.51 | 0.39 | 0.12 |
| 547 | CUNDINAMARCA | TIBIRITA | Andean | 89.5 | 0 | 0.41 | 0 | 0.35 | 0.38 | 0.33 | 0.37 | -0.03 |
| 548 | CUNDINAMARCA | TOCAIMA | Andean | 126 | 0.17 | 0.6 | 0.1 | 0.23 | 0.41 | 0.55 | 0.21 | 0.34 |
| 549 | CUNDINAMARCA | TOCANCIPÁ | Andean | 76.83 | 0.5 | 0.45 | 0.14 | 0.39 | 0.42 | 0.53 | 0.43 | 0.1 |
| 550 | CUNDINAMARCA | TOPAIPÍ | Andean | 127.5 | 0 | 0.52 | 0 | 0.21 | 0.43 | 0.44 | 0.18 | 0.26 |
| 551 | CUNDINAMARCA | UBALÁ | Andean | 143.67 | 0 | 0.73 | 0 | 0.3 | 0.41 | 0.55 | 0.3 | 0.25 |
| 552 | CUNDINAMARCA | UBAQUE | Andean | 94.33 | 0.17 | 0.46 | 0.11 | 0.35 | 0.41 | 0.5 | 0.37 | 0.13 |
| 553 | CUNDINAMARCA | VILLA DE SAN DIEGO DE UBATÉ | Andean | 77.33 | 0.17 | 0.43 | 0.11 | 0.38 | 0.59 | 0.64 | 0.41 | 0.23 |
| 554 | CUNDINAMARCA | UNE | Andean | 87 | 0 | 0.48 | 0 | 0.22 | 0.43 | 0.41 | 0.19 | 0.22 |
| 555 | CUNDINAMARCA | ÚTICA | Andean | 132 | 0 | 0.56 | 0 | 0.24 | 0.38 | 0.42 | 0.22 | 0.19 |
| 556 | CUNDINAMARCA | VERGARA | Andean | 127.67 | 0 | 0.52 | 0 | 0.22 | 0.35 | 0.37 | 0.19 | 0.18 |
| 557 | CUNDINAMARCA | VIANÍ | Andean | 140.5 | 0 | 0.58 | 0 | 0.25 | 0.38 | 0.44 | 0.23 | 0.2 |
| 558 | CUNDINAMARCA | VILLAGÓMEZ | Andean | 123.33 | 0 | 0.49 | 0 | 0.24 | 0.42 | 0.41 | 0.22 | 0.19 |
| 559 | CUNDINAMARCA | VILLAPINZÓN | Andean | 76.83 | 0 | 0.46 | 0 | 0.23 | 0.38 | 0.36 | 0.21 | 0.15 |
| 560 | CUNDINAMARCA | VILLETA | Andean | 131.33 | 0 | 0.57 | 0 | 0.37 | 0.41 | 0.45 | 0.4 | 0.05 |
| 561 | CUNDINAMARCA | VIOTÁ | Andean | 119 | 0 | 0.58 | 0 | 0.33 | 0.4 | 0.45 | 0.34 | 0.1 |
| 562 | CUNDINAMARCA | YACOPÍ | Andean | 142 | 0 | 0.66 | 0 | 0.27 | 0.31 | 0.42 | 0.26 | 0.16 |
| 563 | CUNDINAMARCA | ZIPACÓN | Andean | 116.67 | 0.33 | 0.49 | 0.14 | 0.27 | 0.37 | 0.5 | 0.27 | 0.24 |
| 564 | CUNDINAMARCA | ZIPAQUIRÁ | Andean | 79.83 | 0.33 | 0.49 | 0.11 | 0.49 | 0.37 | 0.47 | 0.56 | -0.09 |
| 565 | CUNDINAMARCA | AGUA DE DIOS | Andean | 112.17 | 0.67 | 0.53 | 0.14 | 0.25 | 0.36 | 0.52 | 0.24 | 0.28 |
| 566 | CUNDINAMARCA | GIRARDOT | Andean | 116 | 0 | 0.52 | 0 | 0.45 | 0.4 | 0.42 | 0.5 | -0.08 |
| 567 | CUNDINAMARCA | SOACHA | Andean | 88 | 0.5 | 0.55 | 0.12 | 0.36 | 0.41 | 0.55 | 0.38 | 0.18 |
| 568 | CUNDINAMARCA | CHÍA | Andean | 79.17 | 0.33 | 0.43 | 0.14 | 0.4 | 0.4 | 0.5 | 0.43 | 0.07 |
| 569 | CUNDINAMARCA | EL ROSAL | Andean | 84.5 | 0 | 0.43 | 0 | 0.26 | 0.35 | 0.32 | 0.25 | 0.07 |
| 570 | CUNDINAMARCA | MOSQUERA | Andean | 108 | 0 | 0.52 | 0 | 0.37 | 0.38 | 0.39 | 0.39 | 0 |
| 571 | CUNDINAMARCA | SIBATÉ | Andean | 85 | 0 | 0.51 | 0 | 0.25 | 0.4 | 0.41 | 0.24 | 0.17 |
| 572 | CUNDINAMARCA | FUSAGASUGÁ | Andean | 128.83 | 0.5 | 0.61 | 0.16 | 0.52 | 0.46 | 0.67 | 0.6 | 0.07 |
| 573 | CUNDINAMARCA | SAN JUAN DE RIOSECO | Andean | 139.83 | 0 | 0.62 | 0 | 0.22 | 0.39 | 0.47 | 0.19 | 0.28 |
| 574 | CHOCÓ | RIOSUCIO | Pacific | 142.33 | 5.5 | 0.63 | 0.48 | 0.54 | 0.4 | 0.94 | 0.63 | 0.31 |
| 575 | CHOCÓ | QUIBDÓ | Pacific | 145.83 | 7 | 0.7 | 0.44 | 0.5 | 0.47 | 1 | 0.58 | 0.42 |
| 576 | CHOCÓ | ACANDÍ | Pacific | 128.67 | 1.17 | 0.63 | 0.14 | 0.45 | 0.31 | 0.53 | 0.51 | 0.03 |
| 577 | CHOCÓ | ALTO BAUDÓ | Pacific | 114.83 | 4 | 0.5 | 0.24 | 0.13 | 0.33 | 0.56 | 0.08 | 0.49 |
| 578 | CHOCÓ | ATRATO | Pacific | 119.33 | 2 | 0.53 | 0.17 | 0.2 | 0.32 | 0.5 | 0.16 | 0.34 |
| 579 | CHOCÓ | BAGADÓ | Pacific | 140.33 | 1.17 | 0.64 | 0.15 | 0.17 | 0.46 | 0.68 | 0.13 | 0.54 |
| 580 | CHOCÓ | BAHÍA SOLANO | Pacific | 115.17 | 2 | 0.55 | 0.17 | 0.49 | 0.37 | 0.57 | 0.56 | 0.01 |
| 581 | CHOCÓ | BAJO BAUDÓ | Pacific | 117.17 | 5.5 | 0.57 | 0.27 | 0.25 | 0.25 | 0.56 | 0.24 | 0.32 |
| 582 | CHOCÓ | BOJAYÁ | Pacific | 121 | 3.33 | 0.57 | 0.22 | 0.15 | 0.23 | 0.5 | 0.11 | 0.4 |
| 583 | CHOCÓ | EL CANTÓN DEL SAN PABLO | Pacific | 110.33 | 1.17 | 0.5 | 0.1 | 0.16 | 0.37 | 0.47 | 0.12 | 0.35 |
| 584 | CHOCÓ | CARMEN DEL DARIÉN | Pacific | 143.17 | 4 | 0.67 | 0.23 | 0.14 | 0.29 | 0.63 | 0.09 | 0.54 |
| 585 | CHOCÓ | CÉRTEGUI | Pacific | 131.83 | 1.5 | 0.58 | 0.21 | 0.32 | 0.36 | 0.61 | 0.33 | 0.28 |
| 586 | CHOCÓ | CONDOTO | Pacific | 141 | 0.83 | 0.65 | 0.12 | 0.27 | 0.39 | 0.6 | 0.26 | 0.35 |
| 587 | CHOCÓ | EL CARMEN DE ATRATO | Pacific | 140.17 | 1.83 | 0.65 | 0.14 | 0.32 | 0.34 | 0.58 | 0.33 | 0.25 |
| 588 | CHOCÓ | EL LITORAL DEL SAN JUAN | Pacific | 145.83 | 3 | 0.68 | 0.2 | 0.35 | 0.21 | 0.53 | 0.37 | 0.16 |
| 589 | CHOCÓ | ISTMINA | Pacific | 112.33 | 2.83 | 0.54 | 0.27 | 0.3 | 0.36 | 0.65 | 0.3 | 0.35 |
| 590 | CHOCÓ | JURADÓ | Pacific | 133.5 | 1.83 | 0.59 | 0.17 | 0.32 | 0.23 | 0.47 | 0.33 | 0.14 |
| 591 | CHOCÓ | LLORÓ | Pacific | 141.33 | 2.5 | 0.66 | 0.2 | 0.16 | 0.28 | 0.58 | 0.11 | 0.46 |
| 592 | CHOCÓ | MEDIO ATRATO | Pacific | 145.17 | 1.83 | 0.68 | 0.14 | 0.15 | 0.24 | 0.5 | 0.09 | 0.41 |
| 593 | CHOCÓ | MEDIO BAUDÓ | Pacific | 105.33 | 3.67 | 0.5 | 0.24 | 0.15 | 0.32 | 0.56 | 0.1 | 0.45 |
| 594 | CHOCÓ | MEDIO SAN JUAN | Pacific | 110.83 | 1 | 0.53 | 0.19 | 0.43 | 0.24 | 0.46 | 0.48 | -0.02 |
| 595 | CHOCÓ | NÓVITA | Pacific | 127.17 | 0.83 | 0.57 | 0.16 | 0.16 | 0.29 | 0.5 | 0.12 | 0.38 |
| 596 | CHOCÓ | NUQUÍ | Pacific | 101.83 | 1.33 | 0.5 | 0.26 | 0.6 | 0.25 | 0.51 | 0.7 | -0.19 |
| 597 | CHOCÓ | RÍO IRÓ | Pacific | 138.33 | 0.83 | 0.64 | 0.12 | 0.11 | 0.35 | 0.56 | 0.05 | 0.51 |
| 598 | CHOCÓ | RÍO QUITO | Pacific | 114.83 | 2.5 | 0.53 | 0.2 | 0.17 | 0.46 | 0.67 | 0.13 | 0.54 |
| 599 | CHOCÓ | SAN JOSÉ DEL PALMAR | Pacific | 142.67 | 1.17 | 0.68 | 0.15 | 0.29 | 0.4 | 0.65 | 0.29 | 0.36 |
| 600 | CHOCÓ | SIPÍ | Pacific | 142.5 | 0.83 | 0.63 | 0.12 | 0.25 | 0.3 | 0.5 | 0.24 | 0.26 |
| 601 | CHOCÓ | TADÓ | Pacific | 142.83 | 2.17 | 0.69 | 0.19 | 0.3 | 0.31 | 0.62 | 0.3 | 0.32 |
| 602 | CHOCÓ | UNGUÍA | Pacific | 138.83 | 1.5 | 0.68 | 0.34 | 0.45 | 0.37 | 0.81 | 0.51 | 0.3 |
| 603 | CHOCÓ | UNIÓN PANAMERICANA | Pacific | 110.5 | 0.83 | 0.54 | 0.09 | 0.22 | 0.39 | 0.49 | 0.19 | 0.3 |
| 604 | HUILA | NEIVA | Andean | 121.67 | 1 | 0.73 | 0.17 | 0.68 | 0.41 | 0.71 | 0.81 | -0.11 |
| 605 | HUILA | GIGANTE | Andean | 112.5 | 0 | 0.63 | 0 | 0.32 | 0.48 | 0.55 | 0.33 | 0.22 |
| 606 | HUILA | GUADALUPE | Andean | 155 | 0 | 0.68 | 0 | 0.2 | 0.44 | 0.54 | 0.17 | 0.38 |
| 607 | HUILA | HOBO | Andean | 118.5 | 0 | 0.65 | 0 | 0.23 | 0.39 | 0.49 | 0.21 | 0.28 |
| 608 | HUILA | ÍQUIRA | Andean | 110.33 | 0.33 | 0.62 | 0.05 | 0.27 | 0.35 | 0.49 | 0.26 | 0.23 |
| 609 | HUILA | ISNOS | Andean | 103 | 0.83 | 0.53 | 0.27 | 0.35 | 0.4 | 0.67 | 0.37 | 0.3 |
| 610 | HUILA | LA ARGENTINA | Andean | 109.83 | 0.67 | 0.63 | 0.16 | 0.28 | 0.35 | 0.59 | 0.27 | 0.32 |
| 611 | HUILA | LA PLATA | Andean | 111.83 | 1.5 | 0.61 | 0.16 | 0.44 | 0.4 | 0.62 | 0.49 | 0.13 |
| 612 | HUILA | NÁTAGA | Andean | 106.17 | 0.5 | 0.57 | 0.07 | 0.3 | 0.36 | 0.48 | 0.3 | 0.18 |
| 613 | HUILA | OPORAPA | Andean | 107.67 | 0.17 | 0.51 | 0.03 | 0.31 | 0.4 | 0.43 | 0.32 | 0.11 |
| 614 | HUILA | PAICOL | Andean | 111.5 | 0.17 | 0.59 | 0.03 | 0.29 | 0.39 | 0.47 | 0.29 | 0.18 |
| 615 | HUILA | PALERMO | Andean | 117 | 0.17 | 0.73 | 0.03 | 0.5 | 0.38 | 0.54 | 0.58 | -0.03 |
| 616 | HUILA | PALESTINA | Andean | 115.5 | 0 | 0.5 | 0 | 0.32 | 0.35 | 0.36 | 0.33 | 0.03 |
| 617 | HUILA | PITAL | Andean | 110.33 | 0.17 | 0.55 | 0.03 | 0.21 | 0.39 | 0.45 | 0.18 | 0.26 |
| 618 | HUILA | PITALITO | Andean | 107.5 | 0.83 | 0.55 | 0.2 | 0.6 | 0.41 | 0.63 | 0.71 | -0.08 |
| 619 | HUILA | RIVERA | Andean | 116.5 | 0.17 | 0.64 | 0.03 | 0.46 | 0.35 | 0.47 | 0.51 | -0.05 |
| 620 | HUILA | SALADOBLANCO | Andean | 107 | 0 | 0.56 | 0 | 0.3 | 0.48 | 0.51 | 0.3 | 0.2 |
| 621 | HUILA | SAN AGUSTÍN | Andean | 106.33 | 0.83 | 0.6 | 0.14 | 0.33 | 0.43 | 0.63 | 0.35 | 0.28 |
| 622 | HUILA | SANTA MARÍA | Andean | 140 | 0 | 0.63 | 0 | 0.31 | 0.39 | 0.47 | 0.32 | 0.15 |
| 623 | HUILA | SUAZA | Andean | 131 | 0 | 0.59 | 0 | 0.29 | 0.36 | 0.42 | 0.29 | 0.12 |
| 624 | HUILA | TARQUI | Andean | 110.83 | 0 | 0.57 | 0 | 0.3 | 0.4 | 0.45 | 0.3 | 0.15 |
| 625 | HUILA | TESALIA | Andean | 111.67 | 0.33 | 0.63 | 0.05 | 0.2 | 0.43 | 0.55 | 0.16 | 0.39 |
| 626 | HUILA | TELLO | Andean | 119.17 | 0 | 0.63 | 0 | 0.27 | 0.35 | 0.44 | 0.26 | 0.18 |
| 627 | HUILA | TERUEL | Andean | 106.17 | 0 | 0.63 | 0 | 0.28 | 0.38 | 0.47 | 0.28 | 0.19 |
| 628 | HUILA | TIMANÁ | Andean | 108.5 | 0 | 0.57 | 0 | 0.33 | 0.38 | 0.43 | 0.35 | 0.08 |
| 629 | HUILA | VILLAVIEJA | Andean | 120.33 | 0.67 | 0.63 | 0.18 | 0.55 | 0.39 | 0.64 | 0.65 | -0.01 |
| 630 | HUILA | YAGUARÁ | Andean | 110.67 | 0 | 0.66 | 0 | 0.41 | 0.38 | 0.47 | 0.45 | 0.02 |
| 631 | HUILA | GARZÓN | Andean | 130 | 0.17 | 0.7 | 0.08 | 0.4 | 0.45 | 0.64 | 0.44 | 0.2 |
| 632 | HUILA | ELÍAS | Andean | 106.33 | 0 | 0.51 | 0 | 0.22 | 0.47 | 0.47 | 0.2 | 0.27 |
| 633 | HUILA | COLOMBIA | Andean | 109.83 | 0.17 | 0.63 | 0.06 | 0.28 | 0.33 | 0.47 | 0.28 | 0.2 |
| 634 | HUILA | CAMPOALEGRE | Andean | 119.5 | 0 | 0.67 | 0 | 0.49 | 0.37 | 0.48 | 0.56 | -0.08 |
| 635 | HUILA | BARAYA | Andean | 116.33 | 0 | 0.62 | 0 | 0.18 | 0.4 | 0.48 | 0.14 | 0.34 |
| 636 | HUILA | ALTAMIRA | Andean | 108.67 | 0 | 0.58 | 0 | 0.39 | 0.4 | 0.45 | 0.43 | 0.02 |
| 637 | HUILA | ALGECIRAS | Andean | 117.83 | 0 | 0.58 | 0 | 0.28 | 0.25 | 0.31 | 0.27 | 0.04 |
| 638 | HUILA | AIPE | Andean | 119.67 | 0 | 0.66 | 0 | 0.39 | 0.35 | 0.46 | 0.42 | 0.04 |
| 639 | HUILA | AGRADO | Andean | 111.5 | 0.17 | 0.59 | 0 | 0.31 | 0.44 | 0.49 | 0.32 | 0.17 |
| 640 | HUILA | ACEVEDO | Andean | 116.33 | 0 | 0.57 | 0 | 0.31 | 0.41 | 0.45 | 0.32 | 0.13 |
| 641 | LA GUAJIRA | BARRANCAS | Andean | 119.67 | 1.33 | 0.67 | 0.11 | 0.38 | 0.46 | 0.65 | 0.41 | 0.24 |
| 642 | LA GUAJIRA | DISTRACCIÓN | Andean | 111.67 | 0.5 | 0.64 | 0.07 | 0.22 | 0.53 | 0.67 | 0.19 | 0.48 |
| 643 | LA GUAJIRA | EL MOLINO | Caribbean | 115 | 0 | 0.58 | 0 | 0.2 | 0.39 | 0.44 | 0.17 | 0.27 |
| 644 | LA GUAJIRA | FONSECA | Andean | 119.83 | 1.17 | 0.65 | 0.18 | 0.29 | 0.35 | 0.62 | 0.29 | 0.33 |
| 645 | LA GUAJIRA | HATONUEVO | Andean | 111 | 1.17 | 0.61 | 0.1 | 0.3 | 0.41 | 0.57 | 0.31 | 0.26 |
| 646 | LA GUAJIRA | LA JAGUA DEL PILAR | Caribbean | 111.17 | 0 | 0.55 | 0 | 0.22 | 0.38 | 0.42 | 0.19 | 0.23 |
| 647 | LA GUAJIRA | URUMITA | Caribbean | 114 | 0 | 0.58 | 0 | 0.28 | 0.37 | 0.42 | 0.28 | 0.14 |
| 648 | LA GUAJIRA | VILLANUEVA | Caribbean | 178.83 | 0.17 | 0.72 | 0.06 | 0.41 | 0.39 | 0.58 | 0.45 | 0.13 |
| 649 | LA GUAJIRA | RIOHACHA | Caribbean | 112.83 | 3.33 | 0.72 | 0.42 | 0.73 | 0.38 | 0.91 | 0.88 | 0.03 |
| 650 | LA GUAJIRA | ALBANIA | Andean | 161.67 | 0.67 | 0.7 | 0.12 | 0.55 | 0.39 | 0.63 | 0.64 | -0.01 |
| 651 | LA GUAJIRA | DIBULLA | Andean | 110.33 | 0.33 | 0.7 | 0.08 | 0.42 | 0.39 | 0.6 | 0.47 | 0.13 |
| 652 | LA GUAJIRA | SAN JUAN DEL CESAR | Andean | 120.33 | 1 | 0.7 | 0.25 | 0.23 | 0.39 | 0.75 | 0.21 | 0.54 |
| 653 | LA GUAJIRA | URIBIA | Caribbean | 54.83 | 0.33 | 0.55 | 0.14 | 0.57 | 0.38 | 0.55 | 0.67 | -0.12 |
| 654 | LA GUAJIRA | MAICAO | Caribbean | 111 | 1.33 | 0.63 | 0.16 | 0.46 | 0.38 | 0.61 | 0.52 | 0.09 |
| 655 | LA GUAJIRA | MANAURE | Caribbean | 96.33 | 0.33 | 0.58 | 0.08 | 0.47 | 0.36 | 0.5 | 0.53 | -0.03 |
| 656 | MAGDALENA | PEDRAZA | Caribbean | 92.17 | 0 | 0.45 | 0 | 0.17 | 0.47 | 0.44 | 0.13 | 0.31 |
| 657 | MAGDALENA | ALGARROBO | Caribbean | 89.17 | 0 | 0.45 | 0 | 0.25 | 0.39 | 0.37 | 0.24 | 0.13 |
| 658 | MAGDALENA | ARIGUANÍ | Caribbean | 90.67 | 0 | 0.5 | 0 | 0.25 | 0.34 | 0.35 | 0.24 | 0.11 |
| 659 | MAGDALENA | CERRO DE SAN ANTONIO | Caribbean | 91.17 | 0 | 0.45 | 0 | 0.21 | 0.41 | 0.38 | 0.18 | 0.2 |
| 660 | MAGDALENA | CHIVOLO | Caribbean | 87 | 0 | 0.45 | 0 | 0.17 | 0.37 | 0.35 | 0.13 | 0.22 |
| 661 | MAGDALENA | CONCORDIA | Caribbean | 119.67 | 0 | 0.56 | 0 | 0.14 | 0.39 | 0.43 | 0.09 | 0.34 |
| 662 | MAGDALENA | EL PIÑÓN | Caribbean | 89.83 | 0 | 0.49 | 0 | 0.18 | 0.36 | 0.36 | 0.14 | 0.23 |
| 663 | MAGDALENA | EL RETÉN | Caribbean | 101.67 | 0.17 | 0.54 | 0.03 | 0.31 | 0.49 | 0.53 | 0.32 | 0.21 |
| 664 | MAGDALENA | GUAMAL | Andean | 156.5 | 0 | 0.71 | 0 | 0.35 | 0.39 | 0.52 | 0.37 | 0.15 |
| 665 | MAGDALENA | NUEVA GRANADA | Caribbean | 86 | 0 | 0.41 | 0 | 0.18 | 0.35 | 0.31 | 0.14 | 0.17 |
| 666 | MAGDALENA | PIJIÑO DEL CARMEN | Caribbean | 91.33 | 0 | 0.5 | 0 | 0.18 | 0.37 | 0.37 | 0.14 | 0.23 |
| 667 | MAGDALENA | PIVIJAY | Caribbean | 95.83 | 0.17 | 0.53 | 0 | 0.15 | 0.33 | 0.36 | 0.1 | 0.25 |
| 668 | MAGDALENA | PLATO | Caribbean | 91.33 | 0 | 0.57 | 0.06 | 0.4 | 0.51 | 0.59 | 0.44 | 0.15 |
| 669 | MAGDALENA | PUEBLOVIEJO | Caribbean | 95.83 | 0.17 | 0.54 | 0.03 | 0.32 | 0.38 | 0.43 | 0.33 | 0.1 |
| 670 | MAGDALENA | REMOLINO | Caribbean | 91.67 | 0 | 0.5 | 0 | 0.29 | 0.37 | 0.38 | 0.29 | 0.09 |
| 671 | MAGDALENA | SABANAS DE SAN ÁNGEL | Caribbean | 89.33 | 0.33 | 0.47 | 0.05 | 0.17 | 0.4 | 0.43 | 0.13 | 0.31 |
| 672 | MAGDALENA | SALAMINA | Caribbean | 124.5 | 0 | 0.57 | 0 | 0.2 | 0.38 | 0.43 | 0.17 | 0.26 |
| 673 | MAGDALENA | SAN SEBASTIÁN DE BUENAVISTA | Caribbean | 91.17 | 0.17 | 0.5 | 0.06 | 0.32 | 0.39 | 0.44 | 0.33 | 0.11 |
| 674 | MAGDALENA | SAN ZENÓN | Caribbean | 89 | 0 | 0.48 | 0 | 0.32 | 0.38 | 0.38 | 0.33 | 0.04 |
| 675 | MAGDALENA | SANTA ANA | Caribbean | 91.5 | 0 | 0.53 | 0 | 0.24 | 0.34 | 0.37 | 0.22 | 0.15 |
| 676 | MAGDALENA | SANTA BÁRBARA DE PINTO | Caribbean | 89.5 | 0 | 0.49 | 0 | 0.17 | 0.38 | 0.38 | 0.13 | 0.25 |
| 677 | MAGDALENA | SITIONUEVO | Caribbean | 92 | 0 | 0.49 | 0 | 0.33 | 0.47 | 0.46 | 0.34 | 0.12 |
| 678 | MAGDALENA | TENERIFE | Andean | 91.33 | 0.17 | 0.53 | 0.03 | 0.2 | 0.39 | 0.44 | 0.17 | 0.27 |
| 679 | MAGDALENA | ZAPAYÁN | Caribbean | 90 | 0 | 0.49 | 0 | 0.27 | 0.4 | 0.4 | 0.26 | 0.14 |
| 680 | MAGDALENA | ZONA BANANERA | Caribbean | 107.17 | 0.17 | 0.62 | 0.03 | 0.28 | 0.45 | 0.54 | 0.28 | 0.26 |
| 681 | MAGDALENA | ARACATACA | Caribbean | 112.17 | 1 | 0.68 | 0.25 | 0.24 | 0.4 | 0.74 | 0.23 | 0.52 |
| 682 | MAGDALENA | CIÉNAGA | Caribbean | 112.83 | 0.67 | 0.72 | 0.22 | 0.45 | 0.5 | 0.83 | 0.51 | 0.32 |
| 683 | MAGDALENA | FUNDACIÓN | Caribbean | 110.17 | 0.5 | 0.66 | 0.11 | 0.27 | 0.39 | 0.59 | 0.26 | 0.33 |
| 684 | MAGDALENA | SANTA MARTA | Caribbean | 112.17 | 1.67 | 0.72 | 0.23 | 0.7 | 0.55 | 0.88 | 0.84 | 0.04 |
| 685 | MAGDALENA | EL BANCO | Caribbean | 113.33 | 0.17 | 0.67 | 0.13 | 0.51 | 0.37 | 0.6 | 0.59 | 0.01 |
| 686 | META | VILLAVICENCIO | Orinoquia | 179 | 0.5 | 0.76 | 0.15 | 0.79 | 0.51 | 0.79 | 0.96 | -0.17 |
| 687 | META | ACACÍAS | Orinoquia | 157.83 | 0 | 0.72 | 0.06 | 0.46 | 0.36 | 0.56 | 0.51 | 0.04 |
| 688 | META | BARRANCA DE UPÍA | Orinoquia | 147.33 | 0 | 0.64 | 0 | 0.36 | 0.4 | 0.49 | 0.39 | 0.09 |
| 689 | META | CABUYARO | Orinoquia | 134.33 | 0 | 0.57 | 0 | 0.37 | 0.42 | 0.46 | 0.4 | 0.06 |
| 690 | META | CASTILLA LA NUEVA | Orinoquia | 158 | 0 | 0.65 | 0 | 0.39 | 0.42 | 0.51 | 0.43 | 0.09 |
| 691 | META | CUBARRAL | Orinoquia | 147.17 | 0.17 | 0.75 | 0.07 | 0.32 | 0.42 | 0.63 | 0.33 | 0.3 |
| 692 | META | CUMARAL | Orinoquia | 156.83 | 0 | 0.71 | 0.06 | 0.63 | 0.39 | 0.57 | 0.75 | -0.18 |
| 693 | META | EL CALVARIO | Orinoquia | 141.5 | 0 | 0.63 | 0 | 0.32 | 0.4 | 0.48 | 0.33 | 0.15 |
| 694 | META | EL CASTILLO | Orinoquia | 162.5 | 0 | 0.71 | 0 | 0.18 | 0.33 | 0.47 | 0.15 | 0.32 |
| 695 | META | EL DORADO | Orinoquia | 143 | 0 | 0.65 | 0 | 0.31 | 0.36 | 0.46 | 0.32 | 0.14 |
| 696 | META | FUENTE DE ORO | Orinoquia | 162.33 | 0.17 | 0.67 | 0.08 | 0.48 | 0.4 | 0.58 | 0.54 | 0.04 |
| 697 | META | GRANADA | Orinoquia | 162.33 | 0 | 0.64 | 0 | 0.43 | 0.55 | 0.62 | 0.48 | 0.13 |
| 698 | META | GUAMAL | Orinoquia | 156.67 | 0 | 0.72 | 0 | 0.29 | 0.32 | 0.47 | 0.29 | 0.17 |
| 699 | META | MESETAS | Orinoquia | 165.5 | 0.33 | 0.78 | 0.05 | 0.26 | 0.37 | 0.59 | 0.25 | 0.35 |
| 700 | META | URIBE | Orinoquia | 169.83 | 0.5 | 0.79 | 0.07 | 0.45 | 0.31 | 0.56 | 0.5 | 0.05 |
| 701 | META | LEJANÍAS | Orinoquia | 172 | 0.17 | 0.77 | 0.03 | 0.28 | 0.42 | 0.61 | 0.28 | 0.33 |
| 702 | META | PUERTO CONCORDIA | Orinoquia | 166.33 | 0.33 | 0.72 | 0.05 | 0.39 | 0.37 | 0.56 | 0.42 | 0.13 |
| 703 | META | PUERTO LÓPEZ | Orinoquia | 139.17 | 0.17 | 0.62 | 0.08 | 0.63 | 0.31 | 0.48 | 0.75 | -0.27 |
| 704 | META | PUERTO LLERAS | Orinoquia | 163 | 0 | 0.72 | 0 | 0.45 | 0.35 | 0.49 | 0.51 | -0.01 |
| 705 | META | PUERTO RICO | Orinoquia | 190.5 | 0 | 0.83 | 0 | 0.33 | 0.36 | 0.57 | 0.34 | 0.23 |
| 706 | META | RESTREPO | Orinoquia | 159.67 | 0 | 0.74 | 0 | 0.45 | 0.37 | 0.52 | 0.51 | 0.01 |
| 707 | META | SAN CARLOS DE GUAROA | Orinoquia | 133.33 | 0 | 0.5 | 0 | 0.38 | 0.39 | 0.39 | 0.41 | -0.02 |
| 708 | META | SAN JUAN DE ARAMA | Orinoquia | 168 | 0 | 0.73 | 0 | 0.28 | 0.36 | 0.5 | 0.28 | 0.22 |
| 709 | META | SAN JUANITO | Orinoquia | 137 | 0 | 0.62 | 0 | 0.21 | 0.51 | 0.57 | 0.19 | 0.38 |
| 710 | META | SAN MARTÍN | Orinoquia | 180 | 0.17 | 0.76 | 0.06 | 0.6 | 0.4 | 0.61 | 0.71 | -0.1 |
| 711 | META | VISTAHERMOSA | Orinoquia | 172.83 | 0 | 0.77 | 0 | 0.45 | 0.46 | 0.61 | 0.5 | 0.11 |
| 712 | META | LA MACARENA | Orinoquia | 182.67 | 0.33 | 0.78 | 0.16 | 0.53 | 0.38 | 0.7 | 0.62 | 0.09 |
| 713 | META | MAPIRIPÁN | Orinoquia | 165.5 | 1.5 | 0.7 | 0.11 | 0.36 | 0.39 | 0.62 | 0.38 | 0.24 |
| 714 | META | PUERTO GAITÁN | Orinoquia | 159 | 1.83 | 0.66 | 0.17 | 0.62 | 0.32 | 0.59 | 0.73 | -0.15 |
| 715 | NARIÑO | SANDONÁ | Pacific | 89.83 | 0 | 0.47 | 0 | 0.33 | 0.5 | 0.47 | 0.35 | 0.12 |
| 716 | NARIÑO | SAN BERNARDO | Pacific | 114.17 | 0 | 0.52 | 0 | 0.21 | 0.48 | 0.49 | 0.19 | 0.3 |
| 717 | NARIÑO | SAN LORENZO | Pacific | 93.17 | 0 | 0.54 | 0 | 0.2 | 0.37 | 0.41 | 0.17 | 0.23 |
| 718 | NARIÑO | POTOSÍ | Pacific | 144.33 | 0.33 | 0.68 | 0.05 | 0.3 | 0.32 | 0.49 | 0.3 | 0.19 |
| 719 | NARIÑO | SAN PABLO | Pacific | 129 | 0 | 0.55 | 0 | 0.25 | 0.37 | 0.41 | 0.23 | 0.17 |
| 720 | NARIÑO | PROVIDENCIA | Pacific | 85.83 | 0.17 | 0.19 | 0.06 | 0.17 | 0.44 | 0.31 | 0.13 | 0.18 |
| 721 | NARIÑO | SAN PEDRO DE CARTAGO | Pacific | 89.67 | 0 | 0.45 | 0 | 0.23 | 0.33 | 0.31 | 0.21 | 0.11 |
| 722 | NARIÑO | PUERRES | Pacific | 93.67 | 0.33 | 0.48 | 0.05 | 0.28 | 0.38 | 0.42 | 0.28 | 0.14 |
| 723 | NARIÑO | SANTACRUZ | Pacific | 107.17 | 0.5 | 0.55 | 0.07 | 0.18 | 0.28 | 0.4 | 0.14 | 0.26 |
| 724 | NARIÑO | PUPIALES | Pacific | 67.67 | 0.17 | 0.18 | 0.11 | 0.22 | 0.35 | 0.27 | 0.19 | 0.08 |
| 725 | NARIÑO | SAMANIEGO | Pacific | 110.33 | 0.83 | 0.54 | 0.11 | 0.17 | 0.37 | 0.49 | 0.13 | 0.36 |
| 726 | NARIÑO | CONSACÁ | Pacific | 88.17 | 0 | 0.49 | 0 | 0.34 | 0.49 | 0.47 | 0.35 | 0.12 |
| 727 | NARIÑO | CONTADERO | Pacific | 74.33 | 0 | 0.2 | 0 | 0.26 | 0.39 | 0.22 | 0.24 | -0.03 |
| 728 | NARIÑO | CÓRDOBA | Pacific | 120.83 | 0.5 | 0.56 | 0.07 | 0.28 | 0.44 | 0.54 | 0.27 | 0.27 |
| 729 | NARIÑO | PASTO | Pacific | 116.17 | 3.17 | 0.71 | 0.32 | 0.61 | 0.33 | 0.77 | 0.73 | 0.04 |
| 730 | NARIÑO | ALBÁN | Pacific | 125 | 0 | 0.51 | 0 | 0.17 | 0.41 | 0.41 | 0.13 | 0.28 |
| 731 | NARIÑO | ALDANA | Pacific | 60 | 0 | 0.09 | 0 | 0.33 | 0.36 | 0.12 | 0.34 | -0.22 |
| 732 | NARIÑO | CUASPUD CARLOSAMA | Pacific | 60.17 | 0.17 | 0.07 | 0.03 | 0.25 | 0.39 | 0.16 | 0.23 | -0.07 |
| 733 | NARIÑO | ANCUYA | Pacific | 88.83 | 0 | 0.4 | 0 | 0.23 | 0.32 | 0.28 | 0.21 | 0.07 |
| 734 | NARIÑO | ARBOLEDA | Pacific | 90 | 0.17 | 0.44 | 0.12 | 0.22 | 0.33 | 0.42 | 0.19 | 0.23 |
| 735 | NARIÑO | BELÉN | Pacific | 94.67 | 0 | 0.47 | 0 | 0.17 | 0.5 | 0.48 | 0.13 | 0.35 |
| 736 | NARIÑO | BUESACO | Pacific | 99 | 0.67 | 0.61 | 0.08 | 0.29 | 0.35 | 0.5 | 0.29 | 0.21 |
| 737 | NARIÑO | COLÓN | Amazon | 95.67 | 0 | 0.45 | 0 | 0.23 | 0.38 | 0.36 | 0.2 | 0.16 |
| 738 | NARIÑO | NARIÑO | Pacific | 149 | 0 | 0.62 | 0 | 0.26 | 0.39 | 0.47 | 0.25 | 0.22 |
| 739 | NARIÑO | OSPINA | Pacific | 72.83 | 0 | 0.18 | 0 | 0.21 | 0.4 | 0.21 | 0.19 | 0.02 |
| 740 | NARIÑO | SAPUYES | Pacific | 63.67 | 0 | 0.4 | 0 | 0.34 | 0.39 | 0.34 | 0.36 | -0.02 |
| 741 | NARIÑO | TAMINANGO | Pacific | 89.83 | 0 | 0.48 | 0 | 0.33 | 0.37 | 0.37 | 0.34 | 0.03 |
| 742 | NARIÑO | TANGUA | Pacific | 92.33 | 0.17 | 0.55 | 0.03 | 0.22 | 0.33 | 0.39 | 0.2 | 0.19 |
| 743 | NARIÑO | CHACHAGÜÍ | Pacific | 91.67 | 0.17 | 0.55 | 0.11 | 0.51 | 0.33 | 0.47 | 0.59 | -0.12 |
| 744 | NARIÑO | EL PEÑOL | Pacific | 89.83 | 0 | 0.46 | 0 | 0.32 | 0.4 | 0.38 | 0.33 | 0.05 |
| 745 | NARIÑO | EL TABLÓN DE GÓMEZ | Pacific | 98.33 | 0.5 | 0.56 | 0.07 | 0.26 | 0.28 | 0.39 | 0.25 | 0.14 |
| 746 | NARIÑO | TÚQUERRES | Andean | 86.83 | 0 | 0.46 | 0 | 0.32 | 0.4 | 0.38 | 0.33 | 0.04 |
| 747 | NARIÑO | LINARES | Pacific | 92.17 | 0 | 0.48 | 0 | 0.23 | 0.47 | 0.46 | 0.21 | 0.25 |
| 748 | NARIÑO | YACUANQUER | Pacific | 91.67 | 0 | 0.49 | 0 | 0.34 | 0.31 | 0.32 | 0.36 | -0.04 |
| 749 | NARIÑO | MALLAMA | Pacific | 106.67 | 0.5 | 0.55 | 0.07 | 0.28 | 0.34 | 0.44 | 0.28 | 0.16 |
| 750 | NARIÑO | LA CRUZ | Pacific | 89.5 | 0 | 0.49 | 0 | 0.19 | 0.31 | 0.32 | 0.16 | 0.16 |
| 751 | NARIÑO | LA FLORIDA | Pacific | 90.5 | 0 | 0.51 | 0 | 0.23 | 0.39 | 0.4 | 0.21 | 0.19 |
| 752 | NARIÑO | EL TAMBO | Pacific | 127.5 | 0 | 0.59 | 0 | 0.21 | 0.32 | 0.39 | 0.18 | 0.2 |
| 753 | NARIÑO | FUNES | Andean | 111 | 0.33 | 0.56 | 0.05 | 0.3 | 0.33 | 0.42 | 0.3 | 0.12 |
| 754 | NARIÑO | GUACHUCAL | Andean | 59 | 0.17 | 0.37 | 0.03 | 0.23 | 0.52 | 0.46 | 0.2 | 0.25 |
| 755 | NARIÑO | LA UNIÓN | Andean | 126.5 | 0 | 0.63 | 0 | 0.31 | 0.4 | 0.48 | 0.32 | 0.16 |
| 756 | NARIÑO | GUAITARILLA | Pacific | 87.17 | 0 | 0.43 | 0 | 0.21 | 0.37 | 0.34 | 0.19 | 0.15 |
| 757 | NARIÑO | LEIVA | Pacific | 104.67 | 0.5 | 0.58 | 0.07 | 0.29 | 0.36 | 0.48 | 0.29 | 0.19 |
| 758 | NARIÑO | GUALMATÁN | Pacific | 69 | 0 | 0.17 | 0 | 0.18 | 0.17 | 0 | 0.14 | -0.14 |
| 759 | NARIÑO | ILES | Pacific | 85.67 | 0.33 | 0.23 | 0.14 | 0.24 | 0.36 | 0.34 | 0.23 | 0.12 |
| 760 | NARIÑO | IMUÉS | Pacific | 89.17 | 0.17 | 0.43 | 0.12 | 0.24 | 0.33 | 0.41 | 0.22 | 0.19 |
| 761 | NARIÑO | IPIALES | Pacific | 165.5 | 1.17 | 0.77 | 0.14 | 0.3 | 0.32 | 0.63 | 0.3 | 0.32 |
| 762 | NARIÑO | CUMBAL | Pacific | 112.17 | 0.33 | 0.63 | 0.05 | 0.3 | 0.39 | 0.52 | 0.31 | 0.22 |
| 763 | NARIÑO | CUMBITARA | Pacific | 111.33 | 0.33 | 0.56 | 0.05 | 0.21 | 0.39 | 0.48 | 0.19 | 0.29 |
| 764 | NARIÑO | EL ROSARIO | Pacific | 110.83 | 0.17 | 0.58 | 0.03 | 0.2 | 0.36 | 0.44 | 0.17 | 0.26 |
| 765 | NARIÑO | LA LLANADA | Pacific | 112.67 | 0.17 | 0.57 | 0.03 | 0.21 | 0.28 | 0.36 | 0.18 | 0.17 |
| 766 | NARIÑO | LOS ANDES | Pacific | 115.83 | 0.17 | 0.62 | 0.03 | 0.27 | 0.34 | 0.44 | 0.26 | 0.19 |
| 767 | NARIÑO | POLICARPA | Pacific | 110.33 | 0.17 | 0.58 | 0.03 | 0.27 | 0.32 | 0.4 | 0.26 | 0.15 |
| 768 | NARIÑO | RICAURTE | Andean | 131.67 | 2.83 | 0.59 | 0.2 | 0.27 | 0.35 | 0.6 | 0.27 | 0.33 |
| 769 | NARIÑO | BARBACOAS | Pacific | 119.67 | 6 | 0.63 | 0.27 | 0.27 | 0.4 | 0.73 | 0.26 | 0.47 |
| 770 | NARIÑO | EL CHARCO | Pacific | 114.33 | 2.5 | 0.57 | 0.2 | 0.24 | 0.27 | 0.52 | 0.23 | 0.29 |
| 771 | NARIÑO | LA TOLA | Pacific | 89.67 | 1.5 | 0.38 | 0.15 | 0.38 | 0.36 | 0.44 | 0.42 | 0.03 |
| 772 | NARIÑO | MAGÜÍ | Pacific | 113.67 | 2.17 | 0.59 | 0.15 | 0.13 | 0.19 | 0.41 | 0.07 | 0.33 |
| 773 | NARIÑO | MOSQUERA | Pacific | 108.17 | 1.17 | 0.54 | 0.12 | 0.44 | 0.25 | 0.41 | 0.49 | -0.08 |
| 774 | NARIÑO | OLAYA HERRERA | Pacific | 91.17 | 1.83 | 0.46 | 0.16 | 0.34 | 0.22 | 0.37 | 0.36 | 0.01 |
| 775 | NARIÑO | FRANCISCO PIZARRO | Pacific | 96.17 | 0.33 | 0.5 | 0.05 | 0.27 | 0.31 | 0.37 | 0.26 | 0.11 |
| 776 | NARIÑO | ROBERTO PAYÁN | Pacific | 99.83 | 2.17 | 0.53 | 0.17 | 0.23 | 0.22 | 0.42 | 0.21 | 0.21 |
| 777 | NARIÑO | SANTA BÁRBARA | Pacific | 127.17 | 1.67 | 0.62 | 0.16 | 0.23 | 0.21 | 0.45 | 0.21 | 0.25 |
| 778 | NARIÑO | SAN ANDRÉS DE TUMACO | Pacific | 110 | 6.5 | 0.6 | 0.39 | 0.59 | 0.36 | 0.8 | 0.7 | 0.1 |
| 779 | NORTE DE SANTANDER | SAN JOSÉ DE CÚCUTA | Andean | 120 | 0.33 | 0.68 | 0.07 | 0.49 | 0.39 | 0.56 | 0.56 | 0.01 |
| 780 | NORTE DE SANTANDER | ÁBREGO | Andean | 129.83 | 0 | 0.7 | 0 | 0.24 | 0.38 | 0.5 | 0.23 | 0.27 |
| 781 | NORTE DE SANTANDER | ARBOLEDAS | Andean | 106 | 0 | 0.62 | 0 | 0.3 | 0.29 | 0.37 | 0.31 | 0.07 |
| 782 | NORTE DE SANTANDER | BOCHALEMA | Andean | 112.17 | 0 | 0.62 | 0 | 0.33 | 0.33 | 0.42 | 0.34 | 0.08 |
| 783 | NORTE DE SANTANDER | BUCARASICA | Andean | 104.67 | 0.17 | 0.48 | 0 | 0.18 | 0.36 | 0.35 | 0.15 | 0.21 |
| 784 | NORTE DE SANTANDER | CÁCOTA | Andean | 86.83 | 0 | 0.5 | 0 | 0.23 | 0.34 | 0.35 | 0.21 | 0.14 |
| 785 | NORTE DE SANTANDER | CÁCHIRA | Andean | 132.5 | 0 | 0.64 | 0 | 0.17 | 0.34 | 0.43 | 0.13 | 0.3 |
| 786 | NORTE DE SANTANDER | CHINÁCOTA | Andean | 103 | 0.17 | 0.56 | 0.1 | 0.34 | 0.35 | 0.5 | 0.36 | 0.13 |
| 787 | NORTE DE SANTANDER | CUCUTILLA | Andean | 102 | 0 | 0.54 | 0 | 0.29 | 0.33 | 0.37 | 0.28 | 0.08 |
| 788 | NORTE DE SANTANDER | DURANIA | Andean | 117 | 0 | 0.57 | 0 | 0.28 | 0.37 | 0.42 | 0.27 | 0.15 |
| 789 | NORTE DE SANTANDER | EL ZULIA | Andean | 118.33 | 0 | 0.62 | 0 | 0.19 | 0.33 | 0.41 | 0.16 | 0.26 |
| 790 | NORTE DE SANTANDER | GRAMALOTE | Andean | 113 | 0 | 0.56 | 0 | 0.21 | 0.36 | 0.4 | 0.18 | 0.22 |
| 791 | NORTE DE SANTANDER | HACARÍ | Andean | 108.5 | 0 | 0.61 | 0 | 0.11 | 0.36 | 0.43 | 0.04 | 0.39 |
| 792 | NORTE DE SANTANDER | HERRÁN | Andean | 99.17 | 0 | 0.54 | 0 | 0.2 | 0.33 | 0.36 | 0.17 | 0.19 |
| 793 | NORTE DE SANTANDER | LABATECA | Andean | 101.33 | 0 | 0.58 | 0 | 0.21 | 0.33 | 0.39 | 0.18 | 0.21 |
| 794 | NORTE DE SANTANDER | LA ESPERANZA | Andean | 133.67 | 0 | 0.64 | 0 | 0.17 | 0.34 | 0.43 | 0.12 | 0.31 |
| 795 | NORTE DE SANTANDER | LA PLAYA | Andean | 96.83 | 0 | 0.55 | 0 | 0.22 | 0.34 | 0.38 | 0.2 | 0.18 |
| 796 | NORTE DE SANTANDER | LOS PATIOS | Andean | 115.33 | 0 | 0.6 | 0 | 0.34 | 0.36 | 0.43 | 0.36 | 0.07 |
| 797 | NORTE DE SANTANDER | LOURDES | Andean | 104 | 0 | 0.5 | 0 | 0.21 | 0.3 | 0.32 | 0.18 | 0.14 |
| 798 | NORTE DE SANTANDER | MUTISCUA | Andean | 89 | 0 | 0.51 | 0 | 0.23 | 0.4 | 0.4 | 0.2 | 0.2 |
| 799 | NORTE DE SANTANDER | OCAÑA | Andean | 108.17 | 0.5 | 0.6 | 0.1 | 0.43 | 0.37 | 0.53 | 0.48 | 0.05 |
| 800 | NORTE DE SANTANDER | PAMPLONA | Andean | 99 | 0.83 | 0.6 | 0.12 | 0.37 | 0.35 | 0.54 | 0.4 | 0.14 |
| 801 | NORTE DE SANTANDER | PAMPLONITA | Andean | 102.33 | 0.17 | 0.58 | 0.1 | 0.32 | 0.34 | 0.49 | 0.33 | 0.17 |
| 802 | NORTE DE SANTANDER | PUERTO SANTANDER | Andean | 194.5 | 0 | 0.61 | 0 | 0.25 | 0.38 | 0.45 | 0.23 | 0.22 |
| 803 | NORTE DE SANTANDER | RAGONVALIA | Andean | 101.33 | 0 | 0.53 | 0 | 0.21 | 0.49 | 0.5 | 0.18 | 0.31 |
| 804 | NORTE DE SANTANDER | SALAZAR | Andean | 111 | 0 | 0.58 | 0 | 0.18 | 0.27 | 0.34 | 0.14 | 0.2 |
| 805 | NORTE DE SANTANDER | SAN CALIXTO | Andean | 123 | 0 | 0.64 | 0 | 0.16 | 0.29 | 0.39 | 0.12 | 0.28 |
| 806 | NORTE DE SANTANDER | SAN CAYETANO | Andean | 137.33 | 0 | 0.68 | 0 | 0.21 | 0.35 | 0.47 | 0.18 | 0.28 |
| 807 | NORTE DE SANTANDER | SANTIAGO | Andean | 119.17 | 0 | 0.57 | 0 | 0.13 | 0.33 | 0.38 | 0.07 | 0.31 |
| 808 | NORTE DE SANTANDER | SARDINATA | Andean | 121.67 | 0 | 0.6 | 0 | 0.13 | 0.33 | 0.4 | 0.07 | 0.34 |
| 809 | NORTE DE SANTANDER | SILOS | Andean | 77.67 | 0 | 0.47 | 0 | 0.3 | 0.34 | 0.33 | 0.3 | 0.03 |
| 810 | NORTE DE SANTANDER | VILLA CARO | Andean | 95.83 | 0 | 0.54 | 0 | 0.17 | 0.37 | 0.4 | 0.13 | 0.27 |
| 811 | NORTE DE SANTANDER | VILLA DEL ROSARIO | Andean | 112.67 | 0.5 | 0.58 | 0.13 | 0.36 | 0.53 | 0.69 | 0.38 | 0.31 |
| 812 | NORTE DE SANTANDER | CHITAGÁ | Andean | 114.5 | 0.17 | 0.67 | 0.03 | 0.25 | 0.31 | 0.45 | 0.24 | 0.22 |
| 813 | NORTE DE SANTANDER | CONVENCIÓN | Andean | 130.5 | 0.33 | 0.67 | 0.05 | 0.15 | 0.33 | 0.49 | 0.11 | 0.39 |
| 814 | NORTE DE SANTANDER | EL CARMEN | Andean | 132.67 | 0.17 | 0.69 | 0.03 | 0.21 | 0.32 | 0.47 | 0.18 | 0.29 |
| 815 | NORTE DE SANTANDER | TEORAMA | Andean | 131.17 | 0.33 | 0.64 | 0.05 | 0.16 | 0.34 | 0.48 | 0.11 | 0.37 |
| 816 | NORTE DE SANTANDER | TIBÚ | Andean | 130.67 | 0.17 | 0.64 | 0.03 | 0.28 | 0.42 | 0.53 | 0.27 | 0.25 |
| 817 | NORTE DE SANTANDER | TOLEDO | Andean | 138.17 | 0.33 | 0.73 | 0.05 | 0.26 | 0.36 | 0.56 | 0.25 | 0.31 |
| 818 | NORTE DE SANTANDER | EL TARRA | Andean | 129.83 | 0.17 | 0.62 | 0.03 | 0.16 | 0.32 | 0.43 | 0.11 | 0.32 |
| 819 | QUINDIO | ARMENIA | Andean | 118.33 | 0.83 | 0.52 | 0.26 | 0.67 | 0.43 | 0.69 | 0.8 | -0.11 |
| 820 | QUINDIO | BUENAVISTA | Andean | 131.33 | 0.33 | 0.55 | 0.19 | 0.51 | 0.55 | 0.75 | 0.59 | 0.16 |
| 821 | QUINDIO | CALARCÁ | Andean | 117.33 | 0.83 | 0.58 | 0.3 | 0.55 | 0.41 | 0.74 | 0.65 | 0.09 |
| 822 | QUINDIO | CIRCASIA | Andean | 115.5 | 0.33 | 0.5 | 0.25 | 0.47 | 0.42 | 0.66 | 0.53 | 0.12 |
| 823 | QUINDIO | CÓRDOBA | Andean | 120.67 | 0.33 | 0.55 | 0.16 | 0.36 | 0.36 | 0.54 | 0.38 | 0.17 |
| 824 | QUINDIO | FILANDIA | Andean | 114 | 0.5 | 0.53 | 0.26 | 0.49 | 0.59 | 0.84 | 0.56 | 0.27 |
| 825 | QUINDIO | GÉNOVA | Andean | 108.5 | 0 | 0.56 | 0 | 0.43 | 0.48 | 0.51 | 0.48 | 0.03 |
| 826 | QUINDIO | LA TEBAIDA | Andean | 102.5 | 0.17 | 0.51 | 0.13 | 0.47 | 0.43 | 0.56 | 0.53 | 0.03 |
| 827 | QUINDIO | MONTENEGRO | Andean | 108.17 | 0.17 | 0.49 | 0.13 | 0.55 | 0.42 | 0.54 | 0.65 | -0.1 |
| 828 | QUINDIO | PIJAO | Andean | 113.5 | 0.33 | 0.54 | 0.16 | 0.43 | 0.38 | 0.55 | 0.48 | 0.08 |
| 829 | QUINDIO | QUIMBAYA | Andean | 112.83 | 0.17 | 0.51 | 0.19 | 0.6 | 0.4 | 0.59 | 0.71 | -0.12 |
| 830 | QUINDIO | SALENTO | Andean | 117 | 0.17 | 0.6 | 0.13 | 0.5 | 0.55 | 0.72 | 0.57 | 0.15 |
| 831 | RISARALDA | PEREIRA | Andean | 118.67 | 1.17 | 0.67 | 0.24 | 0.67 | 0.55 | 0.87 | 0.8 | 0.06 |
| 832 | RISARALDA | APÍA | Andean | 122.5 | 0 | 0.58 | 0 | 0.38 | 0.35 | 0.41 | 0.41 | 0 |
| 833 | RISARALDA | BALBOA | Andean | 119.67 | 0 | 0.56 | 0 | 0.34 | 0.52 | 0.54 | 0.35 | 0.19 |
| 834 | RISARALDA | BELÉN DE UMBRÍA | Andean | 122.67 | 0.5 | 0.58 | 0.26 | 0.32 | 0.32 | 0.63 | 0.33 | 0.3 |
| 835 | RISARALDA | DOSQUEBRADAS | Andean | 114.83 | 0.33 | 0.52 | 0.31 | 0.55 | 0.41 | 0.71 | 0.64 | 0.08 |
| 836 | RISARALDA | GUÁTICA | Andean | 120.67 | 0 | 0.51 | 0 | 0.33 | 0.32 | 0.34 | 0.34 | 0 |
| 837 | RISARALDA | LA CELIA | Andean | 121.5 | 0 | 0.57 | 0 | 0.31 | 0.37 | 0.42 | 0.32 | 0.1 |
| 838 | RISARALDA | LA VIRGINIA | Andean | 105 | 0 | 0.47 | 0 | 0.47 | 0.57 | 0.54 | 0.53 | 0 |
| 839 | RISARALDA | MARSELLA | Andean | 113.5 | 0.83 | 0.52 | 0.31 | 0.46 | 0.39 | 0.69 | 0.52 | 0.17 |
| 840 | RISARALDA | MISTRATÓ | Andean | 132.83 | 0.83 | 0.65 | 0.21 | 0.32 | 0.32 | 0.62 | 0.33 | 0.3 |
| 841 | RISARALDA | PUEBLO RICO | Andean | 136 | 1 | 0.65 | 0.14 | 0.34 | 0.42 | 0.64 | 0.36 | 0.27 |
| 842 | RISARALDA | QUINCHÍA | Andean | 121.5 | 0.5 | 0.55 | 0.27 | 0.33 | 0.33 | 0.62 | 0.35 | 0.27 |
| 843 | RISARALDA | SANTA ROSA DE CABAL | Andean | 118.17 | 0.33 | 0.67 | 0.21 | 0.47 | 0.4 | 0.71 | 0.53 | 0.17 |
| 844 | RISARALDA | SANTUARIO | Andean | 122.83 | 0 | 0.61 | 0 | 0.43 | 0.36 | 0.43 | 0.48 | -0.05 |
| 845 | SANTANDER | AGUADA | Andean | 98.17 | 0.17 | 0.47 | 0 | 0.22 | 0.36 | 0.35 | 0.19 | 0.16 |
| 846 | SANTANDER | ALBANIA | Andean | 161.67 | 0 | 0.7 | 0 | 0.2 | 0.23 | 0.37 | 0.17 | 0.19 |
| 847 | SANTANDER | ARATOCA | Andean | 130.83 | 0 | 0.68 | 0 | 0.35 | 0.49 | 0.59 | 0.37 | 0.22 |
| 848 | SANTANDER | BARBOSA | Andean | 128.83 | 0 | 0.52 | 0 | 0.37 | 0.41 | 0.42 | 0.4 | 0.03 |
| 849 | SANTANDER | BARICHARA | Andean | 135 | 0.33 | 0.64 | 0.12 | 0.37 | 0.55 | 0.73 | 0.4 | 0.33 |
| 850 | SANTANDER | BARRANCABERMEJA | Andean | 124.83 | 0.17 | 0.65 | 0.06 | 0.75 | 0.43 | 0.57 | 0.91 | -0.34 |
| 851 | SANTANDER | BETULIA | Andean | 151.67 | 0 | 0.75 | 0 | 0.42 | 0.37 | 0.52 | 0.47 | 0.05 |
| 852 | SANTANDER | CABRERA | Andean | 135.67 | 0 | 0.61 | 0 | 0.25 | 0.52 | 0.57 | 0.24 | 0.33 |
| 853 | SANTANDER | CALIFORNIA | Andean | 90 | 0 | 0.47 | 0 | 0.34 | 0.48 | 0.46 | 0.36 | 0.09 |
| 854 | SANTANDER | CAPITANEJO | Andean | 96 | 0 | 0.49 | 0 | 0.24 | 0.51 | 0.5 | 0.22 | 0.27 |
| 855 | SANTANDER | CARCASÍ | Andean | 92.67 | 0 | 0.48 | 0 | 0.12 | 0.3 | 0.3 | 0.06 | 0.25 |
| 856 | SANTANDER | CEPITÁ | Andean | 128.33 | 0 | 0.66 | 0 | 0.23 | 0.35 | 0.45 | 0.21 | 0.24 |
| 857 | SANTANDER | CERRITO | Andean | 83.33 | 0 | 0.55 | 0 | 0.26 | 0.37 | 0.41 | 0.25 | 0.15 |
| 858 | SANTANDER | CHARALÁ | Andean | 101.5 | 0.33 | 0.59 | 0.1 | 0.31 | 0.38 | 0.53 | 0.32 | 0.21 |
| 859 | SANTANDER | CHARTA | Andean | 125.5 | 0 | 0.62 | 0 | 0.2 | 0.38 | 0.46 | 0.17 | 0.29 |
| 860 | SANTANDER | CHIMA | Andean | 100.17 | 0 | 0.5 | 0 | 0.21 | 0.36 | 0.37 | 0.18 | 0.19 |
| 861 | SANTANDER | CHIPATÁ | Andean | 97.33 | 0 | 0.46 | 0 | 0.23 | 0.33 | 0.31 | 0.21 | 0.1 |
| 862 | SANTANDER | CIMITARRA | Andean | 146 | 0 | 0.73 | 0 | 0.36 | 0.36 | 0.5 | 0.39 | 0.11 |
| 863 | SANTANDER | CONCEPCIÓN | Andean | 131.17 | 0 | 0.67 | 0 | 0.11 | 0.36 | 0.47 | 0.05 | 0.42 |
| 864 | SANTANDER | CONFINES | Andean | 97.83 | 0 | 0.43 | 0 | 0.25 | 0.38 | 0.35 | 0.23 | 0.11 |
| 865 | SANTANDER | RIONEGRO | Andean | 142.17 | 0 | 0.74 | 0 | 0.37 | 0.38 | 0.53 | 0.39 | 0.14 |
| 866 | SANTANDER | SABANA DE TORRES | Andean | 132.67 | 0 | 0.72 | 0 | 0.48 | 0.41 | 0.54 | 0.54 | 0 |
| 867 | SANTANDER | SAN ANDRÉS | Andean | 127.5 | 0 | 0.69 | 0 | 0.2 | 0.35 | 0.47 | 0.17 | 0.3 |
| 868 | SANTANDER | SAN BENITO | Andean | 92 | 0 | 0.41 | 0 | 0.23 | 0.23 | 0.2 | 0.21 | -0.01 |
| 869 | SANTANDER | SAN GIL | Andean | 132.83 | 0 | 0.66 | 0 | 0.46 | 0.44 | 0.53 | 0.52 | 0.01 |
| 870 | SANTANDER | SAN JOAQUÍN | Andean | 98.33 | 0 | 0.53 | 0 | 0.21 | 0.38 | 0.4 | 0.18 | 0.22 |
| 871 | SANTANDER | SAN JOSÉ DE MIRANDA | Andean | 98.67 | 0 | 0.55 | 0 | 0.24 | 0.35 | 0.39 | 0.22 | 0.16 |
| 872 | SANTANDER | SAN MIGUEL | Andean | 170 | 0 | 0.73 | 0 | 0.16 | 0.34 | 0.49 | 0.11 | 0.38 |
| 873 | SANTANDER | SAN VICENTE DE CHUCURÍ | Andean | 150.83 | 0.33 | 0.73 | 0.07 | 0.52 | 0.34 | 0.55 | 0.6 | -0.06 |
| 874 | SANTANDER | SANTA BÁRBARA | Andean | 127.5 | 0 | 0.65 | 0 | 0.3 | 0.52 | 0.6 | 0.3 | 0.3 |
| 875 | SANTANDER | SANTA HELENA DEL OPÓN | Andean | 134 | 0 | 0.6 | 0 | 0.17 | 0.35 | 0.42 | 0.12 | 0.3 |
| 876 | SANTANDER | SIMACOTA | Andean | 140.17 | 0 | 0.78 | 0 | 0.28 | 0.34 | 0.51 | 0.27 | 0.24 |
| 877 | SANTANDER | SOCORRO | Andean | 129.17 | 0 | 0.59 | 0 | 0.36 | 0.38 | 0.44 | 0.39 | 0.05 |
| 878 | SANTANDER | SUAITA | Andean | 97.83 | 0 | 0.49 | 0 | 0.22 | 0.39 | 0.39 | 0.2 | 0.19 |
| 879 | SANTANDER | SUCRE | Andean | 138.33 | 0 | 0.67 | 0 | 0.19 | 0.36 | 0.47 | 0.15 | 0.32 |
| 880 | SANTANDER | SURATÁ | Andean | 97.83 | 0 | 0.54 | 0 | 0.19 | 0.37 | 0.41 | 0.15 | 0.25 |
| 881 | SANTANDER | TONA | Andean | 128.67 | 0 | 0.66 | 0 | 0.22 | 0.38 | 0.48 | 0.19 | 0.29 |
| 882 | SANTANDER | VALLE DE SAN JOSÉ | Andean | 99.83 | 0 | 0.51 | 0 | 0.24 | 0.48 | 0.48 | 0.22 | 0.26 |
| 883 | SANTANDER | VÉLEZ | Andean | 130.67 | 0 | 0.57 | 0.06 | 0.34 | 0.42 | 0.51 | 0.36 | 0.16 |
| 884 | SANTANDER | VETAS | Andean | 74.83 | 0 | 0.42 | 0 | 0.33 | 0.39 | 0.35 | 0.35 | 0 |
| 885 | SANTANDER | VILLANUEVA | Andean | 178.67 | 0.17 | 0.7 | 0.06 | 0.38 | 0.39 | 0.57 | 0.41 | 0.16 |
| 886 | SANTANDER | ZAPATOCA | Andean | 141.33 | 0.17 | 0.69 | 0.09 | 0.52 | 0.39 | 0.59 | 0.6 | 0 |
| 887 | SANTANDER | JORDÁN | Andean | 130.33 | 0 | 0.6 | 0 | 0.27 | 0.39 | 0.46 | 0.26 | 0.2 |
| 888 | SANTANDER | LA BELLEZA | Andean | 135 | 0 | 0.59 | 0 | 0.19 | 0.4 | 0.46 | 0.15 | 0.3 |
| 889 | SANTANDER | LANDÁZURI | Andean | 138.33 | 0 | 0.62 | 0 | 0.27 | 0.32 | 0.4 | 0.27 | 0.13 |
| 890 | SANTANDER | LA PAZ | Andean | 133.67 | 0 | 0.59 | 0 | 0.12 | 0.36 | 0.42 | 0.05 | 0.36 |
| 891 | SANTANDER | LEBRIJA | Andean | 150 | 0 | 0.76 | 0 | 0.52 | 0.36 | 0.52 | 0.61 | -0.09 |
| 892 | SANTANDER | LOS SANTOS | Andean | 136 | 0 | 0.67 | 0 | 0.34 | 0.38 | 0.49 | 0.36 | 0.13 |
| 893 | SANTANDER | MACARAVITA | Andean | 90.83 | 0 | 0.49 | 0 | 0.2 | 0.42 | 0.42 | 0.17 | 0.25 |
| 894 | SANTANDER | MÁLAGA | Andean | 92.83 | 0 | 0.48 | 0 | 0.48 | 0.4 | 0.4 | 0.55 | -0.15 |
| 895 | SANTANDER | MATANZA | Andean | 128.67 | 0 | 0.66 | 0 | 0.21 | 0.4 | 0.5 | 0.19 | 0.32 |
| 896 | SANTANDER | MOGOTES | Andean | 101.83 | 0 | 0.57 | 0 | 0.2 | 0.33 | 0.38 | 0.17 | 0.21 |
| 897 | SANTANDER | MOLAGAVITA | Andean | 126.83 | 0 | 0.67 | 0 | 0.2 | 0.37 | 0.48 | 0.17 | 0.31 |
| 898 | SANTANDER | OCAMONTE | Andean | 99.5 | 0 | 0.49 | 0 | 0.23 | 0.37 | 0.38 | 0.2 | 0.17 |
| 899 | SANTANDER | OIBA | Andean | 100.33 | 0.17 | 0.51 | 0.09 | 0.23 | 0.41 | 0.5 | 0.2 | 0.3 |
| 900 | SANTANDER | ONZAGA | Andean | 99.67 | 0 | 0.58 | 0 | 0.18 | 0.38 | 0.44 | 0.15 | 0.29 |
| 901 | SANTANDER | PALMAR | Andean | 122.33 | 0 | 0.52 | 0 | 0.27 | 0.38 | 0.4 | 0.26 | 0.14 |
| 902 | SANTANDER | PALMAS DEL SOCORRO | Andean | 96.67 | 0 | 0.44 | 0 | 0.34 | 0.38 | 0.35 | 0.35 | 0 |
| 903 | SANTANDER | PÁRAMO | Andean | 97.83 | 0 | 0.43 | 0 | 0.26 | 0.43 | 0.39 | 0.25 | 0.14 |
| 904 | SANTANDER | PINCHOTE | Andean | 131.5 | 0 | 0.59 | 0.06 | 0.38 | 0.42 | 0.53 | 0.42 | 0.12 |
| 905 | SANTANDER | PUENTE NACIONAL | Andean | 93.83 | 0 | 0.49 | 0 | 0.23 | 0.39 | 0.39 | 0.21 | 0.18 |
| 906 | SANTANDER | PUERTO PARRA | Andean | 127.17 | 0 | 0.66 | 0 | 0.35 | 0.43 | 0.52 | 0.37 | 0.15 |
| 907 | SANTANDER | PUERTO WILCHES | Andean | 125.67 | 0 | 0.68 | 0 | 0.47 | 0.37 | 0.49 | 0.53 | -0.05 |
| 908 | SANTANDER | CONTRATACIÓN | Andean | 100.5 | 0 | 0.51 | 0 | 0.23 | 0.39 | 0.4 | 0.21 | 0.19 |
| 909 | SANTANDER | COROMORO | Andean | 100.83 | 0 | 0.59 | 0 | 0.18 | 0.33 | 0.39 | 0.14 | 0.25 |
| 910 | SANTANDER | CURITÍ | Andean | 130.33 | 0 | 0.72 | 0 | 0.24 | 0.38 | 0.52 | 0.22 | 0.3 |
| 911 | SANTANDER | EL CARMEN DE CHUCURÍ | Andean | 144.5 | 0 | 0.72 | 0 | 0.16 | 0.36 | 0.49 | 0.12 | 0.37 |
| 912 | SANTANDER | EL GUACAMAYO | Andean | 99.83 | 0 | 0.49 | 0 | 0.2 | 0.5 | 0.49 | 0.17 | 0.31 |
| 913 | SANTANDER | EL PLAYÓN | Andean | 134.17 | 0 | 0.64 | 0 | 0.19 | 0.35 | 0.45 | 0.15 | 0.29 |
| 914 | SANTANDER | ENCINO | Andean | 99.67 | 0 | 0.55 | 0 | 0.19 | 0.55 | 0.57 | 0.16 | 0.41 |
| 915 | SANTANDER | ENCISO | Andean | 93.33 | 0 | 0.29 | 0 | 0.21 | 0.4 | 0.28 | 0.18 | 0.1 |
| 916 | SANTANDER | FLORIÁN | Andean | 130.17 | 0 | 0.62 | 0 | 0.22 | 0.4 | 0.48 | 0.19 | 0.29 |
| 917 | SANTANDER | GALÁN | Andean | 133.17 | 0.17 | 0.67 | 0 | 0.31 | 0.31 | 0.43 | 0.32 | 0.11 |
| 918 | SANTANDER | GÁMBITA | Andean | 97.33 | 0 | 0.56 | 0 | 0.28 | 0.5 | 0.53 | 0.28 | 0.25 |
| 919 | SANTANDER | GUACA | Andean | 98.5 | 0 | 0.55 | 0 | 0.2 | 0.46 | 0.49 | 0.17 | 0.32 |
| 920 | SANTANDER | GUADALUPE | Andean | 154.33 | 0.17 | 0.6 | 0.11 | 0.32 | 0.35 | 0.52 | 0.33 | 0.19 |
| 921 | SANTANDER | GUAPOTÁ | Andean | 95.83 | 0 | 0.44 | 0 | 0.24 | 0.39 | 0.35 | 0.22 | 0.13 |
| 922 | SANTANDER | GUAVATÁ | Andean | 92.17 | 0 | 0.36 | 0 | 0.23 | 0.38 | 0.3 | 0.2 | 0.1 |
| 923 | SANTANDER | GÜEPSA | Andean | 93.17 | 0 | 0.38 | 0 | 0.28 | 0.37 | 0.31 | 0.27 | 0.03 |
| 924 | SANTANDER | HATO | Andean | 106 | 0 | 0.52 | 0 | 0.21 | 0.45 | 0.46 | 0.18 | 0.28 |
| 925 | SANTANDER | JESÚS MARÍA | Andean | 97.17 | 0 | 0.47 | 0 | 0.23 | 0.33 | 0.32 | 0.2 | 0.12 |
| 926 | SANTANDER | GIRÓN | Andean | 151 | 0 | 0.72 | 0 | 0.43 | 0.41 | 0.54 | 0.48 | 0.06 |
| 927 | SANTANDER | FLORIDABLANCA | Andean | 129.33 | 0.17 | 0.61 | 0.11 | 0.39 | 0.45 | 0.62 | 0.43 | 0.19 |
| 928 | SANTANDER | BOLÍVAR | Andean | 151.33 | 0 | 0.68 | 0 | 0.28 | 0.34 | 0.45 | 0.28 | 0.17 |
| 929 | SANTANDER | EL PEÑÓN | Andean | 140.5 | 0 | 0.61 | 0 | 0.27 | 0.31 | 0.39 | 0.27 | 0.12 |
| 930 | SANTANDER | PIEDECUESTA | Andean | 135.67 | 0.33 | 0.75 | 0.14 | 0.49 | 0.54 | 0.81 | 0.55 | 0.25 |
| 931 | SANTANDER | BUCARAMANGA | Andean | 134.33 | 1.5 | 0.62 | 0.15 | 0.41 | 0.48 | 0.68 | 0.46 | 0.22 |
| 932 | SUCRE | SINCELEJO | Caribbean | 92.67 | 0.67 | 0.46 | 0.3 | 0.4 | 0.39 | 0.65 | 0.44 | 0.21 |
| 933 | SUCRE | BUENAVISTA | Caribbean | 131.33 | 0 | 0.55 | 0.06 | 0.32 | 0.4 | 0.49 | 0.33 | 0.16 |
| 934 | SUCRE | CAIMITO | Caribbean | 111 | 0 | 0.63 | 0 | 0.29 | 0.41 | 0.48 | 0.28 | 0.2 |
| 935 | SUCRE | COLOSÓ | Caribbean | 93.67 | 0.17 | 0.47 | 0.03 | 0.31 | 0.35 | 0.37 | 0.32 | 0.05 |
| 936 | SUCRE | COROZAL | Caribbean | 91.67 | 0 | 0.44 | 0 | 0.36 | 0.4 | 0.36 | 0.39 | -0.02 |
| 937 | SUCRE | COVEÑAS | Caribbean | 89.5 | 0.17 | 0.46 | 0.12 | 0.38 | 0.41 | 0.51 | 0.42 | 0.09 |
| 938 | SUCRE | CHALÁN | Caribbean | 93 | 0 | 0.46 | 0 | 0.31 | 0.34 | 0.33 | 0.32 | 0.01 |
| 939 | SUCRE | EL ROBLE | Caribbean | 108.83 | 0 | 0.52 | 0 | 0.13 | 0.35 | 0.38 | 0.07 | 0.31 |
| 940 | SUCRE | GALERAS | Caribbean | 108.83 | 0 | 0.54 | 0 | 0.19 | 0.38 | 0.4 | 0.16 | 0.25 |
| 941 | SUCRE | GUARANDA | Caribbean | 113.33 | 0 | 0.57 | 0 | 0.18 | 0.19 | 0.25 | 0.14 | 0.11 |
| 942 | SUCRE | LA UNIÓN | Caribbean | 126.33 | 0 | 0.61 | 0 | 0.12 | 0.48 | 0.54 | 0.06 | 0.48 |
| 943 | SUCRE | LOS PALMITOS | Caribbean | 91.67 | 0 | 0.44 | 0 | 0.21 | 0.4 | 0.37 | 0.18 | 0.19 |
| 944 | SUCRE | MAJAGUAL | Caribbean | 112.17 | 0 | 0.55 | 0 | 0.14 | 0.21 | 0.26 | 0.09 | 0.17 |
| 945 | SUCRE | OVEJAS | Caribbean | 92.5 | 0.33 | 0.47 | 0.14 | 0.33 | 0.39 | 0.51 | 0.34 | 0.17 |
| 946 | SUCRE | PALMITO | Caribbean | 90.5 | 0.17 | 0.42 | 0.03 | 0.19 | 0.37 | 0.36 | 0.15 | 0.2 |
| 947 | SUCRE | SAMPUÉS | Caribbean | 90.83 | 0.17 | 0.42 | 0.03 | 0.22 | 0.4 | 0.38 | 0.19 | 0.19 |
| 948 | SUCRE | SAN BENITO ABAD | Caribbean | 113.5 | 0 | 0.62 | 0 | 0.25 | 0.35 | 0.43 | 0.23 | 0.2 |
| 949 | SUCRE | SAN JUAN DE BETULIA | Caribbean | 91.5 | 0 | 0.42 | 0 | 0.22 | 0.38 | 0.34 | 0.19 | 0.15 |
| 950 | SUCRE | SAN MARCOS | Caribbean | 113.33 | 0 | 0.64 | 0 | 0.26 | 0.38 | 0.46 | 0.26 | 0.21 |
| 951 | SUCRE | SAN ONOFRE | Caribbean | 93.67 | 0 | 0.62 | 0 | 0.4 | 0.42 | 0.49 | 0.44 | 0.05 |
| 952 | SUCRE | SAN PEDRO | Caribbean | 117.17 | 0 | 0.54 | 0 | 0.18 | 0.39 | 0.42 | 0.14 | 0.27 |
| 953 | SUCRE | SAN LUIS DE SINCÉ | Caribbean | 109 | 0 | 0.48 | 0 | 0.2 | 0.38 | 0.38 | 0.17 | 0.21 |
| 954 | SUCRE | SUCRE | Caribbean | 137.67 | 0 | 0.6 | 0 | 0.27 | 0.29 | 0.36 | 0.26 | 0.1 |
| 955 | SUCRE | SANTIAGO DE TOLÚ | Caribbean | 91.83 | 0 | 0.51 | 0 | 0.37 | 0.33 | 0.35 | 0.4 | -0.05 |
| 956 | SUCRE | SAN JOSÉ DE TOLUVIEJO | Caribbean | 93.33 | 0.17 | 0.49 | 0.03 | 0.2 | 0.41 | 0.43 | 0.17 | 0.26 |
| 957 | SUCRE | MORROA | Caribbean | 93.67 | 0 | 0.46 | 0.06 | 0.33 | 0.38 | 0.42 | 0.35 | 0.07 |
| 958 | TOLIMA | IBAGUÉ | Andean | 127.67 | 1.17 | 0.67 | 0.21 | 0.69 | 0.54 | 0.83 | 0.83 | 0 |
| 959 | TOLIMA | ALPUJARRA | Andean | 121.67 | 0.17 | 0.67 | 0.03 | 0.19 | 0.37 | 0.51 | 0.16 | 0.35 |
| 960 | TOLIMA | ALVARADO | Andean | 117.67 | 0 | 0.55 | 0 | 0.4 | 0.43 | 0.46 | 0.43 | 0.02 |
| 961 | TOLIMA | AMBALEMA | Andean | 106.83 | 0 | 0.48 | 0 | 0.41 | 0.44 | 0.42 | 0.46 | -0.03 |
| 962 | TOLIMA | ANZOÁTEGUI | Andean | 120.17 | 0 | 0.57 | 0 | 0.3 | 0.39 | 0.43 | 0.31 | 0.12 |
| 963 | TOLIMA | ARMERO | Andean | 139.17 | 0 | 0.69 | 0 | 0.32 | 0.41 | 0.52 | 0.33 | 0.19 |
| 964 | TOLIMA | ATACO | Andean | 121.67 | 0.5 | 0.67 | 0.07 | 0.16 | 0.37 | 0.55 | 0.11 | 0.43 |
| 965 | TOLIMA | CAJAMARCA | Andean | 104 | 0 | 0.53 | 0 | 0.33 | 0.35 | 0.38 | 0.35 | 0.03 |
| 966 | TOLIMA | CARMEN DE APICALÁ | Andean | 120.17 | 0 | 0.56 | 0 | 0.33 | 0.43 | 0.47 | 0.34 | 0.13 |
| 967 | TOLIMA | CASABIANCA | Andean | 128.67 | 0 | 0.6 | 0 | 0.33 | 0.35 | 0.42 | 0.35 | 0.07 |
| 968 | TOLIMA | CHAPARRAL | Andean | 118.17 | 0.5 | 0.68 | 0.07 | 0.34 | 0.36 | 0.54 | 0.36 | 0.18 |
| 969 | TOLIMA | COELLO | Andean | 112.17 | 0 | 0.51 | 0 | 0.31 | 0.44 | 0.44 | 0.31 | 0.13 |
| 970 | TOLIMA | COYAIMA | Andean | 113.5 | 6.33 | 0.59 | 0.17 | 0.27 | 0.39 | 0.61 | 0.26 | 0.35 |
| 971 | TOLIMA | CUNDAY | Andean | 124.33 | 0 | 0.6 | 0 | 0.28 | 0.37 | 0.43 | 0.27 | 0.16 |
| 972 | TOLIMA | DOLORES | Andean | 121 | 0.17 | 0.67 | 0.03 | 0.18 | 0.33 | 0.47 | 0.14 | 0.33 |
| 973 | TOLIMA | ESPINAL | Andean | 105.83 | 0 | 0.49 | 0.06 | 0.54 | 0.45 | 0.49 | 0.62 | -0.13 |
| 974 | TOLIMA | FALAN | Andean | 141.67 | 0 | 0.59 | 0 | 0.33 | 0.38 | 0.44 | 0.34 | 0.1 |
| 975 | TOLIMA | FLANDES | Andean | 101.67 | 0 | 0.43 | 0 | 0.37 | 0.45 | 0.4 | 0.4 | 0 |
| 976 | TOLIMA | FRESNO | Andean | 141.83 | 0 | 0.61 | 0 | 0.33 | 0.35 | 0.43 | 0.35 | 0.08 |
| 977 | TOLIMA | GUAMO | Andean | 109 | 0 | 0.5 | 0 | 0.21 | 0.43 | 0.43 | 0.18 | 0.25 |
| 978 | TOLIMA | HERVEO | Andean | 131.83 | 0 | 0.64 | 0 | 0.43 | 0.35 | 0.44 | 0.48 | -0.03 |
| 979 | TOLIMA | ICONONZO | Andean | 120.67 | 0 | 0.57 | 0 | 0.33 | 0.39 | 0.44 | 0.35 | 0.09 |
| 980 | TOLIMA | LÉRIDA | Andean | 124.17 | 0 | 0.56 | 0 | 0.3 | 0.39 | 0.43 | 0.3 | 0.13 |
| 981 | TOLIMA | LÍBANO | Andean | 130.67 | 0 | 0.58 | 0 | 0.36 | 0.41 | 0.46 | 0.38 | 0.08 |
| 982 | TOLIMA | SAN SEBASTIÁN DE MARIQUITA | Andean | 144.5 | 0 | 0.61 | 0.06 | 0.69 | 0.37 | 0.5 | 0.83 | -0.33 |
| 983 | TOLIMA | MELGAR | Andean | 123.33 | 0 | 0.61 | 0 | 0.46 | 0.41 | 0.48 | 0.52 | -0.04 |
| 984 | TOLIMA | MURILLO | Andean | 117.67 | 0 | 0.65 | 0 | 0.32 | 0.42 | 0.51 | 0.33 | 0.18 |
| 985 | TOLIMA | NATAGAIMA | Andean | 121.5 | 5 | 0.63 | 0.16 | 0.36 | 0.34 | 0.57 | 0.39 | 0.19 |
| 986 | TOLIMA | ORTEGA | Andean | 119 | 4 | 0.64 | 0.15 | 0.18 | 0.38 | 0.61 | 0.15 | 0.46 |
| 987 | TOLIMA | PALOCABILDO | Andean | 138.5 | 0 | 0.61 | 0 | 0.23 | 0.41 | 0.47 | 0.21 | 0.26 |
| 988 | TOLIMA | PIEDRAS | Andean | 113 | 0.17 | 0.51 | 0 | 0.4 | 0.37 | 0.38 | 0.44 | -0.07 |
| 989 | TOLIMA | PLANADAS | Andean | 108 | 0.17 | 0.6 | 0.03 | 0.27 | 0.4 | 0.49 | 0.26 | 0.23 |
| 990 | TOLIMA | PRADO | Andean | 123.17 | 0.5 | 0.63 | 0.14 | 0.31 | 0.39 | 0.6 | 0.32 | 0.28 |
| 991 | TOLIMA | PURIFICACIÓN | Andean | 123.5 | 0 | 0.65 | 0 | 0.31 | 0.39 | 0.48 | 0.32 | 0.16 |
| 992 | TOLIMA | RIOBLANCO | Andean | 116.83 | 0.5 | 0.65 | 0.07 | 0.24 | 0.37 | 0.53 | 0.22 | 0.32 |
| 993 | TOLIMA | RONCESVALLES | Andean | 115.17 | 0 | 0.63 | 0 | 0.31 | 0.36 | 0.44 | 0.32 | 0.12 |
| 994 | TOLIMA | ROVIRA | Andean | 122.67 | 0 | 0.67 | 0 | 0.27 | 0.46 | 0.56 | 0.26 | 0.3 |
| 995 | TOLIMA | SALDAÑA | Andean | 100.33 | 0 | 0.47 | 0 | 0.3 | 0.38 | 0.37 | 0.3 | 0.07 |
| 996 | TOLIMA | SAN ANTONIO | Andean | 112.67 | 0.17 | 0.6 | 0.03 | 0.3 | 0.4 | 0.49 | 0.31 | 0.18 |
| 997 | TOLIMA | SAN LUIS | Andean | 141 | 0 | 0.64 | 0 | 0.37 | 0.38 | 0.47 | 0.39 | 0.07 |
| 998 | TOLIMA | SANTA ISABEL | Andean | 124.67 | 0 | 0.62 | 0 | 0.31 | 0.52 | 0.58 | 0.31 | 0.27 |
| 999 | TOLIMA | SUÁREZ | Andean | 124.67 | 0 | 0.57 | 0 | 0.22 | 0.36 | 0.41 | 0.2 | 0.21 |
| 1000 | TOLIMA | VALLE DE SAN JUAN | Andean | 120.33 | 0 | 0.61 | 0 | 0.31 | 0.53 | 0.58 | 0.31 | 0.27 |
| 1001 | TOLIMA | VENADILLO | Andean | 126.67 | 0 | 0.59 | 0 | 0.3 | 0.4 | 0.46 | 0.3 | 0.16 |
| 1002 | TOLIMA | VILLAHERMOSA | Andean | 138.33 | 0 | 0.7 | 0 | 0.32 | 0.41 | 0.53 | 0.33 | 0.2 |
| 1003 | TOLIMA | VILLARRICA | Andean | 118.83 | 0 | 0.59 | 0 | 0.21 | 0.33 | 0.4 | 0.18 | 0.22 |
| 1004 | TOLIMA | HONDA | Andean | 137 | 0.33 | 0.62 | 0.1 | 0.53 | 0.35 | 0.53 | 0.61 | -0.08 |
| 1005 | VALLE DEL CAUCA | CALI | Pacific | 125.17 | 4.17 | 0.63 | 0.31 | 0.82 | 0.54 | 0.9 | 1 | -0.1 |
| 1006 | VALLE DEL CAUCA | ANDALUCÍA | Pacific | 109.67 | 0 | 0.51 | 0 | 0.34 | 0.39 | 0.4 | 0.36 | 0.04 |
| 1007 | VALLE DEL CAUCA | ANSERMANUEVO | Pacific | 124 | 0.17 | 0.59 | 0.03 | 0.41 | 0.39 | 0.48 | 0.46 | 0.02 |
| 1008 | VALLE DEL CAUCA | ARGELIA | Pacific | 149.17 | 0.17 | 0.6 | 0.03 | 0.21 | 0.39 | 0.48 | 0.19 | 0.3 |
| 1009 | VALLE DEL CAUCA | GUADALAJARA DE BUGA | Pacific | 116.5 | 0.5 | 0.69 | 0.09 | 0.54 | 0.5 | 0.69 | 0.62 | 0.06 |
| 1010 | VALLE DEL CAUCA | BUGALAGRANDE | Pacific | 113 | 0.33 | 0.57 | 0.05 | 0.4 | 0.35 | 0.45 | 0.43 | 0.02 |
| 1011 | VALLE DEL CAUCA | CAICEDONIA | Pacific | 108.83 | 0.17 | 0.54 | 0.03 | 0.34 | 0.43 | 0.48 | 0.36 | 0.12 |
| 1012 | VALLE DEL CAUCA | CALIMA | Pacific | 146.33 | 0.83 | 0.69 | 0.17 | 0.45 | 0.47 | 0.73 | 0.51 | 0.23 |
| 1013 | VALLE DEL CAUCA | CANDELARIA | Pacific | 103.5 | 0 | 0.46 | 0 | 0.53 | 0.41 | 0.38 | 0.62 | -0.23 |
| 1014 | VALLE DEL CAUCA | CARTAGO | Pacific | 112.17 | 0.33 | 0.53 | 0.11 | 0.58 | 0.38 | 0.51 | 0.68 | -0.17 |
| 1015 | VALLE DEL CAUCA | EL ÁGUILA | Pacific | 124.17 | 0 | 0.59 | 0 | 0.3 | 0.35 | 0.41 | 0.3 | 0.11 |
| 1016 | VALLE DEL CAUCA | EL CAIRO | Pacific | 119.17 | 0.33 | 0.51 | 0.05 | 0.34 | 0.52 | 0.57 | 0.36 | 0.21 |
| 1017 | VALLE DEL CAUCA | EL CERRITO | Pacific | 113.67 | 0.33 | 0.63 | 0.11 | 0.45 | 0.38 | 0.57 | 0.5 | 0.07 |
| 1018 | VALLE DEL CAUCA | EL DOVIO | Pacific | 127.83 | 0.33 | 0.61 | 0.05 | 0.22 | 0.45 | 0.56 | 0.19 | 0.37 |
| 1019 | VALLE DEL CAUCA | FLORIDA | Andean | 113.5 | 0.67 | 0.63 | 0.08 | 0.29 | 0.38 | 0.54 | 0.28 | 0.26 |
| 1020 | VALLE DEL CAUCA | GINEBRA | Andean | 112.67 | 0.17 | 0.54 | 0.08 | 0.46 | 0.34 | 0.45 | 0.52 | -0.06 |
| 1021 | VALLE DEL CAUCA | GUACARÍ | Andean | 113 | 0 | 0.57 | 0 | 0.41 | 0.42 | 0.46 | 0.45 | 0.01 |
| 1022 | VALLE DEL CAUCA | ALCALÁ | Pacific | 111.67 | 0 | 0.49 | 0 | 0.46 | 0.38 | 0.38 | 0.52 | -0.13 |
| 1023 | VALLE DEL CAUCA | OBANDO | Pacific | 112 | 0 | 0.53 | 0 | 0.41 | 0.36 | 0.39 | 0.46 | -0.07 |
| 1024 | VALLE DEL CAUCA | PALMIRA | Pacific | 116 | 0 | 0.67 | 0 | 0.63 | 0.43 | 0.53 | 0.75 | -0.22 |
| 1025 | VALLE DEL CAUCA | PRADERA | Pacific | 113 | 0.5 | 0.6 | 0.07 | 0.33 | 0.38 | 0.51 | 0.34 | 0.17 |
| 1026 | VALLE DEL CAUCA | RESTREPO | Pacific | 159 | 0.17 | 0.67 | 0.03 | 0.24 | 0.37 | 0.5 | 0.22 | 0.28 |
| 1027 | VALLE DEL CAUCA | RIOFRÍO | Pacific | 123.17 | 0.17 | 0.68 | 0.03 | 0.41 | 0.36 | 0.5 | 0.45 | 0.06 |
| 1028 | VALLE DEL CAUCA | ROLDANILLO | Pacific | 120.17 | 0.17 | 0.6 | 0.1 | 0.42 | 0.53 | 0.67 | 0.47 | 0.2 |
| 1029 | VALLE DEL CAUCA | SAN PEDRO | Pacific | 117.5 | 0 | 0.58 | 0 | 0.42 | 0.51 | 0.55 | 0.46 | 0.09 |
| 1030 | VALLE DEL CAUCA | SEVILLA | Pacific | 114.5 | 0.17 | 0.61 | 0.08 | 0.55 | 0.4 | 0.55 | 0.65 | -0.1 |
| 1031 | VALLE DEL CAUCA | TORO | Andean | 123 | 0 | 0.59 | 0 | 0.35 | 0.39 | 0.45 | 0.37 | 0.08 |
| 1032 | VALLE DEL CAUCA | TRUJILLO | Pacific | 120.67 | 0.5 | 0.58 | 0.08 | 0.41 | 0.36 | 0.49 | 0.46 | 0.03 |
| 1033 | VALLE DEL CAUCA | TULUÁ | Andean | 115.17 | 0.33 | 0.63 | 0.1 | 0.52 | 0.37 | 0.55 | 0.61 | -0.06 |
| 1034 | VALLE DEL CAUCA | ULLOA | Pacific | 111.67 | 0 | 0.49 | 0 | 0.37 | 0.4 | 0.39 | 0.39 | 0 |
| 1035 | VALLE DEL CAUCA | VERSALLES | Pacific | 127.17 | 0.33 | 0.6 | 0.05 | 0.27 | 0.39 | 0.51 | 0.27 | 0.24 |
| 1036 | VALLE DEL CAUCA | VIJES | Pacific | 121.83 | 0.33 | 0.64 | 0.14 | 0.23 | 0.53 | 0.73 | 0.21 | 0.52 |
| 1037 | VALLE DEL CAUCA | YOTOCO | Pacific | 123.33 | 0 | 0.71 | 0 | 0.44 | 0.38 | 0.51 | 0.49 | 0.02 |
| 1038 | VALLE DEL CAUCA | YUMBO | Pacific | 121.17 | 0.5 | 0.63 | 0.12 | 0.49 | 0.42 | 0.61 | 0.56 | 0.05 |
| 1039 | VALLE DEL CAUCA | ZARZAL | Pacific | 107 | 0 | 0.55 | 0 | 0.33 | 0.36 | 0.39 | 0.35 | 0.05 |
| 1040 | VALLE DEL CAUCA | LA VICTORIA | Pacific | 157.83 | 0 | 0.62 | 0 | 0.39 | 0.35 | 0.43 | 0.42 | 0.01 |
| 1041 | VALLE DEL CAUCA | LA UNIÓN | Pacific | 126.33 | 0.17 | 0.61 | 0.11 | 0.34 | 0.39 | 0.57 | 0.36 | 0.2 |
| 1042 | VALLE DEL CAUCA | LA CUMBRE | Pacific | 123.17 | 0.17 | 0.62 | 0.09 | 0.36 | 0.38 | 0.55 | 0.39 | 0.16 |
| 1043 | VALLE DEL CAUCA | JAMUNDÍ | Pacific | 121.67 | 0.17 | 0.63 | 0.03 | 0.46 | 0.4 | 0.51 | 0.52 | -0.01 |
| 1044 | VALLE DEL CAUCA | BOLÍVAR | Pacific | 151.67 | 0.67 | 0.71 | 0.1 | 0.38 | 0.51 | 0.72 | 0.41 | 0.31 |
| 1045 | VALLE DEL CAUCA | DAGUA | Pacific | 147.33 | 1.67 | 0.69 | 0.21 | 0.59 | 0.36 | 0.68 | 0.69 | -0.01 |
| 1046 | VALLE DEL CAUCA | BUENAVENTURA | Pacific | 145 | 9.33 | 0.75 | 0.38 | 0.71 | 0.38 | 0.89 | 0.85 | 0.03 |
| 1047 | ARAUCA | ARAUCA | Orinoquia | 132.67 | 0.67 | 0.66 | 0.18 | 0.58 | 0.33 | 0.6 | 0.69 | -0.08 |
| 1048 | ARAUCA | PUERTO RONDÓN | Orinoquia | 128.83 | 0.33 | 0.64 | 0.05 | 0.24 | 0.32 | 0.47 | 0.22 | 0.25 |
| 1049 | ARAUCA | CRAVO NORTE | Orinoquia | 126.83 | 0.5 | 0.57 | 0.11 | 0.49 | 0.38 | 0.53 | 0.56 | -0.04 |
| 1050 | ARAUCA | ARAUQUITA | Orinoquia | 121.33 | 1 | 0.66 | 0.11 | 0.31 | 0.34 | 0.55 | 0.31 | 0.24 |
| 1051 | ARAUCA | FORTUL | Orinoquia | 142.17 | 1 | 0.76 | 0.1 | 0.14 | 0.38 | 0.63 | 0.09 | 0.54 |
| 1052 | ARAUCA | SARAVENA | Orinoquia | 127.5 | 0.5 | 0.68 | 0.07 | 0.36 | 0.38 | 0.56 | 0.39 | 0.17 |
| 1053 | ARAUCA | TAME | Orinoquia | 149 | 3 | 0.8 | 0.22 | 0.53 | 0.37 | 0.77 | 0.61 | 0.16 |
| 1054 | CASANARE | YOPAL | Orinoquia | 172 | 0.5 | 0.74 | 0.12 | 0.73 | 0.4 | 0.65 | 0.89 | -0.23 |
| 1055 | CASANARE | AGUAZUL | Orinoquia | 147.67 | 0.17 | 0.69 | 0.06 | 0.61 | 0.4 | 0.57 | 0.73 | -0.16 |
| 1056 | CASANARE | CHÁMEZA | Orinoquia | 138.67 | 0 | 0.62 | 0 | 0.34 | 0.37 | 0.45 | 0.36 | 0.09 |
| 1057 | CASANARE | LA SALINA | Orinoquia | 108.5 | 0 | 0.54 | 0 | 0.13 | 0.46 | 0.48 | 0.07 | 0.41 |
| 1058 | CASANARE | MANÍ | Orinoquia | 136.17 | 0.5 | 0.53 | 0.12 | 0.59 | 0.33 | 0.48 | 0.69 | -0.21 |
| 1059 | CASANARE | MONTERREY | Orinoquia | 145.83 | 0 | 0.67 | 0 | 0.35 | 0.43 | 0.53 | 0.37 | 0.16 |
| 1060 | CASANARE | NUNCHÍA | Orinoquia | 145.5 | 0 | 0.65 | 0 | 0.34 | 0.39 | 0.49 | 0.36 | 0.13 |
| 1061 | CASANARE | OROCUÉ | Orinoquia | 132.83 | 1.17 | 0.55 | 0.13 | 0.48 | 0.4 | 0.56 | 0.55 | 0.01 |
| 1062 | CASANARE | PORE | Orinoquia | 173.33 | 0 | 0.7 | 0 | 0.26 | 0.45 | 0.57 | 0.25 | 0.32 |
| 1063 | CASANARE | RECETOR | Orinoquia | 138 | 0 | 0.63 | 0 | 0.21 | 0.46 | 0.54 | 0.18 | 0.35 |
| 1064 | CASANARE | SABANALARGA | Orinoquia | 147.83 | 0 | 0.66 | 0 | 0.35 | 0.4 | 0.5 | 0.38 | 0.12 |
| 1065 | CASANARE | SÁCAMA | Orinoquia | 142.5 | 0.17 | 0.65 | 0.03 | 0.2 | 0.36 | 0.49 | 0.17 | 0.31 |
| 1066 | CASANARE | SAN LUIS DE PALENQUE | Orinoquia | 136.67 | 0.33 | 0.53 | 0.05 | 0.49 | 0.51 | 0.57 | 0.56 | 0.01 |
| 1067 | CASANARE | TÁMARA | Orinoquia | 148.17 | 0.17 | 0.63 | 0.03 | 0.31 | 0.32 | 0.44 | 0.32 | 0.12 |
| 1068 | CASANARE | TAURAMENA | Andean | 171.17 | 0 | 0.73 | 0.06 | 0.59 | 0.42 | 0.61 | 0.69 | -0.08 |
| 1069 | CASANARE | TRINIDAD | Orinoquia | 136 | 0 | 0.51 | 0 | 0.5 | 0.37 | 0.39 | 0.57 | -0.18 |
| 1070 | CASANARE | VILLANUEVA | Orinoquia | 178.83 | 0 | 0.72 | 0.06 | 0.63 | 0.4 | 0.59 | 0.76 | -0.17 |
| 1071 | CASANARE | HATO COROZAL | Orinoquia | 153.17 | 0.33 | 0.71 | 0.05 | 0.43 | 0.39 | 0.57 | 0.47 | 0.1 |
| 1072 | CASANARE | PAZ DE ARIPORO | Orinoquia | 170.5 | 0.33 | 0.74 | 0.05 | 0.47 | 0.5 | 0.68 | 0.54 | 0.15 |
| 1073 | PUTUMAYO | MOCOA | Amazon | 164 | 3.33 | 0.77 | 0.24 | 0.47 | 0.45 | 0.83 | 0.54 | 0.3 |
| 1074 | PUTUMAYO | COLÓN | Amazon | 96.33 | 0.67 | 0.53 | 0.08 | 0.23 | 0.45 | 0.54 | 0.21 | 0.34 |
| 1075 | PUTUMAYO | ORITO | Amazon | 172.33 | 3.5 | 0.75 | 0.22 | 0.49 | 0.36 | 0.73 | 0.56 | 0.17 |
| 1076 | PUTUMAYO | SIBUNDOY | Amazon | 91.5 | 0.67 | 0.51 | 0.12 | 0.5 | 0.34 | 0.47 | 0.58 | -0.1 |
| 1077 | PUTUMAYO | SAN FRANCISCO | Amazon | 157.67 | 0.67 | 0.71 | 0.08 | 0.34 | 0.33 | 0.54 | 0.36 | 0.18 |
| 1078 | PUTUMAYO | SAN MIGUEL | Amazon | 169.67 | 0.33 | 0.69 | 0.05 | 0.2 | 0.29 | 0.47 | 0.16 | 0.31 |
| 1079 | PUTUMAYO | SANTIAGO | Amazon | 119.33 | 1 | 0.59 | 0.1 | 0.32 | 0.34 | 0.49 | 0.33 | 0.16 |
| 1080 | PUTUMAYO | VALLE DEL GUAMUEZ | Amazon | 175 | 1.67 | 0.72 | 0.23 | 0.36 | 0.32 | 0.67 | 0.38 | 0.29 |
| 1081 | PUTUMAYO | VILLAGARZÓN | Amazon | 165.83 | 3.17 | 0.77 | 0.14 | 0.42 | 0.46 | 0.74 | 0.47 | 0.28 |
| 1082 | PUTUMAYO | PUERTO CAICEDO | Amazon | 160.83 | 1 | 0.67 | 0.1 | 0.27 | 0.44 | 0.63 | 0.26 | 0.37 |
| 1083 | PUTUMAYO | PUERTO ASÍS | Amazon | 159.83 | 2.17 | 0.66 | 0.17 | 0.54 | 0.32 | 0.58 | 0.63 | -0.04 |
| 1084 | PUTUMAYO | PUERTO GUZMÁN | Amazon | 186 | 1.67 | 0.76 | 0.17 | 0.33 | 0.29 | 0.62 | 0.34 | 0.27 |
| 1085 | PUTUMAYO | PUERTO LEGUÍZAMO | Amazon | 198 | 3 | 0.76 | 0.14 | 0.37 | 0.3 | 0.6 | 0.4 | 0.2 |
| 1086 | ARCHIPIÉLAGO DE SAN ANDRÉS, PROVIDENCIA Y SANTA CATALINA | PROVIDENCIA | Caribbean | 85.83 | 0 | 0.19 | 0 | 0.58 | 0.43 | 0.25 | 0.69 | -0.44 |
| 1087 | ARCHIPIÉLAGO DE SAN ANDRÉS, PROVIDENCIA Y SANTA CATALINA | SAN ANDRÉS | Caribbean | 126.67 | 0.17 | 0.6 | 0 | 0.63 | 0.6 | 0.64 | 0.74 | -0.1 |
| 1088 | AMAZONAS | LETICIA | Amazon | 252.17 | 2.83 | 0.77 | 0.22 | 0.61 | 0.42 | 0.79 | 0.72 | 0.08 |
| 1089 | AMAZONAS | EL ENCANTO | Amazon | 199.83 | 0.17 | 0.74 | 0.03 | 0.27 | 0.27 | 0.45 | 0.27 | 0.19 |
| 1090 | AMAZONAS | LA CHORRERA | Amazon | 185.33 | 0.33 | 0.7 | 0.05 | 0.22 | 0.24 | 0.43 | 0.19 | 0.24 |
| 1091 | AMAZONAS | LA PEDRERA | Amazon | 202.17 | 1.33 | 0.73 | 0.1 | 0.42 | 0.35 | 0.59 | 0.47 | 0.12 |
| 1092 | AMAZONAS | LA VICTORIA | Amazon | 158 | 0.5 | 0.63 | 0.07 | 0.25 | 0.29 | 0.45 | 0.24 | 0.22 |
| 1093 | AMAZONAS | MIRITÍ - PARANÁ | Amazon | 195.17 | 1 | 0.69 | 0.22 | 0.09 | 0.25 | 0.59 | 0.02 | 0.57 |
| 1094 | AMAZONAS | PUERTO ALEGRÍA | Amazon | 194.5 | 0.17 | 0.71 | 0.03 | 0.09 | 0.23 | 0.4 | 0.02 | 0.38 |
| 1095 | AMAZONAS | PUERTO ARICA | Amazon | 199.67 | 0.33 | 0.69 | 0.05 | 0.11 | 0.25 | 0.43 | 0.04 | 0.39 |
| 1096 | AMAZONAS | PUERTO NARIÑO | Amazon | 250.33 | 0.5 | 0.75 | 0.11 | 0.32 | 0.48 | 0.73 | 0.34 | 0.39 |
| 1097 | AMAZONAS | PUERTO SANTANDER | Amazon | 195.33 | 0.67 | 0.71 | 0.08 | 0.2 | 0.27 | 0.49 | 0.17 | 0.31 |
| 1098 | AMAZONAS | TARAPACÁ | Amazon | 207.67 | 0.33 | 0.7 | 0.05 | 0.32 | 0.34 | 0.52 | 0.33 | 0.19 |
| 1099 | GUAINÍA | INÍRIDA | Orinoquia | 174.33 | 2.83 | 0.72 | 0.2 | 0.65 | 0.35 | 0.68 | 0.78 | -0.1 |
| 1100 | GUAINÍA | BARRANCOMINAS | Orinoquia | 165 | 2.5 | 0.72 | 0.13 | 0.42 | 0.29 | 0.55 | 0.47 | 0.09 |
| 1101 | GUAINÍA | SAN FELIPE | Orinoquia | 174.33 | 0.17 | 0.68 | 0.03 | 0.32 | 0.3 | 0.45 | 0.33 | 0.11 |
| 1102 | GUAINÍA | PUERTO COLOMBIA | Orinoquia | 151.33 | 1 | 0.68 | 0.1 | 0.08 | 0.35 | 0.56 | 0 | 0.55 |
| 1103 | GUAINÍA | LA GUADALUPE | Orinoquia | 175.17 | 0.17 | 0.64 | 0.03 | 0.08 | 0.25 | 0.38 | 0 | 0.38 |
| 1104 | GUAINÍA | CACAHUAL | Orinoquia | 140.83 | 0.17 | 0.61 | 0.03 | 0.23 | 0.28 | 0.39 | 0.2 | 0.18 |
| 1105 | GUAINÍA | PANA PANA | Orinoquia | 140.17 | 0.67 | 0.63 | 0.08 | 0.19 | 0.3 | 0.46 | 0.16 | 0.3 |
| 1106 | GUAINÍA | MORICHAL | Orinoquia | 167.5 | 0.67 | 0.72 | 0.08 | 0.09 | 0.22 | 0.45 | 0.03 | 0.42 |
| 1107 | GUAVIARE | SAN JOSÉ DEL GUAVIARE | Orinoquia | 176.5 | 2.5 | 0.77 | 0.31 | 0.64 | 0.42 | 0.87 | 0.76 | 0.11 |
| 1108 | GUAVIARE | CALAMAR | Orinoquia | 171.33 | 0.83 | 0.76 | 0.21 | 0.3 | 0.28 | 0.65 | 0.3 | 0.35 |
| 1109 | GUAVIARE | EL RETORNO | Orinoquia | 144.33 | 1 | 0.67 | 0.22 | 0.29 | 0.35 | 0.67 | 0.29 | 0.38 |
| 1110 | GUAVIARE | MIRAFLORES | Orinoquia | 153.33 | 2.33 | 0.73 | 0.26 | 0.33 | 0.29 | 0.68 | 0.35 | 0.34 |
| 1111 | VAUPÉS | MITÚ | Amazon | 173.67 | 0.5 | 0.75 | 0.05 | 0.38 | 0.44 | 0.64 | 0.41 | 0.23 |
| 1112 | VAUPÉS | CARURÚ | Amazon | 147 | 0.5 | 0.69 | 0.07 | 0.24 | 0.36 | 0.55 | 0.23 | 0.32 |
| 1113 | VAUPÉS | PACOA | Amazon | 154 | 0.5 | 0.69 | 0.19 | 0.22 | 0.36 | 0.65 | 0.2 | 0.45 |
| 1114 | VAUPÉS | TARAIRA | Amazon | 168.67 | 0.33 | 0.69 | 0.05 | 0.41 | 0.38 | 0.55 | 0.46 | 0.09 |
| 1115 | VAUPÉS | PAPUNAHUA | Amazon | 150.33 | 0.5 | 0.66 | 0.07 | 0.22 | 0.35 | 0.52 | 0.19 | 0.33 |
| 1116 | VAUPÉS | YAVARATÉ | Amazon | 168.67 | 0.17 | 0.66 | 0.03 | 0.24 | 0.45 | 0.57 | 0.22 | 0.35 |
| 1117 | VICHADA | SANTA ROSALÍA | Orinoquia | 137 | 1.17 | 0.57 | 0.1 | 0.45 | 0.48 | 0.61 | 0.5 | 0.11 |
| 1118 | VICHADA | PUERTO CARREÑO | Orinoquia | 171.17 | 1.5 | 0.68 | 0.16 | 0.67 | 0.37 | 0.63 | 0.81 | -0.17 |
| 1119 | VICHADA | LA PRIMAVERA | Orinoquia | 137.33 | 0.67 | 0.59 | 0.08 | 0.43 | 0.36 | 0.49 | 0.48 | 0.01 |
| 1120 | VICHADA | CUMARIBO | Orinoquia | 187.67 | 5.5 | 0.76 | 0.16 | 0.46 | 0.36 | 0.67 | 0.52 | 0.15 |

### 3. Supporting information on data

#### 3.1. Predictor and response variables descriptive statistics and distributions

**Table S2. A list of variables before transformation and their summary statistics**

| **Metric** | **Predictor variable** | **Meaning** | **Min.** | **1st Quantile** | **Median** | **Mean** | **3rd Quantile** | **Max.** |
| --- | --- | --- | --- | --- | --- | --- | --- | --- |
| Biodiversity richness index | Bird species richness | Maximum number of bird species found in a municipality | 140 | 342 | 408 | 411 | 491 | 648 |
| Biodiversity richness index | Reptile species richness | Maximum number of reptile species found in a municipality | 53 | 84 | 94 | 96 | 108 | 169 |
| Biodiversity richness index | Amphibian species richness | Maximum number of fish species found in a municipality | 5 | 15 | 23 | 27 | 33 | 131 |
| Biodiversity richness index | Fish species richness | Maximum number of fish species found in a municipality | 0 | 16 | 40 | 54 | 68 | 538 |
| Biodiversity richness index | Mammal species richness | Maximum number of mammal species found in a municipality | 41 | 108 | 117 | 114 | 125 | 147 |
| Biodiversity richness index | Ecosystems richness | Maximum number of ecosystems found in a municipality | 1 | 7 | 8 | 8 | 10 | 19 |
| Institutionalized cultural richness index | World Heritage Sites | # of UNESCO world heritage sites in a municipality | 0 | 0 | 0 | 0 | 0 | 2 |
| Institutionalized cultural richness index | Intangible Cultural Heritage Sites | # of UNESCO intangible cultural heritage sites in a municipality | 0 | 0 | 0 | 0 | 0 | 3 |
| Institutionalized cultural richness index | Music festivals of endemic musical rhythms | # music festivals from endemic musical rhythms in a municipality | 0 | 0 | 0 | 0 | 0 | 3 |
| Institutionalized cultural richness index | Afro-Colombian Lands | # of Afro-Colombian territories in a municipality | 1 | 1 | 1 | 1 | 1 | 42 |
| Institutionalized cultural richness index | Indigenous Lands | # of Indigenous reserves in a municipality | 1 | 1 | 1 | 2 | 1 | 38 |
| Institutionalized cultural richness index | Museums density | # of museums per km^2^ in each municipality | 0 | 0 | 0 | 0 | 0 | 0 |
| Accessibility index | Bird lodging density | # of bird lodges per km^2^ in each municipality | 0 | 0 | 0 | 0 | 0 | 0 |
| Accessibility index | Road density | built roads per km^2^ in each municipality | 0 | 103 | 231 | 457 | 511 | 10679 |
| Accessibility index | Distance to nearest airport | Distance to nearest airport in each municipality | 0 | 12 | 20 | 22 | 30 | 105 |
| Accessibility index | Lodging density | # of hotels and lodges per km^2^ in each municipality | 0 | 0 | 0 | 0 | 0 | 2 |
| Accessibility index | Conflict density | # of armed conflict cases per municipality divided by the area | 0 | 0 | 0 | 1 | 1 | 62 |
| Demand index | eBird hotspots | # eBird hotspots per km^2^ in each municipality | 0 | 0 | 0 | 2 | 2 | 107 |
| Demand index | Sport fisheries | # sport fisheries per km^2^ in each municipality | 0 | 0 | 0 | 0 | 0 | 5 |
| Demand index | Flickr PUD | # of Flicker Photo-User-Days | 0 | 1 | 2 | 14 | 5 | 4240 |
| Demand index | Airport visitors | # of passengers arrived at each airport | 0 | 0 | 0 | 19796 | 0 | 10697700 |
| Demand index | Music festivals visitors | # of visitors at music festivals | 0 | 0 | 0 | 10230 | 0 | 4975448 |

#### 3.2. Pairwise correlations between variables used for each index

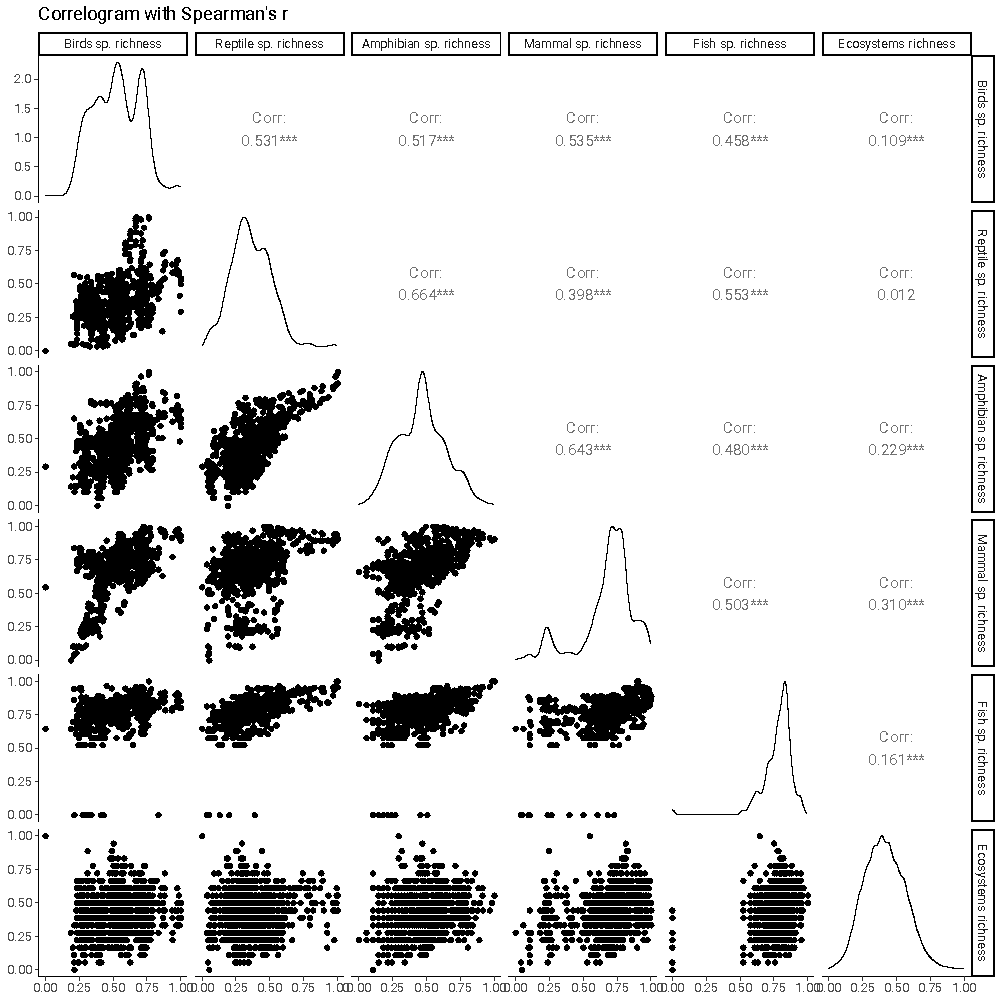

**Figure S3. Correlogram between biodiversity richness predictor variables.**

Many predictor variables used for the biodiversity index are correlated with one another, as shown by this correlogram (Spearman’s r>0.5). Therefore, we dropped the species richness of reptiles, amphibians and mammals, and only did the index with fish, birds, and ecosystems richness.

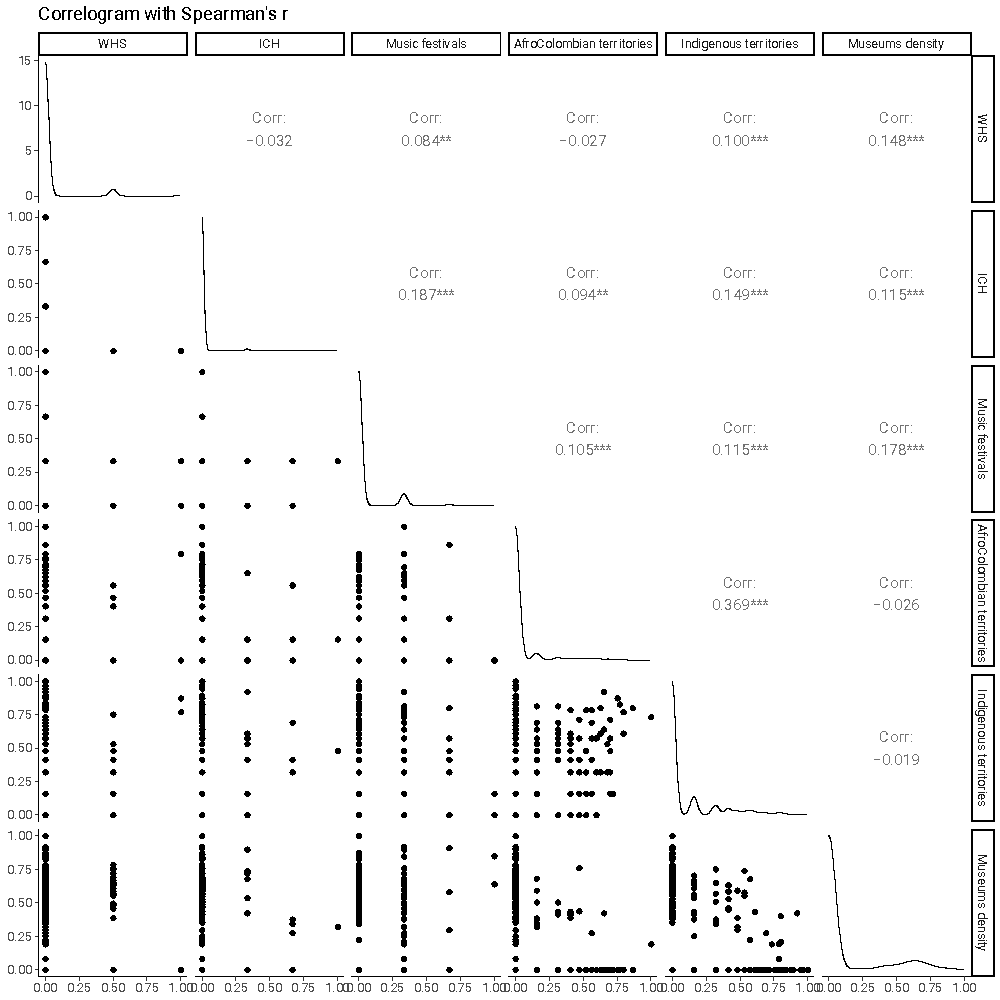

**Figure S4. Correlogram between institutionalized cultural richness predictor variables.**

None of the predictor variables used for the cultural pull factors index are highly correlated with one another, as shown by this correlogram (Spearman’s r<0.5). All of these were included in the composite index.

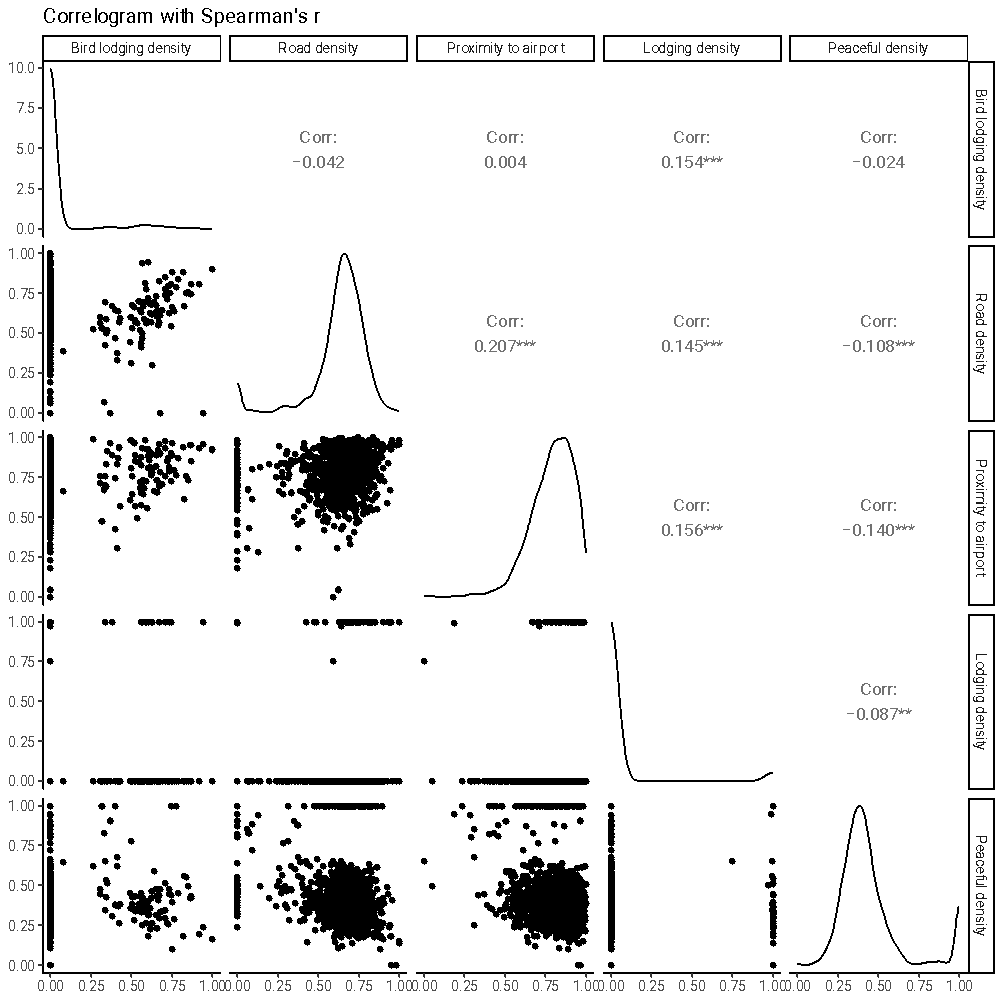

**Figure S5. Correlogram between accessibility index predictor variables.**

None of the predictor variables used for the accessibility index are highly correlated with one another, as shown by this correlogram (Spearman’s r<0.5).

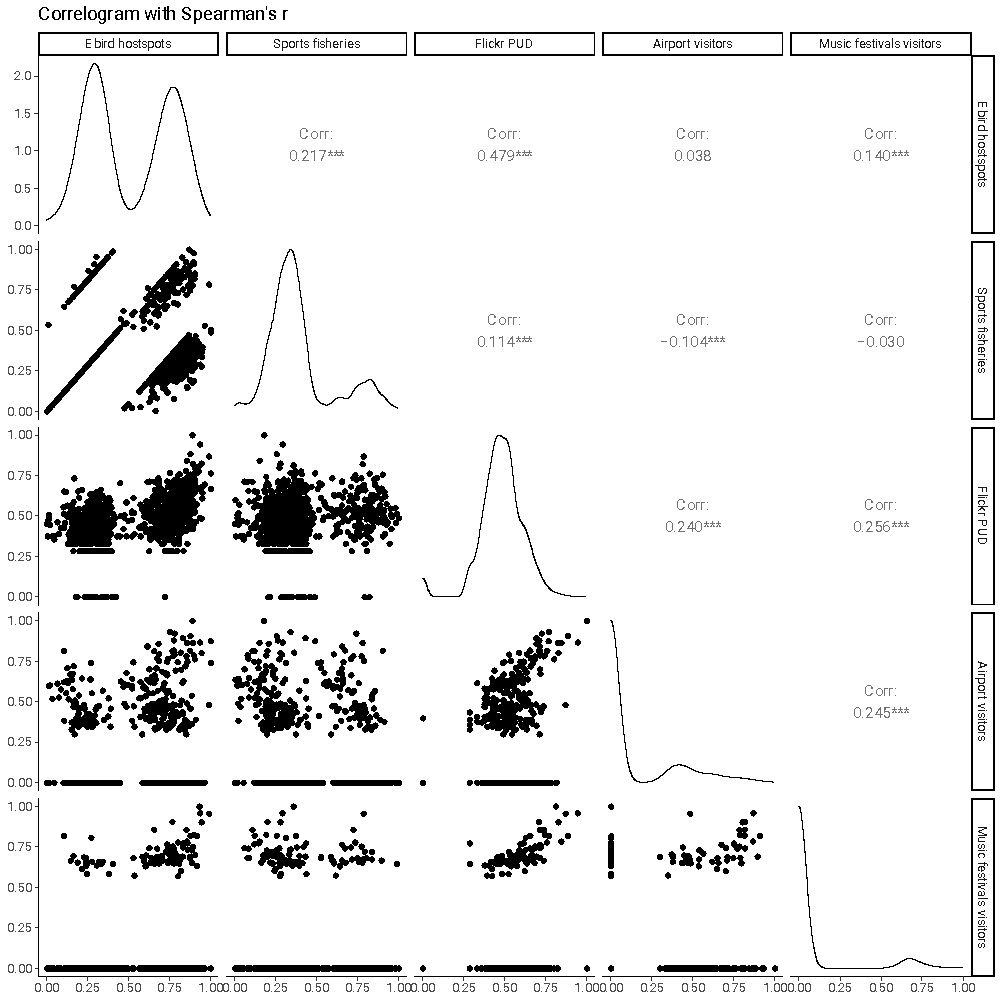

**Figure S6. Correlogram between tourism demand predictor variables.**

None of the predictor variables used for the demand index are highly correlated with one another, as shown by this correlogram (Spearman’s r<0.5).

#### 3.3. Distribution of individual variables

##### 3.3.1. Biodiversity richness variables

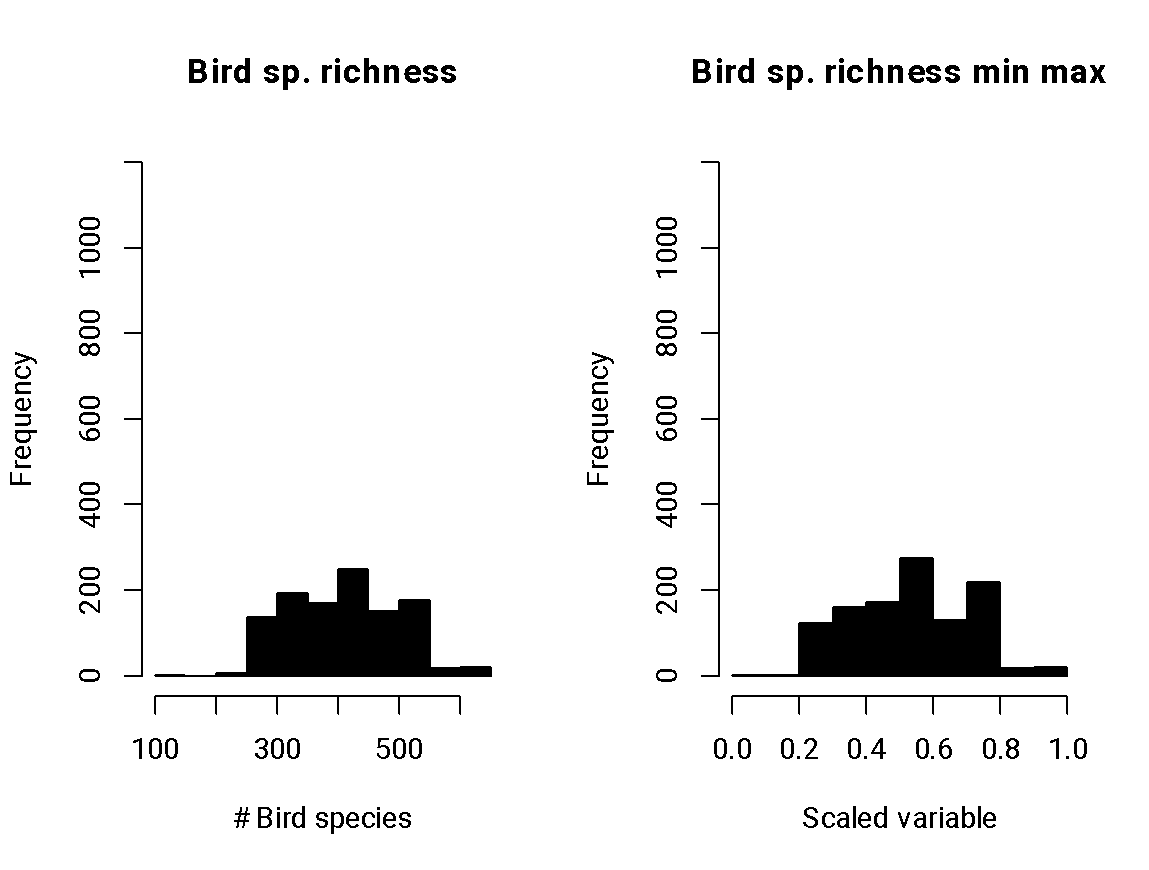

***Figure S7. Histograms of bird species richness.***

Left panel is the untransformed variable distribution, and the right panel is the transformed variable distribution.

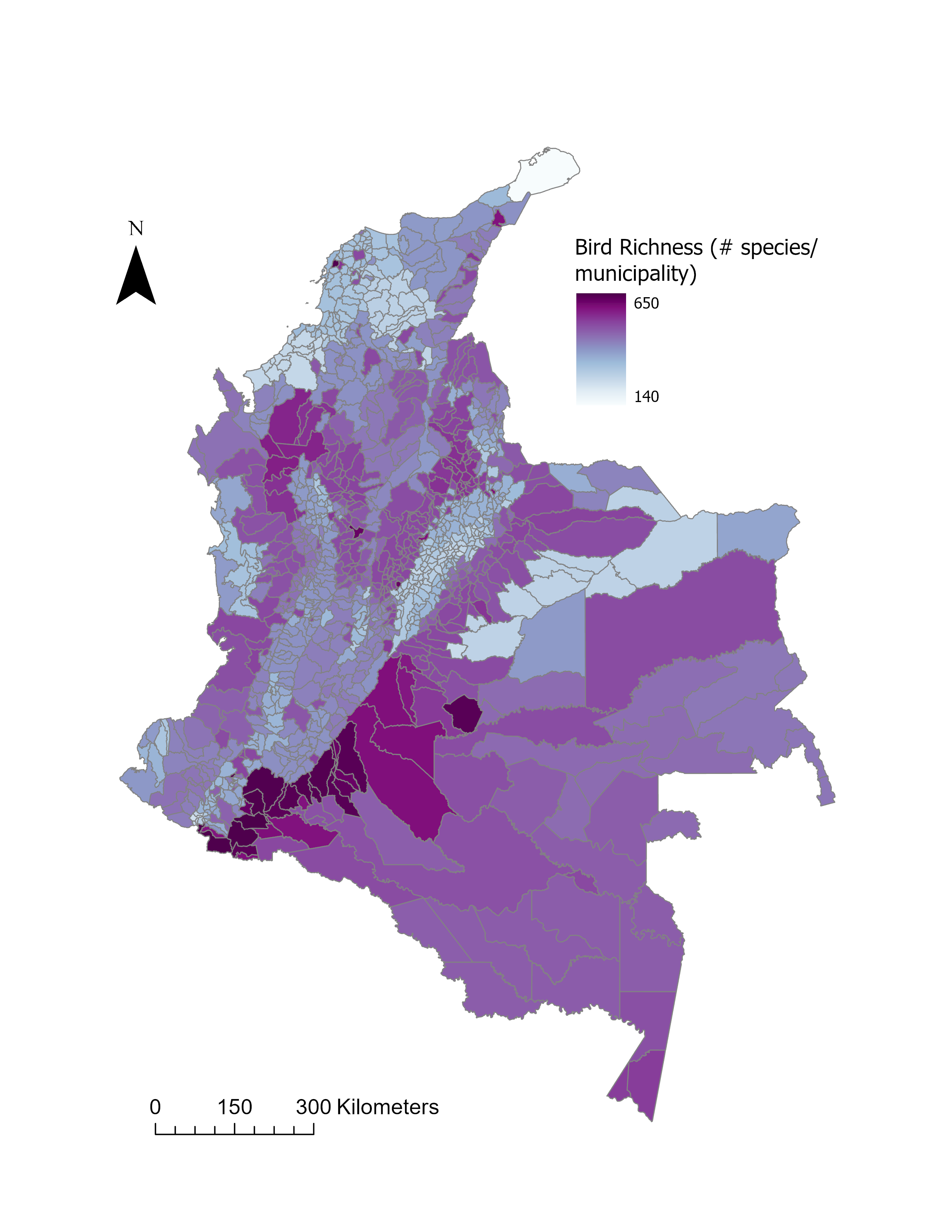

**Figure S8. Spatial distribution of the bird richness layer**

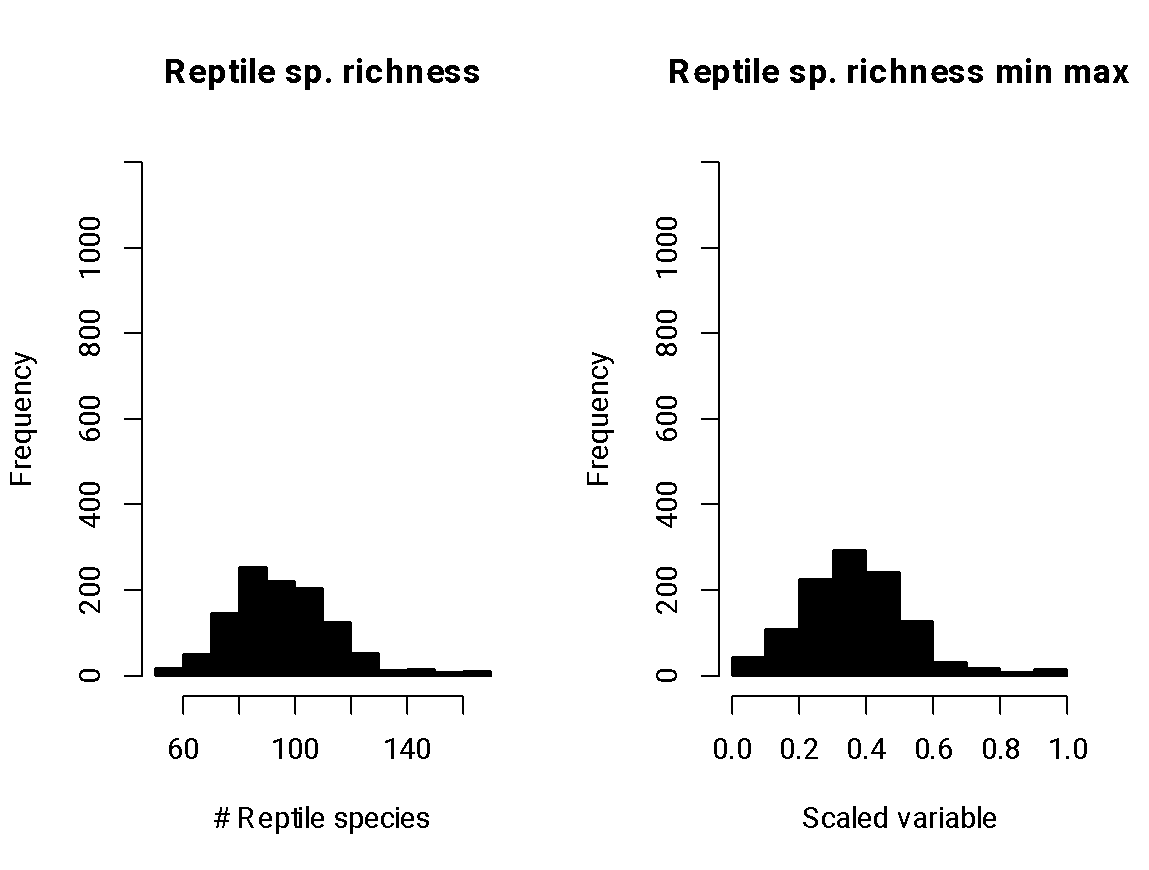

***Figure S9. Histograms of reptile species richness.***

Left panel is the untransformed variable distribution, and the right panel is the transformed variable distribution.

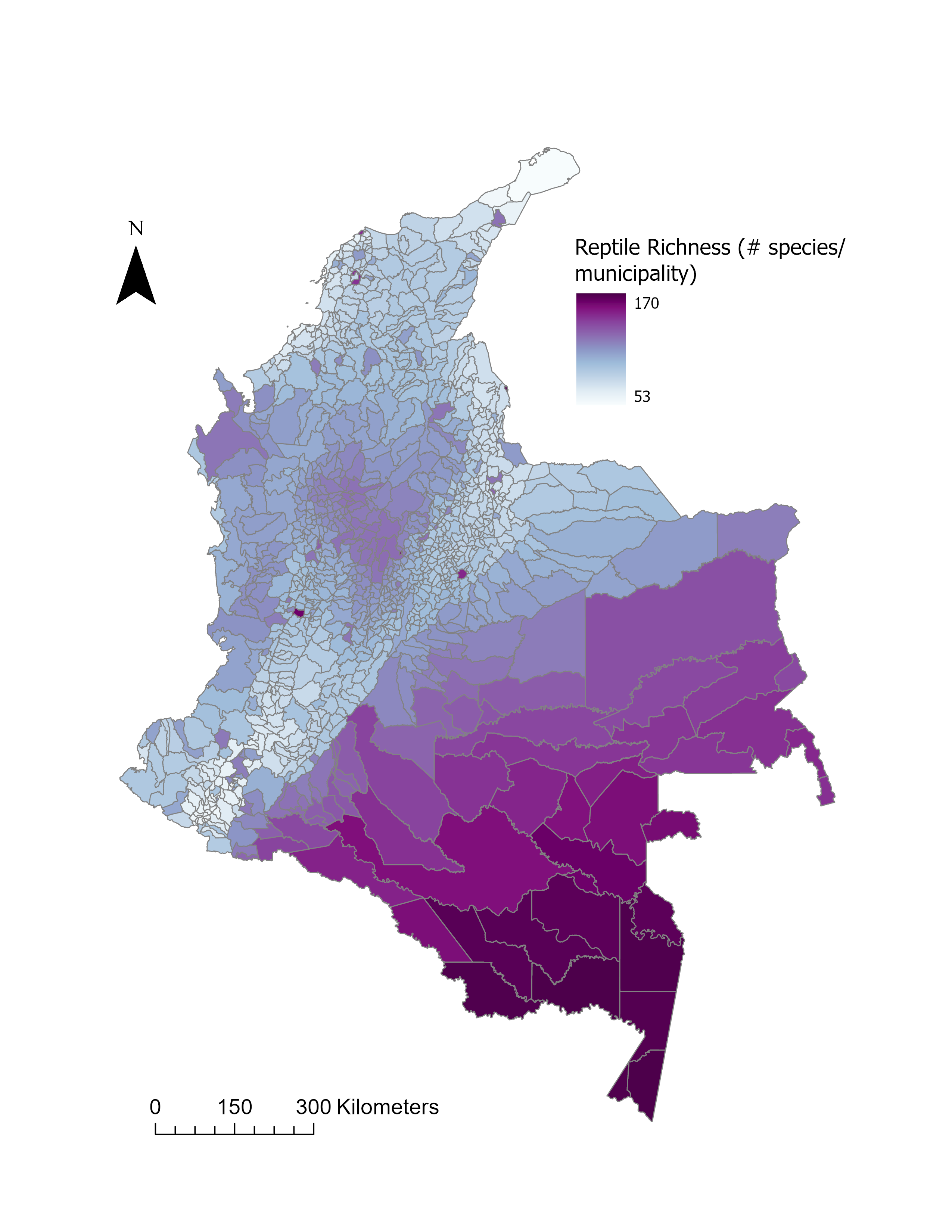

**Figure S10.** **Spatial distribution of the reptile richness layer.**

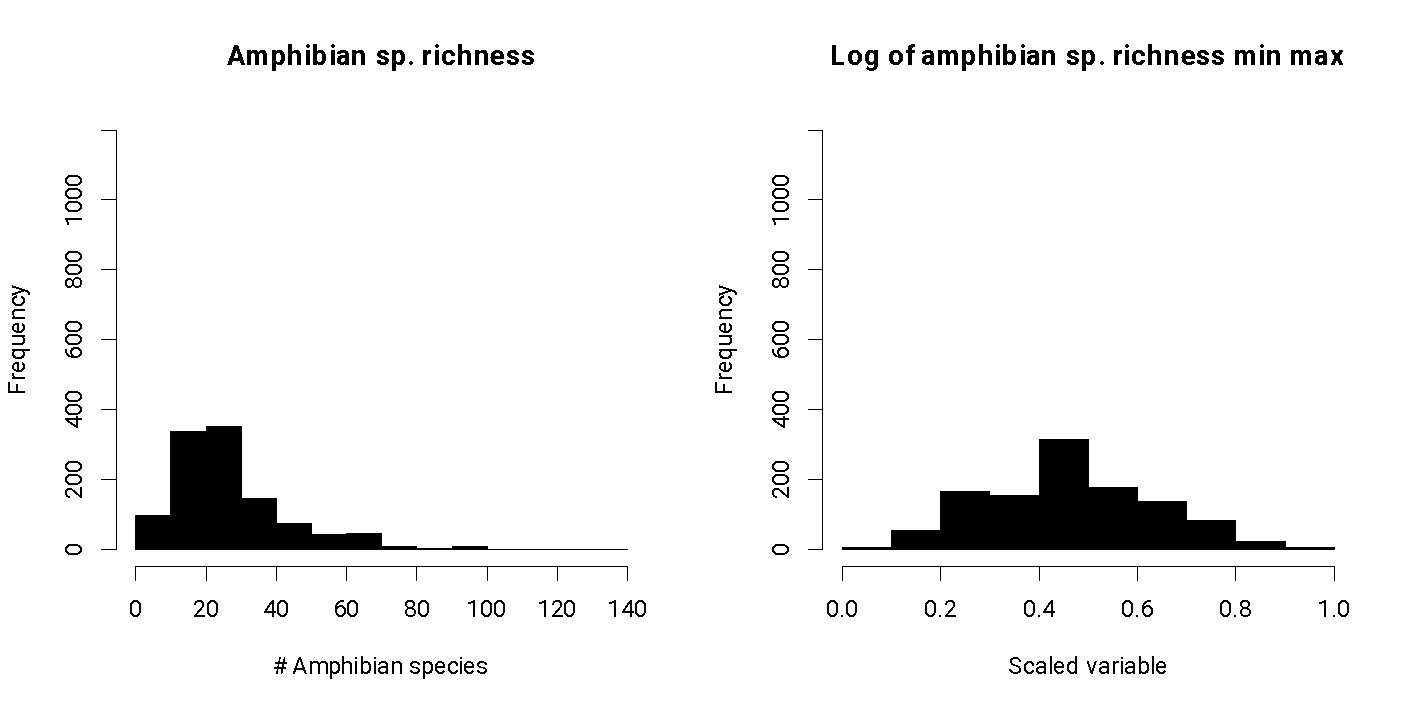

***Figure S11. Histograms of amphibian species richness.***

Left panel is the untransformed variable distribution, and the right panel is the transformed variable distribution.

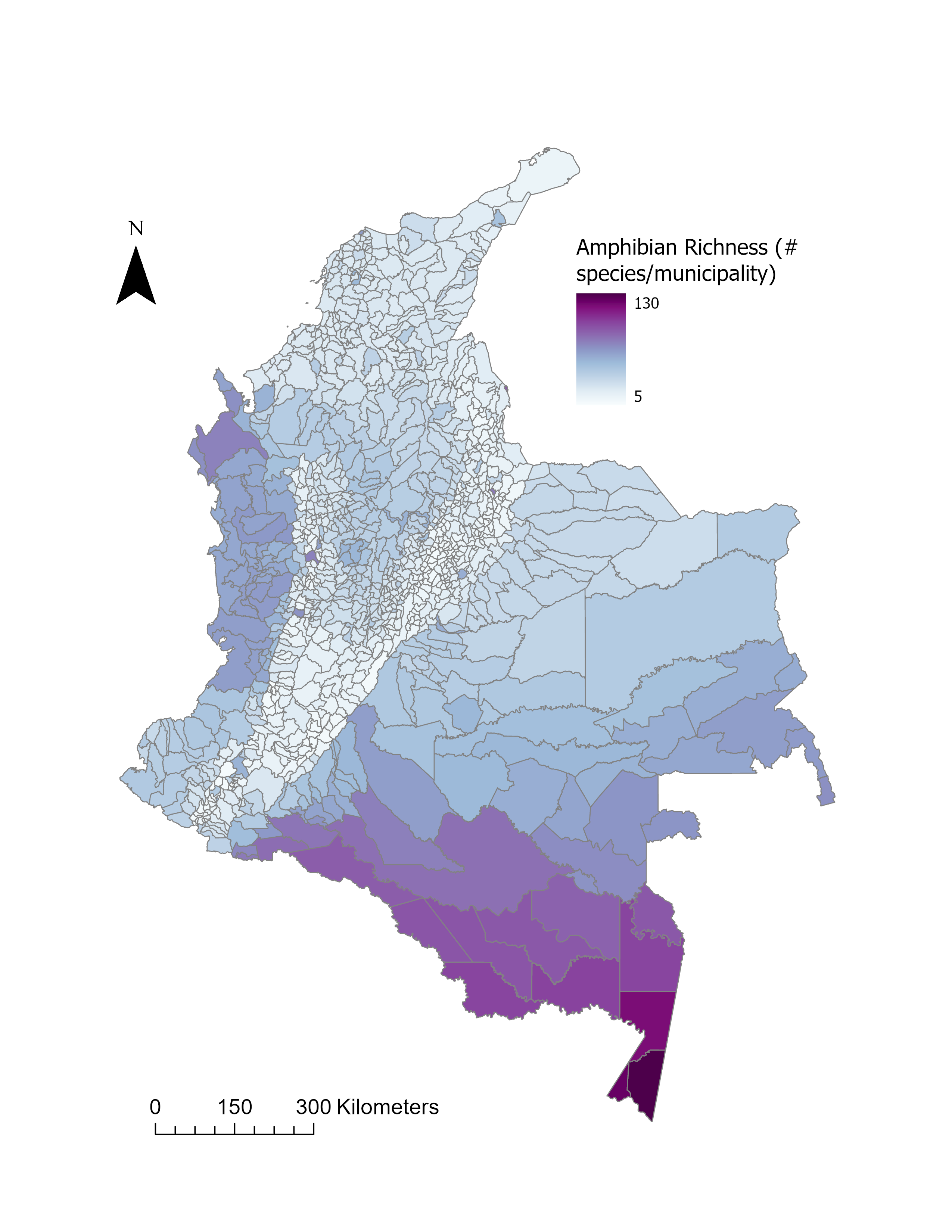

**Figure S12. Spatial distribution of the amphibian richness layer.**

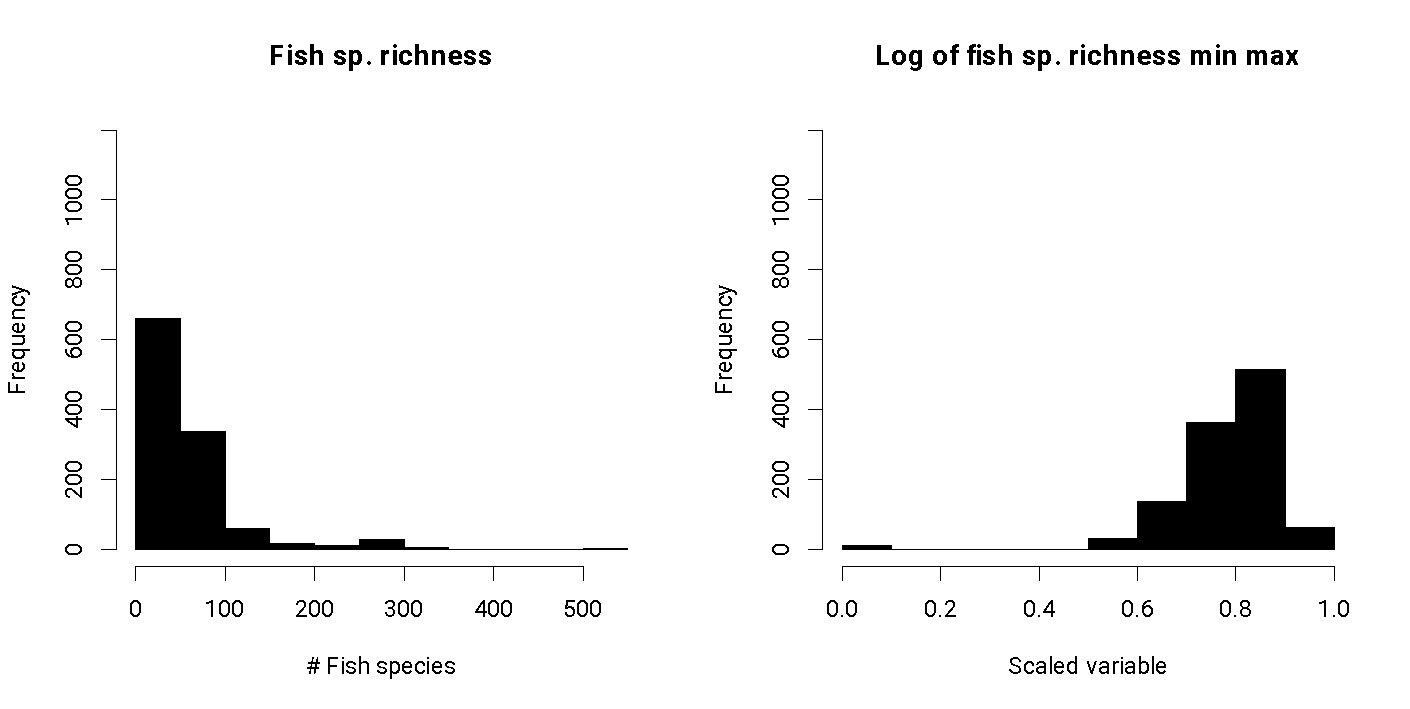

***Figure S13. Histograms of fish species richness.***

Left panel is the untransformed variable distribution, and the right panel is the transformed variable distribution.

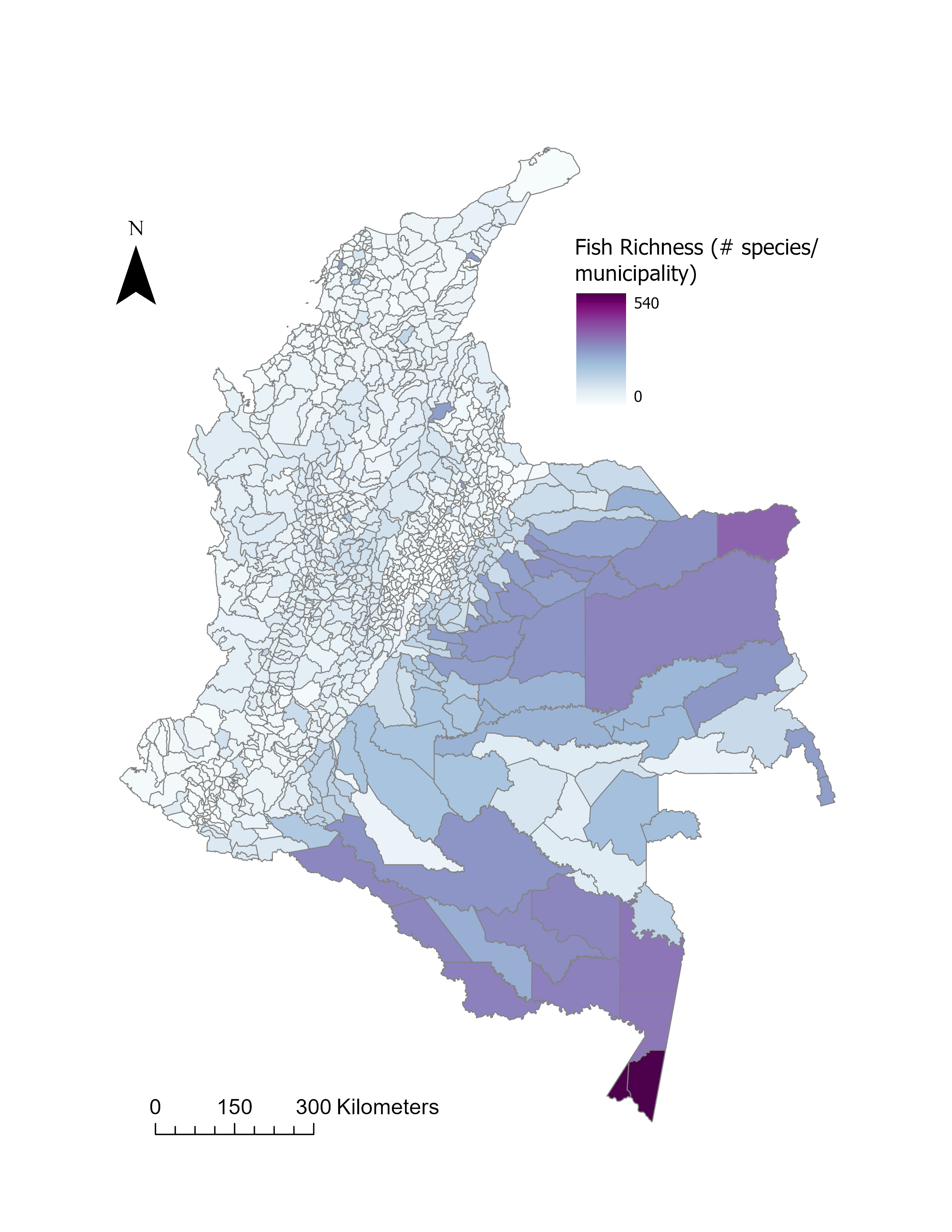

**Figure S14. Spatial distribution of the fish richness layer.**

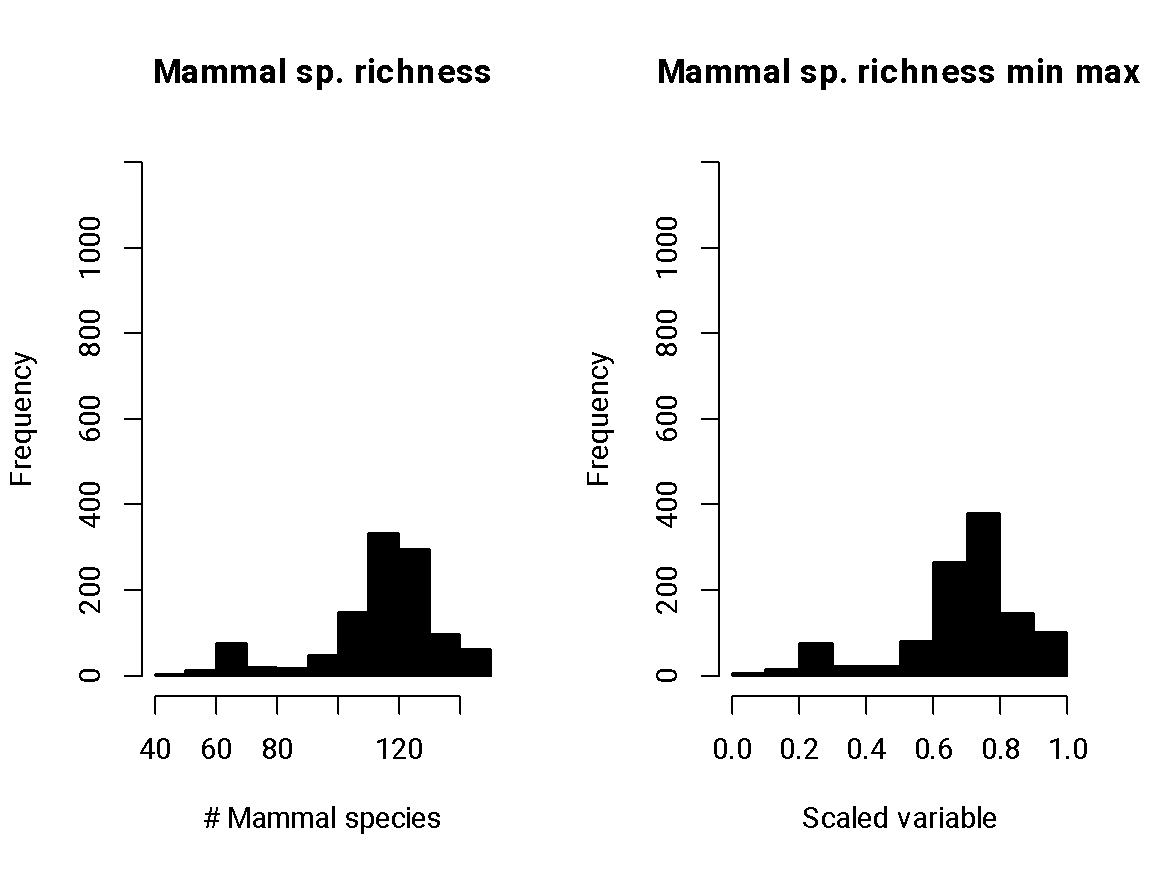

***Figure S15. Histograms of mammal species richness.***

Left panel is the untransformed variable distribution, and the right panel is the transformed variable distribution.

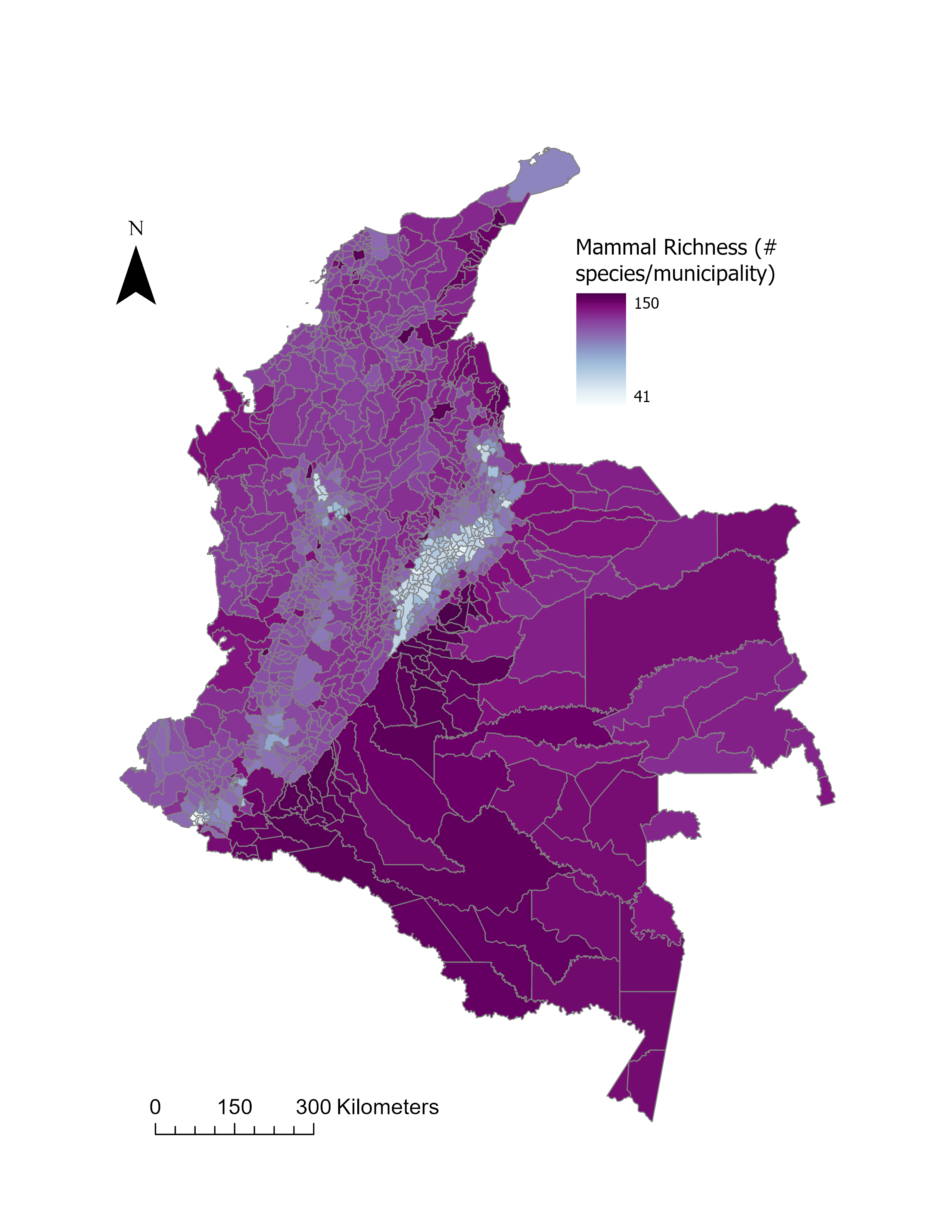

**Figure S16. Spatial distribution of the mammal richness layer.**

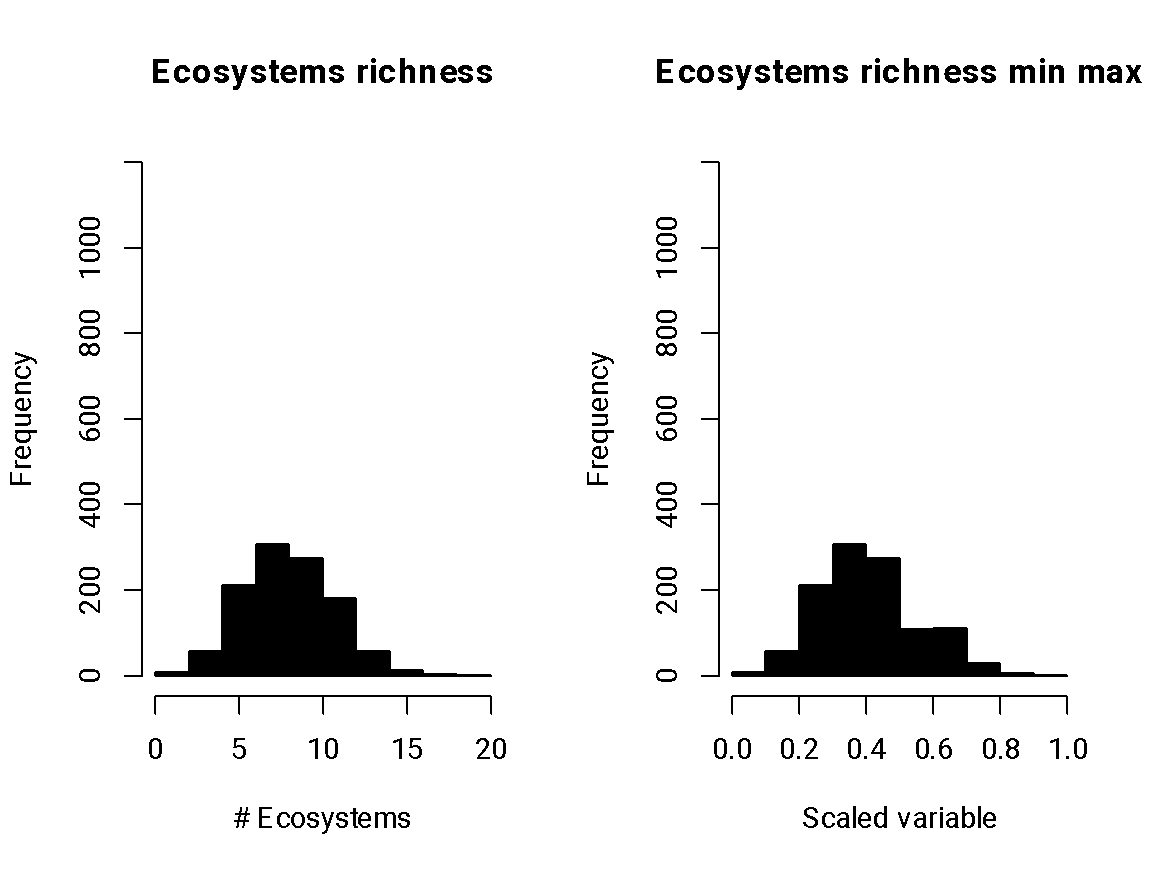

***Figure S17. Histograms of ecosystems richness*.**

Left panel is the untransformed variable distribution, and the right panel is the transformed variable distribution.

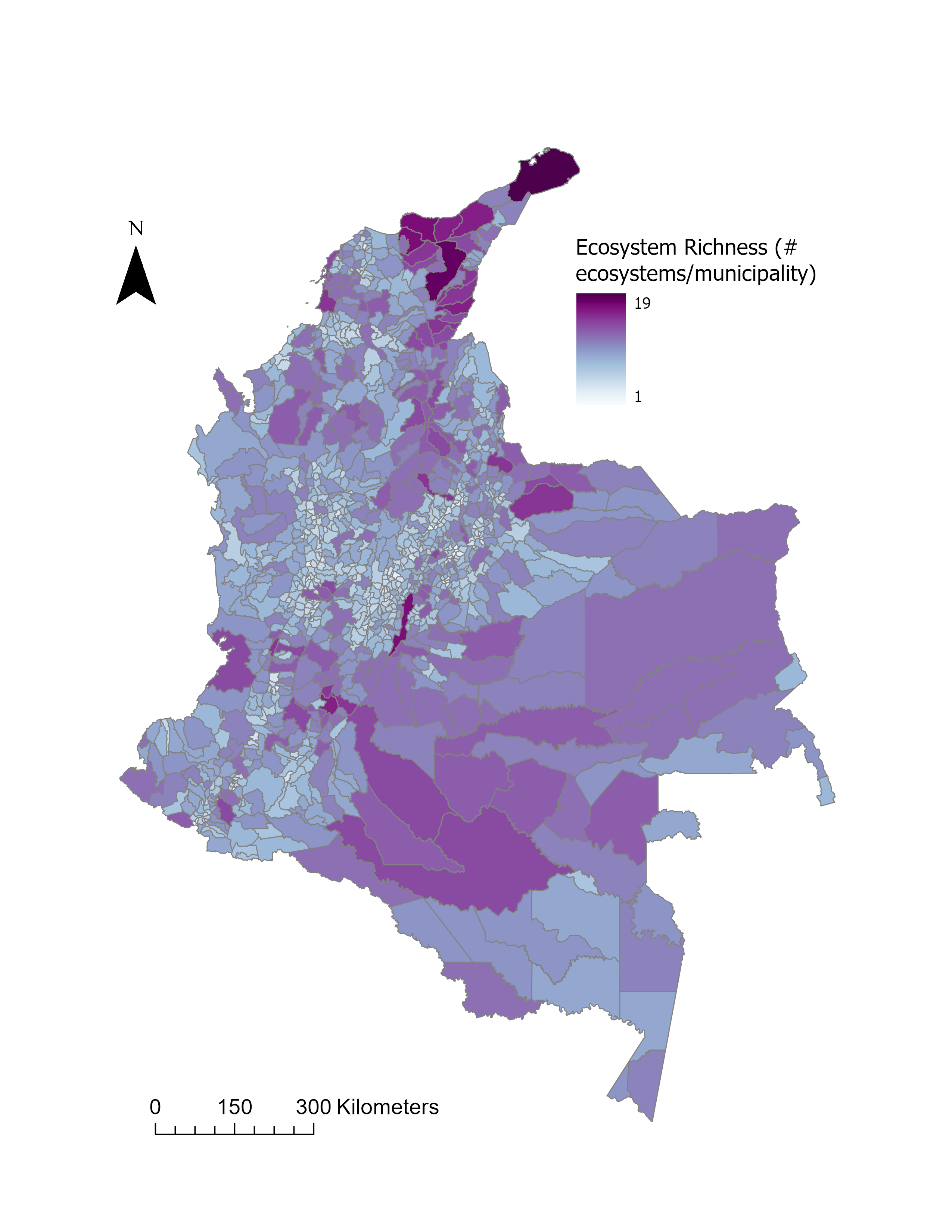

**Figure S18. Spatial distribution of the ecosystem richness layer.**

##### 3.3.2. Institutionalized cultural richness variables

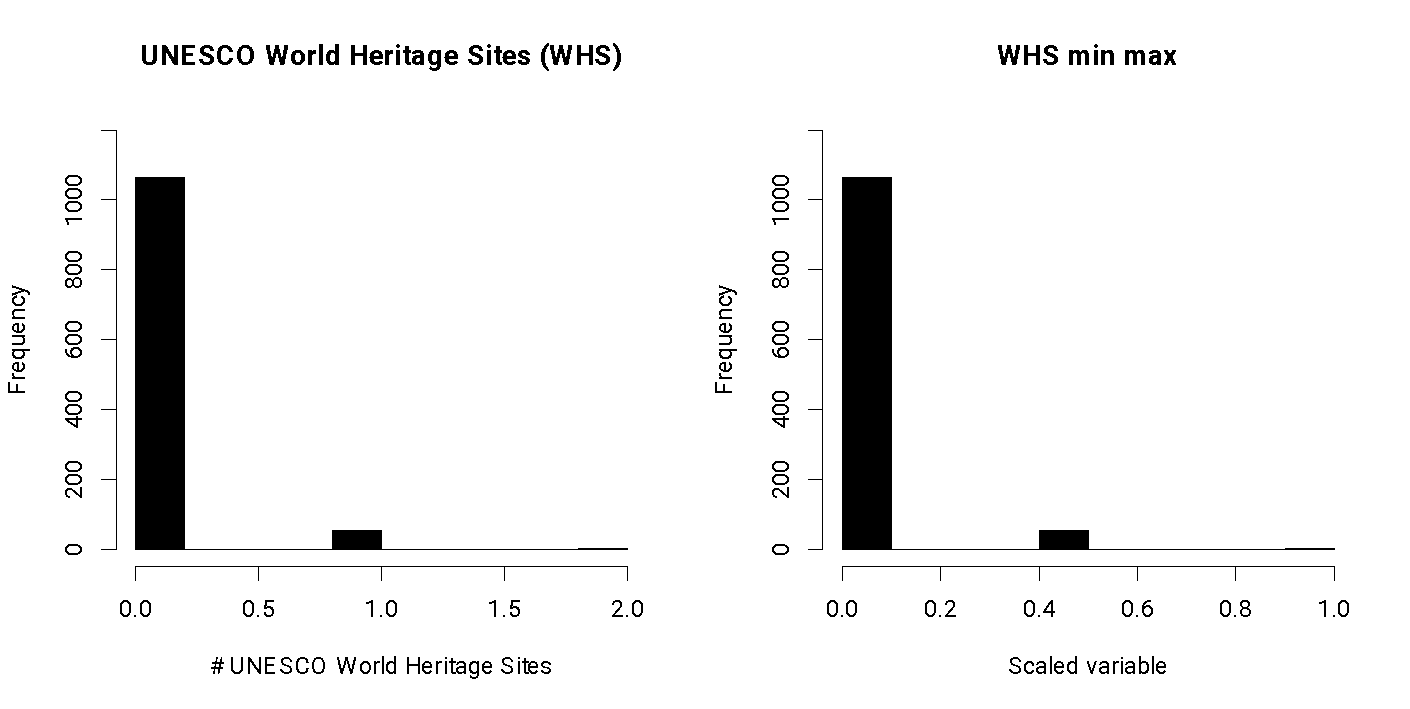

**Figure S19.** **Histograms of UNESCO World Heritage Sites.**

Left panel is the untransformed variable distribution, and the right panel is the transformed variable distribution.

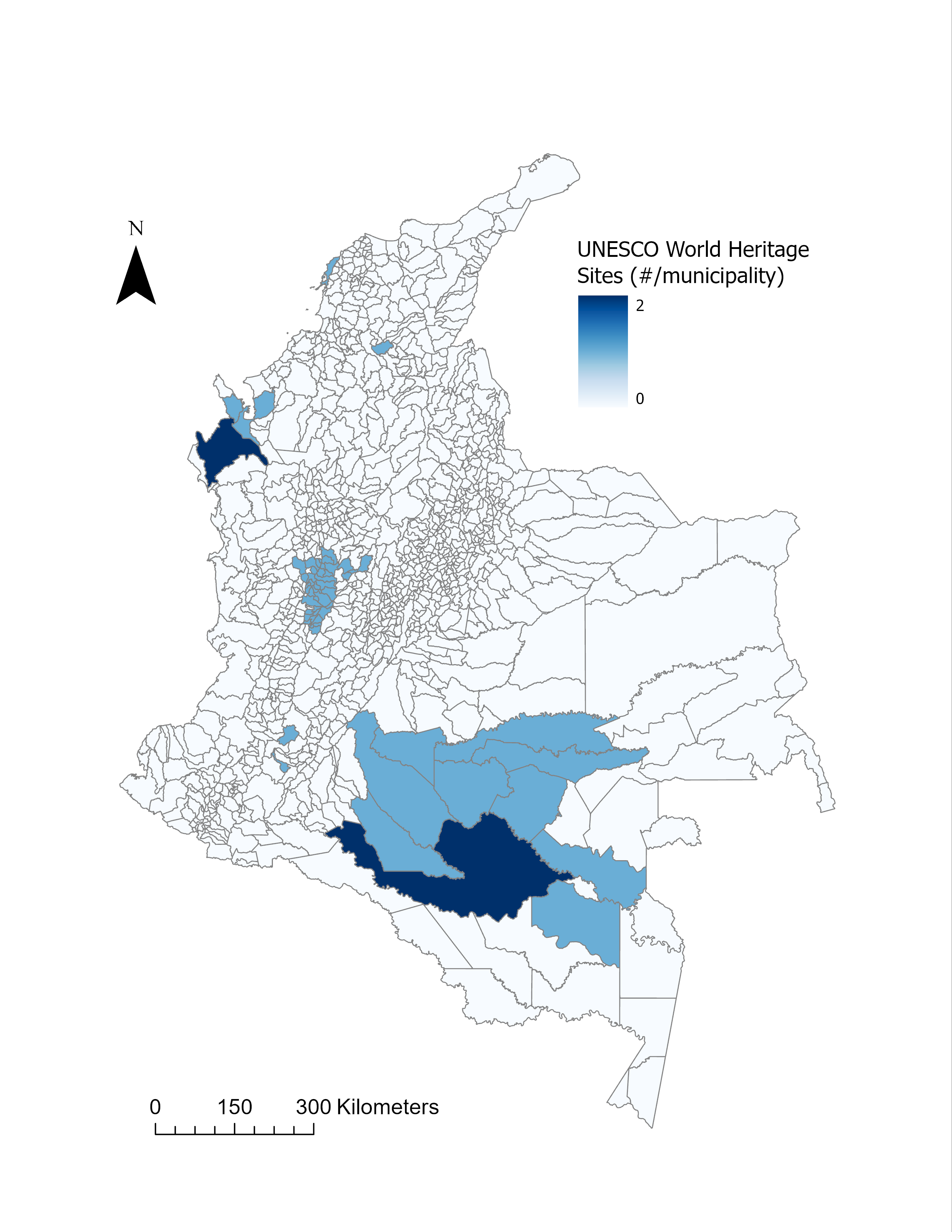

**Figure S20. Spatial distribution of UNESCO World Heritage Sites**

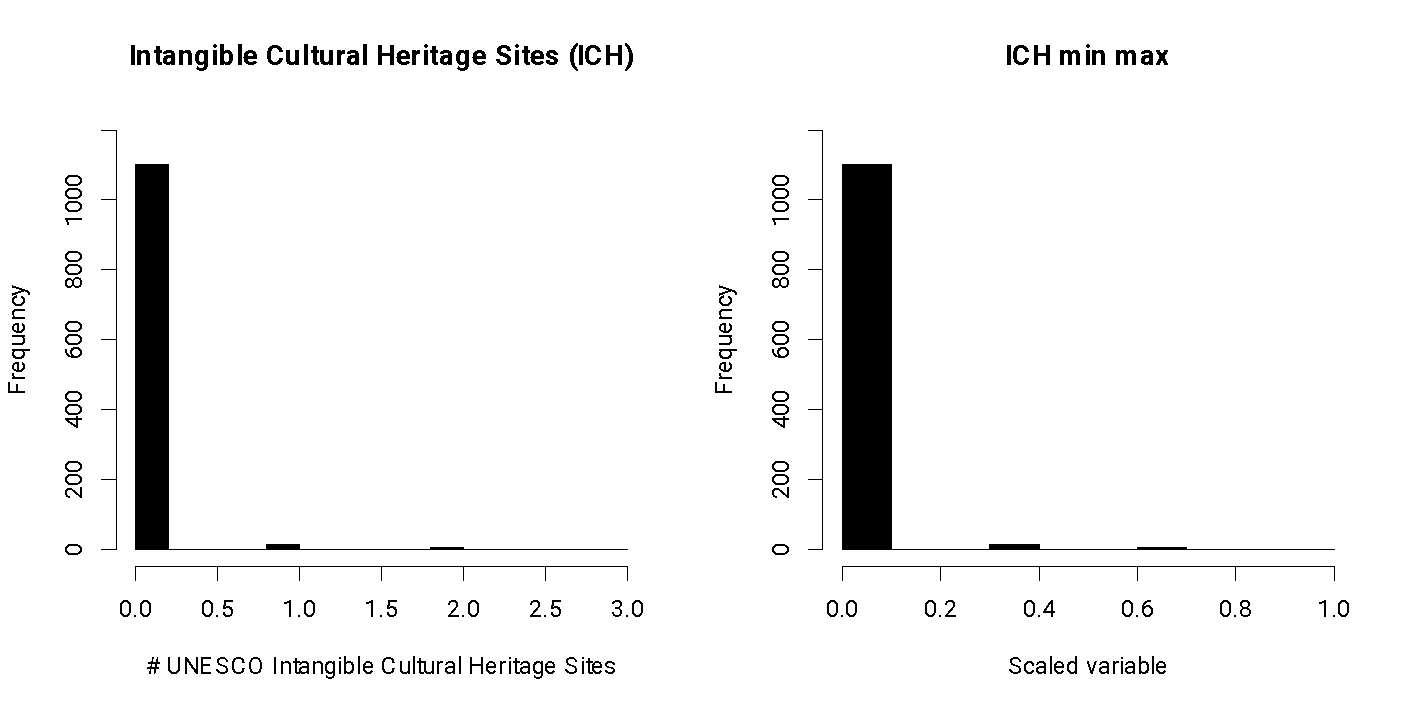

**Figure S21.** **Histograms of UNESCO Intangible Heritage Sites.**

Left panel is the untransformed variable distribution, and the right panel is the transformed variable distribution.

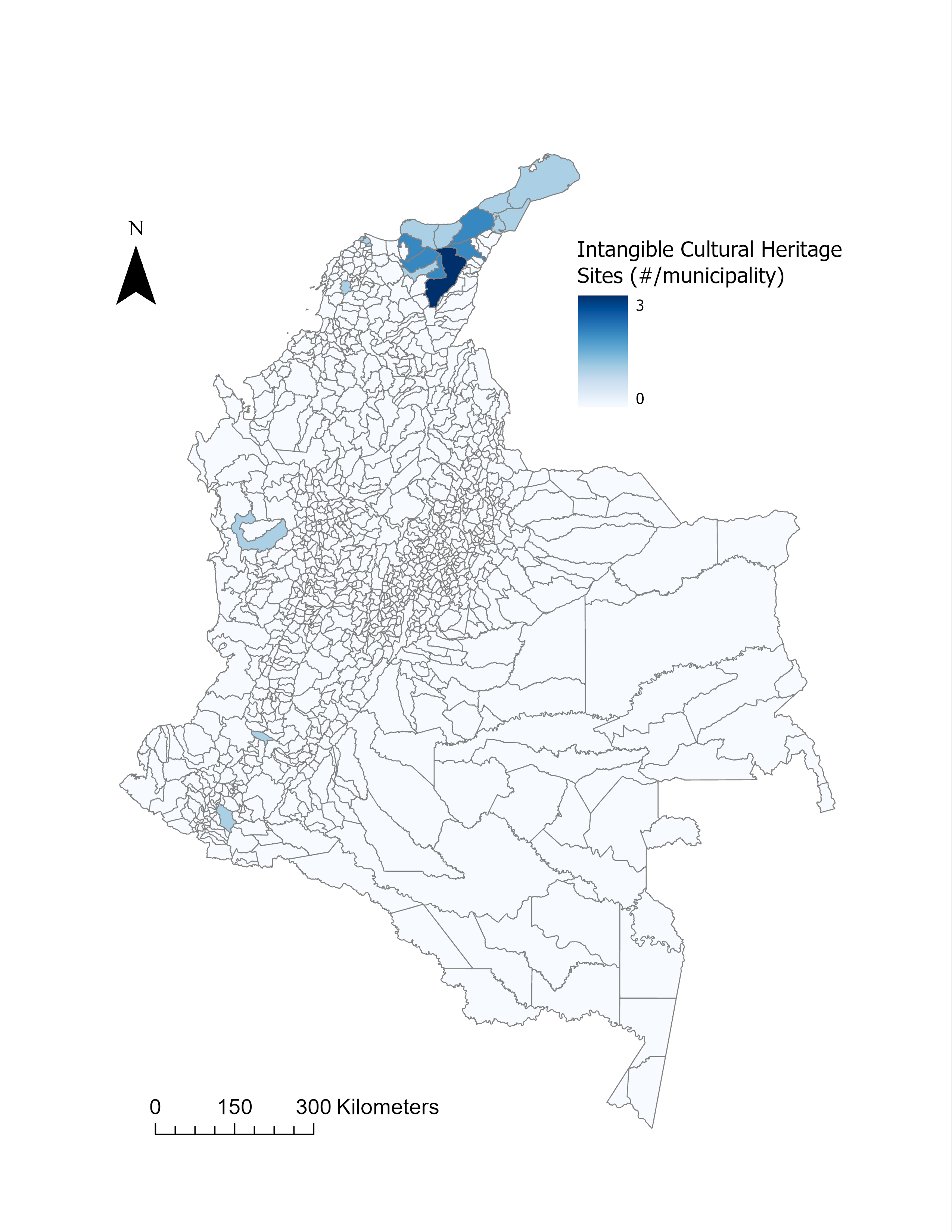

**Figure S22. Spatial distribution of UNESCO intangible cultural heritage sites**

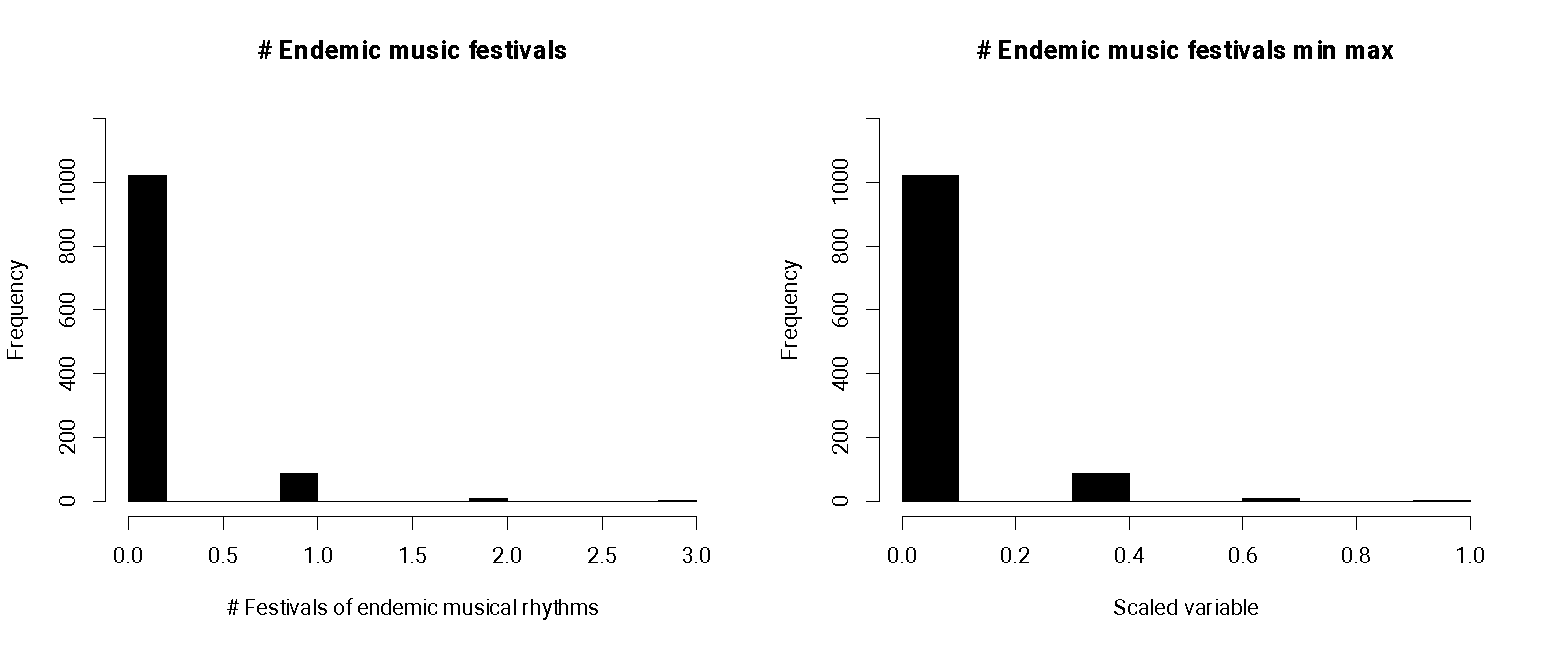

**Figure S23.** **Histograms of endemic rhythm music festivals.**

Left panel is the untransformed variable distribution, and the right panel is the transformed variable distribution.

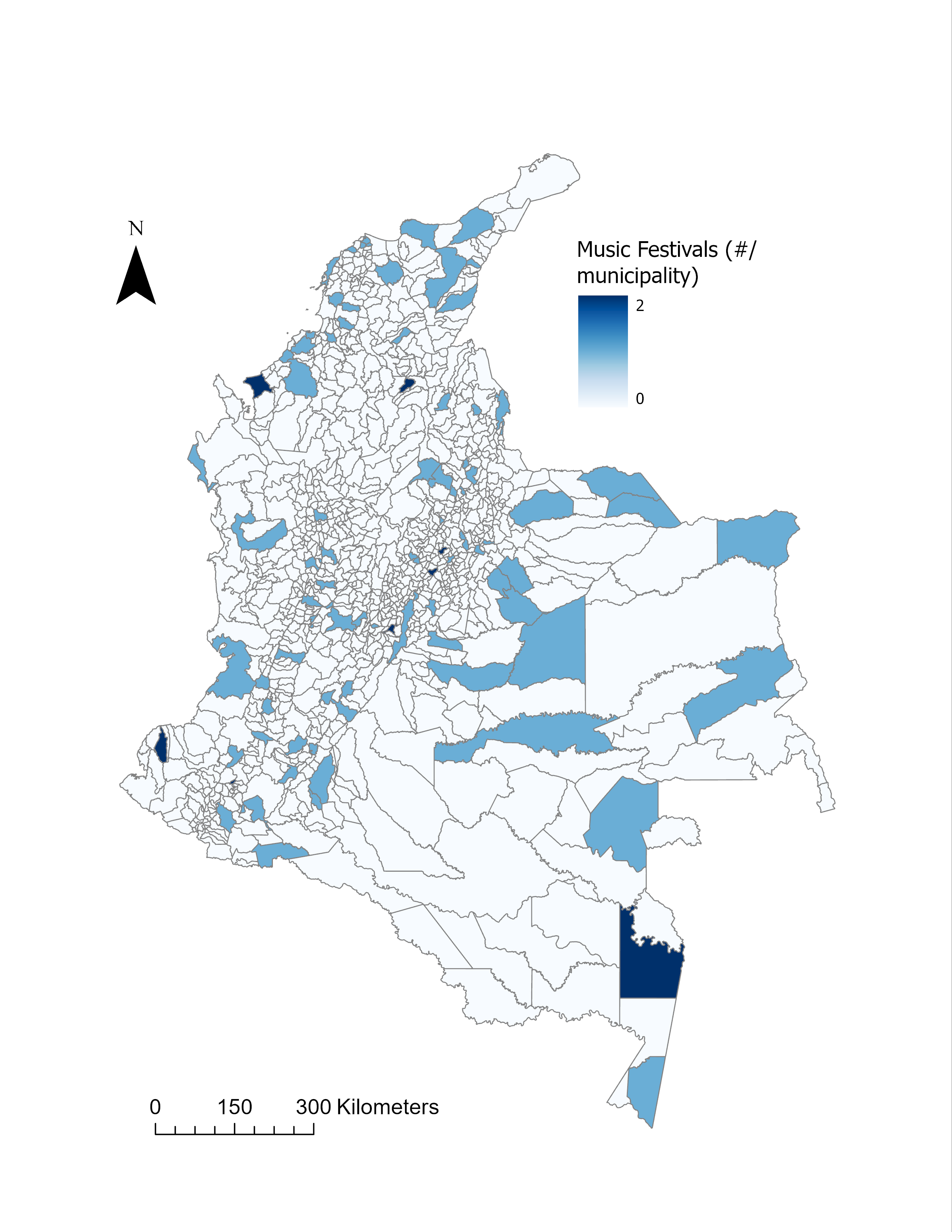

**Figure S24. Spatial distribution of endemic music festivals.**

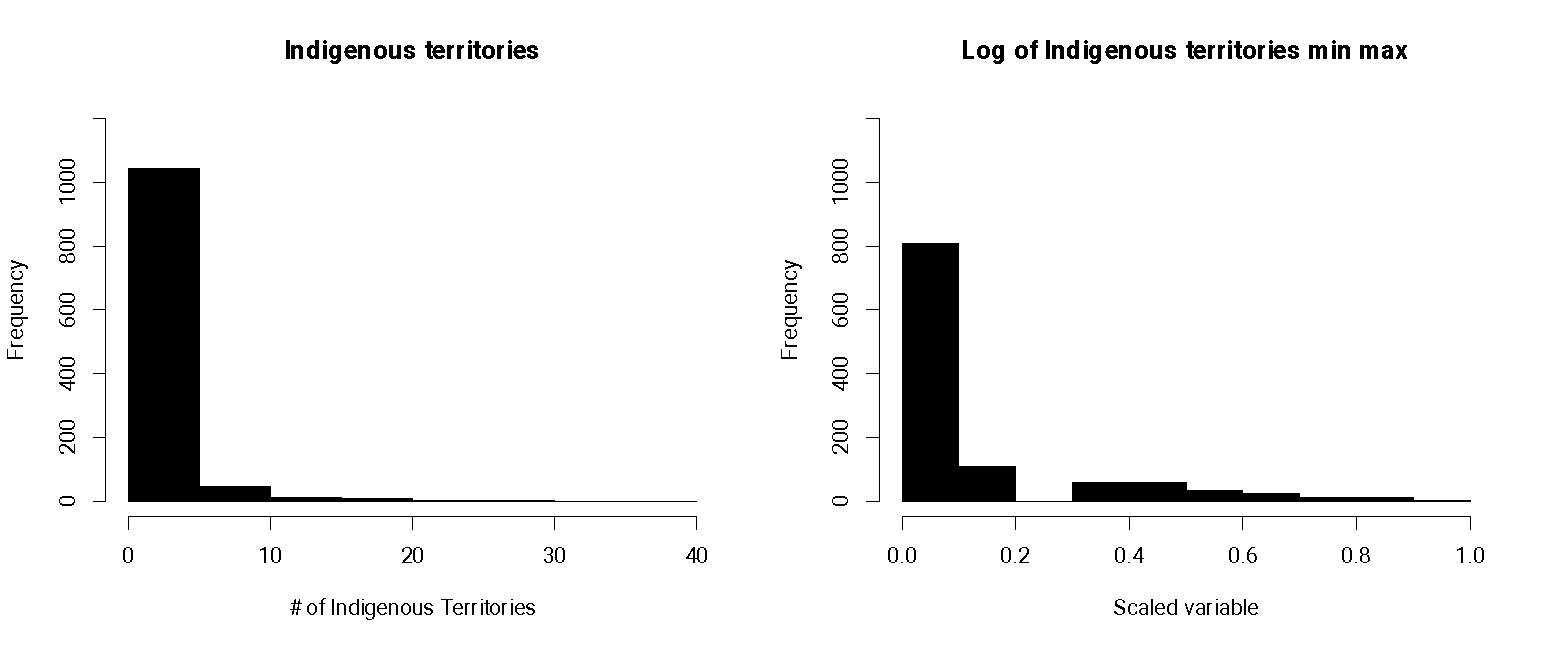

**Figure S25.** **Histograms of Indigenous territories.**

Left panel is the untransformed variable distribution, and the right panel is the transformed variable distribution.

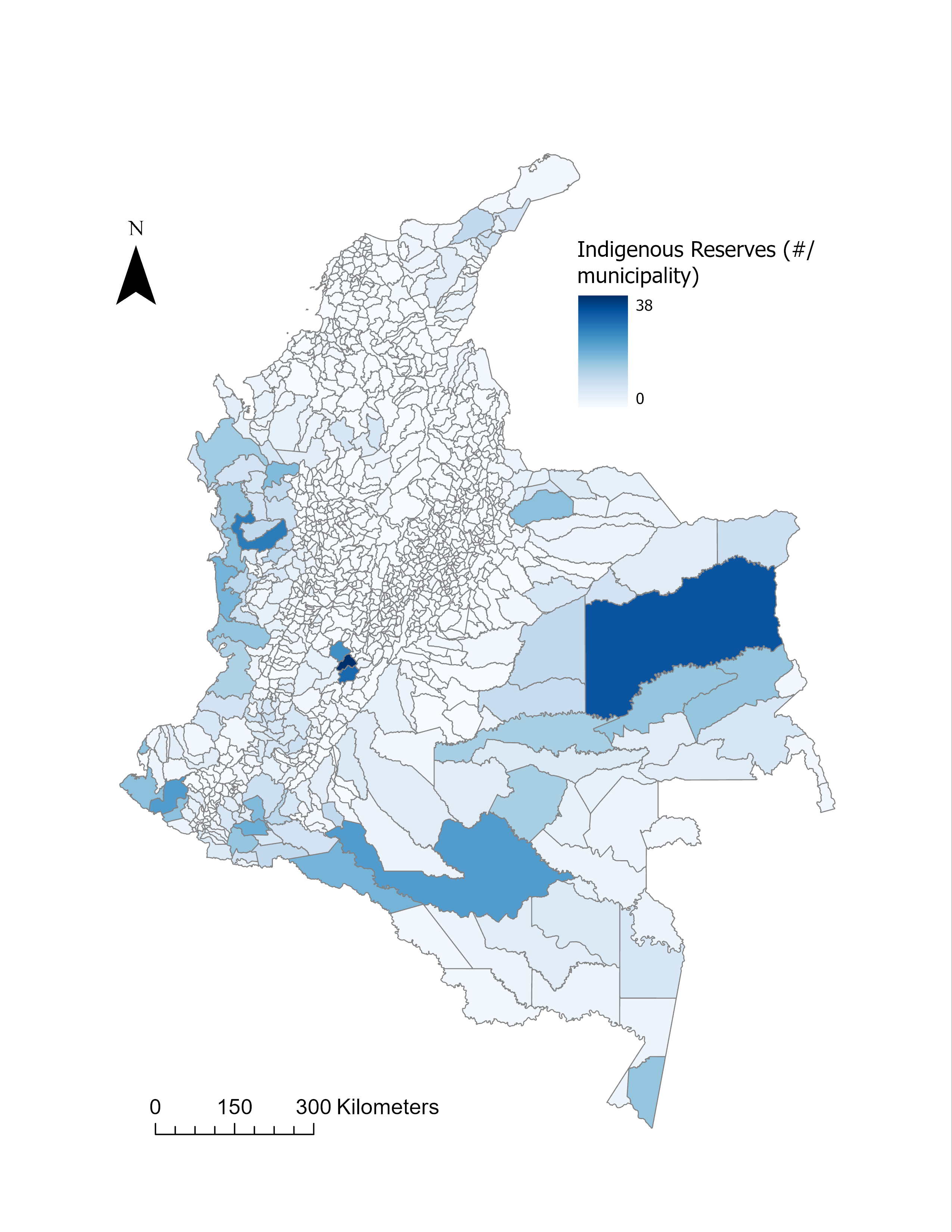

**Figure S26. Spatial distribution of Indigenous reserves.**

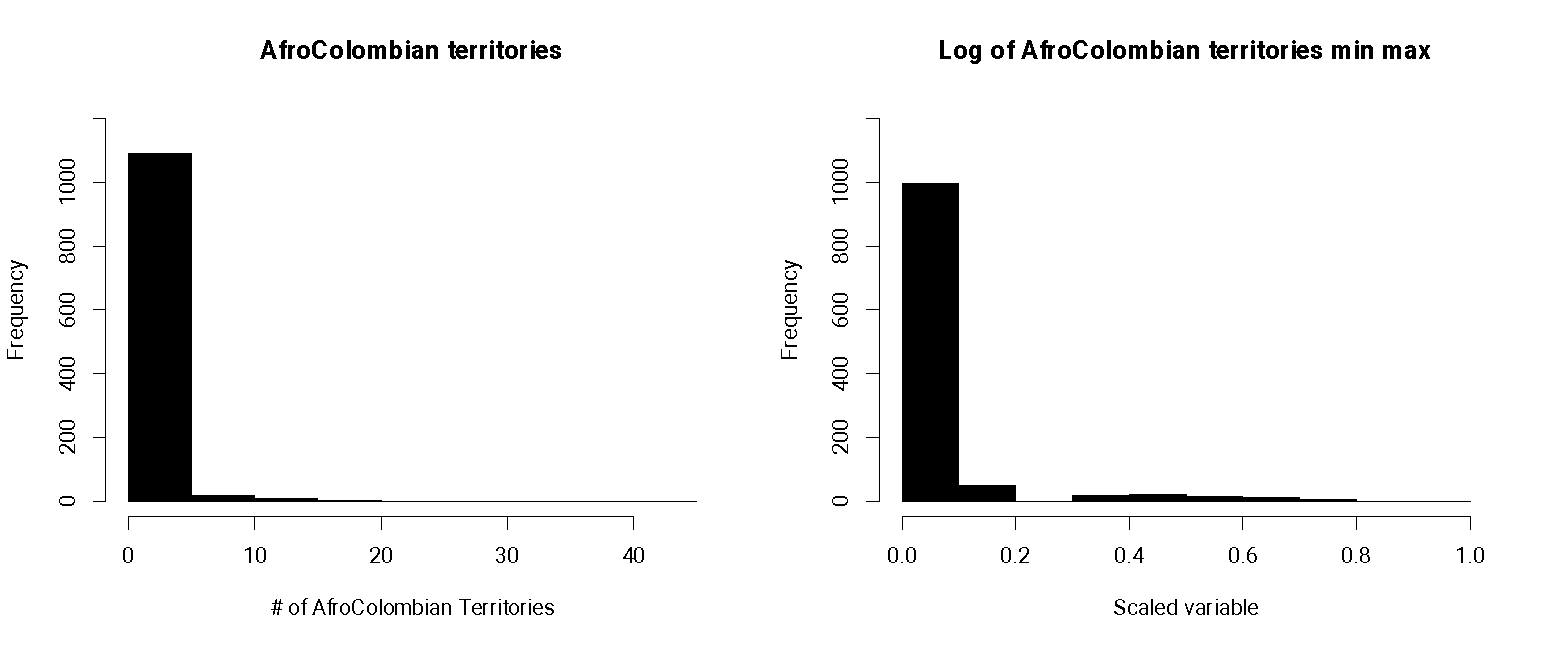

**Figure S27.** **Histograms of Afro-Colombian territories.**

Left panel is the untransformed variable distribution, and the right panel is the transformed variable distribution.

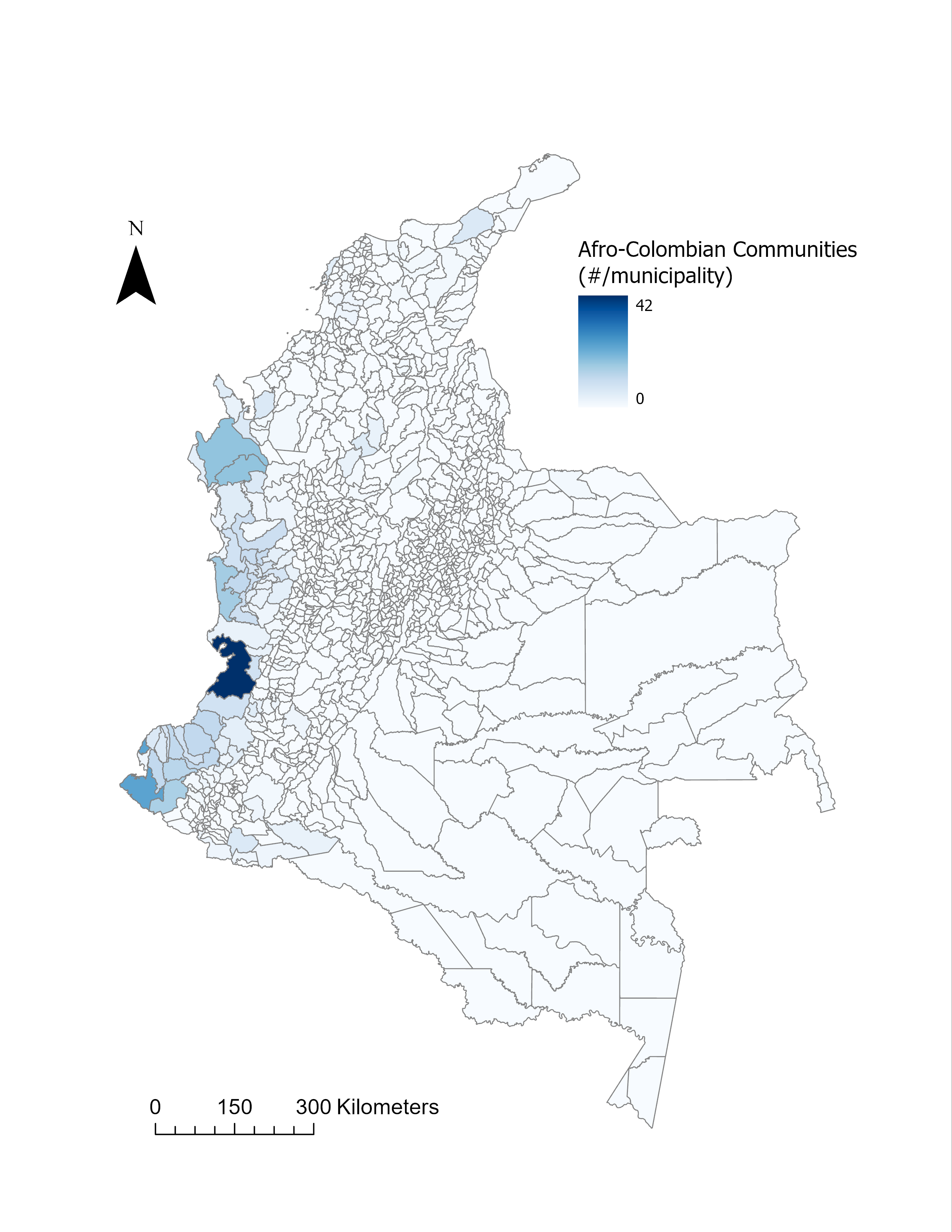

**Figure S28. Spatial distribution of Afro-Colombian territories.**

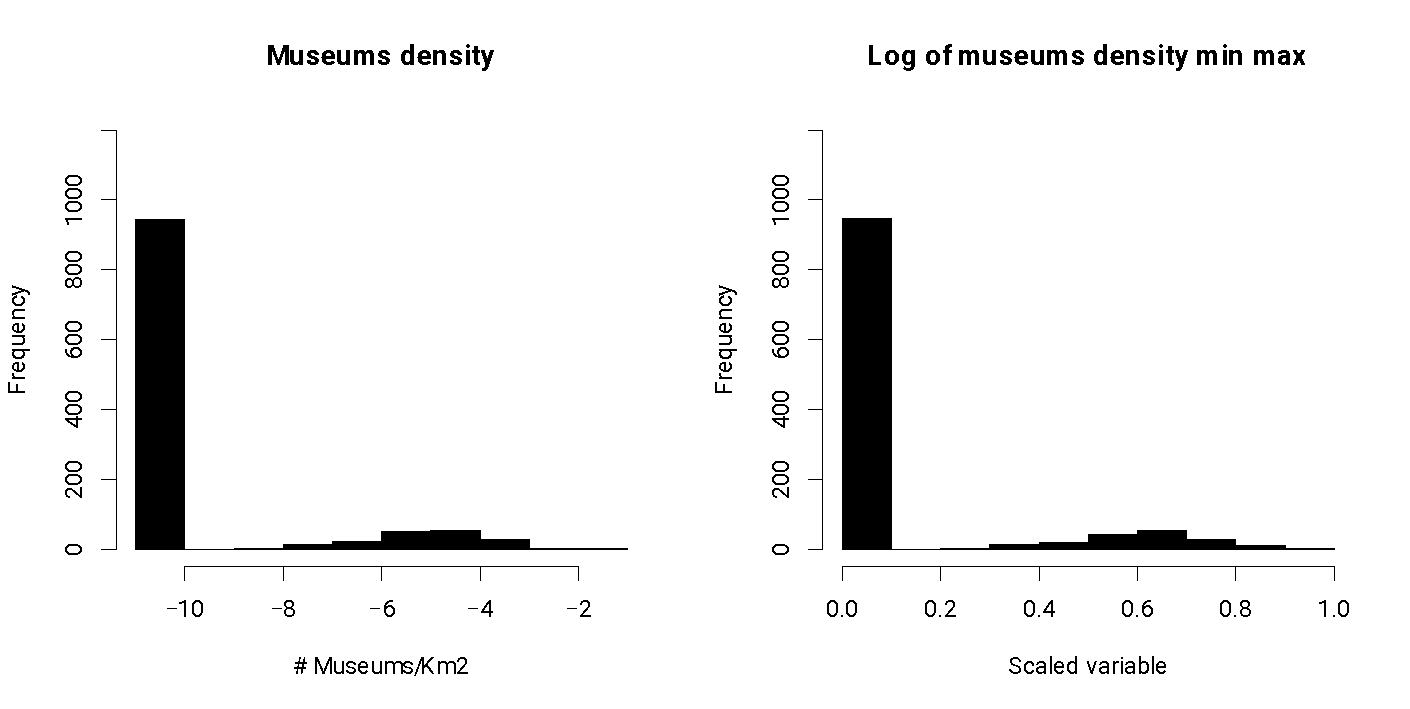

**Figure S29. Histograms of museum density.**

Left panel is the untransformed variable distribution, and the right panel is the transformed variable distribution.

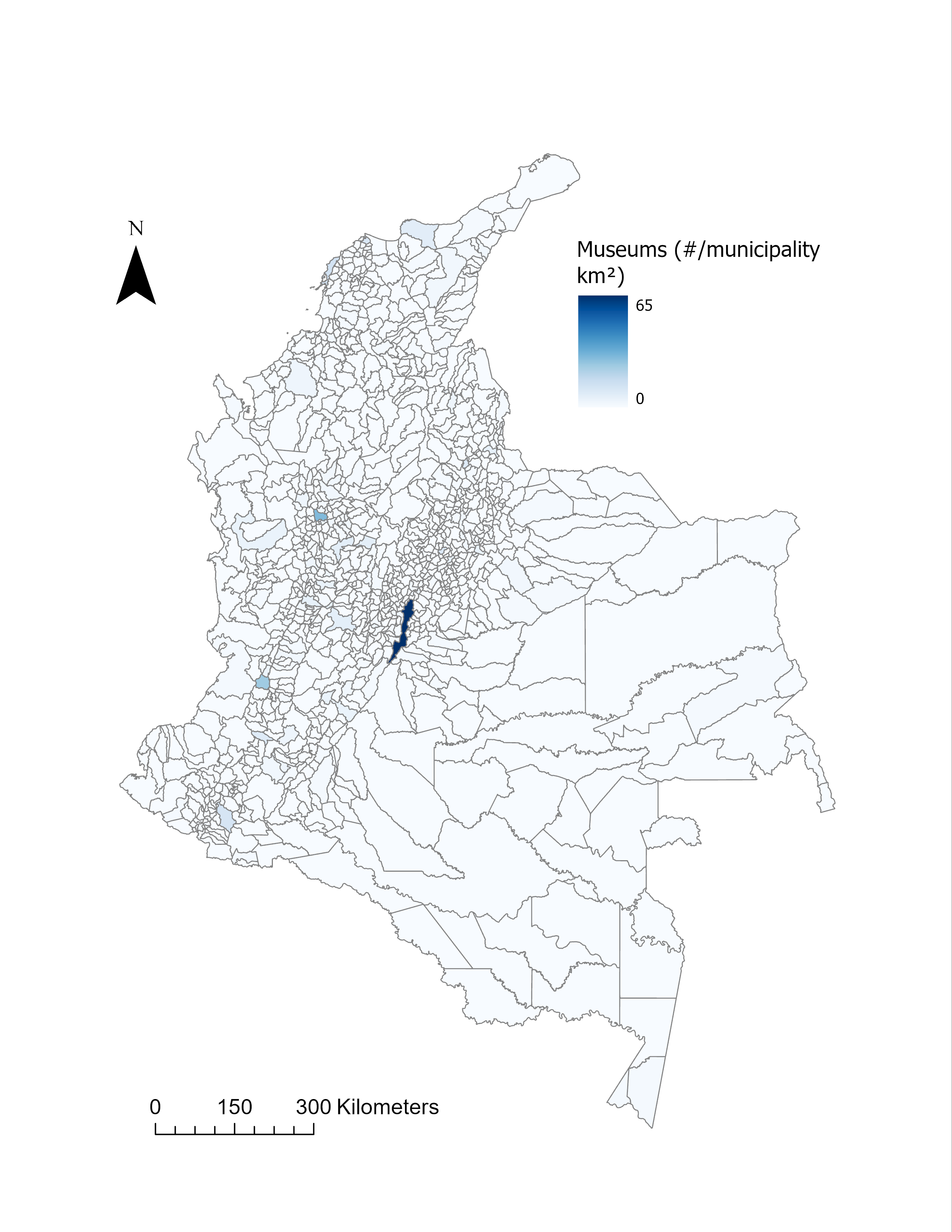

**Figure S30. Spatial distribution of museums.**

#### 3.4. Accessibility variables

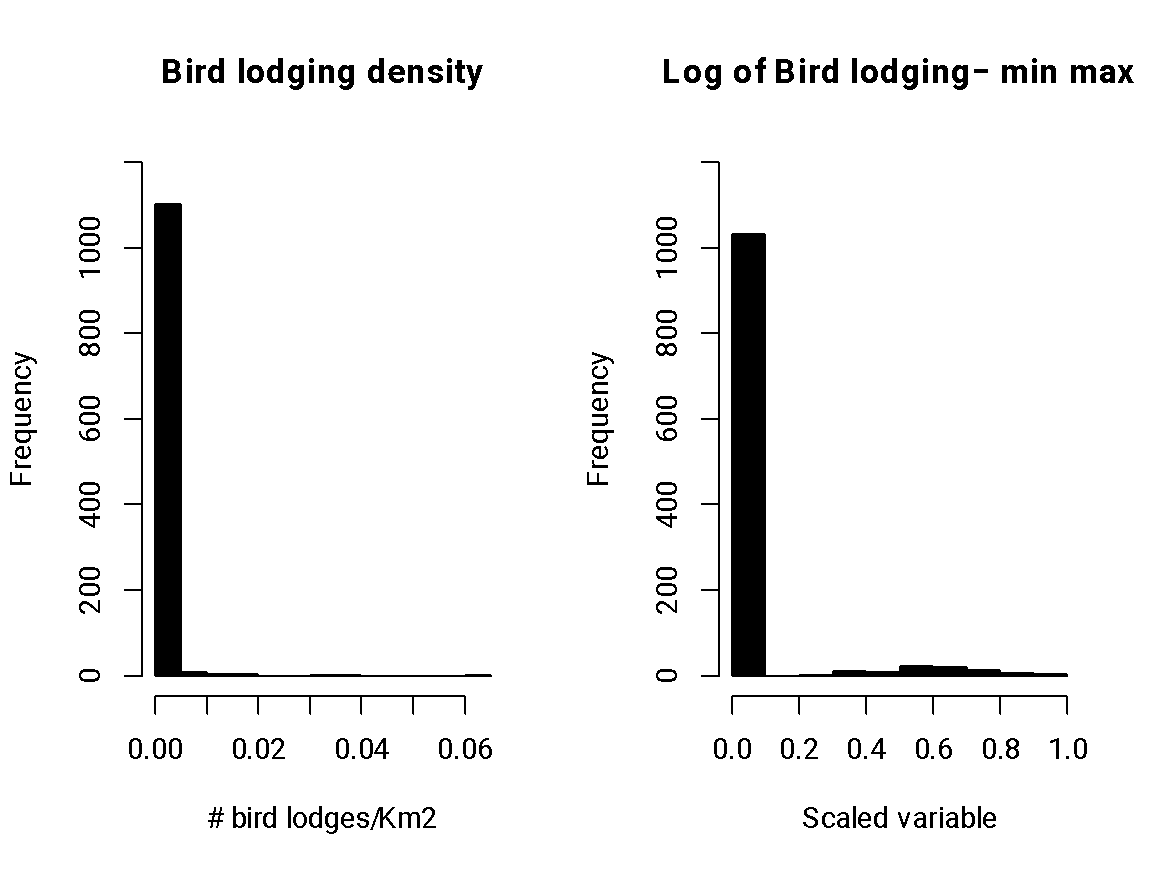

**Figure S31.** **Histograms of bird lodges.**

Left panel is the untransformed variable distribution, and the right panel is the transformed variable distribution.

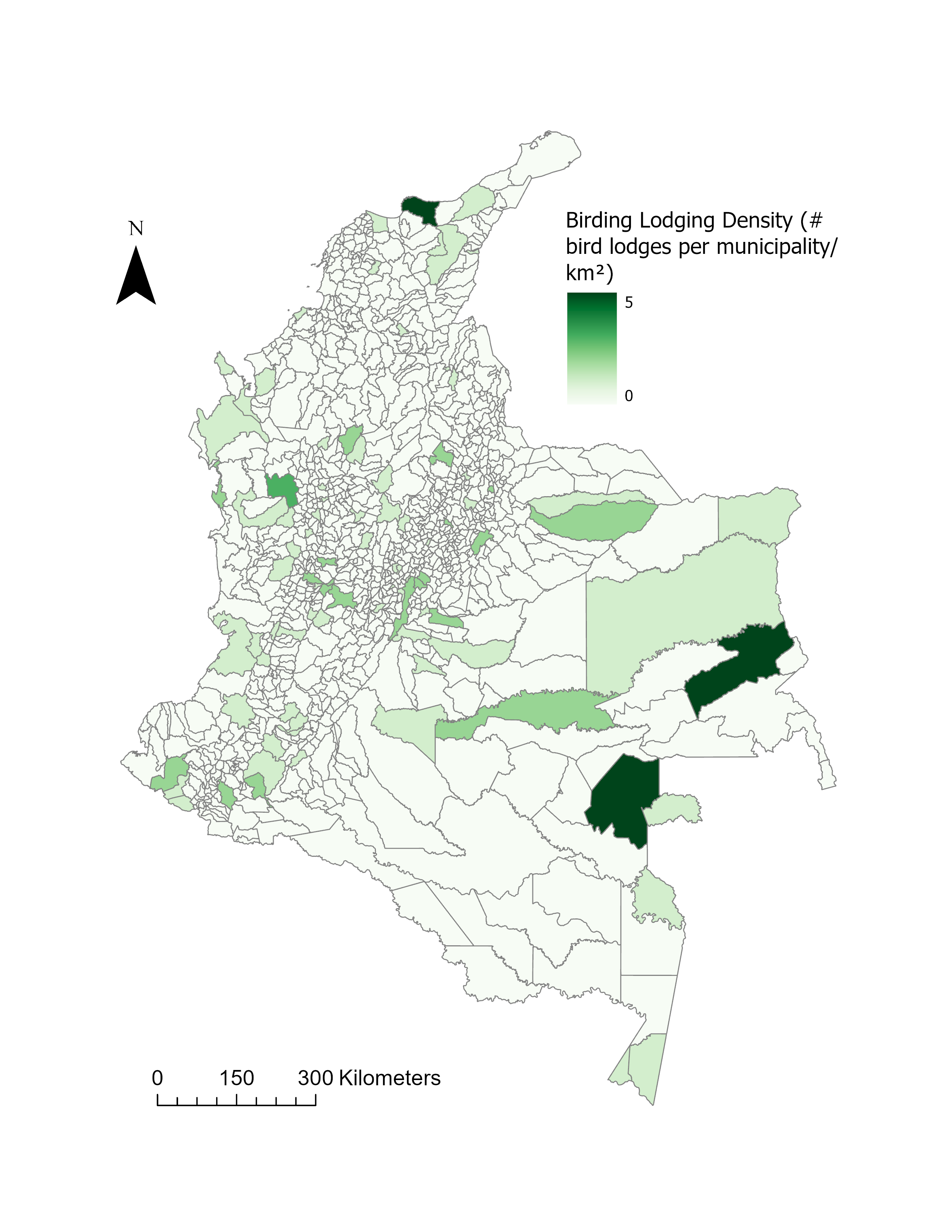

**Figure S32. Spatial distribution of bird lodging locations.**

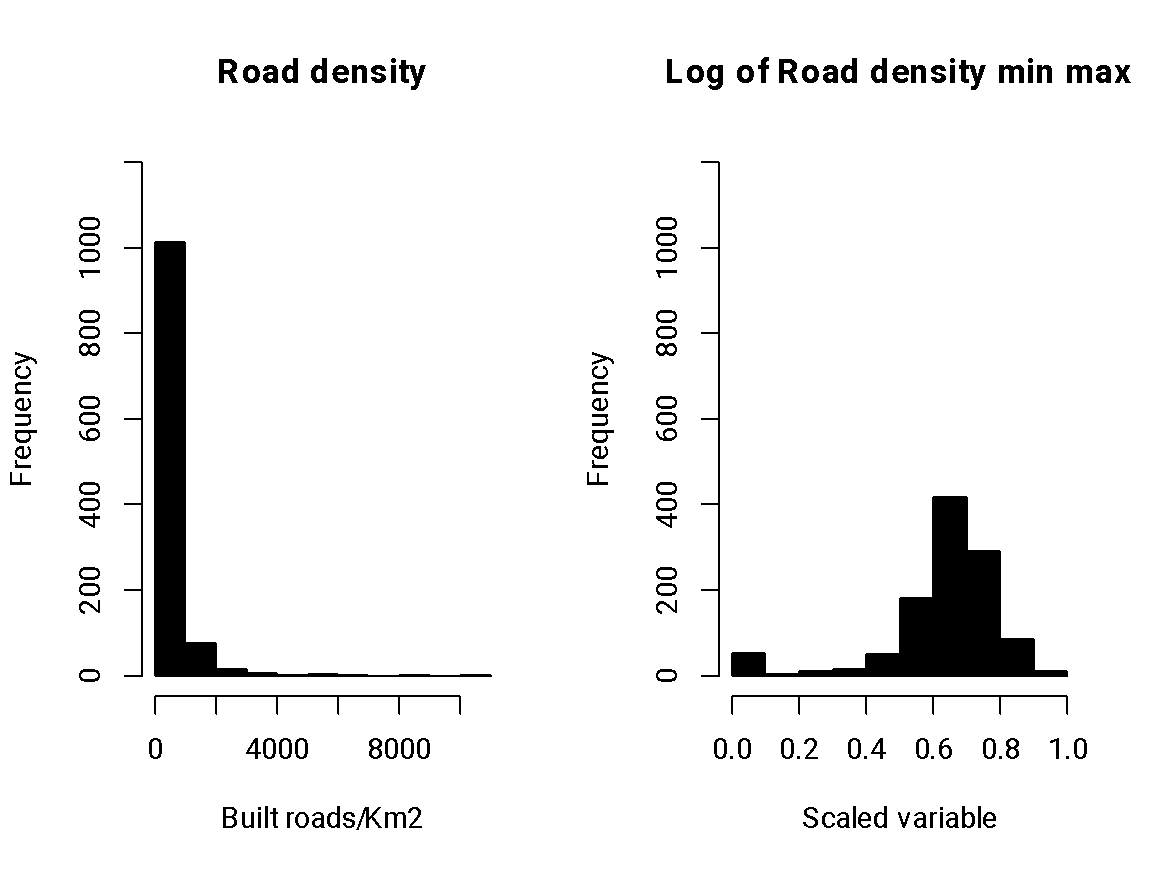

**Figure S33.** **Histograms of road density.**

Left panel is the untransformed variable distribution, and the right panel is the transformed variable distribution.

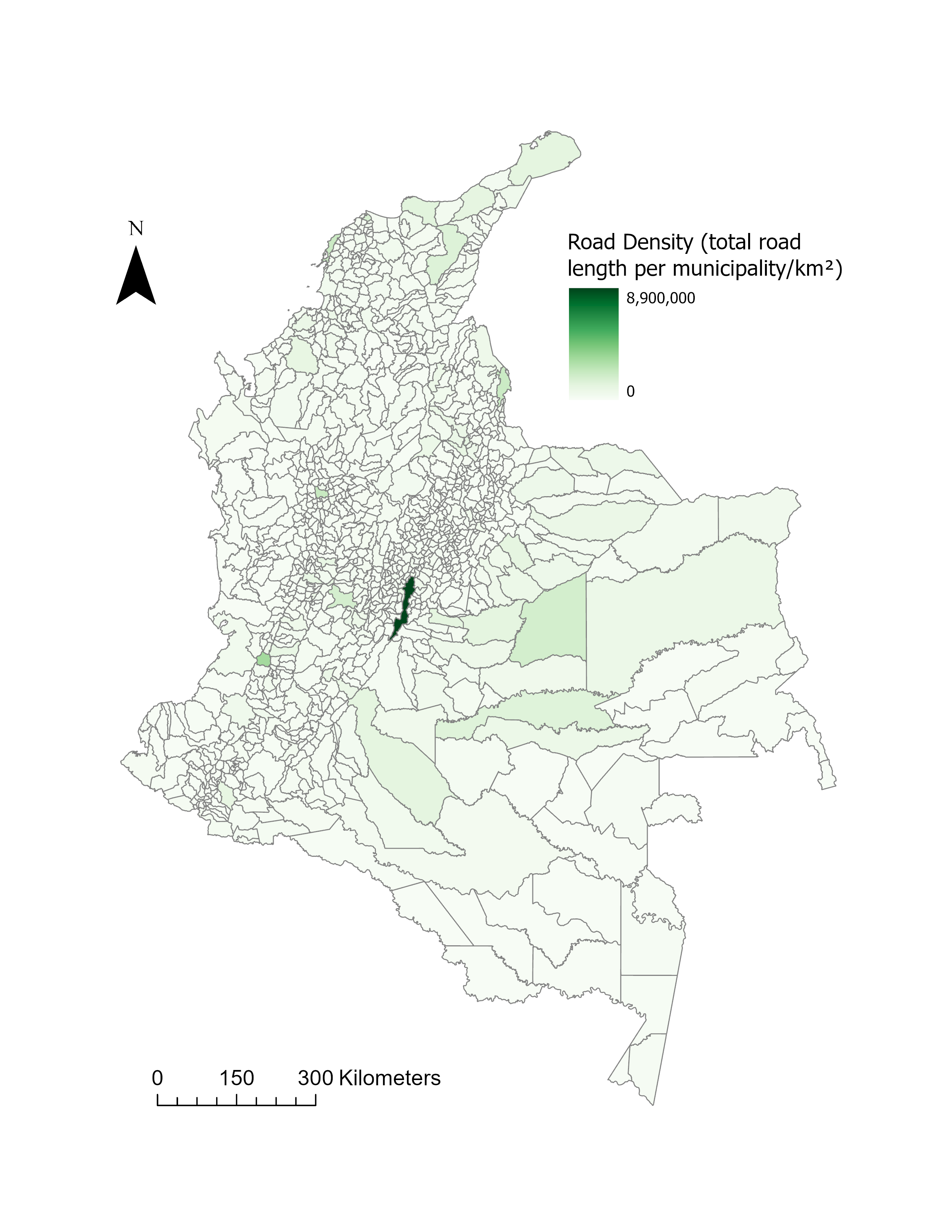

**Figure S34. Spatial distribution of road density**

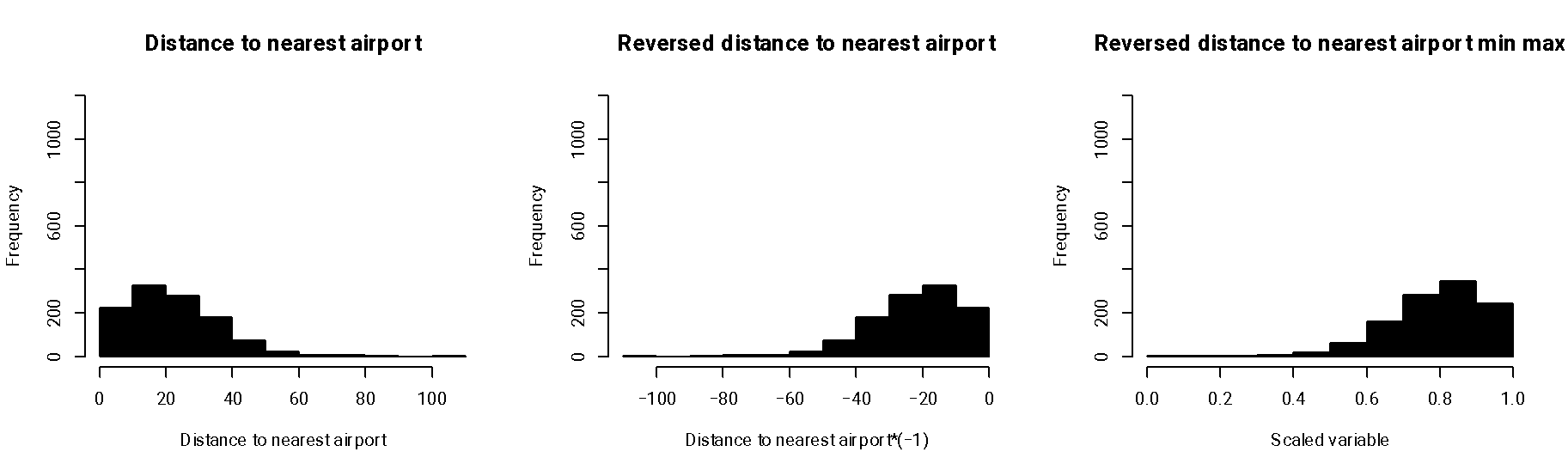

**Figure S35.** **Histograms of distance to nearest airport.**

Left panel is the untransformed variable distribution, middle panel is the reversed variable to express proximity to an airport, and right panel is the transformed variable distribution.

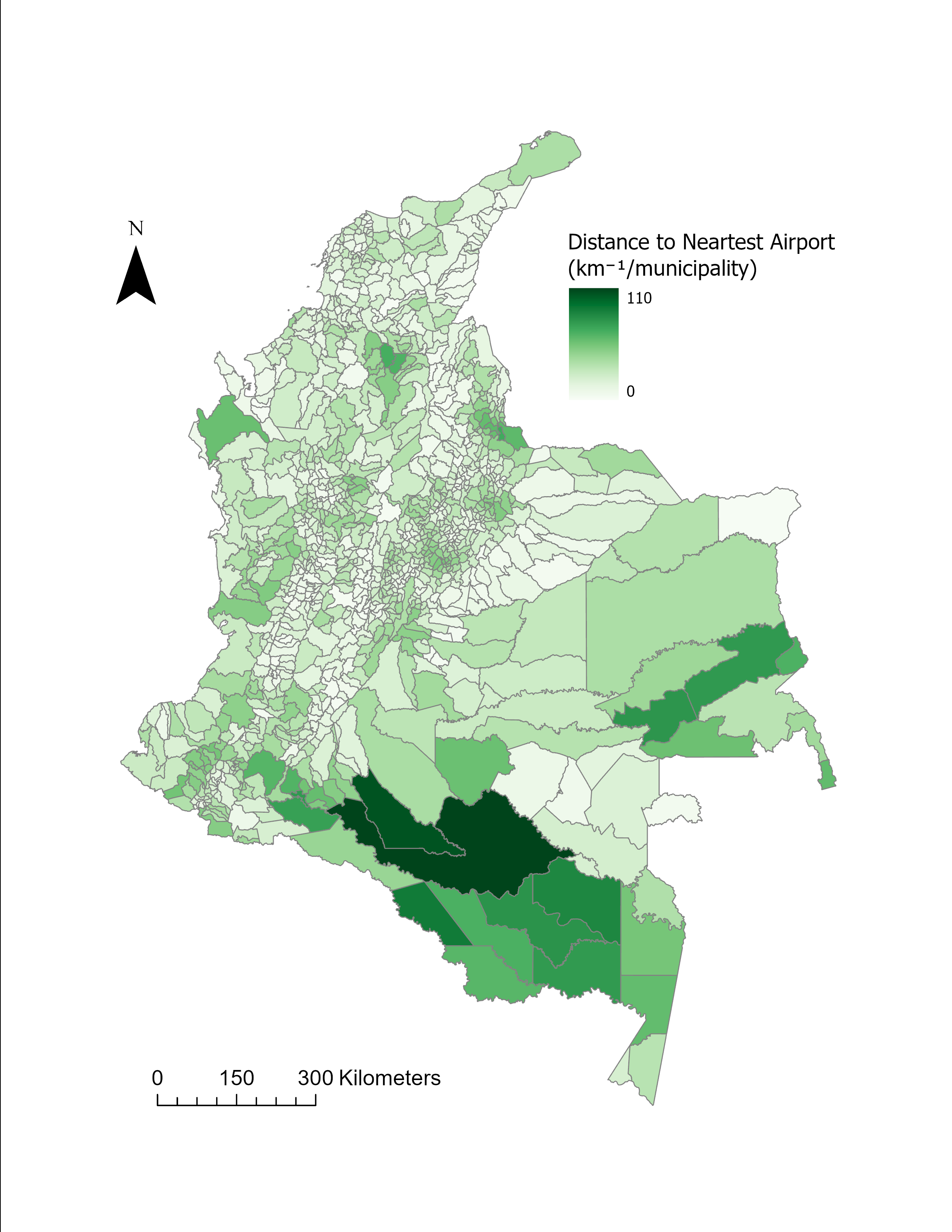

**Figure S36. Spatial distribution of the distance to the nearest airport.**

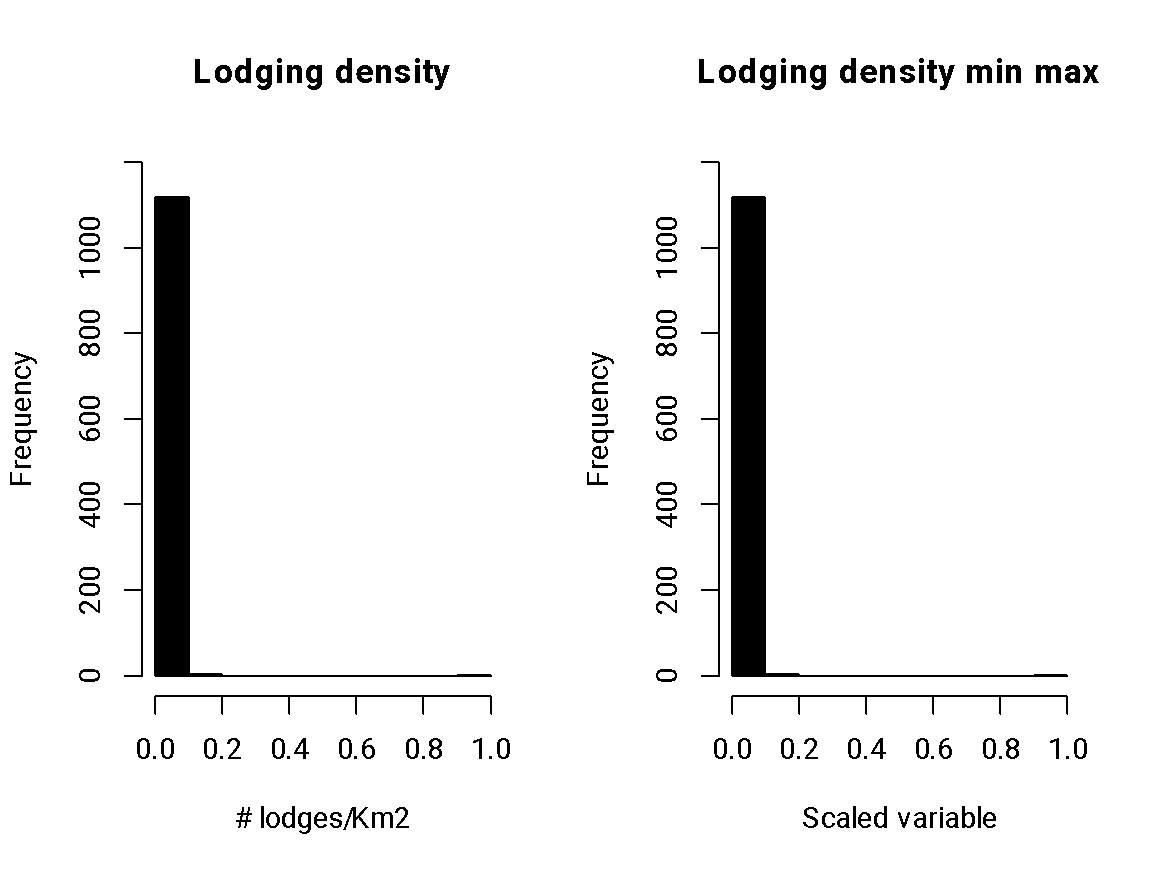

**Figure S37.** **Histograms of lodging density.**

Left panel is the untransformed variable distribution, and the right panel is the transformed variable distribution.

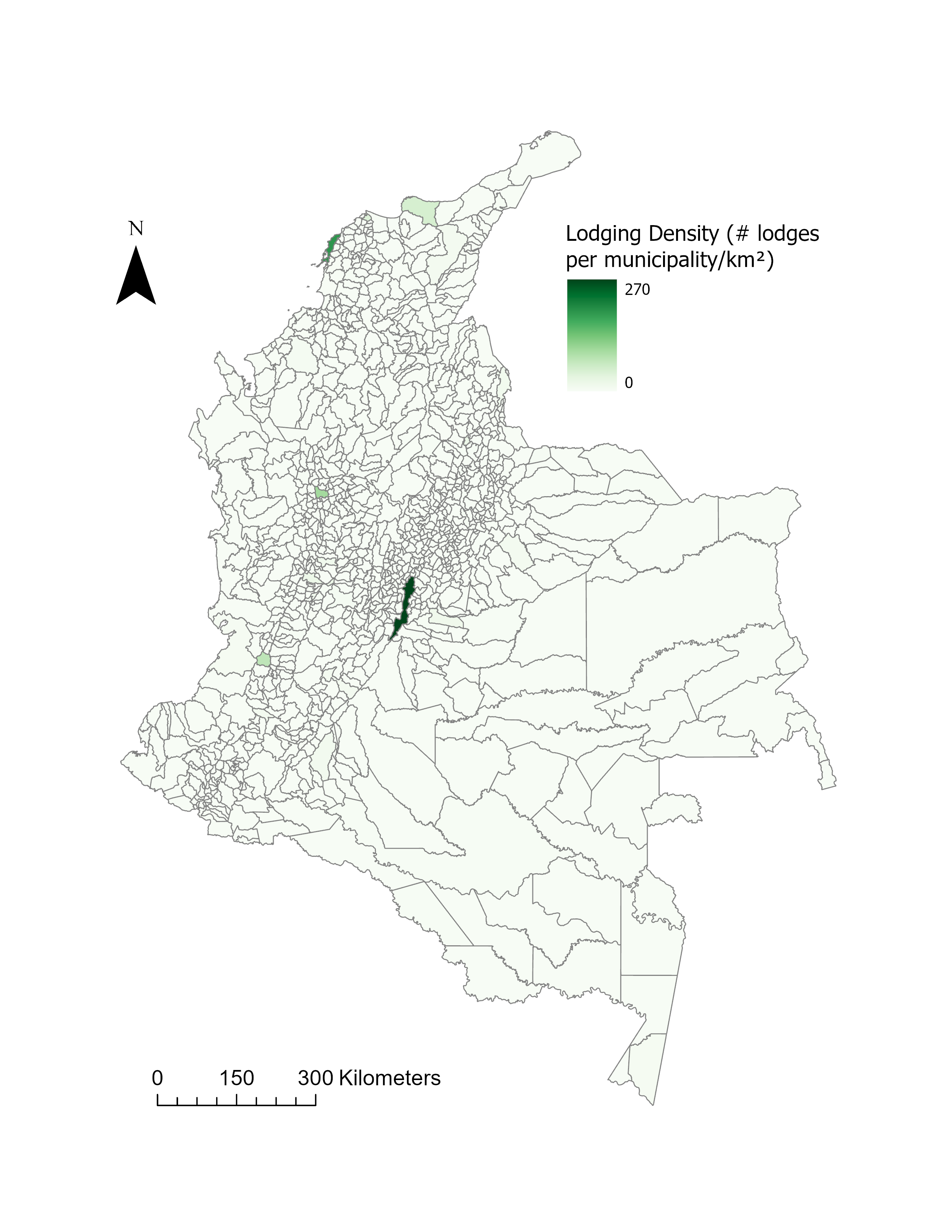

**Figure S38. Spatial distribution of lodging density.**

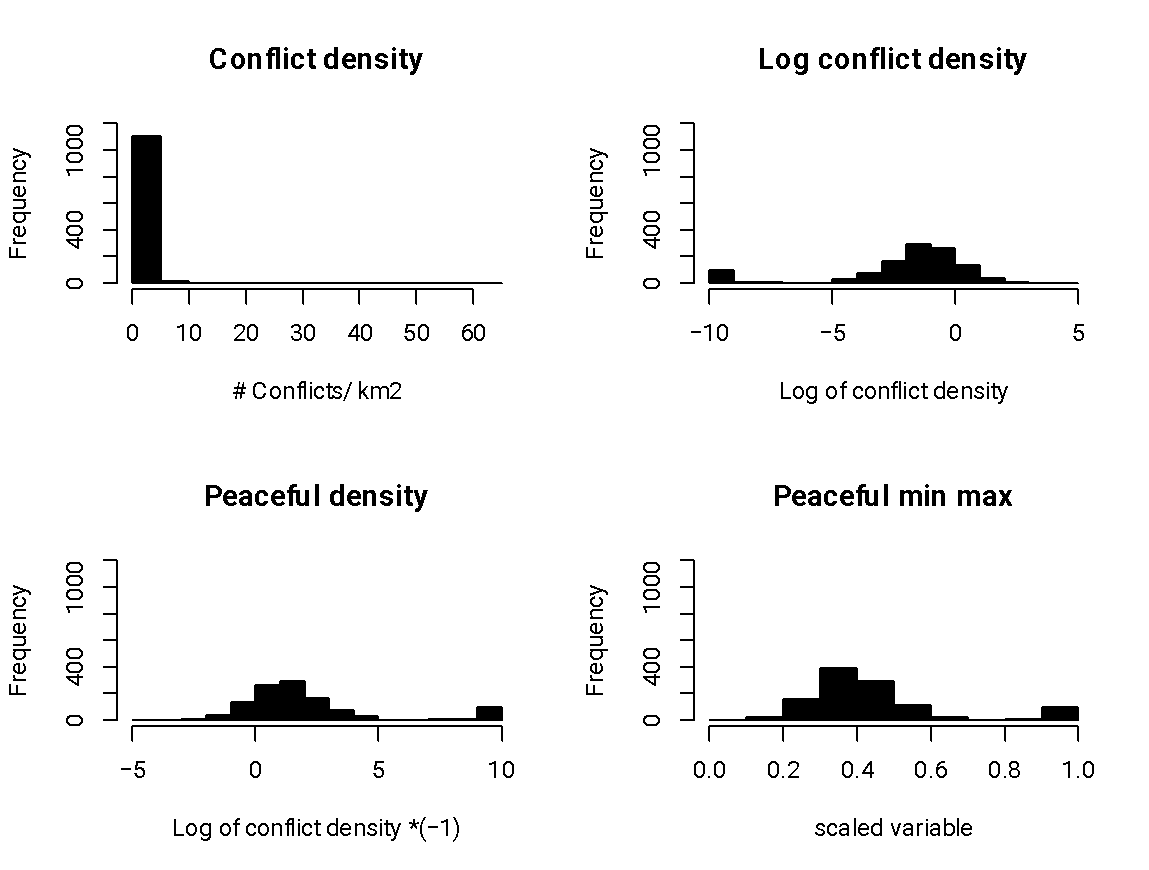

**Figure S39. Histograms of conflict density and peaceful variable.**

Top left panel is the untransformed variable distribution, the top right panel is the transformed variable with logarithmic transformation, bottom left panel is calculated as a density, and bottom right panel is the density transformed with a min max transformation, which was used in subsequent analyses.

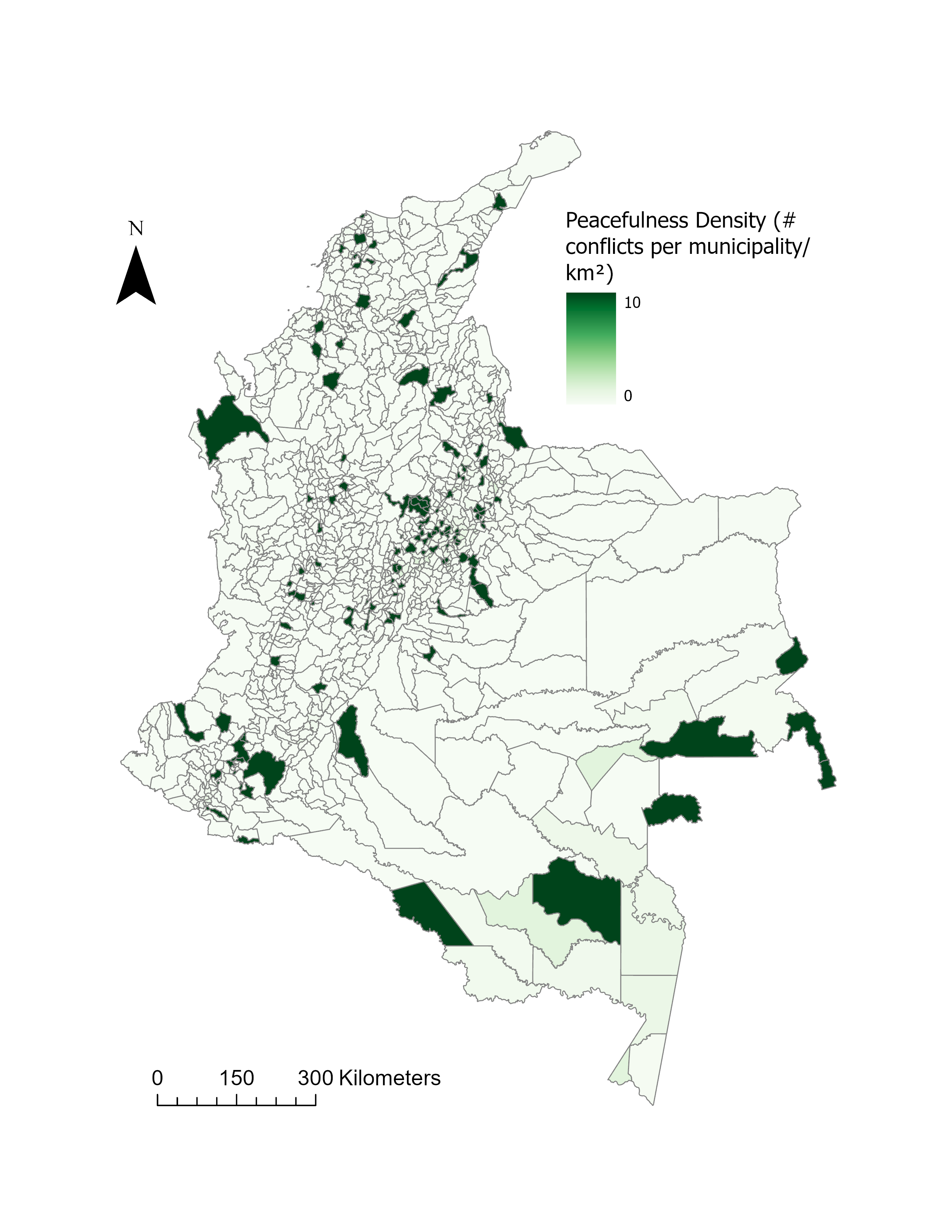

**Figure S40. Spatial distribution of peacefulness.**

#### 3.5. Demand index variables

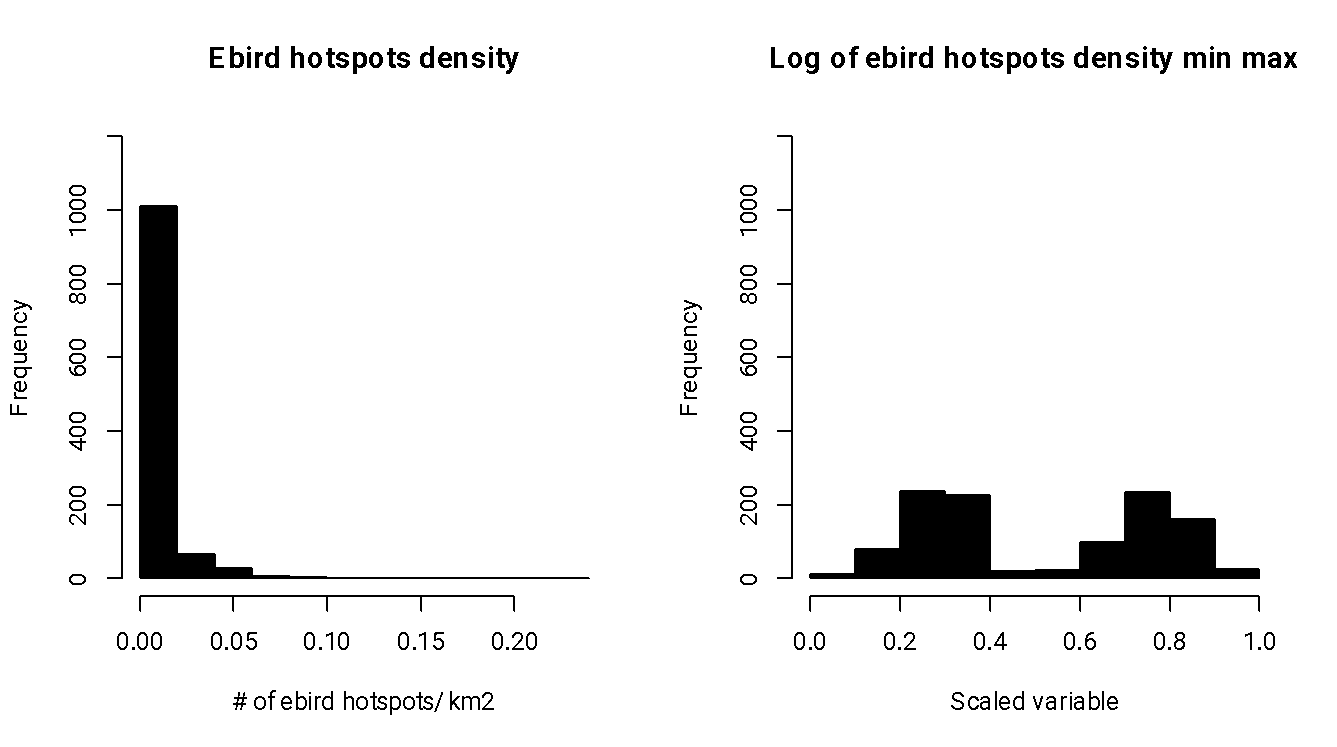

**Figure S41.** **Histograms of eBird hotspots.**

Left panel is the untransformed variable distribution, and the right panel is the transformed variable distribution.

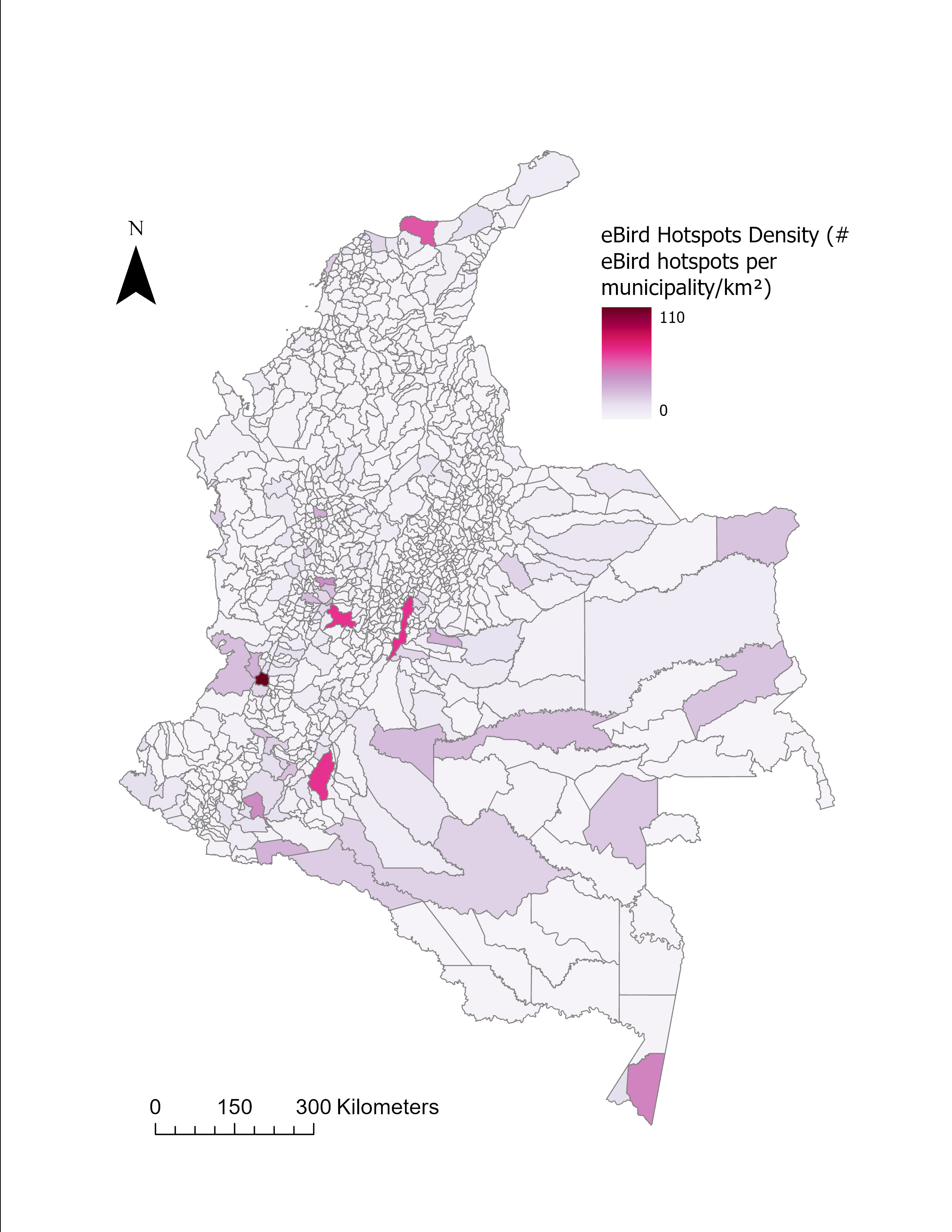

**Figure S42. Spatial distribution of eBird hotspots.**

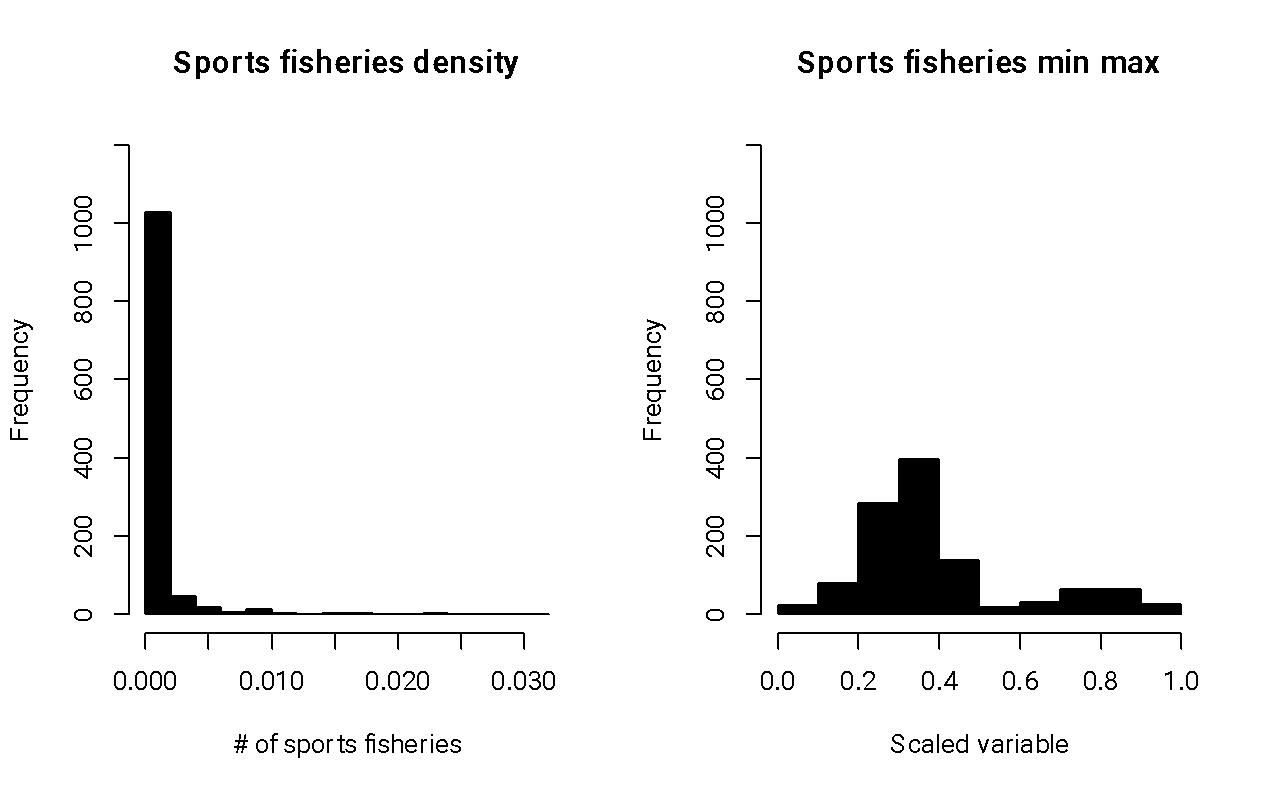

**Figure S43.** **Histograms of sports fisheries.**

Left panel is the untransformed variable distribution, and the right panel is the transformed variable distribution.

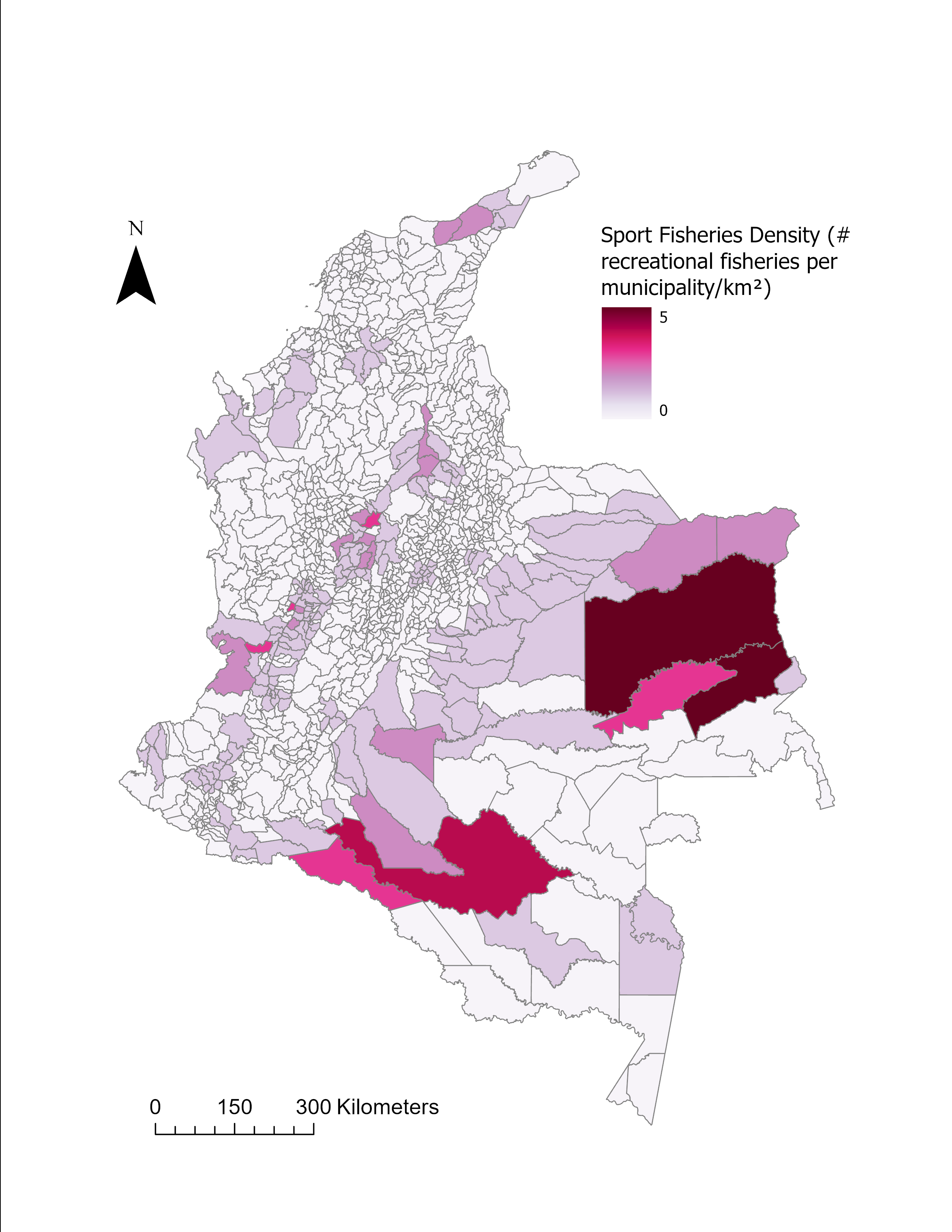

**Figure S44. Spatial distribution of sports fisheries.**

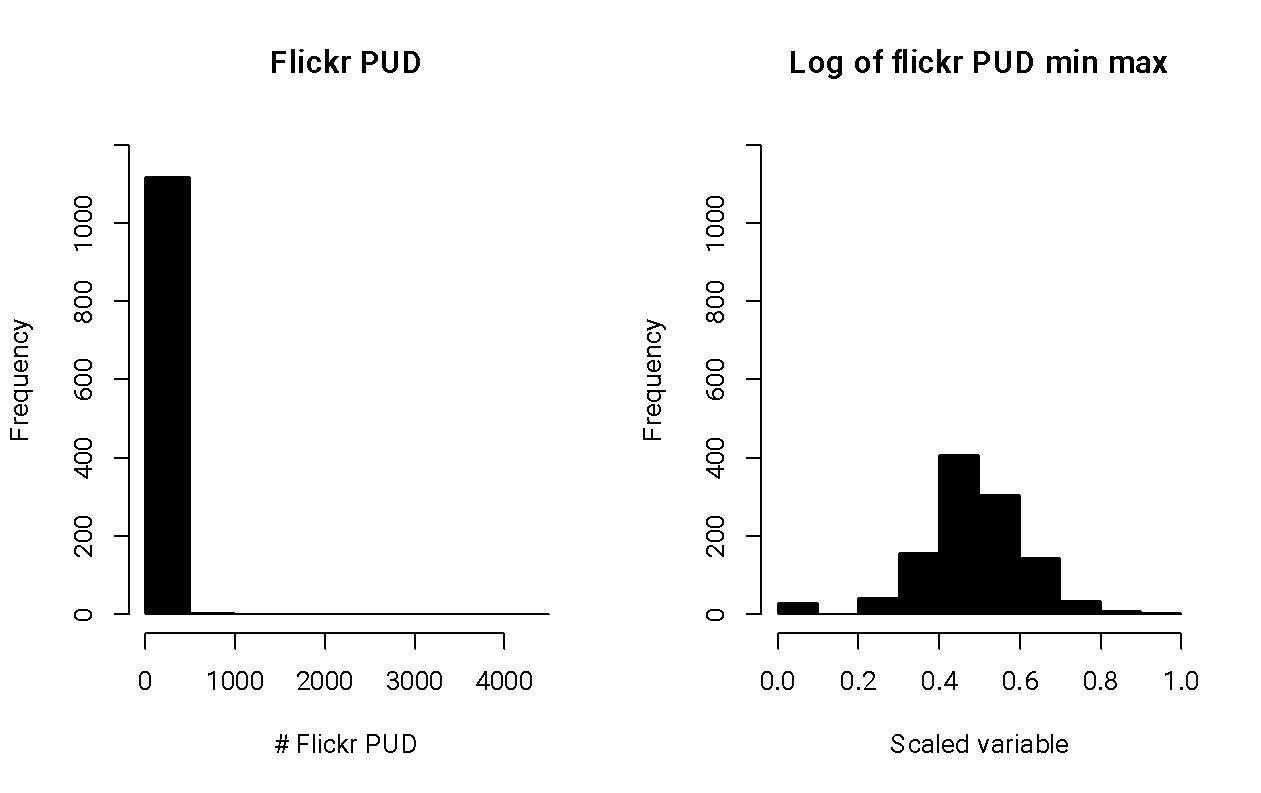

**Figure S45.** **Histograms of Flickr Photo-User-Days.**

Left panel is the untransformed variable distribution, and the right panel is the transformed variable distribution.

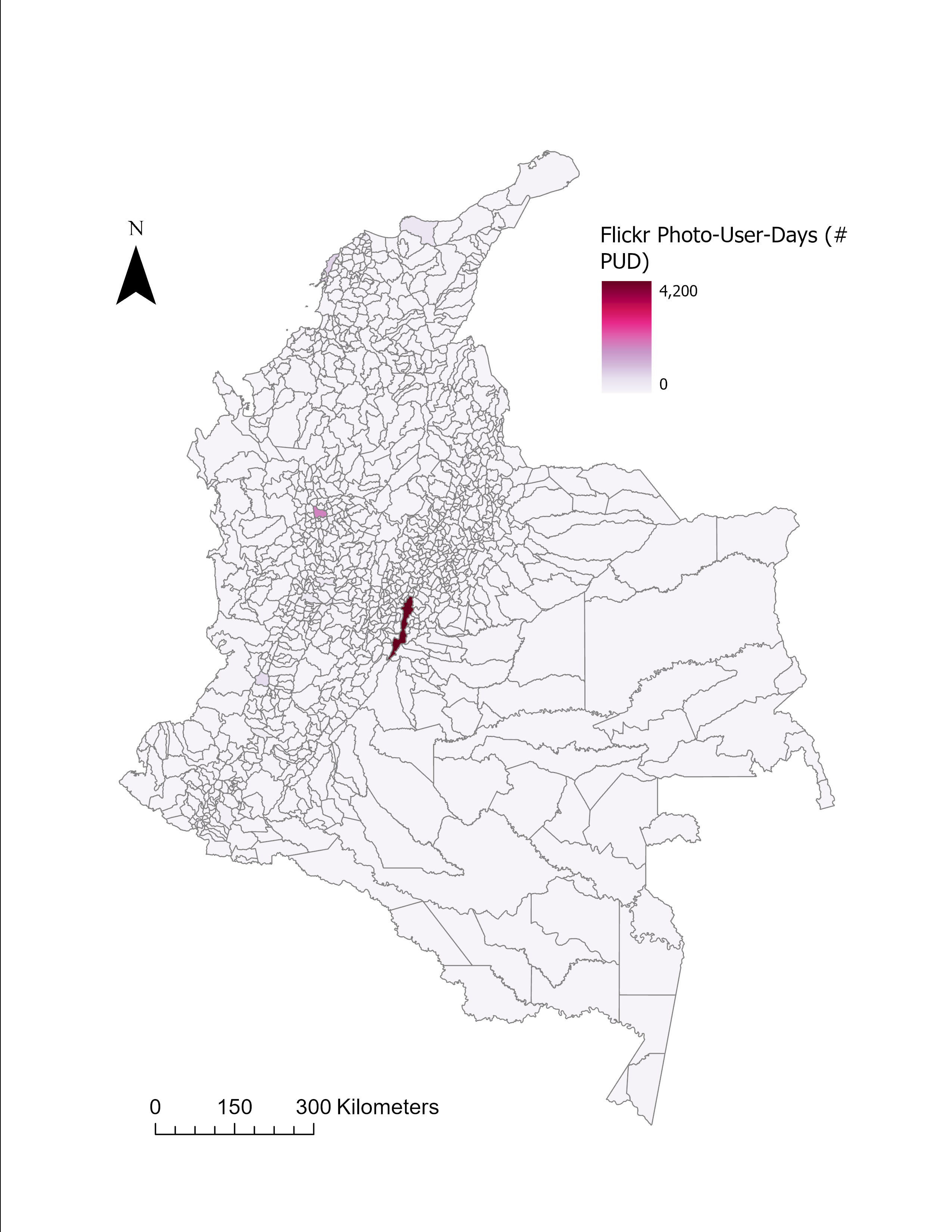

**Figure S46. Spatial distribution of Flickr Photo-User-Days.**

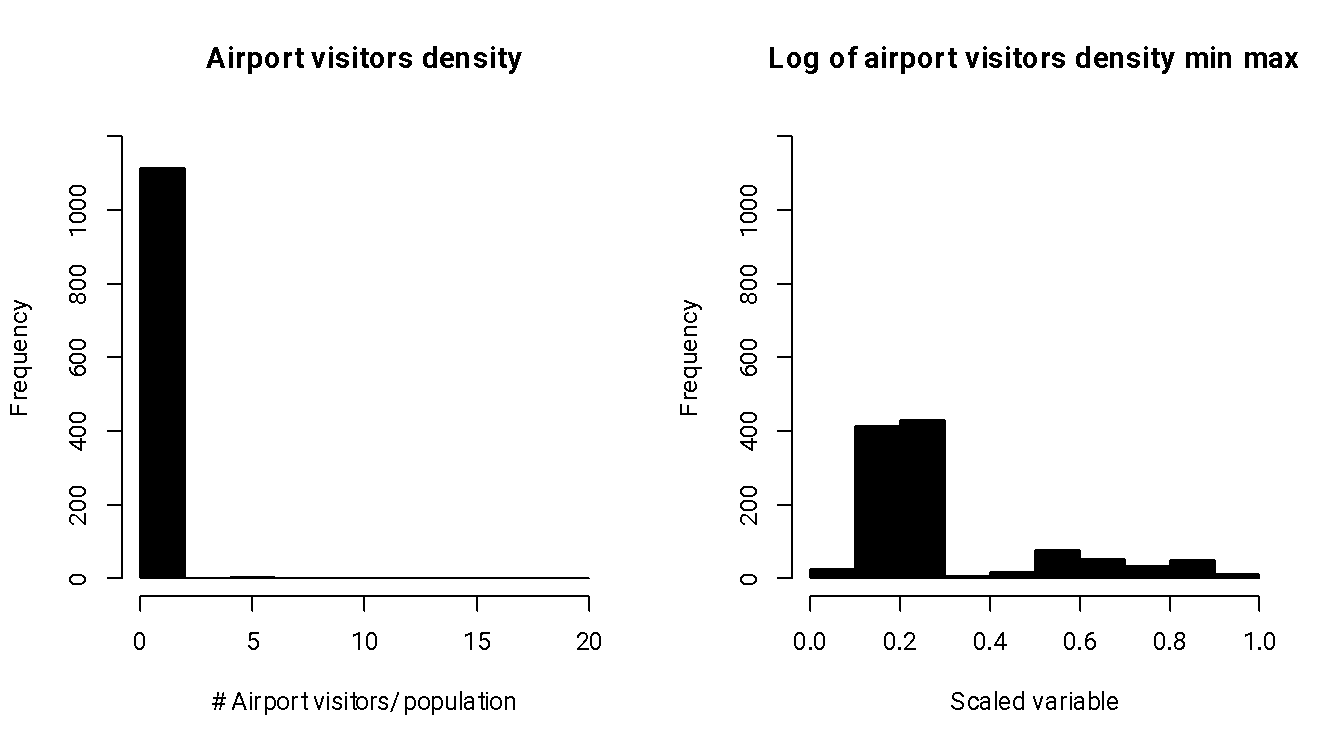

**Figure S47.** **Histograms of airports visits.**

Left panel is the untransformed variable distribution, and the right panel is the transformed variable distribution.

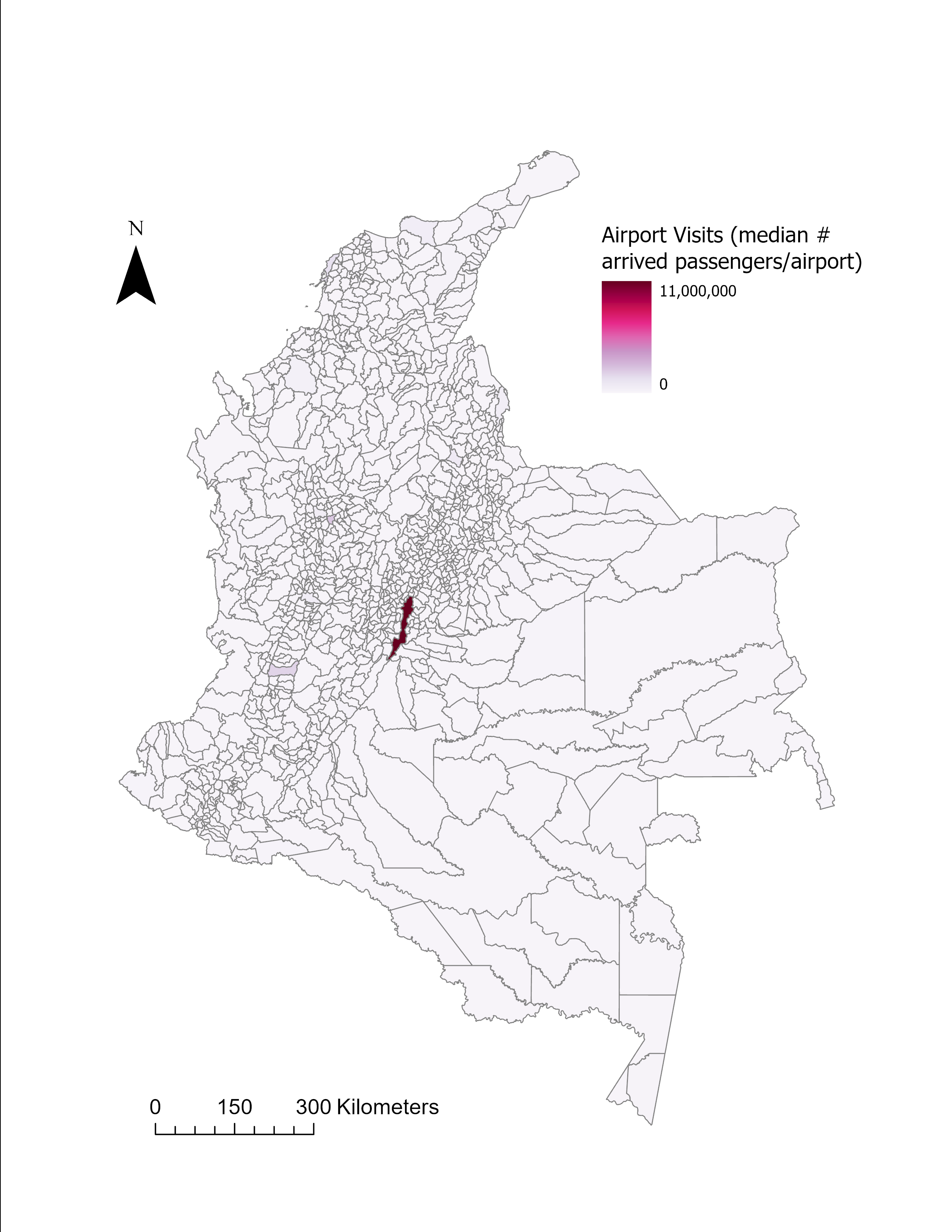

**Figure S48. Spatial distribution of airport visits.**

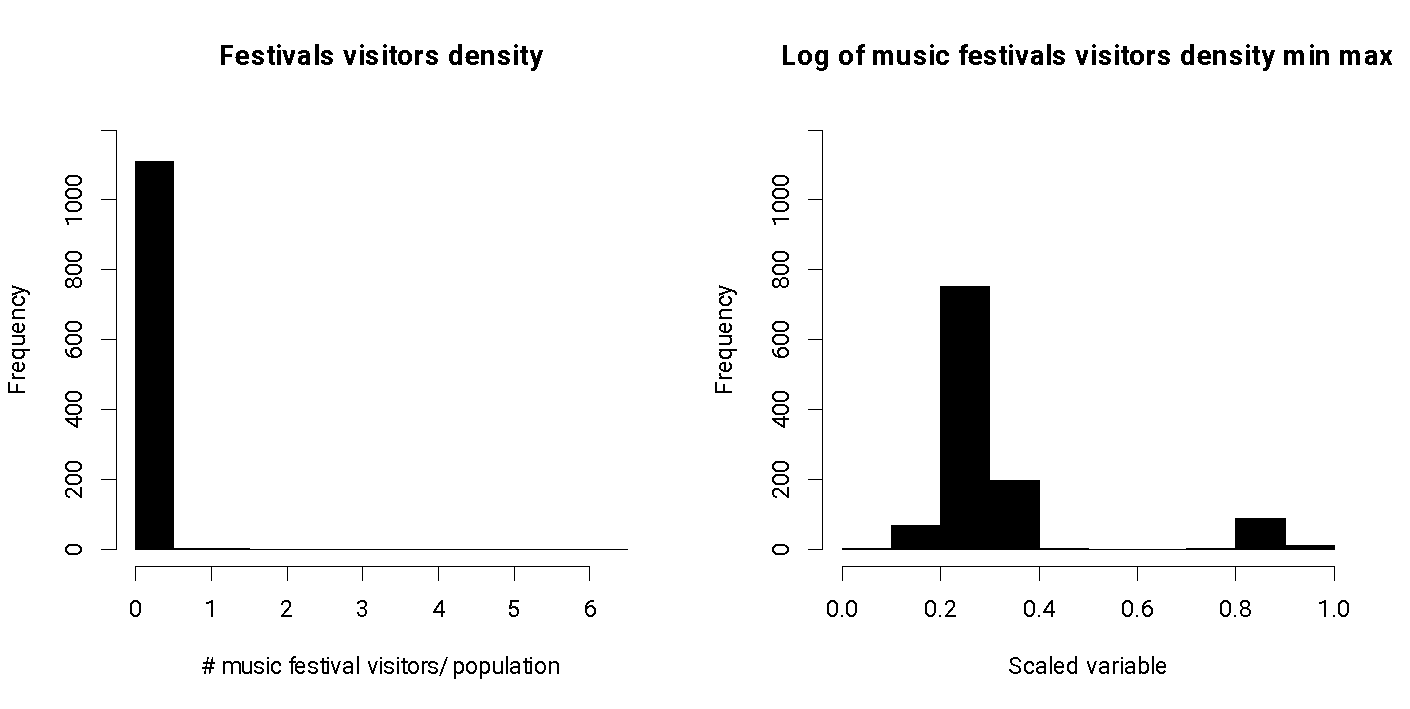

**Figure S49.** **Histograms of visitors to music festivals.**

Left panel is the untransformed variable distribution, and the right panel is the transformed variable distribution.

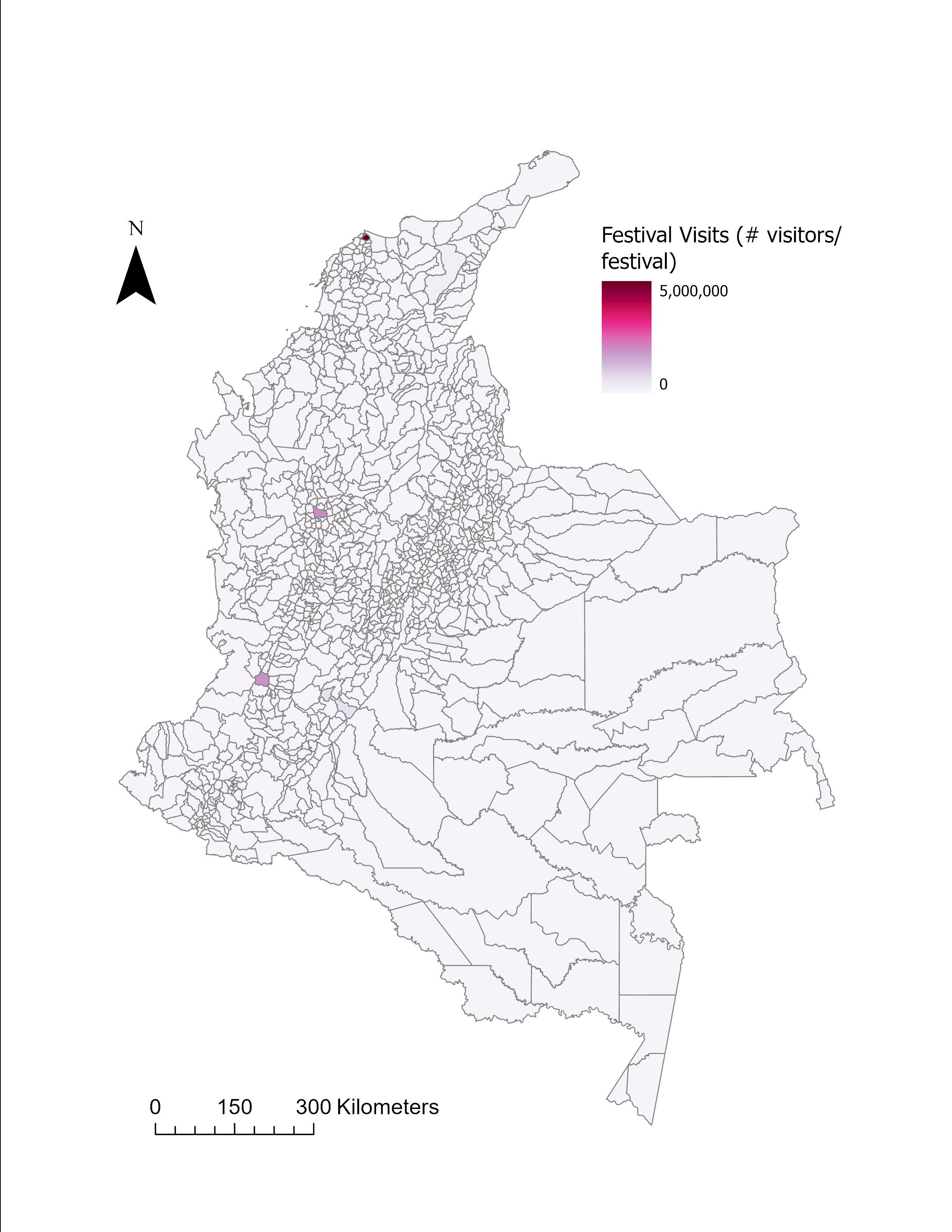

**Figure S50. Spatial distribution of music festival visitation rate**

### 4. Species and ecosystems list

**Table S3. List of bird species, fish species, and ecosystems included in the biodiversity richness index**

| **Group** | **Species or ecosystem type** |
| --- | --- |
| Birds | *Aburria aburri* |
| Birds | *Accipiter bicolor* |
| Birds | *Accipiter striatus* |
| Birds | *Acropternis orthonyx* |
| Birds | *Adelomyia melanogenys* |
| Birds | *Aegolius harrisii* |
| Birds | *Aeronautes montivagus* |
| Birds | *Agamia agami* |
| Birds | *Aglaeactis cupripennis* |
| Birds | *Aglaiocercus coelestis* |
| Birds | *Aglaiocercus kingii* |
| Birds | *Agriornis montanus* |
| Birds | *Akletos melanoceps* |
| Birds | *Amaurolimnas concolor* |
| Birds | *Amaurospiza concolor* |
| Birds | *Amazilia tzacatl* |
| Birds | *Amazona amazonica* |
| Birds | *Amazona autumnalis* |
| Birds | *Amazona farinosa* |
| Birds | *Amazona festiva* |
| Birds | *Amazona kawalli* |
| Birds | *Amazona mercenarius* |
| Birds | *Amazona ochrocephala* |
| Birds | *Amazonetta brasiliensis* |
| Birds | *Amblycercus holosericeus* |
| Birds | *Ammodramus aurifrons* |
| Birds | *Ammodramus humeralis* |
| Birds | *Ammodramus savannarum* |
| Birds | *Ammonastes pelzelni* |
| Birds | *Ampelioides tschudii* |
| Birds | *Ampelion rubrocristatus* |
| Birds | *Ampelion rufaxilla* |
| Birds | *Anabacerthia ruficaudata* |
| Birds | *Anabacerthia striaticollis* |
| Birds | *Anabacerthia variegaticeps* |
| Birds | *Anabazenops dorsalis* |
| Birds | *Anairetes parulus* |
| Birds | *Anas andium* |
| Birds | *Anas bahamensis* |
| Birds | *Anas georgica* |
| Birds | *Ancistrops strigilatus* |
| Birds | *Andigena hypoglauca* |
| Birds | *Andigena laminirostris* |
| Birds | *Andigena nigrirostris* |
| Birds | *Androdon aequatorialis* |
| Birds | *Anhima cornuta* |
| Birds | *Anhinga anhinga* |
| Birds | *Anisognathus igniventris* |
| Birds | *Anisognathus lacrymosus* |
| Birds | *Anisognathus melanogenys* |
| Birds | *Anisognathus notabilis* |
| Birds | *Anisognathus somptuosus* |
| Birds | *Anous minutus* |
| Birds | *Anous stolidus* |
| Birds | *Anthocephala berlepschi* |
| Birds | *Anthocephala floriceps* |
| Birds | *Anthracothorax nigricollis* |
| Birds | *Anthracothorax prevostii* |
| Birds | *Anthus bogotensis* |
| Birds | *Anthus chii* |
| Birds | *Antrostomus rufus* |
| Birds | *Anurolimnas castaneiceps* |
| Birds | *Anurolimnas fasciatus* |
| Birds | *Anurolimnas viridis* |
| Birds | *Aphanotriccus audax* |
| Birds | *Aprositornis disjuncta* |
| Birds | *Ara ambiguus* |
| Birds | *Ara ararauna* |
| Birds | *Ara chloropterus* |
| Birds | *Ara macao* |
| Birds | *Ara militaris* |
| Birds | *Ara severus* |
| Birds | *Aramides axillaris* |
| Birds | *Aramides cajaneus* |
| Birds | *Aramides wolfi* |
| Birds | *Aramus guarauna* |
| Birds | *Aratinga weddellii* |
| Birds | *Ardea alba* |
| Birds | *Ardea cocoi* |
| Birds | *Arremon assimilis* |
| Birds | *Arremon atricapillus* |
| Birds | *Arremon aurantiirostris* |
| Birds | *Arremon basilicus* |
| Birds | *Arremon brunneinucha* |
| Birds | *Arremon castaneiceps* |
| Birds | *Arremon crassirostris* |
| Birds | *Arremon perijanus* |
| Birds | *Arremon schlegeli* |
| Birds | *Arremon taciturnus* |
| Birds | *Arremonops conirostris* |
| Birds | *Arremonops tocuyensis* |
| Birds | *Arundinicola leucocephala* |
| Birds | *Asemospiza fuliginosa* |
| Birds | *Asemospiza obscura* |
| Birds | *Asio clamator* |
| Birds | *Asio flammeus* |
| Birds | *Asio stygius* |
| Birds | *Asthenes flammulata* |
| Birds | *Asthenes fuliginosa* |
| Birds | *Asthenes perijana* |
| Birds | *Asthenes wyatti* |
| Birds | *Atalotriccus pilaris* |
| Birds | *Athene cunicularia* |
| Birds | *Atlapetes albinucha* |
| Birds | *Atlapetes albofrenatus* |
| Birds | *Atlapetes blancae* |
| Birds | *Atlapetes flaviceps* |
| Birds | *Atlapetes fuscoolivaceus* |
| Birds | *Atlapetes latinuchus* |
| Birds | *Atlapetes leucopis* |
| Birds | *Atlapetes melanocephalus* |
| Birds | *Atlapetes pallidinucha* |
| Birds | *Atlapetes schistaceus* |
| Birds | *Atlapetes semirufus* |
| Birds | *Atlapetes tricolor* |
| Birds | *Attagis gayi* |
| Birds | *Atticora fasciata* |
| Birds | *Atticora tibialis* |
| Birds | *Attila bolivianus* |
| Birds | *Attila cinnamomeus* |
| Birds | *Attila citriniventris* |
| Birds | *Attila spadiceus* |
| Birds | *Attila torridus* |
| Birds | *Aulacorhynchus albivitta* |
| Birds | *Aulacorhynchus derbianus* |
| Birds | *Aulacorhynchus haematopygus* |
| Birds | *Aulacorhynchus sulcatus* |
| Birds | *Automolus infuscatus* |
| Birds | *Automolus melanopezus* |
| Birds | *Automolus ochrolaemus* |
| Birds | *Automolus rufipileatus* |
| Birds | *Automolus subulatus* |
| Birds | *Bangsia arcaei* |
| Birds | *Bangsia aureocincta* |
| Birds | *Bangsia edwardsi* |
| Birds | *Bangsia flavovirens* |
| Birds | *Bangsia melanochlamys* |
| Birds | *Bangsia rothschildi* |
| Birds | *Baryphthengus martii* |
| Birds | *Basileuterus culicivorus* |
| Birds | *Basileuterus ignotus* |
| Birds | *Basileuterus rufifrons* |
| Birds | *Basileuterus tristriatus* |
| Birds | *Berlepschia rikeri* |
| Birds | *Boissonneaua flavescens* |
| Birds | *Boissonneaua jardini* |
| Birds | *Boissonneaua matthewsii* |
| Birds | *Bolborhynchus ferrugineifrons* |
| Birds | *Bolborhynchus lineola* |
| Birds | *Botaurus pinnatus* |
| Birds | *Brachygalba goeringi* |
| Birds | *Brachygalba lugubris* |
| Birds | *Brachygalba salmoni* |
| Birds | *Brotogeris cyanoptera* |
| Birds | *Brotogeris jugularis* |
| Birds | *Brotogeris sanctithomae* |
| Birds | *Brotogeris versicolurus* |
| Birds | *Bubo virginianus* |
| Birds | *Bubulcus ibis* |
| Birds | *Bucco capensis* |
| Birds | *Bucco macrodactylus* |
| Birds | *Bucco noanamae* |
| Birds | *Bucco tamatia* |
| Birds | *Burhinus bistriatus* |
| Birds | *Busarellus nigricollis* |
| Birds | *Buteo albigula* |
| Birds | *Buteo albonotatus* |
| Birds | *Buteo brachyurus* |
| Birds | *Buteo nitidus* |
| Birds | *Buteogallus anthracinus* |
| Birds | *Buteogallus meridionalis* |
| Birds | *Buteogallus schistaceus* |
| Birds | *Buteogallus solitarius* |
| Birds | *Buteogallus urubitinga* |
| Birds | *Buthraupis montana* |
| Birds | *Butorides striata* |
| Birds | *Cacicus cela* |
| Birds | *Cacicus chrysonotus* |
| Birds | *Cacicus haemorrhous* |
| Birds | *Cacicus latirostris* |
| Birds | *Cacicus oseryi* |
| Birds | *Cacicus sclateri* |
| Birds | *Cacicus solitarius* |
| Birds | *Cacicus uropygialis* |
| Birds | *Cairina moschata* |
| Birds | *Calliphlox amethystina* |
| Birds | *Calochaetes coccineus* |
| Birds | *Campephilus gayaquilensis* |
| Birds | *Campephilus haematogaster* |
| Birds | *Campephilus melanoleucos* |
| Birds | *Campephilus pollens* |
| Birds | *Campephilus rubricollis* |
| Birds | *Camptostoma obsoletum* |
| Birds | *Campylopterus falcatus* |
| Birds | *Campylopterus largipennis* |
| Birds | *Campylopterus phainopeplus* |
| Birds | *Campylopterus villaviscensio* |
| Birds | *Campylorhamphus procurvoides* |
| Birds | *Campylorhamphus pusillus* |
| Birds | *Campylorhamphus trochilirostris* |
| Birds | *Campylorhynchus albobrunneus* |
| Birds | *Campylorhynchus griseus* |
| Birds | *Campylorhynchus nuchalis* |
| Birds | *Campylorhynchus turdinus* |
| Birds | *Campylorhynchus zonatus* |
| Birds | *Cantorchilus leucopogon* |
| Birds | *Cantorchilus leucotis* |
| Birds | *Cantorchilus nigricapillus* |
| Birds | *Capito auratus* |
| Birds | *Capito aurovirens* |
| Birds | *Capito hypoleucus* |
| Birds | *Capito maculicoronatus* |
| Birds | *Capito quinticolor* |
| Birds | *Capito squamatus* |
| Birds | *Capsiempis flaveola* |
| Birds | *Caracara plancus* |
| Birds | *Cardinalis phoeniceus* |
| Birds | *Carpodectes hopkei* |
| Birds | *Caryothraustes canadensis* |
| Birds | *Catamblyrhynchus diadema* |
| Birds | *Catamenia analis* |
| Birds | *Catamenia homochroa* |
| Birds | *Catamenia inornata* |
| Birds | *Cathartes aura* |
| Birds | *Cathartes burrovianus* |
| Birds | *Cathartes melambrotus* |
| Birds | *Catharus aurantiirostris* |
| Birds | *Catharus fuscater* |
| Birds | *Catharus maculatus* |
| Birds | *Celeus elegans* |
| Birds | *Celeus flavus* |
| Birds | *Celeus grammicus* |
| Birds | *Celeus loricatus* |
| Birds | *Celeus spectabilis* |
| Birds | *Celeus torquatus* |
| Birds | *Cephalopterus ornatus* |
| Birds | *Cephalopterus penduliger* |
| Birds | *Ceratopipra erythrocephala* |
| Birds | *Ceratopipra mentalis* |
| Birds | *Cercibis oxycerca* |
| Birds | *Cercomacra cinerascens* |
| Birds | *Cercomacra nigricans* |
| Birds | *Cercomacroides fuscicauda* |
| Birds | *Cercomacroides nigrescens* |
| Birds | *Cercomacroides parkeri* |
| Birds | *Cercomacroides serva* |
| Birds | *Cercomacroides tyrannina* |
| Birds | *Certhiasomus stictolaemus* |
| Birds | *Certhiaxis cinnamomeus* |
| Birds | *Certhiaxis mustelinus* |
| Birds | *Chaetocercus astreans* |
| Birds | *Chaetocercus heliodor* |
| Birds | *Chaetocercus jourdanii* |
| Birds | *Chaetocercus mulsant* |
| Birds | *Chaetura brachyura* |
| Birds | *Chaetura chapmani* |
| Birds | *Chaetura cinereiventris* |
| Birds | *Chaetura spinicaudus* |
| Birds | *Chalcostigma herrani* |
| Birds | *Chalcostigma heteropogon* |
| Birds | *Chalcostigma stanleyi* |
| Birds | *Chalcothraupis ruficervix* |
| Birds | *Chalybura buffonii* |
| Birds | *Chalybura urochrysia* |
| Birds | *Chamaepetes goudotii* |
| Birds | *Chamaeza campanisona* |
| Birds | *Chamaeza mollissima* |
| Birds | *Chamaeza nobilis* |
| Birds | *Chamaeza turdina* |
| Birds | *Charadrius collaris* |
| Birds | *Charadrius nivosus* |
| Birds | *Charadrius vociferus* |
| Birds | *Charadrius wilsonia* |
| Birds | *Chauna chavaria* |
| Birds | *Chelidoptera tenebrosa* |
| Birds | *Chionomesa fimbriata* |
| Birds | *Chiroxiphia lanceolata* |
| Birds | *Chiroxiphia pareola* |
| Birds | *Chlorestes cyanus* |
| Birds | *Chlorestes eliciae* |
| Birds | *Chlorestes julie* |
| Birds | *Chlorestes notata* |
| Birds | *Chloroceryle aenea* |
| Birds | *Chloroceryle amazona* |
| Birds | *Chloroceryle americana* |
| Birds | *Chloroceryle inda* |
| Birds | *Chlorochrysa calliparaea* |
| Birds | *Chlorochrysa nitidissima* |
| Birds | *Chlorochrysa phoenicotis* |
| Birds | *Chlorophanes spiza* |
| Birds | *Chlorophonia cyanea* |
| Birds | *Chlorophonia cyanocephala* |
| Birds | *Chlorophonia flavirostris* |
| Birds | *Chlorophonia pyrrhophrys* |
| Birds | *Chloropipo flavicapilla* |
| Birds | *Chlorornis riefferii* |
| Birds | *Chlorospingus canigularis* |
| Birds | *Chlorospingus flavigularis* |
| Birds | *Chlorospingus flavopectus* |
| Birds | *Chlorospingus inornatus* |
| Birds | *Chlorospingus parvirostris* |
| Birds | *Chlorospingus semifuscus* |
| Birds | *Chlorospingus tacarcunae* |
| Birds | *Chlorostilbon gibsoni* |
| Birds | *Chlorostilbon melanorhynchus* |
| Birds | *Chlorostilbon mellisugus* |
| Birds | *Chlorostilbon olivaresi* |
| Birds | *Chlorostilbon poortmani* |
| Birds | *Chlorostilbon russatus* |
| Birds | *Chlorostilbon stenurus* |
| Birds | *Chlorothraupis carmioli* |
| Birds | *Chlorothraupis olivacea* |
| Birds | *Chlorothraupis stolzmanni* |
| Birds | *Chondrohierax uncinatus* |
| Birds | *Chordeiles acutipennis* |
| Birds | *Chordeiles nacunda* |
| Birds | *Chordeiles pusillus* |
| Birds | *Chordeiles rupestris* |
| Birds | *Chroicocephalus serranus* |
| Birds | *Chrysolampis mosquitus* |
| Birds | *Chrysomus icterocephalus* |
| Birds | *Chrysothlypis chrysomelas* |
| Birds | *Chrysothlypis salmoni* |
| Birds | *Chrysuronia coeruleogularis* |
| Birds | *Chrysuronia goudoti* |
| Birds | *Chrysuronia grayi* |
| Birds | *Chrysuronia humboldtii* |
| Birds | *Chrysuronia lilliae* |
| Birds | *Chrysuronia oenone* |
| Birds | *Chrysuronia versicolor* |
| Birds | *Cichlopsis leucogenys* |
| Birds | *Ciconia maguari* |
| Birds | *Cinclodes albidiventris* |
| Birds | *Cinclodes excelsior* |
| Birds | *Cinclus leucocephalus* |
| Birds | *Cinnycerthia olivascens* |
| Birds | *Cinnycerthia unirufa* |
| Birds | *Circus buffoni* |
| Birds | *Circus cinereus* |
| Birds | *Cissopis leverianus* |
| Birds | *Cistothorus apolinari* |
| Birds | *Cistothorus platensis* |
| Birds | *Claravis pretiosa* |
| Birds | *Clibanornis rubiginosus* |
| Birds | *Clibanornis rufipectus* |
| Birds | *Clytoctantes alixii* |
| Birds | *Cnemarchus erythropygius* |
| Birds | *Cnemathraupis eximia* |
| Birds | *Cnemoscopus rubrirostris* |
| Birds | *Cnemotriccus fuscatus* |
| Birds | *Cnipodectes subbrunneus* |
| Birds | *Coccycua minuta* |
| Birds | *Coccycua pumila* |
| Birds | *Coccyzus lansbergi* |
| Birds | *Coccyzus melacoryphus* |
| Birds | *Cochlearius cochlearius* |
| Birds | *Coeligena bonapartei* |
| Birds | *Coeligena coeligena* |
| Birds | *Coeligena helianthea* |
| Birds | *Coeligena lutetiae* |
| Birds | *Coeligena orina* |
| Birds | *Coeligena phalerata* |
| Birds | *Coeligena prunellei* |
| Birds | *Coeligena torquata* |
| Birds | *Coeligena wilsoni* |
| Birds | *Coereba flaveola* |
| Birds | *Colaptes punctigula* |
| Birds | *Colaptes rivolii* |
| Birds | *Colaptes rubiginosus* |
| Birds | *Colibri coruscans* |
| Birds | *Colibri cyanotus* |
| Birds | *Colibri delphinae* |
| Birds | *Colinus cristatus* |
| Birds | *Colonia colonus* |
| Birds | *Columbina buckleyi* |
| Birds | *Columbina cruziana* |
| Birds | *Columbina minuta* |
| Birds | *Columbina passerina* |
| Birds | *Columbina squammata* |
| Birds | *Columbina talpacoti* |
| Birds | *Conirostrum albifrons* |
| Birds | *Conirostrum bicolor* |
| Birds | *Conirostrum binghami* |
| Birds | *Conirostrum cinereum* |
| Birds | *Conirostrum leucogenys* |
| Birds | *Conirostrum margaritae* |
| Birds | *Conirostrum rufum* |
| Birds | *Conirostrum sitticolor* |
| Birds | *Conirostrum speciosum* |
| Birds | *Conopias albovittatus* |
| Birds | *Conopias cinchoneti* |
| Birds | *Conopias parvus* |
| Birds | *Conopophaga aurita* |
| Birds | *Conopophaga castaneiceps* |
| Birds | *Contopus cinereus* |
| Birds | *Contopus fumigatus* |
| Birds | *Coragyps atratus* |
| Birds | *Corapipo altera* |
| Birds | *Corapipo leucorrhoa* |
| Birds | *Coryphospingus cucullatus* |
| Birds | *Coryphospingus pileatus* |
| Birds | *Corythopis torquatus* |
| Birds | *Cotinga cayana* |
| Birds | *Cotinga cotinga* |
| Birds | *Cotinga maynana* |
| Birds | *Cotinga nattererii* |
| Birds | *Cranioleuca curtata* |
| Birds | *Cranioleuca erythrops* |
| Birds | *Cranioleuca gutturata* |
| Birds | *Cranioleuca hellmayri* |
| Birds | *Cranioleuca subcristata* |
| Birds | *Cranioleuca vulpecula* |
| Birds | *Cranioleuca vulpina* |
| Birds | *Crax alberti* |
| Birds | *Crax alector* |
| Birds | *Crax daubentoni* |
| Birds | *Crax globulosa* |
| Birds | *Crax rubra* |
| Birds | *Creagrus furcatus* |
| Birds | *Creurgops verticalis* |
| Birds | *Crotophaga ani* |
| Birds | *Crotophaga major* |
| Birds | *Crotophaga sulcirostris* |
| Birds | *Cryptoleucopteryx plumbea* |
| Birds | *Cryptopipo holochlora* |
| Birds | *Crypturellus berlepschi* |
| Birds | *Crypturellus casiquiare* |
| Birds | *Crypturellus cinereus* |
| Birds | *Crypturellus duidae* |
| Birds | *Crypturellus erythropus* |
| Birds | *Crypturellus kerriae* |
| Birds | *Crypturellus obsoletus* |
| Birds | *Crypturellus soui* |
| Birds | *Crypturellus undulatus* |
| Birds | *Crypturellus variegatus* |
| Birds | *Cyanerpes caeruleus* |
| Birds | *Cyanerpes cyaneus* |
| Birds | *Cyanerpes lucidus* |
| Birds | *Cyanerpes nitidus* |
| Birds | *Cyanocorax affinis* |
| Birds | *Cyanocorax heilprini* |
| Birds | *Cyanocorax violaceus* |
| Birds | *Cyanocorax yncas* |
| Birds | *Cyanoloxia brissonii* |
| Birds | *Cyanoloxia cyanoides* |
| Birds | *Cyanoloxia rothschildii* |
| Birds | *Cyanolyca armillata* |
| Birds | *Cyanolyca pulchra* |
| Birds | *Cyanolyca turcosa* |
| Birds | *Cyclarhis gujanensis* |
| Birds | *Cyclarhis nigrirostris* |
| Birds | *Cymbilaimus lineatus* |
| Birds | *Cyphorhinus arada* |
| Birds | *Cyphorhinus phaeocephalus* |
| Birds | *Cyphorhinus thoracicus* |
| Birds | *Cypseloides cherriei* |
| Birds | *Cypseloides cryptus* |
| Birds | *Cypseloides lemosi* |
| Birds | *Dacnis albiventris* |
| Birds | *Dacnis berlepschi* |
| Birds | *Dacnis cayana* |
| Birds | *Dacnis flaviventer* |
| Birds | *Dacnis hartlaubi* |
| Birds | *Dacnis lineata* |
| Birds | *Dacnis venusta* |
| Birds | *Dacnis viguieri* |
| Birds | *Daptrius ater* |
| Birds | *Deconychura longicauda* |
| Birds | *Dendrexetastes rufigula* |
| Birds | *Dendrocincla fuliginosa* |
| Birds | *Dendrocincla homochroa* |
| Birds | *Dendrocincla merula* |
| Birds | *Dendrocincla tyrannina* |
| Birds | *Dendrocolaptes certhia* |
| Birds | *Dendrocolaptes picumnus* |
| Birds | *Dendrocolaptes sanctithomae* |
| Birds | *Dendrocygna autumnalis* |
| Birds | *Dendrocygna bicolor* |
| Birds | *Dendrocygna viduata* |
| Birds | *Dendroma erythroptera* |
| Birds | *Dendroma rufa* |
| Birds | *Dendroplex kienerii* |
| Birds | *Dendroplex picus* |
| Birds | *Deroptyus accipitrinus* |
| Birds | *Dichrozona cincta* |
| Birds | *Diglossa albilatera* |
| Birds | *Diglossa brunneiventris* |
| Birds | *Diglossa caerulescens* |
| Birds | *Diglossa cyanea* |
| Birds | *Diglossa glauca* |
| Birds | *Diglossa gloriosissima* |
| Birds | *Diglossa humeralis* |
| Birds | *Diglossa indigotica* |
| Birds | *Diglossa lafresnayii* |
| Birds | *Diglossa sittoides* |
| Birds | *Discosura conversii* |
| Birds | *Discosura langsdorffi* |
| Birds | *Discosura longicaudus* |
| Birds | *Discosura popelairii* |
| Birds | *Doliornis remseni* |
| Birds | *Donacobius atricapilla* |
| Birds | *Doryfera johannae* |
| Birds | *Doryfera ludovicae* |
| Birds | *Dromococcyx pavoninus* |
| Birds | *Dromococcyx phasianellus* |
| Birds | *Drymophila caudata* |
| Birds | *Drymophila devillei* |
| Birds | *Drymophila hellmayri* |
| Birds | *Drymophila klagesi* |
| Birds | *Drymophila striaticeps* |
| Birds | *Drymotoxeres pucheranii* |
| Birds | *Dryobates affinis* |
| Birds | *Dryobates callonotus* |
| Birds | *Dryobates chocoensis* |
| Birds | *Dryobates dignus* |
| Birds | *Dryobates fumigatus* |
| Birds | *Dryobates kirkii* |
| Birds | *Dryobates nigriceps* |
| Birds | *Dryobates passerinus* |
| Birds | *Dryocopus lineatus* |
| Birds | *Dubusia taeniata* |
| Birds | *Dysithamnus leucostictus* |
| Birds | *Dysithamnus mentalis* |
| Birds | *Dysithamnus occidentalis* |
| Birds | *Dysithamnus puncticeps* |
| Birds | *Egretta caerulea* |
| Birds | *Egretta rufescens* |
| Birds | *Egretta thula* |
| Birds | *Egretta tricolor* |
| Birds | *Elaenia albiceps* |
| Birds | *Elaenia brachyptera* |
| Birds | *Elaenia chiriquensis* |
| Birds | *Elaenia cristata* |
| Birds | *Elaenia flavogaster* |
| Birds | *Elaenia frantzii* |
| Birds | *Elaenia gigas* |
| Birds | *Elaenia martinica* |
| Birds | *Elaenia pallatangae* |
| Birds | *Elaenia ruficeps* |
| Birds | *Elanoides forficatus* |
| Birds | *Elanus leucurus* |
| Birds | *Electron platyrhynchum* |
| Birds | *Emberizoides herbicola* |
| Birds | *Ensifera ensifera* |
| Birds | *Entomodestes coracinus* |
| Birds | *Epinecrophylla erythrura* |
| Birds | *Epinecrophylla fulviventris* |
| Birds | *Epinecrophylla haematonota* |
| Birds | *Epinecrophylla ornata* |
| Birds | *Epinecrophylla spodionota* |
| Birds | *Eremophila alpestris* |
| Birds | *Eriocnemis aline* |
| Birds | *Eriocnemis cupreoventris* |
| Birds | *Eriocnemis derbyi* |
| Birds | *Eriocnemis godini* |
| Birds | *Eriocnemis isabellae* |
| Birds | *Eriocnemis luciani* |
| Birds | *Eriocnemis mirabilis* |
| Birds | *Eriocnemis mosquera* |
| Birds | *Eriocnemis vestita* |
| Birds | *Eubucco bourcierii* |
| Birds | *Eubucco richardsoni* |
| Birds | *Euchrepomis callinota* |
| Birds | *Euchrepomis spodioptila* |
| Birds | *Eucometis penicillata* |
| Birds | *Eudocimus albus* |
| Birds | *Eudocimus ruber* |
| Birds | *Euphonia anneae* |
| Birds | *Euphonia chlorotica* |
| Birds | *Euphonia chrysopasta* |
| Birds | *Euphonia concinna* |
| Birds | *Euphonia fulvicrissa* |
| Birds | *Euphonia laniirostris* |
| Birds | *Euphonia mesochrysa* |
| Birds | *Euphonia minuta* |
| Birds | *Euphonia plumbea* |
| Birds | *Euphonia rufiventris* |
| Birds | *Euphonia saturata* |
| Birds | *Euphonia trinitatis* |
| Birds | *Euphonia xanthogaster* |
| Birds | *Eupsittula pertinax* |
| Birds | *Eurypyga helias* |
| Birds | *Euscarthmus meloryphus* |
| Birds | *Eutoxeres aquila* |
| Birds | *Eutoxeres condamini* |
| Birds | *Falco deiroleucus* |
| Birds | *Falco femoralis* |
| Birds | *Falco rufigularis* |
| Birds | *Falco sparverius* |
| Birds | *Florisuga mellivora* |
| Birds | *Fluvicola nengeta* |
| Birds | *Fluvicola pica* |
| Birds | *Formicarius analis* |
| Birds | *Formicarius colma* |
| Birds | *Formicarius nigricapillus* |
| Birds | *Formicarius rufipectus* |
| Birds | *Formicivora grisea* |
| Birds | *Forpus coelestis* |
| Birds | *Forpus conspicillatus* |
| Birds | *Forpus crassirostris* |
| Birds | *Forpus modestus* |
| Birds | *Forpus passerinus* |
| Birds | *Forpus spengeli* |
| Birds | *Frederickena fulva* |
| Birds | *Fregata magnificens* |
| Birds | *Fregata minor* |
| Birds | *Fulica americana* |
| Birds | *Fulica ardesiaca* |
| Birds | *Furnarius leucopus* |
| Birds | *Furnarius minor* |
| Birds | *Furnarius torridus* |
| Birds | *Galbalcyrhynchus leucotis* |
| Birds | *Galbula albirostris* |
| Birds | *Galbula chalcothorax* |
| Birds | *Galbula dea* |
| Birds | *Galbula galbula* |
| Birds | *Galbula leucogastra* |
| Birds | *Galbula pastazae* |
| Birds | *Galbula ruficauda* |
| Birds | *Galbula tombacea* |
| Birds | *Gallinago imperialis* |
| Birds | *Gallinago jamesoni* |
| Birds | *Gallinago nobilis* |
| Birds | *Gallinago paraguaiae* |
| Birds | *Gallinago undulata* |
| Birds | *Gallinula galeata* |
| Birds | *Gampsonyx swainsonii* |
| Birds | *Gelochelidon nilotica* |
| Birds | *Geospizopsis unicolor* |
| Birds | *Geothlypis aequinoctialis* |
| Birds | *Geothlypis semiflava* |
| Birds | *Geotrygon montana* |
| Birds | *Geotrygon purpurata* |
| Birds | *Geotrygon saphirina* |
| Birds | *Geotrygon violacea* |
| Birds | *Geranoaetus albicaudatus* |
| Birds | *Geranoaetus melanoleucus* |
| Birds | *Geranospiza caerulescens* |
| Birds | *Glaucidium brasilianum* |
| Birds | *Glaucidium griseiceps* |
| Birds | *Glaucidium jardinii* |
| Birds | *Glaucidium nubicola* |
| Birds | *Glaucidium parkeri* |
| Birds | *Glaucis aeneus* |
| Birds | *Glaucis hirsutus* |
| Birds | *Glyphorynchus spirurus* |
| Birds | *Goldmania bella* |
| Birds | *Goldmania violiceps* |
| Birds | *Grallaria alleni* |
| Birds | *Grallaria alvarezi* |
| Birds | *Grallaria bangsi* |
| Birds | *Grallaria dignissima* |
| Birds | *Grallaria flavotincta* |
| Birds | *Grallaria gigantea* |
| Birds | *Grallaria guatimalensis* |
| Birds | *Grallaria haplonota* |
| Birds | *Grallaria hypoleuca* |
| Birds | *Grallaria kaestneri* |
| Birds | *Grallaria milleri* |
| Birds | *Grallaria nuchalis* |
| Birds | *Grallaria quitensis* |
| Birds | *Grallaria ruficapilla* |
| Birds | *Grallaria rufocinerea* |
| Birds | *Grallaria rufula* |
| Birds | *Grallaria saltuensis* |
| Birds | *Grallaria saturata* |
| Birds | *Grallaria spatiator* |
| Birds | *Grallaria squamigera* |
| Birds | *Grallaria urraoensis* |
| Birds | *Grallaria varia* |
| Birds | *Grallaricula cucullata* |
| Birds | *Grallaricula ferrugineipectus* |
| Birds | *Grallaricula flavirostris* |
| Birds | *Grallaricula lineifrons* |
| Birds | *Grallaricula nana* |
| Birds | *Granatellus pelzelni* |
| Birds | *Graydidascalus brachyurus* |
| Birds | *Gygis alba* |
| Birds | *Gymnocichla nudiceps* |
| Birds | *Gymnoderus foetidus* |
| Birds | *Gymnomystax mexicanus* |
| Birds | *Gymnopithys bicolor* |
| Birds | *Gymnopithys leucaspis* |
| Birds | *Habia cristata* |
| Birds | *Habia fuscicauda* |
| Birds | *Habia gutturalis* |
| Birds | *Habia rubica* |
| Birds | *Haematopus palliatus* |
| Birds | *Hafferia fortis* |
| Birds | *Hafferia immaculata* |
| Birds | *Hafferia zeledoni* |
| Birds | *Hapalopsittaca amazonina* |
| Birds | *Hapalopsittaca fuertesi* |
| Birds | *Hapaloptila castanea* |
| Birds | *Haplophaedia aureliae* |
| Birds | *Haplophaedia lugens* |
| Birds | *Haplospiza rustica* |
| Birds | *Harpagus bidentatus* |
| Birds | *Harpia harpyja* |
| Birds | *Heliangelus amethysticollis* |
| Birds | *Heliangelus exortis* |
| Birds | *Heliangelus mavors* |
| Birds | *Heliangelus strophianus* |
| Birds | *Helicolestes hamatus* |
| Birds | *Heliodoxa aurescens* |
| Birds | *Heliodoxa gularis* |
| Birds | *Heliodoxa imperatrix* |
| Birds | *Heliodoxa jacula* |
| Birds | *Heliodoxa leadbeateri* |
| Birds | *Heliodoxa rubinoides* |
| Birds | *Heliodoxa schreibersii* |
| Birds | *Heliomaster furcifer* |
| Birds | *Heliomaster longirostris* |
| Birds | *Heliornis fulica* |
| Birds | *Heliothryx auritus* |
| Birds | *Heliothryx barroti* |
| Birds | *Hellmayrea gularis* |
| Birds | *Hemithraupis flavicollis* |
| Birds | *Hemithraupis guira* |
| Birds | *Hemitriccus granadensis* |
| Birds | *Hemitriccus iohannis* |
| Birds | *Hemitriccus margaritaceiventer* |
| Birds | *Hemitriccus rufigularis* |
| Birds | *Hemitriccus striaticollis* |
| Birds | *Hemitriccus zosterops* |
| Birds | *Henicorhina anachoreta* |
| Birds | *Henicorhina leucophrys* |
| Birds | *Henicorhina leucosticta* |
| Birds | *Henicorhina negreti* |
| Birds | *Herpetotheres cachinnans* |
| Birds | *Herpsilochmus axillaris* |
| Birds | *Herpsilochmus dorsimaculatus* |
| Birds | *Herpsilochmus dugandi* |
| Birds | *Herpsilochmus frater* |
| Birds | *Heterocercus flavivertex* |
| Birds | *Heterospingus xanthopygius* |
| Birds | *Himantopus mexicanus* |
| Birds | *Hirundinea ferruginea* |
| Birds | *Hydropsalis cayennensis* |
| Birds | *Hydropsalis climacocerca* |
| Birds | *Hydropsalis maculicaudus* |
| Birds | *Hylexetastes stresemanni* |
| Birds | *Hylocharis sapphirina* |
| Birds | *Hylomanes momotula* |
| Birds | *Hylopezus macularius* |
| Birds | *Hylopezus perspicillatus* |
| Birds | *Hylophilus brunneiceps* |
| Birds | *Hylophilus flavipes* |
| Birds | *Hylophilus thoracicus* |
| Birds | *Hylophylax naevioides* |
| Birds | *Hylophylax naevius* |
| Birds | *Hylophylax punctulatus* |
| Birds | *Hypnelus ruficollis* |
| Birds | *Hypocnemis flavescens* |
| Birds | *Hypocnemis hypoxantha* |
| Birds | *Hypocnemis peruviana* |
| Birds | *Hypocnemoides maculicauda* |
| Birds | *Hypocnemoides melanopogon* |
| Birds | *Hypopyrrhus pyrohypogaster* |
| Birds | *Ibycter americanus* |
| Birds | *Icterus auricapillus* |
| Birds | *Icterus cayanensis* |
| Birds | *Icterus chrysater* |
| Birds | *Icterus croconotus* |
| Birds | *Icterus icterus* |
| Birds | *Icterus leucopteryx* |
| Birds | *Icterus mesomelas* |
| Birds | *Icterus nigrogularis* |
| Birds | *Ictinia plumbea* |
| Birds | *Inezia caudata* |
| Birds | *Inezia subflava* |
| Birds | *Inezia tenuirostris* |
| Birds | *Iodopleura isabellae* |
| Birds | *Iridophanes pulcherrimus* |
| Birds | *Iridosornis analis* |
| Birds | *Iridosornis porphyrocephalus* |
| Birds | *Iridosornis rufivertex* |
| Birds | *Isleria hauxwelli* |
| Birds | *Ixobrychus exilis* |
| Birds | *Ixobrychus involucris* |
| Birds | *Ixothraupis guttata* |
| Birds | *Ixothraupis punctata* |
| Birds | *Ixothraupis rufigula* |
| Birds | *Ixothraupis varia* |
| Birds | *Ixothraupis xanthogastra* |
| Birds | *Jabiru mycteria* |
| Birds | *Jacamerops aureus* |
| Birds | *Jacana jacana* |
| Birds | *Klais guimeti* |
| Birds | *Kleinothraupis atropileus* |
| Birds | *Knipolegus orenocensis* |
| Birds | *Knipolegus poecilocercus* |
| Birds | *Knipolegus poecilurus* |
| Birds | *Lafresnaya lafresnayi* |
| Birds | *Lampropsar tanagrinus* |
| Birds | *Laniisoma elegans* |
| Birds | *Lanio fulvus* |
| Birds | *Laniocera hypopyrra* |
| Birds | *Laniocera rufescens* |
| Birds | *Laterallus albigularis* |
| Birds | *Laterallus exilis* |
| Birds | *Laterallus melanophaius* |
| Birds | *Lathrotriccus euleri* |
| Birds | *Legatus leucophaius* |
| Birds | *Leistes bellicosus* |
| Birds | *Leistes militaris* |
| Birds | *Lepidocolaptes lacrymiger* |
| Birds | *Lepidocolaptes souleyetii* |
| Birds | *Lepidothrix coronata* |
| Birds | *Lepidothrix isidorei* |
| Birds | *Lepidothrix velutina* |
| Birds | *Leptasthenura andicola* |
| Birds | *Leptodon cayanensis* |
| Birds | *Leptopogon amaurocephalus* |
| Birds | *Leptopogon rufipectus* |
| Birds | *Leptopogon superciliaris* |
| Birds | *Leptosittaca branickii* |
| Birds | *Leptotila cassinii* |
| Birds | *Leptotila conoveri* |
| Birds | *Leptotila jamaicensis* |
| Birds | *Leptotila pallida* |
| Birds | *Leptotila plumbeiceps* |
| Birds | *Leptotila rufaxilla* |
| Birds | *Leptotila verreauxi* |
| Birds | *Leptotrygon veraguensis* |
| Birds | *Lesbia nuna* |
| Birds | *Lesbia victoriae* |
| Birds | *Leucippus fallax* |
| Birds | *Leucopternis melanops* |
| Birds | *Leucopternis semiplumbeus* |
| Birds | *Liosceles thoracicus* |
| Birds | *Lipaugus fuscocinereus* |
| Birds | *Lipaugus unirufus* |
| Birds | *Lipaugus vociferans* |
| Birds | *Lipaugus weberi* |
| Birds | *Lochmias nematura* |
| Birds | *Lophornis chalybeus* |
| Birds | *Lophornis delattrei* |
| Birds | *Lophornis stictolophus* |
| Birds | *Lophostrix cristata* |
| Birds | *Lophotriccus galeatus* |
| Birds | *Lophotriccus pileatus* |
| Birds | *Lophotriccus vitiosus* |
| Birds | *Loriotus cristatus* |
| Birds | *Loriotus luctuosus* |
| Birds | *Lurocalis rufiventris* |
| Birds | *Lurocalis semitorquatus* |
| Birds | *Machaeropterus deliciosus* |
| Birds | *Machaeropterus striolatus* |
| Birds | *Machetornis rixosa* |
| Birds | *Macroagelaius subalaris* |
| Birds | *Malacoptila fulvogularis* |
| Birds | *Malacoptila fusca* |
| Birds | *Malacoptila mystacalis* |
| Birds | *Malacoptila panamensis* |
| Birds | *Manacus manacus* |
| Birds | *Margarornis bellulus* |
| Birds | *Margarornis squamiger* |
| Birds | *Margarornis stellatus* |
| Birds | *Masius chrysopterus* |
| Birds | *Mazaria propinqua* |
| Birds | *Mecocerculus leucophrys* |
| Birds | *Mecocerculus minor* |
| Birds | *Mecocerculus poecilocercus* |
| Birds | *Mecocerculus stictopterus* |
| Birds | *Megaceryle torquata* |
| Birds | *Megarynchus pitangua* |
| Birds | *Megascops albogularis* |
| Birds | *Megascops centralis* |
| Birds | *Megascops choliba* |
| Birds | *Megascops clarkii* |
| Birds | *Megascops gilesi* |
| Birds | *Megascops ingens* |
| Birds | *Megascops petersoni* |
| Birds | *Megascops roraimae* |
| Birds | *Megascops watsonii* |
| Birds | *Megastictus margaritatus* |
| Birds | *Melanerpes cruentatus* |
| Birds | *Melanerpes formicivorus* |
| Birds | *Melanerpes pucherani* |
| Birds | *Melanerpes pulcher* |
| Birds | *Melanerpes rubricapillus* |
| Birds | *Melanospiza bicolor* |
| Birds | *Merganetta armata* |
| Birds | *Mesembrinibis cayennensis* |
| Birds | *Metallura iracunda* |
| Birds | *Metallura tyrianthina* |
| Birds | *Metallura williami* |
| Birds | *Metopothrix aurantiaca* |
| Birds | *Metriopelia melanoptera* |
| Birds | *Micrastur gilvicollis* |
| Birds | *Micrastur mirandollei* |
| Birds | *Micrastur plumbeus* |
| Birds | *Micrastur ruficollis* |
| Birds | *Micrastur semitorquatus* |
| Birds | *Microbates cinereiventris* |
| Birds | *Microbates collaris* |
| Birds | *Microcerculus bambla* |
| Birds | *Microcerculus marginatus* |
| Birds | *Micromonacha lanceolata* |
| Birds | *Micropygia schomburgkii* |
| Birds | *Microrhopias quixensis* |
| Birds | *Microspizias collaris* |
| Birds | *Microspizias superciliosus* |
| Birds | *Microxenops milleri* |
| Birds | *Milvago chimachima* |
| Birds | *Mimus gilvus* |
| Birds | *Mionectes oleagineus* |
| Birds | *Mionectes olivaceus* |
| Birds | *Mionectes striaticollis* |
| Birds | *Mitrephanes phaeocercus* |
| Birds | *Mitrospingus cassinii* |
| Birds | *Mitu salvini* |
| Birds | *Mitu tomentosum* |
| Birds | *Mitu tuberosum* |
| Birds | *Molothrus aeneus* |
| Birds | *Molothrus bonariensis* |
| Birds | *Molothrus oryzivorus* |
| Birds | *Momotus aequatorialis* |
| Birds | *Momotus momota* |
| Birds | *Momotus subrufescens* |
| Birds | *Monasa flavirostris* |
| Birds | *Monasa morphoeus* |
| Birds | *Monasa nigrifrons* |
| Birds | *Morphnarchus princeps* |
| Birds | *Morphnus guianensis* |
| Birds | *Muscisaxicola alpinus* |
| Birds | *Muscisaxicola maculirostris* |
| Birds | *Mustelirallus albicollis* |
| Birds | *Mustelirallus colombianus* |
| Birds | *Mustelirallus erythrops* |
| Birds | *Myadestes coloratus* |
| Birds | *Myadestes ralloides* |
| Birds | *Mycteria americana* |
| Birds | *Myiarchus apicalis* |
| Birds | *Myiarchus cephalotes* |
| Birds | *Myiarchus ferox* |
| Birds | *Myiarchus panamensis* |
| Birds | *Myiarchus tuberculifer* |
| Birds | *Myiarchus tyrannulus* |
| Birds | *Myiarchus venezuelensis* |
| Birds | *Myiobius atricaudus* |
| Birds | *Myiobius barbatus* |
| Birds | *Myiobius villosus* |
| Birds | *Myioborus flavivertex* |
| Birds | *Myioborus melanocephalus* |
| Birds | *Myioborus miniatus* |
| Birds | *Myioborus ornatus* |
| Birds | *Myiodynastes chrysocephalus* |
| Birds | *Myiodynastes maculatus* |
| Birds | *Myiopagis caniceps* |
| Birds | *Myiopagis flavivertex* |
| Birds | *Myiopagis gaimardii* |
| Birds | *Myiopagis olallai* |
| Birds | *Myiopagis viridicata* |
| Birds | *Myiophobus fasciatus* |
| Birds | *Myiophobus flavicans* |
| Birds | *Myiophobus phoenicomitra* |
| Birds | *Myiornis atricapillus* |
| Birds | *Myiornis ecaudatus* |
| Birds | *Myiotheretes fumigatus* |
| Birds | *Myiotheretes pernix* |
| Birds | *Myiotheretes striaticollis* |
| Birds | *Myiothlypis basilica* |
| Birds | *Myiothlypis chrysogaster* |
| Birds | *Myiothlypis cinereicollis* |
| Birds | *Myiothlypis conspicillata* |
| Birds | *Myiothlypis coronata* |
| Birds | *Myiothlypis flaveola* |
| Birds | *Myiothlypis fulvicauda* |
| Birds | *Myiothlypis luteoviridis* |
| Birds | *Myiothlypis nigrocristata* |
| Birds | *Myiotriccus ornatus* |
| Birds | *Myiozetetes cayanensis* |
| Birds | *Myiozetetes granadensis* |
| Birds | *Myiozetetes luteiventris* |
| Birds | *Myiozetetes similis* |
| Birds | *Myornis senilis* |
| Birds | *Myrmeciza longipes* |
| Birds | *Myrmelastes hyperythrus* |
| Birds | *Myrmelastes leucostigma* |
| Birds | *Myrmelastes schistaceus* |
| Birds | *Myrmoborus leucophrys* |
| Birds | *Myrmoborus lugubris* |
| Birds | *Myrmoborus melanurus* |
| Birds | *Myrmoborus myotherinus* |
| Birds | *Myrmochanes hemileucus* |
| Birds | *Myrmophylax atrothorax* |
| Birds | *Myrmornis torquata* |
| Birds | *Myrmothera campanisona* |
| Birds | *Myrmothera dives* |
| Birds | *Myrmothera fulviventris* |
| Birds | *Myrmotherula ambigua* |
| Birds | *Myrmotherula assimilis* |
| Birds | *Myrmotherula axillaris* |
| Birds | *Myrmotherula behni* |
| Birds | *Myrmotherula brachyura* |
| Birds | *Myrmotherula cherriei* |
| Birds | *Myrmotherula ignota* |
| Birds | *Myrmotherula longicauda* |
| Birds | *Myrmotherula longipennis* |
| Birds | *Myrmotherula menetriesii* |
| Birds | *Myrmotherula multostriata* |
| Birds | *Myrmotherula pacifica* |
| Birds | *Myrmotherula schisticolor* |
| Birds | *Myrmotherula sunensis* |
| Birds | *Nasica longirostris* |
| Birds | *Nemosia pileata* |
| Birds | *Neoctantes niger* |
| Birds | *Neomorphus geoffroyi* |
| Birds | *Neomorphus pucheranii* |
| Birds | *Neomorphus radiolosus* |
| Birds | *Neopelma chrysocephalum* |
| Birds | *Neopipo cinnamomea* |
| Birds | *Nephelomyias pulcher* |
| Birds | *Netta erythrophthalma* |
| Birds | *Nomonyx dominicus* |
| Birds | *Nonnula brunnea* |
| Birds | *Nonnula frontalis* |
| Birds | *Nonnula rubecula* |
| Birds | *Nonnula ruficapilla* |
| Birds | *Notharchus hyperrhynchus* |
| Birds | *Notharchus ordii* |
| Birds | *Notharchus pectoralis* |
| Birds | *Notharchus tectus* |
| Birds | *Nothocercus bonapartei* |
| Birds | *Nothocercus julius* |
| Birds | *Nothocrax urumutum* |
| Birds | *Nothoprocta curvirostris* |
| Birds | *Nyctanassa violacea* |
| Birds | *Nyctibius aethereus* |
| Birds | *Nyctibius grandis* |
| Birds | *Nyctibius griseus* |
| Birds | *Nyctibius leucopterus* |
| Birds | *Nyctibius maculosus* |
| Birds | *Nycticorax nycticorax* |
| Birds | *Nyctidromus albicollis* |
| Birds | *Nyctiphrynus ocellatus* |
| Birds | *Nyctiphrynus rosenbergi* |
| Birds | *Nyctipolus nigrescens* |
| Birds | *Nyctiprogne leucopyga* |
| Birds | *Nystalus obamai* |
| Birds | *Nystalus radiatus* |
| Birds | *Ochthoeca cinnamomeiventris* |
| Birds | *Ochthoeca diadema* |
| Birds | *Ochthoeca frontalis* |
| Birds | *Ochthoeca fumicolor* |
| Birds | *Ochthoeca rufipectoralis* |
| Birds | *Ochthornis littoralis* |
| Birds | *Ocreatus underwoodii* |
| Birds | *Odontophorus atrifrons* |
| Birds | *Odontophorus dialeucos* |
| Birds | *Odontophorus erythrops* |
| Birds | *Odontophorus gujanensis* |
| Birds | *Odontophorus hyperythrus* |
| Birds | *Odontophorus melanonotus* |
| Birds | *Odontophorus speciosus* |
| Birds | *Odontophorus strophium* |
| Birds | *Odontorchilus branickii* |
| Birds | *Ognorhynchus icterotis* |
| Birds | *Oncostoma cinereigulare* |
| Birds | *Oncostoma olivaceum* |
| Birds | *Onychoprion anaethetus* |
| Birds | *Onychorhynchus coronatus* |
| Birds | *Opisthocomus hoazin* |
| Birds | *Opisthoprora euryptera* |
| Birds | *Oreothraupis arremonops* |
| Birds | *Oreotrochilus chimborazo* |
| Birds | *Oressochen jubatus* |
| Birds | *Ornithion brunneicapillus* |
| Birds | *Ornithion inerme* |
| Birds | *Orochelidon flavipes* |
| Birds | *Orochelidon murina* |
| Birds | *Ortalis cinereiceps* |
| Birds | *Ortalis columbiana* |
| Birds | *Ortalis erythroptera* |
| Birds | *Ortalis garrula* |
| Birds | *Ortalis guttata* |
| Birds | *Ortalis ruficauda* |
| Birds | *Orthopsittaca manilatus* |
| Birds | *Oxypogon cyanolaemus* |
| Birds | *Oxypogon guerinii* |
| Birds | *Oxypogon stuebelii* |
| Birds | *Oxyruncus cristatus* |
| Birds | *Oxyura jamaicensis* |
| Birds | *Pachyramphus albogriseus* |
| Birds | *Pachyramphus castaneus* |
| Birds | *Pachyramphus cinnamomeus* |
| Birds | *Pachyramphus homochrous* |
| Birds | *Pachyramphus marginatus* |
| Birds | *Pachyramphus minor* |
| Birds | *Pachyramphus polychopterus* |
| Birds | *Pachyramphus rufus* |
| Birds | *Pachyramphus versicolor* |
| Birds | *Pachysylvia aurantiifrons* |
| Birds | *Pachysylvia decurtata* |
| Birds | *Pachysylvia hypoxantha* |
| Birds | *Pachysylvia semibrunnea* |
| Birds | *Panyptila cayennensis* |
| Birds | *Parabuteo leucorrhous* |
| Birds | *Parabuteo unicinctus* |
| Birds | *Paraclaravis mondetoura* |
| Birds | *Pardirallus maculatus* |
| Birds | *Pardirallus nigricans* |
| Birds | *Parkerthraustes humeralis* |
| Birds | *Paroaria gularis* |
| Birds | *Paroaria nigrogenis* |
| Birds | *Patagioenas cayennensis* |
| Birds | *Patagioenas corensis* |
| Birds | *Patagioenas fasciata* |
| Birds | *Patagioenas goodsoni* |
| Birds | *Patagioenas leucocephala* |
| Birds | *Patagioenas nigrirostris* |
| Birds | *Patagioenas plumbea* |
| Birds | *Patagioenas speciosa* |
| Birds | *Patagioenas subvinacea* |
| Birds | *Patagona gigas* |
| Birds | *Pauxi pauxi* |
| Birds | *Pelecanus occidentalis* |
| Birds | *Penelope argyrotis* |
| Birds | *Penelope jacquacu* |
| Birds | *Penelope montagnii* |
| Birds | *Penelope ortoni* |
| Birds | *Penelope perspicax* |
| Birds | *Penelope purpurascens* |
| Birds | *Percnostola rufifrons* |
| Birds | *Perissocephalus tricolor* |
| Birds | *Phacellodomus rufifrons* |
| Birds | *Phaenostictus mcleannani* |
| Birds | *Phaeochroa cuvierii* |
| Birds | *Phaeomyias murina* |
| Birds | *Phaethon aethereus* |
| Birds | *Phaethornis anthophilus* |
| Birds | *Phaethornis atrimentalis* |
| Birds | *Phaethornis augusti* |
| Birds | *Phaethornis bourcieri* |
| Birds | *Phaethornis griseogularis* |
| Birds | *Phaethornis guy* |
| Birds | *Phaethornis hispidus* |
| Birds | *Phaethornis longirostris* |
| Birds | *Phaethornis malaris* |
| Birds | *Phaethornis ruber* |
| Birds | *Phaethornis rupurumii* |
| Birds | *Phaethornis striigularis* |
| Birds | *Phaethornis syrmatophorus* |
| Birds | *Phaethornis yaruqui* |
| Birds | *Phaetusa simplex* |
| Birds | *Phalacrocorax brasilianus* |
| Birds | *Phalcoboenus carunculatus* |
| Birds | *Pharomachrus antisianus* |
| Birds | *Pharomachrus auriceps* |
| Birds | *Pharomachrus fulgidus* |
| Birds | *Pharomachrus pavoninus* |
| Birds | *Phelpsia inornata* |
| Birds | *Pheucticus aureoventris* |
| Birds | *Pheucticus chrysogaster* |
| Birds | *Pheugopedius coraya* |
| Birds | *Pheugopedius euophrys* |
| Birds | *Pheugopedius fasciatoventris* |
| Birds | *Pheugopedius mystacalis* |
| Birds | *Pheugopedius rutilus* |
| Birds | *Pheugopedius sclateri* |
| Birds | *Pheugopedius spadix* |
| Birds | *Philodice mitchellii* |
| Birds | *Philydor erythrocercum* |
| Birds | *Philydor fuscipenne* |
| Birds | *Philydor pyrrhodes* |
| Birds | *Phimosus infuscatus* |
| Birds | *Phlegopsis erythroptera* |
| Birds | *Phlegopsis nigromaculata* |
| Birds | *Phlogophilus hemileucurus* |
| Birds | *Phoenicircus nigricollis* |
| Birds | *Phoenicopterus ruber* |
| Birds | *Phyllaemulor bracteatus* |
| Birds | *Phyllomyias burmeisteri* |
| Birds | *Phyllomyias cinereiceps* |
| Birds | *Phyllomyias griseiceps* |
| Birds | *Phyllomyias nigrocapillus* |
| Birds | *Phyllomyias plumbeiceps* |
| Birds | *Phyllomyias uropygialis* |
| Birds | *Phylloscartes lanyoni* |
| Birds | *Phylloscartes ophthalmicus* |
| Birds | *Phylloscartes orbitalis* |
| Birds | *Phylloscartes poecilotis* |
| Birds | *Phylloscartes superciliaris* |
| Birds | *Piaya cayana* |
| Birds | *Piaya melanogaster* |
| Birds | *Piculus chrysochloros* |
| Birds | *Piculus flavigula* |
| Birds | *Piculus leucolaemus* |
| Birds | *Piculus litae* |
| Birds | *Picumnus castelnau* |
| Birds | *Picumnus cinnamomeus* |
| Birds | *Picumnus exilis* |
| Birds | *Picumnus granadensis* |
| Birds | *Picumnus lafresnayi* |
| Birds | *Picumnus olivaceus* |
| Birds | *Picumnus pumilus* |
| Birds | *Picumnus rufiventris* |
| Birds | *Picumnus spilogaster* |
| Birds | *Picumnus squamulatus* |
| Birds | *Pilherodius pileatus* |
| Birds | *Pionites leucogaster* |
| Birds | *Pionites melanocephalus* |
| Birds | *Pionus chalcopterus* |
| Birds | *Pionus fuscus* |
| Birds | *Pionus menstruus* |
| Birds | *Pionus sordidus* |
| Birds | *Pionus tumultuosus* |
| Birds | *Pipile cumanensis* |
| Birds | *Pipra filicauda* |
| Birds | *Pipraeidea melanonota* |
| Birds | *Pipreola arcuata* |
| Birds | *Pipreola aureopectus* |
| Birds | *Pipreola chlorolepidota* |
| Birds | *Pipreola jucunda* |
| Birds | *Pipreola lubomirskii* |
| Birds | *Pipreola riefferii* |
| Birds | *Piprites chloris* |
| Birds | *Piranga flava* |
| Birds | *Piranga leucoptera* |
| Birds | *Piranga rubriceps* |
| Birds | *Pitangus lictor* |
| Birds | *Pitangus sulphuratus* |
| Birds | *Pithys albifrons* |
| Birds | *Pittasoma michleri* |
| Birds | *Pittasoma rufopileatum* |
| Birds | *Platalea ajaja* |
| Birds | *Platyrinchus coronatus* |
| Birds | *Platyrinchus flavigularis* |
| Birds | *Platyrinchus mystaceus* |
| Birds | *Platyrinchus platyrhynchos* |
| Birds | *Platyrinchus saturatus* |
| Birds | *Plegadis falcinellus* |
| Birds | *Podiceps andinus* |
| Birds | *Podiceps occipitalis* |
| Birds | *Podilymbus podiceps* |
| Birds | *Poecilostreptus palmeri* |
| Birds | *Poecilotriccus calopterus* |
| Birds | *Poecilotriccus capitalis* |
| Birds | *Poecilotriccus latirostris* |
| Birds | *Poecilotriccus ruficeps* |
| Birds | *Poecilotriccus sylvia* |
| Birds | *Poliocrania exsul* |
| Birds | *Polioptila facilis* |
| Birds | *Polioptila plumbea* |
| Birds | *Polioptila schistaceigula* |
| Birds | *Polyerata amabilis* |
| Birds | *Polyerata rosenbergi* |
| Birds | *Polystictus pectoralis* |
| Birds | *Polytmus guainumbi* |
| Birds | *Polytmus theresiae* |
| Birds | *Porphyrio martinica* |
| Birds | *Porphyriops melanops* |
| Birds | *Porphyrolaema porphyrolaema* |
| Birds | *Porzana flaviventer* |
| Birds | *Premnoplex brunnescens* |
| Birds | *Premnornis guttuliger* |
| Birds | *Procnias averano* |
| Birds | *Progne chalybea* |
| Birds | *Progne tapera* |
| Birds | *Psarocolius angustifrons* |
| Birds | *Psarocolius bifasciatus* |
| Birds | *Psarocolius cassini* |
| Birds | *Psarocolius decumanus* |
| Birds | *Psarocolius guatimozinus* |
| Birds | *Psarocolius viridis* |
| Birds | *Psarocolius wagleri* |
| Birds | *Pseudastur albicollis* |
| Birds | *Pseudocolaptes boissonneautii* |
| Birds | *Pseudocolaptes johnstoni* |
| Birds | *Pseudocolopteryx acutipennis* |
| Birds | *Pseudopipra pipra* |
| Birds | *Pseudospingus verticalis* |
| Birds | *Pseudotriccus pelzelni* |
| Birds | *Pseudotriccus ruficeps* |
| Birds | *Psittacara leucophthalmus* |
| Birds | *Psittacara wagleri* |
| Birds | *Psophia crepitans* |
| Birds | *Pteroglossus azara* |
| Birds | *Pteroglossus castanotis* |
| Birds | *Pteroglossus inscriptus* |
| Birds | *Pteroglossus pluricinctus* |
| Birds | *Pteroglossus torquatus* |
| Birds | *Pterophanes cyanopterus* |
| Birds | *Puffinus lherminieri* |
| Birds | *Pulsatrix melanota* |
| Birds | *Pulsatrix perspicillata* |
| Birds | *Pygiptila stellaris* |
| Birds | *Pygochelidon cyanoleuca* |
| Birds | *Pygochelidon melanoleuca* |
| Birds | *Pyriglena leuconota* |
| Birds | *Pyrilia barrabandi* |
| Birds | *Pyrilia haematotis* |
| Birds | *Pyrilia pulchra* |
| Birds | *Pyrilia pyrilia* |
| Birds | *Pyrocephalus rubinus* |
| Birds | *Pyroderus scutatus* |
| Birds | *Pyrrhomyias cinnamomeus* |
| Birds | *Pyrrhura calliptera* |
| Birds | *Pyrrhura melanura* |
| Birds | *Pyrrhura picta* |
| Birds | *Pyrrhura viridicata* |
| Birds | *Querula purpurata* |
| Birds | *Quiscalus lugubris* |
| Birds | *Quiscalus mexicanus* |
| Birds | *Rallus limicola* |
| Birds | *Rallus longirostris* |
| Birds | *Rallus semiplumbeus* |
| Birds | *Ramphastos ambiguus* |
| Birds | *Ramphastos brevis* |
| Birds | *Ramphastos sulfuratus* |
| Birds | *Ramphastos tucanus* |
| Birds | *Ramphastos vitellinus* |
| Birds | *Ramphocaenus melanurus* |
| Birds | *Ramphocelus carbo* |
| Birds | *Ramphocelus dimidiatus* |
| Birds | *Ramphocelus flammigerus* |
| Birds | *Ramphocelus nigrogularis* |
| Birds | *Ramphomicron dorsale* |
| Birds | *Ramphomicron microrhynchum* |
| Birds | *Ramphotrigon fuscicauda* |
| Birds | *Ramphotrigon megacephalum* |
| Birds | *Ramphotrigon ruficauda* |
| Birds | *Rhegmatorhina cristata* |
| Birds | *Rhegmatorhina melanosticta* |
| Birds | *Rhodinocichla rosea* |
| Birds | *Rhynchocyclus brevirostris* |
| Birds | *Rhynchocyclus fulvipectus* |
| Birds | *Rhynchocyclus olivaceus* |
| Birds | *Rhynchocyclus pacificus* |
| Birds | *Rhynchortyx cinctus* |
| Birds | *Rhytipterna holerythra* |
| Birds | *Rhytipterna immunda* |
| Birds | *Rhytipterna simplex* |
| Birds | *Rostrhamus sociabilis* |
| Birds | *Rupicola peruvianus* |
| Birds | *Rupicola rupicola* |
| Birds | *Rupornis magnirostris* |
| Birds | *Rynchops niger* |
| Birds | *Sakesphorus canadensis* |
| Birds | *Saltator atripennis* |
| Birds | *Saltator cinctus* |
| Birds | *Saltator coerulescens* |
| Birds | *Saltator grossus* |
| Birds | *Saltator maximus* |
| Birds | *Saltator olivascens* |
| Birds | *Saltator orenocensis* |
| Birds | *Saltator striatipectus* |
| Birds | *Sapayoa aenigma* |
| Birds | *Sarcoramphus papa* |
| Birds | *Sarkidiornis sylvicola* |
| Birds | *Saucerottia castaneiventris* |
| Birds | *Saucerottia cyanifrons* |
| Birds | *Saucerottia edward* |
| Birds | *Saucerottia saucerottei* |
| Birds | *Saucerottia viridigaster* |
| Birds | *Sayornis nigricans* |
| Birds | *Schiffornis aenea* |
| Birds | *Schiffornis major* |
| Birds | *Schiffornis stenorhyncha* |
| Birds | *Schiffornis turdina* |
| Birds | *Schiffornis veraepacis* |
| Birds | *Schistes albogularis* |
| Birds | *Schistes geoffroyi* |
| Birds | *Schistochlamys melanopis* |
| Birds | *Sciaphylax castanea* |
| Birds | *Sclateria naevia* |
| Birds | *Sclerurus albigularis* |
| Birds | *Sclerurus caudacutus* |
| Birds | *Sclerurus guatemalensis* |
| Birds | *Sclerurus obscurior* |
| Birds | *Sclerurus rufigularis* |
| Birds | *Scytalopus alvarezlopezi* |
| Birds | *Scytalopus atratus* |
| Birds | *Scytalopus canus* |
| Birds | *Scytalopus chocoensis* |
| Birds | *Scytalopus griseicollis* |
| Birds | *Scytalopus latebricola* |
| Birds | *Scytalopus latrans* |
| Birds | *Scytalopus micropterus* |
| Birds | *Scytalopus opacus* |
| Birds | *Scytalopus panamensis* |
| Birds | *Scytalopus perijanus* |
| Birds | *Scytalopus rodriguezi* |
| Birds | *Scytalopus sanctaemartae* |
| Birds | *Scytalopus spillmanni* |
| Birds | *Scytalopus stilesi* |
| Birds | *Scytalopus vicinior* |
| Birds | *Selenidera nattereri* |
| Birds | *Selenidera reinwardtii* |
| Birds | *Selenidera spectabilis* |
| Birds | *Semnornis ramphastinus* |
| Birds | *Sericossypha albocristata* |
| Birds | *Serpophaga cinerea* |
| Birds | *Serpophaga hypoleuca* |
| Birds | *Setopagis heterura* |
| Birds | *Setophaga pitiayumi* |
| Birds | *Sicalis citrina* |
| Birds | *Sicalis columbiana* |
| Birds | *Sicalis flaveola* |
| Birds | *Sicalis luteola* |
| Birds | *Sipia berlepschi* |
| Birds | *Sipia laemosticta* |
| Birds | *Sipia nigricauda* |
| Birds | *Sipia palliata* |
| Birds | *Siptornis striaticollis* |
| Birds | *Sirystes albocinereus* |
| Birds | *Sirystes albogriseus* |
| Birds | *Sittasomus griseicapillus* |
| Birds | *Snowornis cryptolophus* |
| Birds | *Snowornis subalaris* |
| Birds | *Spatula cyanoptera* |
| Birds | *Spatula discors* |
| Birds | *Sphenopsis frontalis* |
| Birds | *Sphenopsis melanotis* |
| Birds | *Spinus cucullatus* |
| Birds | *Spinus magellanicus* |
| Birds | *Spinus psaltria* |
| Birds | *Spinus spinescens* |
| Birds | *Spinus xanthogastrus* |
| Birds | *Spizaetus isidori* |
| Birds | *Spizaetus melanoleucus* |
| Birds | *Spizaetus ornatus* |
| Birds | *Spizaetus tyrannus* |
| Birds | *Sporathraupis cyanocephala* |
| Birds | *Sporophila americana* |
| Birds | *Sporophila angolensis* |
| Birds | *Sporophila atrirostris* |
| Birds | *Sporophila bouvronides* |
| Birds | *Sporophila caerulescens* |
| Birds | *Sporophila castaneiventris* |
| Birds | *Sporophila corvina* |
| Birds | *Sporophila crassirostris* |
| Birds | *Sporophila fringilloides* |
| Birds | *Sporophila funerea* |
| Birds | *Sporophila intermedia* |
| Birds | *Sporophila lineola* |
| Birds | *Sporophila luctuosa* |
| Birds | *Sporophila maximiliani* |
| Birds | *Sporophila minuta* |
| Birds | *Sporophila nigricollis* |
| Birds | *Sporophila plumbea* |
| Birds | *Sporophila schistacea* |
| Birds | *Sporophila telasco* |
| Birds | *Steatornis caripensis* |
| Birds | *Stelgidopteryx ruficollis* |
| Birds | *Sternoclyta cyanopectus* |
| Birds | *Sternula antillarum* |
| Birds | *Sternula superciliaris* |
| Birds | *Stigmatura napensis* |
| Birds | *Stilpnia cayana* |
| Birds | *Stilpnia cyanicollis* |
| Birds | *Stilpnia cyanoptera* |
| Birds | *Stilpnia heinei* |
| Birds | *Stilpnia larvata* |
| Birds | *Stilpnia nigrocincta* |
| Birds | *Stilpnia vitriolina* |
| Birds | *Streptoprocne rutila* |
| Birds | *Streptoprocne zonaris* |
| Birds | *Strix albitarsis* |
| Birds | *Strix huhula* |
| Birds | *Strix nigrolineata* |
| Birds | *Strix virgata* |
| Birds | *Sturnella magna* |
| Birds | *Sublegatus arenarum* |
| Birds | *Sublegatus obscurior* |
| Birds | *Sula dactylatra* |
| Birds | *Sula granti* |
| Birds | *Sula leucogaster* |
| Birds | *Sula nebouxii* |
| Birds | *Sula sula* |
| Birds | *Synallaxis albescens* |
| Birds | *Synallaxis albigularis* |
| Birds | *Synallaxis azarae* |
| Birds | *Synallaxis brachyura* |
| Birds | *Synallaxis candei* |
| Birds | *Synallaxis cherriei* |
| Birds | *Synallaxis cinnamomea* |
| Birds | *Synallaxis fuscorufa* |
| Birds | *Synallaxis gujanensis* |
| Birds | *Synallaxis moesta* |
| Birds | *Synallaxis rutilans* |
| Birds | *Synallaxis subpudica* |
| Birds | *Synallaxis unirufa* |
| Birds | *Syndactyla subalaris* |
| Birds | *Syrigma sibilatrix* |
| Birds | *Systellura longirostris* |
| Birds | *Tachornis furcata* |
| Birds | *Tachornis squamata* |
| Birds | *Tachybaptus dominicus* |
| Birds | *Tachycineta albiventer* |
| Birds | *Tachyphonus delatrii* |
| Birds | *Tachyphonus phoenicius* |
| Birds | *Tachyphonus rufus* |
| Birds | *Tachyphonus surinamus* |
| Birds | *Talaphorus chlorocercus* |
| Birds | *Tangara arthus* |
| Birds | *Tangara callophrys* |
| Birds | *Tangara chilensis* |
| Birds | *Tangara chrysotis* |
| Birds | *Tangara cyanotis* |
| Birds | *Tangara florida* |
| Birds | *Tangara gyrola* |
| Birds | *Tangara icterocephala* |
| Birds | *Tangara inornata* |
| Birds | *Tangara johannae* |
| Birds | *Tangara labradorides* |
| Birds | *Tangara lavinia* |
| Birds | *Tangara mexicana* |
| Birds | *Tangara nigroviridis* |
| Birds | *Tangara parzudakii* |
| Birds | *Tangara schrankii* |
| Birds | *Tangara vassorii* |
| Birds | *Tangara velia* |
| Birds | *Tangara xanthocephala* |
| Birds | *Tapera naevia* |
| Birds | *Taphrospilus hypostictus* |
| Birds | *Taraba major* |
| Birds | *Tephrophilus wetmorei* |
| Birds | *Terenotriccus erythrurus* |
| Birds | *Tersina viridis* |
| Birds | *Thalurania colombica* |
| Birds | *Thalurania furcata* |
| Birds | *Thamnistes anabatinus* |
| Birds | *Thamnomanes ardesiacus* |
| Birds | *Thamnomanes caesius* |
| Birds | *Thamnophilus aethiops* |
| Birds | *Thamnophilus amazonicus* |
| Birds | *Thamnophilus atrinucha* |
| Birds | *Thamnophilus cryptoleucus* |
| Birds | *Thamnophilus doliatus* |
| Birds | *Thamnophilus melanonotus* |
| Birds | *Thamnophilus multistriatus* |
| Birds | *Thamnophilus murinus* |
| Birds | *Thamnophilus nigriceps* |
| Birds | *Thamnophilus nigrocinereus* |
| Birds | *Thamnophilus praecox* |
| Birds | *Thamnophilus punctatus* |
| Birds | *Thamnophilus schistaceus* |
| Birds | *Thamnophilus tenuepunctatus* |
| Birds | *Thamnophilus unicolor* |
| Birds | *Thectocercus acuticaudatus* |
| Birds | *Theristicus caudatus* |
| Birds | *Thlypopsis fulviceps* |
| Birds | *Thlypopsis ornata* |
| Birds | *Thlypopsis sordida* |
| Birds | *Thlypopsis superciliaris* |
| Birds | *Thraupis episcopus* |
| Birds | *Thraupis glaucocolpa* |
| Birds | *Thraupis palmarum* |
| Birds | *Threnetes leucurus* |
| Birds | *Threnetes ruckeri* |
| Birds | *Thripadectes flammulatus* |
| Birds | *Thripadectes holostictus* |
| Birds | *Thripadectes ignobilis* |
| Birds | *Thripadectes melanorhynchus* |
| Birds | *Thripadectes virgaticeps* |
| Birds | *Thripophaga cherriei* |
| Birds | *Thryophilus nicefori* |
| Birds | *Thryophilus rufalbus* |
| Birds | *Thryophilus sernai* |
| Birds | *Tiaris olivaceus* |
| Birds | *Tigrisoma fasciatum* |
| Birds | *Tigrisoma lineatum* |
| Birds | *Tigrisoma mexicanum* |
| Birds | *Tinamus guttatus* |
| Birds | *Tinamus major* |
| Birds | *Tinamus osgoodi* |
| Birds | *Tinamus tao* |
| Birds | *Tityra cayana* |
| Birds | *Tityra inquisitor* |
| Birds | *Tityra semifasciata* |
| Birds | *Todirostrum chrysocrotaphum* |
| Birds | *Todirostrum cinereum* |
| Birds | *Todirostrum maculatum* |
| Birds | *Todirostrum nigriceps* |
| Birds | *Tolmomyias assimilis* |
| Birds | *Tolmomyias flaviventris* |
| Birds | *Tolmomyias poliocephalus* |
| Birds | *Tolmomyias sulphurescens* |
| Birds | *Tolmomyias traylori* |
| Birds | *Topaza pyra* |
| Birds | *Touit batavicus* |
| Birds | *Touit dilectissimus* |
| Birds | *Touit huetii* |
| Birds | *Touit purpuratus* |
| Birds | *Touit stictopterus* |
| Birds | *Troglodytes aedon* |
| Birds | *Troglodytes monticola* |
| Birds | *Troglodytes ochraceus* |
| Birds | *Troglodytes solstitialis* |
| Birds | *Trogon caligatus* |
| Birds | *Trogon chionurus* |
| Birds | *Trogon collaris* |
| Birds | *Trogon comptus* |
| Birds | *Trogon cupreicauda* |
| Birds | *Trogon curucui* |
| Birds | *Trogon massena* |
| Birds | *Trogon melanurus* |
| Birds | *Trogon personatus* |
| Birds | *Trogon ramonianus* |
| Birds | *Trogon rufus* |
| Birds | *Trogon tenellus* |
| Birds | *Trogon viridis* |
| Birds | *Tunchiornis ochraceiceps* |
| Birds | *Turdus albicollis* |
| Birds | *Turdus arthuri* |
| Birds | *Turdus assimilis* |
| Birds | *Turdus flavipes* |
| Birds | *Turdus fulviventris* |
| Birds | *Turdus fumigatus* |
| Birds | *Turdus fuscater* |
| Birds | *Turdus grayi* |
| Birds | *Turdus hauxwelli* |
| Birds | *Turdus ignobilis* |
| Birds | *Turdus lawrencii* |
| Birds | *Turdus leucomelas* |
| Birds | *Turdus leucops* |
| Birds | *Turdus nudigenis* |
| Birds | *Turdus obsoletus* |
| Birds | *Turdus olivater* |
| Birds | *Turdus sanchezorum* |
| Birds | *Turdus serranus* |
| Birds | *Tyranneutes stolzmanni* |
| Birds | *Tyrannopsis sulphurea* |
| Birds | *Tyrannulus elatus* |
| Birds | *Tyrannus melancholicus* |
| Birds | *Tyrannus savana* |
| Birds | *Tyto alba* |
| Birds | *Uranomitra franciae* |
| Birds | *Urochroa bougueri* |
| Birds | *Urochroa leucura* |
| Birds | *Uromyias agilis* |
| Birds | *Uropsalis lyra* |
| Birds | *Uropsalis segmentata* |
| Birds | *Urosticte benjamini* |
| Birds | *Urosticte ruficrissa* |
| Birds | *Urothraupis stolzmanni* |
| Birds | *Vanellus cayanus* |
| Birds | *Vanellus chilensis* |
| Birds | *Vanellus resplendens* |
| Birds | *Vireo approximans* |
| Birds | *Vireo caribaeus* |
| Birds | *Vireo chivi* |
| Birds | *Vireo leucophrys* |
| Birds | *Vireo masteri* |
| Birds | *Vireolanius eximius* |
| Birds | *Vireolanius leucotis* |
| Birds | *Volatinia jacarina* |
| Birds | *Vultur gryphus* |
| Birds | *Willisornis poecilinotus* |
| Birds | *Xenerpestes minlosi* |
| Birds | *Xenopipo atronitens* |
| Birds | *Xenops minutus* |
| Birds | *Xenops rutilans* |
| Birds | *Xenops tenuirostris* |
| Birds | *Xenornis setifrons* |
| Birds | *Xiphocolaptes promeropirhynchus* |
| Birds | *Xipholena punicea* |
| Birds | *Xiphorhynchus elegans* |
| Birds | *Xiphorhynchus erythropygius* |
| Birds | *Xiphorhynchus guttatus* |
| Birds | *Xiphorhynchus lachrymosus* |
| Birds | *Xiphorhynchus obsoletus* |
| Birds | *Xiphorhynchus ocellatus* |
| Birds | *Xiphorhynchus susurrans* |
| Birds | *Xiphorhynchus triangularis* |
| Birds | *Zebrilus undulatus* |
| Birds | *Zenaida auriculata* |
| Birds | *Zentrygon frenata* |
| Birds | *Zentrygon goldmani* |
| Birds | *Zentrygon linearis* |
| Birds | *Zimmerius albigularis* |
| Birds | *Zimmerius chrysops* |
| Birds | *Zimmerius gracilipes* |
| Birds | *Zimmerius improbus* |
| Birds | *Zimmerius vilissimus* |
| Birds | *Zonotrichia capensis* |
| Fishes | *Thalassophryne amazonica* |
| Fishes | *Pseudotylosurus microps* |
| Fishes | *Hyporhamphus brederi* |
| Fishes | *Acestrorhynchus abbreviatus* |
| Fishes | *Acestrorhynchus falcatus* |
| Fishes | *Acestrorhynchus falcirostris* |
| Fishes | *Acestrorhynchus grandoculis* |
| Fishes | *Acestrorhynchus heterolepis* |
| Fishes | *Acestrorhynchus microlepis* |
| Fishes | *Acestrorhynchus minimus* |
| Fishes | *Acestrorhynchus nasutus* |
| Fishes | *Abramites eques* |
| Fishes | *Abramites hypselonotus* |
| Fishes | *Anodus elongatus* |
| Fishes | *Anodus orinocensis* |
| Fishes | *Anostomoides atrianalis* |
| Fishes | *Anostomus anostomus* |
| Fishes | *Anostomus ternetzi* |
| Fishes | *Cheirocerus abuelo* |
| Fishes | *Cheirocerus goeldii* |
| Fishes | *Compsaraia compsa* |
| Fishes | *Dentectus barbarmatus* |
| Fishes | *Engraulisoma taeniatum* |
| Fishes | *Haemomaster venezuelae* |
| Fishes | *Hemiodontichthys acipenserinus* |
| Fishes | *Hemiodus amazonum* |
| Fishes | *Hemiodus argenteus* |
| Fishes | *Hemiodus atranalis* |
| Fishes | *Hemiodus gracilis* |
| Fishes | *Hemiodus immaculatus* |
| Fishes | *Hemiodus microlepis* |
| Fishes | *Hemiodus semitaeniatus* |
| Fishes | *Hemiodus thayeria* |
| Fishes | *Hemiodus unimaculatus* |
| Fishes | *Hemiodus vorderwinkleri* |
| Fishes | *Laemolyta fernandezi* |
| Fishes | *Laemolyta garmani* |
| Fishes | *Laemolyta proxima* |
| Fishes | *Laemolyta taeniata* |
| Fishes | *Leporinus agassizii* |
| Fishes | *Leporinus altipinnis* |
| Fishes | *Leporinus amazonicus* |
| Fishes | *Leporinus arimaspi* |
| Fishes | *Leporinus boehlkei* |
| Fishes | *Leporinus brunneus* |
| Fishes | *Leporinus enyae* |
| Fishes | *Leporinus fasciatus* |
| Fishes | *Leporinus friderici* |
| Fishes | *Leporinus granti* |
| Fishes | *Leporinus jamesi* |
| Fishes | *Leporinus klausewitzi* |
| Fishes | *Leporinus moralesi* |
| Fishes | *Leporinus niceforoi* |
| Fishes | *Leporinus ortomaculatus* |
| Fishes | *Leporinus parae* |
| Fishes | *Leporinus punctatus* |
| Fishes | *Leporinus striatus* |
| Fishes | *Leporinus subniger* |
| Fishes | *Leporinus y-ophorus* |
| Fishes | *Megaleporinus muyscorum* |
| Fishes | *Megaleporinus trifasciatus* |
| Fishes | *Schizodon corti* |
| Fishes | *Schizodon fasciatus* |
| Fishes | *Schizodon scotorhabdotus* |
| Fishes | *Stichonodon insignis* |
| Fishes | *Chalceus epakros* |
| Fishes | *Chalceus erythrurus* |
| Fishes | *Chalceus macrolepidotus* |
| Fishes | *Acestridium colombiensis* |
| Fishes | *Acestridium dichromum* |
| Fishes | *Acestridium martini* |
| Fishes | *Acestrocephalus anomalus* |
| Fishes | *Acestrocephalus boehlkei* |
| Fishes | *Acestrocephalus sardina* |
| Fishes | *Agoniates anchovia* |
| Fishes | *Agoniates halecinus* |
| Fishes | *Amazonsprattus scintilla* |
| Fishes | *Ammocryptocharax elegans* |
| Fishes | *Ammocryptocharax minutus* |
| Fishes | *Aphyocharax colifax* |
| Fishes | *Aphyocharax erythrurus* |
| Fishes | *Aphyocharax pusillus* |
| Fishes | *Apionichthys nattereri* |
| Fishes | *Apionichthys sauli* |
| Fishes | *Argonectes longiceps* |
| Fishes | *Astyanax atratoensis* |
| Fishes | *Astyanax bimaculatus* |
| Fishes | *Astyanax caucanus* |
| Fishes | *Astyanax filiferus* |
| Fishes | *Astyanax gisleni* |
| Fishes | *Astyanax integer* |
| Fishes | *Astyanax magdalenae* |
| Fishes | *Astyanax maximus* |
| Fishes | *Astyanax megaspilura* |
| Fishes | *Astyanax metae* |
| Fishes | *Astyanax microlepis* |
| Fishes | *Astyanax orthodus* |
| Fishes | *Astyanax panamensis* |
| Fishes | *Astyanax scintillans* |
| Fishes | *Astyanax siapae* |
| Fishes | *Astyanax stilbe* |
| Fishes | *Astyanax superbus* |
| Fishes | *Astyanax venezuelae* |
| Fishes | *Astyanax villwocki* |
| Fishes | *Astyanax yariguies* |
| Fishes | *Atopomesus pachyodus* |
| Fishes | *Axelrodia riesei* |
| Fishes | *Axelrodia stigmatias* |
| Fishes | *Bario steindachneri* |
| Fishes | *Boehlkea fredcochui* |
| Fishes | *Brachychalcinus copei* |
| Fishes | *Brachychalcinus nummus* |
| Fishes | *Brittanichthys axelrodi* |
| Fishes | *Brittanichthys myersi* |
| Fishes | *Brycon amazonicus* |
| Fishes | *Brycon argenteus* |
| Fishes | *Brycon dentex* |
| Fishes | *Brycon falcatus* |
| Fishes | *Brycon fowleri* |
| Fishes | *Brycon henni* |
| Fishes | *Brycon hilarii* |
| Fishes | *Brycon labiatus* |
| Fishes | *Brycon medemi* |
| Fishes | *Brycon meeki* |
| Fishes | *Brycon melanopterus* |
| Fishes | *Brycon moorei* |
| Fishes | *Brycon oligolepis* |
| Fishes | *Brycon pesu* |
| Fishes | *Brycon polylepis* |
| Fishes | *Brycon posadae* |
| Fishes | *Brycon rubricauda* |
| Fishes | *Brycon sinuensis* |
| Fishes | *Brycon striatulus* |
| Fishes | *Brycon whitei* |
| Fishes | *Bryconamericus carlosi* |
| Fishes | *Bryconamericus guizae* |
| Fishes | *Bryconamericus icelus* |
| Fishes | *Bryconamericus macrophthalmus* |
| Fishes | *Bryconella pallidifrons* |
| Fishes | *Bryconops affinis* |
| Fishes | *Bryconops alburnoides* |
| Fishes | *Bryconops caudomaculatus* |
| Fishes | *Bryconops collettei* |
| Fishes | *Bryconops giacopinii* |
| Fishes | *Bryconops humeralis* |
| Fishes | *Bryconops inpai* |
| Fishes | *Bryconops magoi* |
| Fishes | *Caenotropus labyrinthicus* |
| Fishes | *Caenotropus mestomorgmatos* |
| Fishes | *Charax apurensis* |
| Fishes | *Charax condei* |
| Fishes | *Charax metae* |
| Fishes | *Charax michaeli* |
| Fishes | *Charax niger* |
| Fishes | *Charax notulatus* |
| Fishes | *Charax pauciradiatus* |
| Fishes | *Charax tectifer* |
| Fishes | *Cheirodontops geayi* |
| Fishes | *Chrysobrycon guahibo* |
| Fishes | *Chrysobrycon hesperus* |
| Fishes | *Chrysobrycon mojicai* |
| Fishes | *Corynopoma riisei* |
| Fishes | *Ctenobrycon oliverai* |
| Fishes | *Ctenobrycon spilurus* |
| Fishes | *Cyanogaster noctivaga* |
| Fishes | *Cynodon gibbus* |
| Fishes | *Cynodon septenarius* |
| Fishes | *Denticetopsis seducta* |
| Fishes | *Duopalatinus peruanus* |
| Fishes | *Dupouyichthys sapito* |
| Fishes | *Elachocharax geryi* |
| Fishes | *Elachocharax mitopterus* |
| Fishes | *Elachocharax pulcher* |
| Fishes | *Entomocorus gameroi* |
| Fishes | *Epapterus blohmi* |
| Fishes | *Epapterus dispilurus* |
| Fishes | *Eremophilus mutisii* |
| Fishes | *Eretmobrycon guaytarae* |
| Fishes | *Eretmobrycon peruanus* |
| Fishes | *Exallodontus aguanai* |
| Fishes | *Exodon paradoxus* |
| Fishes | *Galeocharax gulo* |
| Fishes | *Genycharax tarpon* |
| Fishes | *Gephyrocharax caucanus* |
| Fishes | *Gephyrocharax chocoensis* |
| Fishes | *Gephyrocharax melanocheir* |
| Fishes | *Gephyrocharax sinuensis* |
| Fishes | *Gephyrocharax valencia* |
| Fishes | *Gephyrocharax venezuelae* |
| Fishes | *Gilbertolus alatus* |
| Fishes | *Gilbertolus atratoensis* |
| Fishes | *Gilbertolus maracaiboensis* |
| Fishes | *Gnathocharax steindachneri* |
| Fishes | *Goeldiella eques* |
| Fishes | *Gymnocorymbus bondi* |
| Fishes | *Gymnocorymbus thayeri* |
| Fishes | *Hemibrycon antioquiae* |
| Fishes | *Hemibrycon boquiae* |
| Fishes | *Hemibrycon cardalensis* |
| Fishes | *Hemibrycon carrilloi* |
| Fishes | *Hemibrycon clausen* |
| Fishes | *Hemibrycon colombianus* |
| Fishes | *Hemibrycon dariensis* |
| Fishes | *Hemibrycon decurrens* |
| Fishes | *Hemibrycon dentatus* |
| Fishes | *Hemibrycon fasciatus* |
| Fishes | *Hemibrycon iqueima* |
| Fishes | *Hemibrycon jabonero* |
| Fishes | *Hemibrycon jelskii* |
| Fishes | *Hemibrycon metae* |
| Fishes | *Hemibrycon paez* |
| Fishes | *Hemibrycon palomae* |
| Fishes | *Hemibrycon rafaelense* |
| Fishes | *Hemibrycon raqueliae* |
| Fishes | *Hemibrycon sanjuanensis* |
| Fishes | *Hemibrycon santamartae* |
| Fishes | *Hemibrycon sierraensis* |
| Fishes | *Hemibrycon tolimae* |
| Fishes | *Hemibrycon velox* |
| Fishes | *Hemibrycon virolinica* |
| Fishes | *Hemibrycon yacopiae* |
| Fishes | *Hemigrammus aguaruna* |
| Fishes | *Hemigrammus amacayacu* |
| Fishes | *Hemigrammus analis* |
| Fishes | *Hemigrammus barrigonae* |
| Fishes | *Hemigrammus bellottii* |
| Fishes | *Hemigrammus coeruleus* |
| Fishes | *Hemigrammus diagonicus* |
| Fishes | *Hemigrammus geisleri* |
| Fishes | *Hemigrammus hyanuary* |
| Fishes | *Hemigrammus levis* |
| Fishes | *Hemigrammus luelingi* |
| Fishes | *Hemigrammus lunatus* |
| Fishes | *Hemigrammus melanochrous* |
| Fishes | *Hemigrammus micropterus* |
| Fishes | *Hemigrammus microstomus* |
| Fishes | *Hemigrammus mimus* |
| Fishes | *Hemigrammus newboldi* |
| Fishes | *Hemigrammus ocellifer* |
| Fishes | *Hemigrammus orthus* |
| Fishes | *Hemigrammus pretoensis* |
| Fishes | *Hemigrammus pulcher* |
| Fishes | *Hemigrammus rubrostriatus* |
| Fishes | *Hemigrammus schmardae* |
| Fishes | *Hemigrammus stictus* |
| Fishes | *Hemigrammus unilineatus* |
| Fishes | *Hemigrammus vorderwinkleri* |
| Fishes | *Hemigrammus xaveriellus* |
| Fishes | *Hemigrammus yinyang* |
| Fishes | *Hemus punctatus* |
| Fishes | *Hemus triacanthopomus* |
| Fishes | *Hyphessobrycon acaciae* |
| Fishes | *Hyphessobrycon agulha* |
| Fishes | *Hyphessobrycon amaronensis* |
| Fishes | *Hyphessobrycon bentosi* |
| Fishes | *Hyphessobrycon chiribiquete* |
| Fishes | *Hyphessobrycon condotensis* |
| Fishes | *Hyphessobrycon copelandi* |
| Fishes | *Hyphessobrycon diancistrus* |
| Fishes | *Hyphessobrycon dorsalis* |
| Fishes | *Hyphessobrycon epicharis* |
| Fishes | *Hyphessobrycon erythrostigma* |
| Fishes | *Hyphessobrycon gracilior* |
| Fishes | *Hyphessobrycon klausanni* |
| Fishes | *Hyphessobrycon loretoensis* |
| Fishes | *Hyphessobrycon mavro* |
| Fishes | *Hyphessobrycon metae* |
| Fishes | *Hyphessobrycon natagaima* |
| Fishes | *Hyphessobrycon niger* |
| Fishes | *Hyphessobrycon ocasoensis* |
| Fishes | *Hyphessobrycon oritoensis* |
| Fishes | *Hyphessobrycon otrynus* |
| Fishes | *Hyphessobrycon peruvianus* |
| Fishes | *Hyphessobrycon poecilioides* |
| Fishes | *Hyphessobrycon proteus* |
| Fishes | *Hyphessobrycon rheophilus* |
| Fishes | *Hyphessobrycon saizi* |
| Fishes | *Hyphessobrycon sovichthys* |
| Fishes | *Hyphessobrycon sweglesi* |
| Fishes | *Hyphessobrycon taguae* |
| Fishes | *Hyphessobrycon tropis* |
| Fishes | *Hyphessobrycon tukunai* |
| Fishes | *Hypopygus lepturus* |
| Fishes | *Hypopygus neblinae* |
| Fishes | *Jupiaba abramoides* |
| Fishes | *Jupiaba anterior* |
| Fishes | *Jupiaba anteroides* |
| Fishes | *Jupiaba asymmetrica* |
| Fishes | *Jupiaba polylepis* |
| Fishes | *Jupiaba scologaster* |
| Fishes | *Jupiaba zonata* |
| Fishes | *Knodus breviceps* |
| Fishes | *Knodus cinarucoensis* |
| Fishes | *Knodus gamma* |
| Fishes | *Knodus heteresthes* |
| Fishes | *Knodus meridae* |
| Fishes | *Knodus orteguasae* |
| Fishes | *Knodus tiquiensis* |
| Fishes | *Lebiasina chocoensis* |
| Fishes | *Lebiasina chucuriensis* |
| Fishes | *Lebiasina colombia* |
| Fishes | *Lebiasina elongata* |
| Fishes | *Lebiasina erythrinoides* |
| Fishes | *Lebiasina festae* |
| Fishes | *Lebiasina floridablancaensis* |
| Fishes | *Lebiasina multimaculata* |
| Fishes | *Lebiasina narinensis* |
| Fishes | *Lebiasina ortegai* |
| Fishes | *Leptobrycon jatuaranae* |
| Fishes | *Microschemobrycon callops* |
| Fishes | *Microschemobrycon casiquiare* |
| Fishes | *Microschemobrycon geisleri* |
| Fishes | *Microschemobrycon melanotus* |
| Fishes | *Moenkhausia browni* |
| Fishes | *Moenkhausia ceros* |
| Fishes | *Moenkhausia chrysargyrea* |
| Fishes | *Moenkhausia collettii* |
| Fishes | *Moenkhausia comma* |
| Fishes | *Moenkhausia copei* |
| Fishes | *Moenkhausia cotinho* |
| Fishes | *Moenkhausia dichroura* |
| Fishes | *Moenkhausia diktyota* |
| Fishes | *Moenkhausia eigenmanni* |
| Fishes | *Moenkhausia gracilima* |
| Fishes | *Moenkhausia grandisquamis* |
| Fishes | *Moenkhausia hemigrammoides* |
| Fishes | *Moenkhausia hysterosticta* |
| Fishes | *Moenkhausia intermedia* |
| Fishes | *Moenkhausia jamesi* |
| Fishes | *Moenkhausia justae* |
| Fishes | *Moenkhausia lata* |
| Fishes | *Moenkhausia latissima* |
| Fishes | *Moenkhausia lepidura* |
| Fishes | *Moenkhausia margitae* |
| Fishes | *Moenkhausia megalops* |
| Fishes | *Moenkhausia melogramma* |
| Fishes | *Moenkhausia metae* |
| Fishes | *Moenkhausia mikia* |
| Fishes | *Moenkhausia oligolepis* |
| Fishes | *Moenkhausia orteguasae* |
| Fishes | *Moenkhausia robertsi* |
| Fishes | *Nanocheirodon insignis* |
| Fishes | *Nematobrycon lacortei* |
| Fishes | *Nematobrycon palmeri* |
| Fishes | *Odontocharacidium aphanes* |
| Fishes | *Odontostilbe fugitiva* |
| Fishes | *Odontostilbe pao* |
| Fishes | *Odontostilbe pulchra* |
| Fishes | *Odontostilbe splendida* |
| Fishes | *Oxybrycon parvulus* |
| Fishes | *Paracheirodon axelrodi* |
| Fishes | *Paracheirodon innesi* |
| Fishes | *Paracheirodon simulans* |
| Fishes | *Paragoniates alburnus* |
| Fishes | *Parapristella georgiae* |
| Fishes | *Parastremma album* |
| Fishes | *Parastremma sadina* |
| Fishes | *Paravandellia phaneronema* |
| Fishes | *Parecbasis cyclolepis* |
| Fishes | *Pariolius armillatus* |
| Fishes | *Petitella georgiae* |
| Fishes | *Phenagoniates macrolepis* |
| Fishes | *Poptella brevispina* |
| Fishes | *Poptella compressa* |
| Fishes | *Poptella longipinnis* |
| Fishes | *Priocharax ariel* |
| Fishes | *Priocharax pygmaeus* |
| Fishes | *Prionobrama filigera* |
| Fishes | *Pristella ariporo* |
| Fishes | *Pristella maxillaris* |
| Fishes | *Protocheirodon pi* |
| Fishes | *Pseudanos trimaculatus* |
| Fishes | *Pseudanos winterbottomi* |
| Fishes | *Pseudochalceus longianalis* |
| Fishes | *Pterobrycon landoni* |
| Fishes | *Rhinobrycon negrensis* |
| Fishes | *Roeboides affinis* |
| Fishes | *Roeboides araguaito* |
| Fishes | *Roeboides dayi* |
| Fishes | *Roeboides dientonito* |
| Fishes | *Roeboides myersii* |
| Fishes | *Roeboides numerosus* |
| Fishes | *Roeboides occidentalis* |
| Fishes | *Roestes ogilviei* |
| Fishes | *Saccoderma hastata* |
| Fishes | *Saccoderma melanostigma* |
| Fishes | *Saccoderma robusta* |
| Fishes | *Saccodon dariensis* |
| Fishes | *Salminus affinis* |
| Fishes | *Salminus hilarii* |
| Fishes | *Schultzichthys bondi* |
| Fishes | *Schultzichthys gracilis* |
| Fishes | *Scopaeocharax rhinodus* |
| Fishes | *Serrabrycon magoi* |
| Fishes | *Serrasalmus altuvei* |
| Fishes | *Serrasalmus calmoni* |
| Fishes | *Serrasalmus compressus* |
| Fishes | *Serrasalmus eigenmanni* |
| Fishes | *Serrasalmus elongatus* |
| Fishes | *Serrasalmus gouldingi* |
| Fishes | *Serrasalmus hollandi* |
| Fishes | *Serrasalmus irritans* |
| Fishes | *Serrasalmus maculatus* |
| Fishes | *Serrasalmus manueli* |
| Fishes | *Serrasalmus medinai* |
| Fishes | *Serrasalmus nalseni* |
| Fishes | *Serrasalmus rhombeus* |
| Fishes | *Serrasalmus sanchezi* |
| Fishes | *Serrasalmus striolatus* |
| Fishes | *Steindachnerina argentea* |
| Fishes | *Steindachnerina atratoensis* |
| Fishes | *Steindachnerina bimaculata* |
| Fishes | *Steindachnerina dobula* |
| Fishes | *Steindachnerina guentheri* |
| Fishes | *Steindachnerina hypostoma* |
| Fishes | *Steindachnerina planiventris* |
| Fishes | *Steindachnerina pupula* |
| Fishes | *Stethaprion erythrops* |
| Fishes | *Tenellus leporhinus* |
| Fishes | *Tenellus ternetzi* |
| Fishes | *Tenellus trimaculatus* |
| Fishes | *Tetragonopterus argenteus* |
| Fishes | *Tetragonopterus chalceus* |
| Fishes | *Tetragonopterus daguae* |
| Fishes | *Thayeria obliqua* |
| Fishes | *Thoracocharax securis* |
| Fishes | *Thoracocharax stellatus* |
| Fishes | *Thrissobrycon pectinifer* |
| Fishes | *Tridensimilis venezuelae* |
| Fishes | *Tridentopsis pearsoni* |
| Fishes | *Triportheus albus* |
| Fishes | *Triportheus angulatus* |
| Fishes | *Triportheus auritus* |
| Fishes | *Triportheus brachipomus* |
| Fishes | *Triportheus culter* |
| Fishes | *Triportheus magdalenae* |
| Fishes | *Triportheus orinocensis* |
| Fishes | *Triportheus pictus* |
| Fishes | *Triportheus rotundatus* |
| Fishes | *Triportheus venezuelensis* |
| Fishes | *Trochilocharax ornatus* |
| Fishes | *Tyttobrycon xeruini* |
| Fishes | *Tyttocharax cochui* |
| Fishes | *Tyttocharax madeirae* |
| Fishes | *Tyttocharax metae* |
| Fishes | *Xenagoniates bondi* |
| Fishes | *Xenurobrycon heterodon* |
| Fishes | *Xyliphius kryptos* |
| Fishes | *Xyliphius lepturus* |
| Fishes | *Xyliphius magdalenae* |
| Fishes | *Xyliphius melanopterus* |
| Fishes | *Chilodus gracilis* |
| Fishes | *Chilodus punctatus* |
| Fishes | *Chocoheros microlepis* |
| Fishes | *Acaronia nassa* |
| Fishes | *Acaronia vultuosa* |
| Fishes | *Amaralia hypsiura* |
| Fishes | *Characidium boavistae* |
| Fishes | *Characidium caucanum* |
| Fishes | *Characidium chupa* |
| Fishes | *Characidium declivirostre* |
| Fishes | *Characidium etheostoma* |
| Fishes | *Characidium longum* |
| Fishes | *Characidium pellucidum* |
| Fishes | *Characidium phoxocephalum* |
| Fishes | *Characidium pteroides* |
| Fishes | *Characidium purpuratum* |
| Fishes | *Characidium roesseli* |
| Fishes | *Characidium steindachneri* |
| Fishes | *Characidium zebra* |
| Fishes | *Crenuchus spilurus* |
| Fishes | *Gnathodolus bidens* |
| Fishes | *Heterocharax leptogrammus* |
| Fishes | *Heterocharax macrolepis* |
| Fishes | *Heterocharax virgulatus* |
| Fishes | *Leptocharacidium omospilus* |
| Fishes | *Makunaima guianensis* |
| Fishes | *Melanocharacidium dispilomma* |
| Fishes | *Melanocharacidium nigrum* |
| Fishes | *Melanocharacidium pectorale* |
| Fishes | *Microcharacidium gnomus* |
| Fishes | *Microcharacidium weitzmani* |
| Fishes | *Microgenys lativirgata* |
| Fishes | *Microgenys minuta* |
| Fishes | *Micromoema xiphophora* |
| Fishes | *Microphilypnus ternetzi* |
| Fishes | *Tympanopleura atronasus* |
| Fishes | *Tympanopleura brevis* |
| Fishes | *Tympanopleura piperata* |
| Fishes | *Boulengerella cuvieri* |
| Fishes | *Boulengerella lateristriga* |
| Fishes | *Boulengerella lucius* |
| Fishes | *Boulengerella maculata* |
| Fishes | *Boulengerella xyrekes* |
| Fishes | *Ctenolucius beani* |
| Fishes | *Ctenolucius hujeta* |
| Fishes | *Phenacogaster napoatilis* |
| Fishes | *Creagrutus affinis* |
| Fishes | *Creagrutus amoenus* |
| Fishes | *Creagrutus andaki* |
| Fishes | *Creagrutus atratus* |
| Fishes | *Creagrutus barrigai* |
| Fishes | *Creagrutus bolivari* |
| Fishes | *Creagrutus brevipinnis* |
| Fishes | *Creagrutus calai* |
| Fishes | *Creagrutus caucanus* |
| Fishes | *Creagrutus cochui* |
| Fishes | *Creagrutus dulima* |
| Fishes | *Creagrutus flavescens* |
| Fishes | *Creagrutus guanes* |
| Fishes | *Creagrutus gyrospilus* |
| Fishes | *Creagrutus hildebrandi* |
| Fishes | *Creagrutus machadoi* |
| Fishes | *Creagrutus maculosus* |
| Fishes | *Creagrutus magdalenae* |
| Fishes | *Creagrutus maracaiboensis* |
| Fishes | *Creagrutus maxillaris* |
| Fishes | *Creagrutus melasma* |
| Fishes | *Creagrutus nigrostigmatus* |
| Fishes | *Creagrutus paralacus* |
| Fishes | *Creagrutus phasma* |
| Fishes | *Creagrutus runa* |
| Fishes | *Creagrutus taphorni* |
| Fishes | *Creagrutus tuyuka* |
| Fishes | *Creagrutus vexillapinnus* |
| Fishes | *Creagrutus xiphos* |
| Fishes | *Curimata aspera* |
| Fishes | *Curimata cerasina* |
| Fishes | *Curimata cisandina* |
| Fishes | *Curimata cyprinoides* |
| Fishes | *Curimata incompta* |
| Fishes | *Curimata inornata* |
| Fishes | *Curimata mivartii* |
| Fishes | *Curimata ocellata* |
| Fishes | *Curimata roseni* |
| Fishes | *Curimata vittata* |
| Fishes | *Curimatella alburna* |
| Fishes | *Curimatella dorsalis* |
| Fishes | *Curimatella immaculata* |
| Fishes | *Curimatella meyeri* |
| Fishes | *Curimatopsis cryptica* |
| Fishes | *Curimatopsis evelynae* |
| Fishes | *Curimatopsis macrolepis* |
| Fishes | *Curimatopsis microlepis* |
| Fishes | *Cynopotamus amazonum* |
| Fishes | *Cynopotamus atratoensis* |
| Fishes | *Cynopotamus bipunctatus* |
| Fishes | *Cynopotamus magdalenae* |
| Fishes | *Cynopotamus venezuelae* |
| Fishes | *Cyphocharax abramoides* |
| Fishes | *Cyphocharax aspilos* |
| Fishes | *Cyphocharax festivus* |
| Fishes | *Cyphocharax leucostictus* |
| Fishes | *Cyphocharax magdalenae* |
| Fishes | *Cyphocharax multilineatus* |
| Fishes | *Cyphocharax nigripinnis* |
| Fishes | *Cyphocharax notatus* |
| Fishes | *Cyphocharax oenas* |
| Fishes | *Cyphocharax pantostictos* |
| Fishes | *Cyphocharax spiluropsis* |
| Fishes | *Cyphocharax spilurus* |
| Fishes | *Potamorhina altamazonica* |
| Fishes | *Potamorhina laticeps* |
| Fishes | *Potamorhina latior* |
| Fishes | *Potamorhina pristigaster* |
| Fishes | *Pseudocurimata lineopunctata* |
| Fishes | *Pseudocurimata patiae* |
| Fishes | *Rhytiodus argenteofuscus* |
| Fishes | *Rhytiodus microlepis* |
| Fishes | *Hydrolycus armatus* |
| Fishes | *Hydrolycus scomberoides* |
| Fishes | *Hydrolycus tatauaia* |
| Fishes | *Hydrolycus wallacei* |
| Fishes | *Rhaphiodon vulpinus* |
| Fishes | *Chasmocranus rosae* |
| Fishes | *Hoplocharax goethei* |
| Fishes | *Erythrinus erythrinus* |
| Fishes | *Hoplerythrinus unitaeniatus* |
| Fishes | *Hoplias curupira* |
| Fishes | *Hoplias malabaricus* |
| Fishes | *Carnegiella marthae* |
| Fishes | *Carnegiella myersi* |
| Fishes | *Carnegiella schereri* |
| Fishes | *Carnegiella strigata* |
| Fishes | *Gasteropelecus maculatus* |
| Fishes | *Gasteropelecus sternicla* |
| Fishes | *Stegophilus septentrionalis* |
| Fishes | *Iguanodectes adujai* |
| Fishes | *Iguanodectes geisleri* |
| Fishes | *Iguanodectes purusii* |
| Fishes | *Iguanodectes spilurus* |
| Fishes | *Belonion dibranchodon* |
| Fishes | *Bivibranchia fowleri* |
| Fishes | *Copeina guttata* |
| Fishes | *Copella eigenmanni* |
| Fishes | *Copella nattereri* |
| Fishes | *Copella vilmae* |
| Fishes | *Nannostomus digrammus* |
| Fishes | *Nannostomus eques* |
| Fishes | *Nannostomus harrisoni* |
| Fishes | *Nannostomus marginatus* |
| Fishes | *Nannostomus marilynae* |
| Fishes | *Nannostomus trifasciatus* |
| Fishes | *Nannostomus unifasciatus* |
| Fishes | *Pyrrhulina brevis* |
| Fishes | *Pyrrhulina eleanorae* |
| Fishes | *Pyrrhulina filamentosa* |
| Fishes | *Pyrrhulina laeta* |
| Fishes | *Pyrrhulina lugubris* |
| Fishes | *Pyrrhulina obermulleri* |
| Fishes | *Pyrrhulina semifasciata* |
| Fishes | *Pyrrhulina zigzag* |
| Fishes | *Hoplomyzon papillatus* |
| Fishes | *Hoplomyzon sexpapilostoma* |
| Fishes | *Parodon alfonsoi* |
| Fishes | *Parodon apolinari* |
| Fishes | *Parodon atratoensis* |
| Fishes | *Parodon buckleyi* |
| Fishes | *Parodon caliensis* |
| Fishes | *Parodon magdalenensis* |
| Fishes | *Parodon pongoensis* |
| Fishes | *Parodon suborbitalis* |
| Fishes | *Prochilodus magdalenae* |
| Fishes | *Prochilodus mariae* |
| Fishes | *Prochilodus nigricans* |
| Fishes | *Prochilodus reticulatus* |
| Fishes | *Prochilodus rubrotaeniatus* |
| Fishes | *Semaprochilodus insignis* |
| Fishes | *Semaprochilodus kneri* |
| Fishes | *Semaprochilodus laticeps* |
| Fishes | *Semaprochilodus taeniurus* |
| Fishes | *Catoprion mento* |
| Fishes | *Colossoma macropomum* |
| Fishes | *Metynnis argenteus* |
| Fishes | *Metynnis hypsauchen* |
| Fishes | *Metynnis lippincottianus* |
| Fishes | *Metynnis luna* |
| Fishes | *Myleus setiger* |
| Fishes | *Myloplus asterias* |
| Fishes | *Myloplus rubripinnis* |
| Fishes | *Myloplus schomburgkii* |
| Fishes | *Myloplus torquatus* |
| Fishes | *Mylossoma acanthogaster* |
| Fishes | *Mylossoma albiscopum* |
| Fishes | *Mylossoma aureum* |
| Fishes | *Piaractus brachypomus* |
| Fishes | *Piaractus orinoquensis* |
| Fishes | *Pygocentrus cariba* |
| Fishes | *Pygocentrus nattereri* |
| Fishes | *Pygopristis denticulata* |
| Fishes | *Vandellia beccarii* |
| Fishes | *Vandellia cirrhosa* |
| Fishes | *Acarichthys heckelii* |
| Fishes | *Aequidens diadema* |
| Fishes | *Aequidens metae* |
| Fishes | *Aequidens tetramerus* |
| Fishes | *Andinoacara biseriatus* |
| Fishes | *Andinoacara latifrons* |
| Fishes | *Andinoacara sapayensis* |
| Fishes | *Apistogramma agassizii* |
| Fishes | *Apistogramma alacrina* |
| Fishes | *Apistogramma bitaeniata* |
| Fishes | *Apistogramma cacatuoides* |
| Fishes | *Apistogramma cruzi* |
| Fishes | *Apistogramma diplotaenia* |
| Fishes | *Apistogramma eunotus* |
| Fishes | *Apistogramma flabellicauda* |
| Fishes | *Apistogramma hoignei* |
| Fishes | *Apistogramma hongsloi* |
| Fishes | *Apistogramma iniridae* |
| Fishes | *Apistogramma inornata* |
| Fishes | *Apistogramma lineata* |
| Fishes | *Apistogramma macmasteri* |
| Fishes | *Apistogramma megaptera* |
| Fishes | *Apistogramma minima* |
| Fishes | *Apistogramma personata* |
| Fishes | *Apistogramma piaroa* |
| Fishes | *Apistogramma velifera* |
| Fishes | *Apistogramma viejita* |
| Fishes | *Apistogrammoides pucallpaensis* |
| Fishes | *Biotodoma cupido* |
| Fishes | *Biotodoma wavrini* |
| Fishes | *Biotoecus dicentrarchus* |
| Fishes | *Bujurquina cordemadi* |
| Fishes | *Bujurquina huallagae* |
| Fishes | *Bujurquina mariae* |
| Fishes | *Bujurquina moriorum* |
| Fishes | *Bujurquina peregrinabunda* |
| Fishes | *Bujurquina syspilus* |
| Fishes | *Caquetaia kraussii* |
| Fishes | *Caquetaia myersi* |
| Fishes | *Lonchogenys ilisha* |
| Fishes | *Anchoviella guianensis* |
| Fishes | *Anchoviella jamesi* |
| Fishes | *Jurengraulis juruensis* |
| Fishes | *Lycengraulis batesii* |
| Fishes | *Ichthyoelephas longirostris* |
| Fishes | *Ilisha amazonica* |
| Fishes | *Pellona castelnaeana* |
| Fishes | *Pellona flavipinnis* |
| Fishes | *Pristigaster cayana* |
| Fishes | *Pterengraulis atherinoides* |
| Fishes | *Leporellus vittatus* |
| Fishes | *Synaptolaemus latofasciatus* |
| Fishes | *Gambusia lemaitrei* |
| Fishes | *Grundulus bogotensis* |
| Fishes | *Grundulus cochae* |
| Fishes | *Neoheterandria elegans* |
| Fishes | *Poecilia caucana* |
| Fishes | *Poecilia gillii* |
| Fishes | *Poecilia koperi* |
| Fishes | *Poecilia wandae* |
| Fishes | *Poeciliopsis turrubarensis* |
| Fishes | *Poecilocharax weitzmani* |
| Fishes | *Pseudopoecilia austrocolumbiana* |
| Fishes | *Anablepsoides atratus* |
| Fishes | *Anablepsoides elongatus* |
| Fishes | *Anablepsoides limoncochae* |
| Fishes | *Anablepsoides ophiomimus* |
| Fishes | *Anablepsoides ornatus* |
| Fishes | *Anablepsoides rubrolineatus* |
| Fishes | *Anablepsoides taeniatus* |
| Fishes | *Anablepsoides tessellatus* |
| Fishes | *Cynodonichthys boehlkei* |
| Fishes | *Cynodonichthys elegans* |
| Fishes | *Cynodonichthys leucurus* |
| Fishes | *Cynodonichthys magdalenae* |
| Fishes | *Cynodonichthys pacificus* |
| Fishes | *Fluviphylax obscurus* |
| Fishes | *Fluviphylax pygmaeus* |
| Fishes | *Laimosemion carolinae* |
| Fishes | *Laimosemion flammaecauda* |
| Fishes | *Laimosemion leticia* |
| Fishes | *Laimosemion rectocaudatus* |
| Fishes | *Priapichthys caliensis* |
| Fishes | *Priapichthys nigroventralis* |
| Fishes | *Rachovia brevis* |
| Fishes | *Rachovia hummelincki* |
| Fishes | *Rachovia maculipinnis* |
| Fishes | *Adontosternarchus balaenops* |
| Fishes | *Adontosternarchus clarkae* |
| Fishes | *Adontosternarchus devenanzii* |
| Fishes | *Adontosternarchus sachsi* |
| Fishes | *Apteronotus albifrons* |
| Fishes | *Apteronotus anu* |
| Fishes | *Apteronotus apurensis* |
| Fishes | *Apteronotus bonapartii* |
| Fishes | *Apteronotus cuchillejo* |
| Fishes | *Apteronotus cuchillo* |
| Fishes | *Apteronotus eschmeyeri* |
| Fishes | *Apteronotus galvisi* |
| Fishes | *Apteronotus macrolepis* |
| Fishes | *Apteronotus macrostomus* |
| Fishes | *Apteronotus magdalenensis* |
| Fishes | *Apteronotus magoi* |
| Fishes | *Apteronotus mariae* |
| Fishes | *Apteronotus milesi* |
| Fishes | *Apteronotus rostratus* |
| Fishes | *Apteronotus spurrellii* |
| Fishes | *Microsternarchus bilineatus* |
| Fishes | *Renova oscari* |
| Fishes | *Steatogenys duidae* |
| Fishes | *Steatogenys elegans* |
| Fishes | *Steatogenys ocellatus* |
| Fishes | *Sternarchella orthos* |
| Fishes | *Sternarchella schotti* |
| Fishes | *Sternarchogiton nattereri* |
| Fishes | *Sternarchogiton porcinum* |
| Fishes | *Sternarchorhamphus muelleri* |
| Fishes | *Sternarchorhynchus mormyrus* |
| Fishes | *Sternarchorhynchus oxyrhynchus* |
| Fishes | *Sternarchorhynchus roseni* |
| Fishes | *Electrophorus electricus* |
| Fishes | *Gymnotus anguillaris* |
| Fishes | *Gymnotus ardilai* |
| Fishes | *Gymnotus carapo* |
| Fishes | *Gymnotus cataniapo* |
| Fishes | *Gymnotus choco* |
| Fishes | *Gymnotus coropinae* |
| Fishes | *Gymnotus henni* |
| Fishes | *Gymnotus javari* |
| Fishes | *Gymnotus pedanopterus* |
| Fishes | *Gymnotus stenoleucus* |
| Fishes | *Gymnotus tigre* |
| Fishes | *Gymnotus tiquie* |
| Fishes | *Brachyhypopomus batesi* |
| Fishes | *Brachyhypopomus beebei* |
| Fishes | *Brachyhypopomus bennetti* |
| Fishes | *Brachyhypopomus brevirostris* |
| Fishes | *Brachyhypopomus bullocki* |
| Fishes | *Brachyhypopomus flavipomus* |
| Fishes | *Brachyhypopomus occidentalis* |
| Fishes | *Brachyhypopomus sullivani* |
| Fishes | *Gymnorhamphichthys bogardusae* |
| Fishes | *Gymnorhamphichthys hypostomus* |
| Fishes | *Gymnorhamphichthys rondoni* |
| Fishes | *Rhamphichthys apurensis* |
| Fishes | *Rhamphichthys drepanium* |
| Fishes | *Rhamphichthys marmoratus* |
| Fishes | *Rhamphichthys rostratus* |
| Fishes | *Eigenmannia camposi* |
| Fishes | *Eigenmannia humboldtii* |
| Fishes | *Eigenmannia limbata* |
| Fishes | *Eigenmannia macrops* |
| Fishes | *Eigenmannia magoi* |
| Fishes | *Eigenmannia nigra* |
| Fishes | *Eigenmannia zenuensis* |
| Fishes | *Platyurosternarchus macrostoma* |
| Fishes | *Sternopygus aequilabiatus* |
| Fishes | *Sternopygus astrabes* |
| Fishes | *Sternopygus dariensis* |
| Fishes | *Sternopygus macrurus* |
| Fishes | *Sternopygus pejeraton* |
| Fishes | *Lepidosiren paradoxa* |
| Fishes | *Heliotrygon gomesi* |
| Fishes | *Paratrygon aiereba* |
| Fishes | *Plesiotrygon iwamae* |
| Fishes | *Plesiotrygon nana* |
| Fishes | *Potamotrygon constellata* |
| Fishes | *Potamotrygon magdalenae* |
| Fishes | *Potamotrygon motoro* |
| Fishes | *Potamotrygon orbignyi* |
| Fishes | *Potamotrygon schroederi* |
| Fishes | *Potamotrygon scobina* |
| Fishes | *Arapaima gigas* |
| Fishes | *Osteoglossum bicirrhosum* |
| Fishes | *Osteoglossum ferreirai* |
| Fishes | *Astronotus ocellatus* |
| Fishes | *Cichla intermedia* |
| Fishes | *Cichla monoculus* |
| Fishes | *Cichla orinocensis* |
| Fishes | *Cichla temensis* |
| Fishes | *Cichlasoma amazonarum* |
| Fishes | *Cichlasoma orinocense* |
| Fishes | *Crenicara punctulatum* |
| Fishes | *Crenicichla alta* |
| Fishes | *Crenicichla anthurus* |
| Fishes | *Crenicichla cincta* |
| Fishes | *Crenicichla geayi* |
| Fishes | *Crenicichla johanna* |
| Fishes | *Crenicichla lenticulata* |
| Fishes | *Crenicichla lugubris* |
| Fishes | *Crenicichla marmorata* |
| Fishes | *Crenicichla proteus* |
| Fishes | *Crenicichla reticulata* |
| Fishes | *Crenicichla strigata* |
| Fishes | *Crenicichla sveni* |
| Fishes | *Crenicichla zebrina* |
| Fishes | *Dicrossus filamentosus* |
| Fishes | *Dicrossus gladicauda* |
| Fishes | *Geophagus abalios* |
| Fishes | *Geophagus crassilabris* |
| Fishes | *Geophagus dicrozoster* |
| Fishes | *Geophagus megasema* |
| Fishes | *Geophagus pellegrini* |
| Fishes | *Geophagus steindachneri* |
| Fishes | *Geophagus surinamensis* |
| Fishes | *Geophagus taeniopareius* |
| Fishes | *Geophagus winemilleri* |
| Fishes | *Heroina isonycterina* |
| Fishes | *Heros efasciatus* |
| Fishes | *Heros liberifer* |
| Fishes | *Heros severus* |
| Fishes | *Hoplarchus psittacus* |
| Fishes | *Hypselecara coryphaenoides* |
| Fishes | *Hypselecara temporalis* |
| Fishes | *Kronoheros umbrifer* |
| Fishes | *Laetacara flavilabris* |
| Fishes | *Laetacara fulvipinnis* |
| Fishes | *Laetacara thayeri* |
| Fishes | *Mesoheros atromaculatus* |
| Fishes | *Mesoheros ornatus* |
| Fishes | *Mesonauta egregius* |
| Fishes | *Mesonauta insignis* |
| Fishes | *Mesonauta mirificus* |
| Fishes | *Mikrogeophagus ramirezi* |
| Fishes | *Pterophyllum altum* |
| Fishes | *Pterophyllum scalare* |
| Fishes | *Satanoperca acuticeps* |
| Fishes | *Satanoperca daemon* |
| Fishes | *Satanoperca jurupari* |
| Fishes | *Satanoperca mapiritensis* |
| Fishes | *Symphysodon aequifasciatus* |
| Fishes | *Uaru amphiacanthoides* |
| Fishes | *Uaru fernandezyepezi* |
| Fishes | *Monocirrhus polyacanthus* |
| Fishes | *Plagioscion squamosissimus* |
| Fishes | *Achirus novoae* |
| Fishes | *Trinectes hubbsbollinger* |
| Fishes | *Notarius bonillai* |
| Fishes | *Bunocephalus aleuropsis* |
| Fishes | *Bunocephalus aloikae* |
| Fishes | *Bunocephalus colombianus* |
| Fishes | *Bunocephalus coracoideus* |
| Fishes | *Bunocephalus knerii* |
| Fishes | *Bunocephalus verrucosus* |
| Fishes | *Helogenes castaneus* |
| Fishes | *Helogenes marmoratus* |
| Fishes | *Pseudobunocephalus amazonicus* |
| Fishes | *Pseudobunocephalus bifidus* |
| Fishes | *Pseudobunocephalus lundbergi* |
| Fishes | *Pseudostegophilus haemomyzon* |
| Fishes | *Pseudostegophilus nemurus* |
| Fishes | *Pterobunocephalus depressus* |
| Fishes | *Tetranematichthys wallacei* |
| Fishes | *Astroblepus acostai* |
| Fishes | *Astroblepus ardiladuartei* |
| Fishes | *Astroblepus ardilai* |
| Fishes | *Astroblepus bellezaensis* |
| Fishes | *Astroblepus cacharas* |
| Fishes | *Astroblepus caquetae* |
| Fishes | *Astroblepus chapmani* |
| Fishes | *Astroblepus chotae* |
| Fishes | *Astroblepus curitiensis* |
| Fishes | *Astroblepus floridablancaensis* |
| Fishes | *Astroblepus frenatus* |
| Fishes | *Astroblepus grixalvii* |
| Fishes | *Astroblepus guentheri* |
| Fishes | *Astroblepus heterodon* |
| Fishes | *Astroblepus homodon* |
| Fishes | *Astroblepus itae* |
| Fishes | *Astroblepus jimenezae* |
| Fishes | *Astroblepus jurubidae* |
| Fishes | *Astroblepus latidens* |
| Fishes | *Astroblepus mariae* |
| Fishes | *Astroblepus marmoratus* |
| Fishes | *Astroblepus martinezi* |
| Fishes | *Astroblepus micrescens* |
| Fishes | *Astroblepus mojicai* |
| Fishes | *Astroblepus nettoferreirai* |
| Fishes | *Astroblepus nicefori* |
| Fishes | *Astroblepus onzagaensis* |
| Fishes | *Astroblepus orientalis* |
| Fishes | *Astroblepus pradai* |
| Fishes | *Astroblepus putumayoensis* |
| Fishes | *Astroblepus retropinnus* |
| Fishes | *Astroblepus rivasae* |
| Fishes | *Astroblepus santanderensis* |
| Fishes | *Astroblepus trifasciatus* |
| Fishes | *Astroblepus ventralis* |
| Fishes | *Astroblepus verai* |
| Fishes | *Ageneiosus dentatus* |
| Fishes | *Ageneiosus inermis* |
| Fishes | *Ageneiosus lineatus* |
| Fishes | *Ageneiosus magoi* |
| Fishes | *Ageneiosus pardalis* |
| Fishes | *Ageneiosus polystictus* |
| Fishes | *Ageneiosus ucayalensis* |
| Fishes | *Ageneiosus vittatus* |
| Fishes | *Asterophysus batrachus* |
| Fishes | *Auchenipterichthys coracoideus* |
| Fishes | *Auchenipterichthys longimanus* |
| Fishes | *Auchenipterichthys punctatus* |
| Fishes | *Auchenipterus ambyiacus* |
| Fishes | *Auchenipterus brachyurus* |
| Fishes | *Auchenipterus britskii* |
| Fishes | *Auchenipterus nuchalis* |
| Fishes | *Centromochlus existimatus* |
| Fishes | *Centromochlus heckelii* |
| Fishes | *Centromochlus macracanthus* |
| Fishes | *Ceratobranchia joanae* |
| Fishes | *Chaetobranchus flavescens* |
| Fishes | *Gelanoglanis stroudi* |
| Fishes | *Gladioglanis conquistador* |
| Fishes | *Gladioglanis machadoi* |
| Fishes | *Markiana geayi* |
| Fishes | *Tatia altae* |
| Fishes | *Tatia aulopygia* |
| Fishes | *Tatia brunnea* |
| Fishes | *Tatia caudosignata* |
| Fishes | *Tatia dunni* |
| Fishes | *Tatia galaxias* |
| Fishes | *Tatia gyrina* |
| Fishes | *Tatia intermedia* |
| Fishes | *Tatia marthae* |
| Fishes | *Tatia nigra* |
| Fishes | *Tatia perugiae* |
| Fishes | *Tatia reticulata* |
| Fishes | *Tatia romani* |
| Fishes | *Tatia strigata* |
| Fishes | *Trachelyichthys decaradiatus* |
| Fishes | *Trachelyopterichthys anduzei* |
| Fishes | *Trachelyopterichthys taeniatus* |
| Fishes | *Trachelyopterus fisheri* |
| Fishes | *Trachelyopterus galeatus* |
| Fishes | *Trachelyopterus insignis* |
| Fishes | *Trachelyopterus peloichthys* |
| Fishes | *Trachycorystes trachycorystes* |
| Fishes | *Callichthys callichthys* |
| Fishes | *Callichthys fabricioi* |
| Fishes | *Callichthys oibaensis* |
| Fishes | *Callichthys serralabium* |
| Fishes | *Corydoras aeneus* |
| Fishes | *Corydoras agassizii* |
| Fishes | *Corydoras ambiacus* |
| Fishes | *Corydoras arcuatus* |
| Fishes | *Corydoras armatus* |
| Fishes | *Corydoras axelrodi* |
| Fishes | *Corydoras benattii* |
| Fishes | *Corydoras brevirostris* |
| Fishes | *Corydoras concolor* |
| Fishes | *Corydoras crypticus* |
| Fishes | *Corydoras delphax* |
| Fishes | *Corydoras elegans* |
| Fishes | *Corydoras esperanzae* |
| Fishes | *Corydoras evelynae* |
| Fishes | *Corydoras fowleri* |
| Fishes | *Corydoras gomezi* |
| Fishes | *Corydoras granti* |
| Fishes | *Corydoras habrosus* |
| Fishes | *Corydoras julii* |
| Fishes | *Corydoras leopardus* |
| Fishes | *Corydoras leucomelas* |
| Fishes | *Corydoras loxozonus* |
| Fishes | *Corydoras melanistius* |
| Fishes | *Corydoras melanotaenia* |
| Fishes | *Corydoras melini* |
| Fishes | *Corydoras metae* |
| Fishes | *Corydoras napoensis* |
| Fishes | *Corydoras osteocarus* |
| Fishes | *Corydoras pastazensis* |
| Fishes | *Corydoras pygmaeus* |
| Fishes | *Corydoras rabauti* |
| Fishes | *Corydoras reticulatus* |
| Fishes | *Corydoras reynoldsi* |
| Fishes | *Corydoras semiaquilus* |
| Fishes | *Corydoras septentrionalis* |
| Fishes | *Corydoras simulatus* |
| Fishes | *Corydoras sodalis* |
| Fishes | *Corydoras splendens* |
| Fishes | *Corydoras trilineatus* |
| Fishes | *Corydoras zygatus* |
| Fishes | *Hoplosternum littorale* |
| Fishes | *Hoplosternum magdalenae* |
| Fishes | *Hoplosternum punctatum* |
| Fishes | *Megalechis picta* |
| Fishes | *Megalechis thoracata* |
| Fishes | *Cetopsidium morenoi* |
| Fishes | *Cetopsidium pemon* |
| Fishes | *Cetopsis amphiloxa* |
| Fishes | *Cetopsis baudoensis* |
| Fishes | *Cetopsis candiru* |
| Fishes | *Cetopsis coecutiens* |
| Fishes | *Cetopsis fimbriata* |
| Fishes | *Cetopsis montana* |
| Fishes | *Cetopsis motatanensis* |
| Fishes | *Cetopsis orinoco* |
| Fishes | *Cetopsis othonops* |
| Fishes | *Cetopsis umbrosa* |
| Fishes | *Cetopsis varii* |
| Fishes | *Psectrogaster amazonica* |
| Fishes | *Psectrogaster ciliata* |
| Fishes | *Psectrogaster essequibensis* |
| Fishes | *Psectrogaster rhomboides* |
| Fishes | *Psectrogaster rutiloides* |
| Fishes | *Acanthodoras cataphractus* |
| Fishes | *Acanthodoras depressus* |
| Fishes | *Acanthodoras spinosissimus* |
| Fishes | *Agamyxis albomaculatus* |
| Fishes | *Agamyxis pectinifrons* |
| Fishes | *Amblydoras affinis* |
| Fishes | *Amblydoras gonzalezi* |
| Fishes | *Amblydoras monitor* |
| Fishes | *Amblydoras nauticus* |
| Fishes | *Anadoras grypus* |
| Fishes | *Anadoras regani* |
| Fishes | *Anduzedoras oxyrhynchus* |
| Fishes | *Argopleura chocoensis* |
| Fishes | *Argopleura conventus* |
| Fishes | *Argopleura diquensis* |
| Fishes | *Argopleura magdalenensis* |
| Fishes | *Centrochir crocodili* |
| Fishes | *Centrodoras brachiatus* |
| Fishes | *Centrodoras hasemani* |
| Fishes | *Doraops zuloagai* |
| Fishes | *Doras phlyzakion* |
| Fishes | *Hassar orestis* |
| Fishes | *Hemidoras boulengeri* |
| Fishes | *Hemidoras morrisi* |
| Fishes | *Hemidoras stenopeltis* |
| Fishes | *Hemidoras stuebelii* |
| Fishes | *Hypodoras forficulatus* |
| Fishes | *Leptodoras acipenserinus* |
| Fishes | *Leptodoras juruensis* |
| Fishes | *Leptodoras linnelli* |
| Fishes | *Leptodoras nelsoni* |
| Fishes | *Leptodoras rogersae* |
| Fishes | *Liosomadoras morrowi* |
| Fishes | *Liosomadoras oncinus* |
| Fishes | *Lithodoras dorsalis* |
| Fishes | *Megalodoras uranoscopus* |
| Fishes | *Nemadoras elongatus* |
| Fishes | *Nemadoras hemipeltis* |
| Fishes | *Nemadoras humeralis* |
| Fishes | *Orinocodoras eigenmanni* |
| Fishes | *Ossancora punctata* |
| Fishes | *Oxydoras niger* |
| Fishes | *Oxydoras sifontesi* |
| Fishes | *Oxyropsis acutirostra* |
| Fishes | *Oxyropsis carinata* |
| Fishes | *Oxyropsis wrightiana* |
| Fishes | *Pachypops fourcroi* |
| Fishes | *Pachypops trifilis* |
| Fishes | *Physopyxis ananas* |
| Fishes | *Physopyxis lyra* |
| Fishes | *Platydoras armatulus* |
| Fishes | *Platydoras hancockii* |
| Fishes | *Pterodoras granulosus* |
| Fishes | *Pterodoras rivasi* |
| Fishes | *Rhinodoras boehlkei* |
| Fishes | *Rhinodoras gallagheri* |
| Fishes | *Rhinodoras thomersoni* |
| Fishes | *Scorpiodoras bolivarensis* |
| Fishes | *Scorpiodoras heckelii* |
| Fishes | *Trachydoras microstomus* |
| Fishes | *Trachydoras nattereri* |
| Fishes | *Trachydoras steindachneri* |
| Fishes | *Distocyclus conirostris* |
| Fishes | *Brachyrhamdia imitator* |
| Fishes | *Brachyrhamdia meesi* |
| Fishes | *Brachyrhamdia thayeria* |
| Fishes | *Cetopsorhamdia boquillae* |
| Fishes | *Cetopsorhamdia molinae* |
| Fishes | *Cetopsorhamdia nasus* |
| Fishes | *Cetopsorhamdia orinoco* |
| Fishes | *Cetopsorhamdia picklei* |
| Fishes | *Cetopsorhamdia shermani* |
| Fishes | *Imparfinis microps* |
| Fishes | *Imparfinis nemacheir* |
| Fishes | *Imparfinis pristos* |
| Fishes | *Imparfinis pseudonemacheir* |
| Fishes | *Imparfinis spurrellii* |
| Fishes | *Imparfinis stictonotus* |
| Fishes | *Imparfinis timana* |
| Fishes | *Imparfinis usmai* |
| Fishes | *Mastiglanis asopos* |
| Fishes | *Megalonema orixanthum* |
| Fishes | *Megalonema platycephalum* |
| Fishes | *Megalonema psammium* |
| Fishes | *Megalonema xanthum* |
| Fishes | *Myoglanis koepckei* |
| Fishes | *Nemuroglanis mariai* |
| Fishes | *Nemuroglanis pauciradiatus* |
| Fishes | *Phenacorhamdia anisura* |
| Fishes | *Phenacorhamdia macarenensis* |
| Fishes | *Phenacorhamdia nigrolineata* |
| Fishes | *Phenacorhamdia provenzanoi* |
| Fishes | *Phenacorhamdia taphorni* |
| Fishes | *Pimelodella chagresi* |
| Fishes | *Pimelodella chaparae* |
| Fishes | *Pimelodella conquetaensis* |
| Fishes | *Pimelodella cristata* |
| Fishes | *Pimelodella cruxenti* |
| Fishes | *Pimelodella eutaenia* |
| Fishes | *Pimelodella figueroai* |
| Fishes | *Pimelodella floridablancaensis* |
| Fishes | *Pimelodella geryi* |
| Fishes | *Pimelodella gracilis* |
| Fishes | *Pimelodella grisea* |
| Fishes | *Pimelodella linami* |
| Fishes | *Pimelodella longibarbata* |
| Fishes | *Pimelodella macrocephala* |
| Fishes | *Pimelodella megalops* |
| Fishes | *Pimelodella metae* |
| Fishes | *Pimelodella modestus* |
| Fishes | *Pimelodella odynea* |
| Fishes | *Pimelodella reyesi* |
| Fishes | *Pimelodella serrata* |
| Fishes | *Potamorrhaphis guianensis* |
| Fishes | *Potamorrhaphis petersi* |
| Fishes | *Rhamdia guatemalensis* |
| Fishes | *Rhamdia laukidi* |
| Fishes | *Rhamdia muelleri* |
| Fishes | *Hypophthalmus celiae* |
| Fishes | *Hypophthalmus edentatus* |
| Fishes | *Hypophthalmus fimbriatus* |
| Fishes | *Hypophthalmus marginatus* |
| Fishes | *Hypophthalmus oremaculatus* |
| Fishes | *Acanthicus hystrix* |
| Fishes | *Aguarunichthys inpai* |
| Fishes | *Ancistrus caucanus* |
| Fishes | *Ancistrus centrolepis* |
| Fishes | *Ancistrus dolichopterus* |
| Fishes | *Ancistrus macrophthalmus* |
| Fishes | *Ancistrus malacops* |
| Fishes | *Ancistrus martini* |
| Fishes | *Ancistrus patronus* |
| Fishes | *Ancistrus tolima* |
| Fishes | *Ancistrus triradiatus* |
| Fishes | *Aphanotorulus ammophilus* |
| Fishes | *Aphanotorulus emarginatus* |
| Fishes | *Aphanotorulus horridus* |
| Fishes | *Aphanotorulus unicolor* |
| Fishes | *Baryancistrus beggini* |
| Fishes | *Baryancistrus demantoides* |
| Fishes | *Chaetostoma anale* |
| Fishes | *Chaetostoma anomalum* |
| Fishes | *Chaetostoma breve* |
| Fishes | *Chaetostoma brevilabiatum* |
| Fishes | *Chaetostoma chimu* |
| Fishes | *Chaetostoma dorsale* |
| Fishes | *Chaetostoma fischeri* |
| Fishes | *Chaetostoma formosae* |
| Fishes | *Chaetostoma joropo* |
| Fishes | *Chaetostoma lepturum* |
| Fishes | *Chaetostoma leucomelas* |
| Fishes | *Chaetostoma marginatum* |
| Fishes | *Chaetostoma milesi* |
| Fishes | *Chaetostoma niveum* |
| Fishes | *Chaetostoma palmeri* |
| Fishes | *Chaetostoma patiae* |
| Fishes | *Chaetostoma paucispinis* |
| Fishes | *Chaetostoma platyrhynchus* |
| Fishes | *Chaetostoma sovichthys* |
| Fishes | *Chaetostoma tachiraense* |
| Fishes | *Chaetostoma thomsoni* |
| Fishes | *Cordylancistrus daguae* |
| Fishes | *Cordylancistrus pijao* |
| Fishes | *Cordylancistrus tayrona* |
| Fishes | *Crossoloricaria cephalaspis* |
| Fishes | *Crossoloricaria variegata* |
| Fishes | *Crossoloricaria venezuelae* |
| Fishes | *Cruciglanis pacifici* |
| Fishes | *Daector quadrizonatus* |
| Fishes | *Dasyloricaria filamentosa* |
| Fishes | *Dasyloricaria latiura* |
| Fishes | *Dasyloricaria paucisquama* |
| Fishes | *Dekeyseria amazonica* |
| Fishes | *Dekeyseria picta* |
| Fishes | *Dolichancistrus atratoensis* |
| Fishes | *Dolichancistrus carnegiei* |
| Fishes | *Dolichancistrus cobrensis* |
| Fishes | *Dolichancistrus fuesslii* |
| Fishes | *Farlowella acus* |
| Fishes | *Farlowella amazonum* |
| Fishes | *Farlowella colombiensis* |
| Fishes | *Farlowella curtirostra* |
| Fishes | *Farlowella gracilis* |
| Fishes | *Farlowella mariaelenae* |
| Fishes | *Farlowella mitoupibo* |
| Fishes | *Farlowella nattereri* |
| Fishes | *Farlowella oxyrryncha* |
| Fishes | *Farlowella smithi* |
| Fishes | *Farlowella taphorni* |
| Fishes | *Farlowella vittata* |
| Fishes | *Hemiancistrus guahiborum* |
| Fishes | *Hemiancistrus subviridis* |
| Fishes | *Hypancistrus contradens* |
| Fishes | *Hypancistrus debilittera* |
| Fishes | *Hypancistrus furunculus* |
| Fishes | *Hypancistrus inspector* |
| Fishes | *Hypancistrus lunaorum* |
| Fishes | *Hypoclinemus mentalis* |
| Fishes | *Hypoptopoma bianale* |
| Fishes | *Hypoptopoma brevirostratum* |
| Fishes | *Hypoptopoma gulare* |
| Fishes | *Hypoptopoma machadoi* |
| Fishes | *Hypoptopoma steindachneri* |
| Fishes | *Hypoptopoma thoracatum* |
| Fishes | *Hypostomus annectens* |
| Fishes | *Hypostomus argus* |
| Fishes | *Hypostomus carinatus* |
| Fishes | *Hypostomus hemicochliodon* |
| Fishes | *Hypostomus holostictus* |
| Fishes | *Hypostomus hondae* |
| Fishes | *Hypostomus niceforoi* |
| Fishes | *Hypostomus oculeus* |
| Fishes | *Hypostomus plecostomoides* |
| Fishes | *Hypostomus plecostomus* |
| Fishes | *Hypostomus pyrineusi* |
| Fishes | *Hypostomus robinii* |
| Fishes | *Hypostomus sculpodon* |
| Fishes | *Hypostomus varimaculosus* |
| Fishes | *Hypostomus wilsoni* |
| Fishes | *Isorineloricaria tenuicauda* |
| Fishes | *Isorineloricaria villarsi* |
| Fishes | *Lamontichthys filamentosus* |
| Fishes | *Lamontichthys llanero* |
| Fishes | *Lamontichthys maracaibero* |
| Fishes | *Lasiancistrus caucanus* |
| Fishes | *Lasiancistrus guacharote* |
| Fishes | *Lasiancistrus schomburgkii* |
| Fishes | *Lasiancistrus tentaculatus* |
| Fishes | *Leporacanthicus galaxias* |
| Fishes | *Leporacanthicus triactis* |
| Fishes | *Lepthoplosternum altamazonicum* |
| Fishes | *Leptoancistrus canensis* |
| Fishes | *Leptoancistrus cordobensis* |
| Fishes | *Leptotocinclus ctenistus* |
| Fishes | *Limatulichthys griseus* |
| Fishes | *Loricaria cataphracta* |
| Fishes | *Loricaria nickeriensis* |
| Fishes | *Loricaria simillima* |
| Fishes | *Loricariichthys acutus* |
| Fishes | *Loricariichthys brunneus* |
| Fishes | *Loricariichthys hauxwelli* |
| Fishes | *Loricariichthys stuebelii* |
| Fishes | *Nannoptopoma spectabile* |
| Fishes | *Nannoptopoma sternoptychum* |
| Fishes | *Otocinclus batmani* |
| Fishes | *Otocinclus huaorani* |
| Fishes | *Otocinclus macrospilus* |
| Fishes | *Otocinclus vestitus* |
| Fishes | *Otocinclus vittatus* |
| Fishes | *Panaqolus albomaculatus* |
| Fishes | *Panaqolus maccus* |
| Fishes | *Panaque cochliodon* |
| Fishes | *Panaque nigrolineatus* |
| Fishes | *Panaque suttonorum* |
| Fishes | *Panaque titan* |
| Fishes | *Parapteronotus hasemani* |
| Fishes | *Parotocinclus eppleyi* |
| Fishes | *Parotocinclus longirostris* |
| Fishes | *Parotocinclus variola* |
| Fishes | *Peckoltia brevis* |
| Fishes | *Peckoltia caenosa* |
| Fishes | *Peckoltia lineola* |
| Fishes | *Peckoltia lujani* |
| Fishes | *Peckoltia sabaji* |
| Fishes | *Peckoltia vittata* |
| Fishes | *Peckoltichthys bachi* |
| Fishes | *Platynematichthys notatus* |
| Fishes | *Pseudancistrus sidereus* |
| Fishes | *Pseudohemiodon unillano* |
| Fishes | *Pseudolithoxus anthrax* |
| Fishes | *Pseudolithoxus dumus* |
| Fishes | *Pseudolithoxus kelsorum* |
| Fishes | *Pseudolithoxus tigris* |
| Fishes | *Pseudorinelepis genibarbis* |
| Fishes | *Pterygoplichthys gibbiceps* |
| Fishes | *Pterygoplichthys lituratus* |
| Fishes | *Pterygoplichthys multiradiatus* |
| Fishes | *Pterygoplichthys pardalis* |
| Fishes | *Pterygoplichthys undecimalis* |
| Fishes | *Pterygoplichthys weberi* |
| Fishes | *Pterygoplichthys zuliaensis* |
| Fishes | *Rhabdolichops caviceps* |
| Fishes | *Rhabdolichops eastwardi* |
| Fishes | *Rhabdolichops troscheli* |
| Fishes | *Rhabdolichops zareti* |
| Fishes | *Rhadinoloricaria rhami* |
| Fishes | *Rineloricaria castroi* |
| Fishes | *Rineloricaria daraha* |
| Fishes | *Rineloricaria eigenmanni* |
| Fishes | *Rineloricaria formosa* |
| Fishes | *Rineloricaria jubata* |
| Fishes | *Rineloricaria jurupari* |
| Fishes | *Rineloricaria lanceolata* |
| Fishes | *Rineloricaria magdalenae* |
| Fishes | *Rineloricaria phoxocephala* |
| Fishes | *Rineloricaria rupestris* |
| Fishes | *Spatuloricaria atratoensis* |
| Fishes | *Spatuloricaria caquetae* |
| Fishes | *Spatuloricaria curvispina* |
| Fishes | *Spatuloricaria euacanthagenys* |
| Fishes | *Spatuloricaria fimbriata* |
| Fishes | *Spatuloricaria gymnogaster* |
| Fishes | *Spatuloricaria lagoichthys* |
| Fishes | *Spatuloricaria phelpsi* |
| Fishes | *Spatuloricaria terracanticum* |
| Fishes | *Sturisoma guentheri* |
| Fishes | *Sturisoma tenuirostre* |
| Fishes | *Sturisomatichthys citurensis* |
| Fishes | *Sturisomatichthys leightoni* |
| Fishes | *Sturisomatichthys tamanae* |
| Fishes | *Brachyplatystoma filamentosum* |
| Fishes | *Brachyplatystoma juruense* |
| Fishes | *Brachyplatystoma platynemum* |
| Fishes | *Brachyplatystoma rousseauxii* |
| Fishes | *Brachyplatystoma tigrinum* |
| Fishes | *Brachyplatystoma vaillantii* |
| Fishes | *Calophysus macropterus* |
| Fishes | *Dianema longibarbis* |
| Fishes | *Hemisorubim platyrhynchos* |
| Fishes | *Leiarius marmoratus* |
| Fishes | *Leiarius pictus* |
| Fishes | *Pachyurus gabrielensis* |
| Fishes | *Pachyurus junki* |
| Fishes | *Pachyurus schomburgkii* |
| Fishes | *Phractocephalus hemioliopterus* |
| Fishes | *Pimelodina flavipinnis* |
| Fishes | *Pimelodus albofasciatus* |
| Fishes | *Pimelodus altissimus* |
| Fishes | *Pimelodus blochii* |
| Fishes | *Pimelodus coprophagus* |
| Fishes | *Pimelodus crypticus* |
| Fishes | *Pimelodus garciabarrigai* |
| Fishes | *Pimelodus grosskopfii* |
| Fishes | *Pimelodus navarroi* |
| Fishes | *Pimelodus ornatus* |
| Fishes | *Pimelodus pictus* |
| Fishes | *Pimelodus punctatus* |
| Fishes | *Pimelodus yuma* |
| Fishes | *Pinirampus pirinampu* |
| Fishes | *Platysilurus malarmo* |
| Fishes | *Platysilurus mucosus* |
| Fishes | *Platystomatichthys sturio* |
| Fishes | *Pseudepapterus cucuhyensis* |
| Fishes | *Pseudepapterus hasemani* |
| Fishes | *Pseudopimelodus atricaudus* |
| Fishes | *Pseudopimelodus bufonius* |
| Fishes | *Pseudopimelodus magnus* |
| Fishes | *Pseudopimelodus schultzi* |
| Fishes | *Pseudoplatystoma magdaleniatum* |
| Fishes | *Pseudoplatystoma metaense* |
| Fishes | *Pseudoplatystoma orinocoense* |
| Fishes | *Pseudoplatystoma punctifer* |
| Fishes | *Pseudoplatystoma tigrinum* |
| Fishes | *Sorubim cuspicaudus* |
| Fishes | *Sorubim elongatus* |
| Fishes | *Sorubim lima* |
| Fishes | *Sorubim maniradii* |
| Fishes | *Sorubimichthys planiceps* |
| Fishes | *Zungaro zungaro* |
| Fishes | *Cephalosilurus apurensis* |
| Fishes | *Megalocentor echthrus* |
| Fishes | *Microglanis iheringi* |
| Fishes | *Microglanis poecilus* |
| Fishes | *Microglanis secundus* |
| Fishes | *Scoloplax baileyi* |
| Fishes | *Scoloplax dicra* |
| Fishes | *Ammoglanis pulex* |
| Fishes | *Batrochoglanis acanthochiroides* |
| Fishes | *Batrochoglanis raninus* |
| Fishes | *Batrochoglanis transmontanus* |
| Fishes | *Batrochoglanis villosus* |
| Fishes | *Ituglanis guayaberensis* |
| Fishes | *Ituglanis metae* |
| Fishes | *Malacoglanis gelatinosus* |
| Fishes | *Ochmacanthus alternus* |
| Fishes | *Ochmacanthus orinoco* |
| Fishes | *Ochmacanthus reinhardtii* |
| Fishes | *Potamoglanis hasemani* |
| Fishes | *Rhyacoglanis annulatus* |
| Fishes | *Trichomycterus arhuaco* |
| Fishes | *Trichomycterus ballesterosi* |
| Fishes | *Trichomycterus banneaui* |
| Fishes | *Trichomycterus bogotensis* |
| Fishes | *Trichomycterus cachiraensis* |
| Fishes | *Trichomycterus calai* |
| Fishes | *Trichomycterus caliensis* |
| Fishes | *Trichomycterus chapmani* |
| Fishes | *Trichomycterus donascimientoi* |
| Fishes | *Trichomycterus dorsostriatum* |
| Fishes | *Trichomycterus emanueli* |
| Fishes | *Trichomycterus garciamarquezi* |
| Fishes | *Trichomycterus gorgona* |
| Fishes | *Trichomycterus kankuamo* |
| Fishes | *Trichomycterus knerii* |
| Fishes | *Trichomycterus latidens* |
| Fishes | *Trichomycterus latistriatus* |
| Fishes | *Trichomycterus maldonadoi* |
| Fishes | *Trichomycterus manaurensis* |
| Fishes | *Trichomycterus maracaiboensis* |
| Fishes | *Trichomycterus migrans* |
| Fishes | *Trichomycterus mogotensis* |
| Fishes | *Trichomycterus nietoi* |
| Fishes | *Trichomycterus nigromaculatus* |
| Fishes | *Trichomycterus ocanaensis* |
| Fishes | *Trichomycterus regani* |
| Fishes | *Trichomycterus retropinnis* |
| Fishes | *Trichomycterus romeroi* |
| Fishes | *Trichomycterus rosablanca* |
| Fishes | *Trichomycterus ruitoquensis* |
| Fishes | *Trichomycterus sandovali* |
| Fishes | *Trichomycterus sketi* |
| Fishes | *Trichomycterus spectrum* |
| Fishes | *Trichomycterus spilosoma* |
| Fishes | *Trichomycterus steindachneri* |
| Fishes | *Trichomycterus stellatus* |
| Fishes | *Trichomycterus straminius* |
| Fishes | *Trichomycterus taenia* |
| Fishes | *Trichomycterus tetuanensis* |
| Fishes | *Trichomycterus torcoromaensis* |
| Fishes | *Trichomycterus transandianus* |
| Fishes | *Trichomycterus uisae* |
| Fishes | *Synbranchus marmoratus* |
| Fishes | *Rhizosomichthys totae* |
| Fishes | *Colomesus asellus* |
| Ecosystem | Agroecosystem |
| Ecosystem | Shrub |
| Ecosystem | Forest |
| Ecosystem | Fragmented forest |
| Ecosystem | Rocky complex |
| Ecosystem | Coral reef |
| Ecosystem | Artificial water body |
| Ecosystem | Desert |
| Ecosystem | Soft water bottoms |
| Ecosystem | Hard water bottoms, not coral reefs |
| Ecosystem | Glaciers and snowfields |
| Ecosystem | Grassland |
| Ecosystem | Lake |
| Ecosystem | Lagoon |
| Ecosystem | Tidal pool |
| Ecosystem | Other areas |
| Ecosystem | Paramo |
| Ecosystem | Beaches |
| Ecosystem | Seagrass meadow |
| Ecosystem | River |
| Ecosystem | Savannah |
| Ecosystem | No information |
| Ecosystem | Subxerophytia |
| Ecosystem | Artificial territory |
| Ecosystem | Transformed transition |
| Ecosystem | Peatland |
| Ecosystem | Secondary vegetation |
| Ecosystem | Xerophytia |
| Ecosystem | Swampy area |
| Ecosystem | Natural sandy areas |
